# Evolution of the regulation of developmental gene expression in blind Mexican cavefish

**DOI:** 10.1101/2022.07.12.499770

**Authors:** Julien Leclercq, Jorge Torres-Paz, Maxime Policarpo, François Agnès, Sylvie Rétaux

## Abstract

Changes in gene expression regulation during development are considered the main drivers of morphological evolution and diversification. Here, we analysed the embryonic transcriptomes of surface-dwelling and blind cave-adapted morphs of the species *Astyanax mexicanus* and their reciprocal F1 hybrids at tailbud stage. Comparing gene expression in parents and allelic expression ratios in hybrids, we found that ∼20% of the transcriptome is differentially expressed and that *cis*-regulatory changes are the main contributors to variations in early developmental gene expression in the two morphs. We provide a list of 108 *cis*-regulated genes that could contribute to cavefish developmental evolution, and further explore the regulatory mechanisms controlling the cellular and regional expression of *rx3*, a “master eye gene”. Using quantitative embryology approaches after fluorescent *in situ* hybridisation, cell transplantations and interference with signalling pathways, we show that *rx3* cellular levels -controlling optic cell fates-are regulated in *cis* and in a cell-autonomous manner, whereas the size of *rx3* domain -controlling eye size-depends on non-autonomous Wnt signalling. Altogether, we reveal how distinct mechanisms and regulatory modules can regulate developmental gene expression and shape developmental evolution, with negligible contribution of coding mutations.

## INTRODUCTION

Changes in the spatial and temporal regulation of gene expression are considered the main drivers of morphological evolution and diversification [1]. As developmental gene regulatory networks deploy and progressively establish embryonic axes and germ layers, control regional and cell-type specification and drive terminal differentiation, gene expression is tightly regulated in time, in space and in intensity. Thus, any viable change in non-coding *cis-* regulatory DNA elements and/or *trans*-acting factors binding them can affect gene expression patterns and in turn lead to developmental variations. As opposed to such regulatory evolution, modification in protein-coding sequences supposedly have a greater functional impact, especially for developmental genes with pleiotropic functions, and are thought to be negatively selected [1, 2].

Teleost fish are the most diversified vertebrate group with more than 30,000 species adapted to various environments and displaying a wide range of morphologies [3]. However little is known on the genome-wide regulatory evolution accompanying their diversification [4, 5]. The Mexican tetra, *Astyanax mexicanus*, has emerged as a model to study microevolution and adaptation to a drastic environmental change. This species comes in two eco-morphotypes: the sighted, river-dwelling surface fish (SF) and the blind dark-adapted cavefish (CF) [6, 7]. Cavefishes present a variety of anatomical, behavioural and physiological phenotypes whose genetic bases are mostly unknown [8], yet evidences point towards multiple changes in gene expression profiles during early development that could explain those traits, e.g. [9-14]. As the divergence between the two morphs is recent [15-17], they are inter-fertile and crosses can generate viable F1 hybrids. We used this property to explore the mechanisms underlying the evolution of developmental gene regulation in cavefish, first by transcriptome-wide analysis of allelic expression ratios in F1 hybrids, then by embryological manipulations including cell transplantations between the two morphs.

## RESULTS

### *Cis*- versus *trans*-regulatory changes in the evolution of developmental gene expression in cavefish

We generated transcriptomes of Pachón CF, Texas SF as well as reciprocal F1 hybrid embryos at neural plate stage (tailbud/10hours post-fertilization). This stage corresponds to the end of gastrulation, a time when extensive variations in gene expression occur to set out neurulation and brain/eye development [9, 13, 18, 19]. Thus, 4483 transcripts were differentially expressed between SF and CF (DEG; Fold change>1.5; FDR<1%), representing 19% of the 10hpf *A. mexicanus* transcriptome (**Figure 1AB)**. This result, comparable to previous transcriptomic analyses at various stages [12, 13], illustrates the extensive regulatory variations that occur between SF and CF during precocious development.

**Figure 1:**
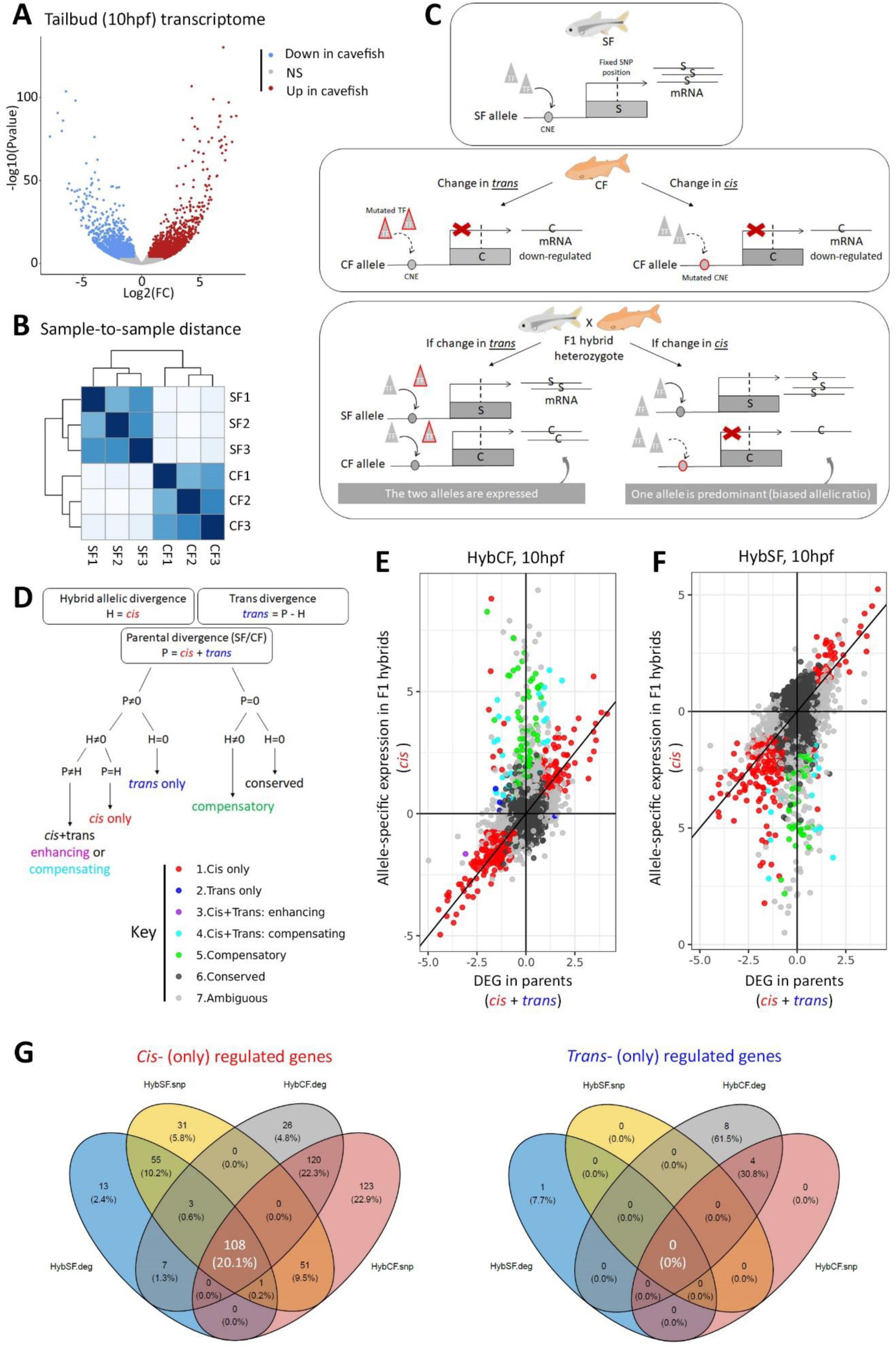
the evolution of cavefish tailbud transcriptome is mainly regulated in *cis*. **A**, Volcano plot showing differentially-expressed genes (DEG) between the two *Astyanax* morphs at tailbud stage. Red dots, upregulated in cavefish; blue dots down-regulated in cavefish; grey dots, non DEG. **B**, Sample to sample distance. **C**, Principle of the F1 hybrids analyses to uncover *cis* from *trans* gene regulation. The example shows a gene that is down-regulated in cavefish, either because of a defective transcription factor (TF, triangles) that cannot bind a conserved non-coding enhancer element (CNE) in *trans*, or because of a defective CNE in *cis*. In heterozygote F1 hybrids, the presence of fixed SNP position(s) allows recognizing the origin of expressed alleles and calculating allelic expression ratios. **D**, Principle of attribution to the different regulatory categories. Adapted and modified from [20]. **EF**, Scatter plots of *cis* regulatory divergence (y axis; from allelic expression ratios in hybrids) versus parental divergence (x-axis; from DEG in parents), where the different categories of genes are color-coded. E is from HybSF analysis and F is from HybSF analysis (computed from reads exclusively encompassing the variant position). **G**, Venn diagrams showing the consistence of results obtained from HybCF or HybSF datasets and with the two calculation methods (at the variant position/.snp, or on the whole transcript/.deg).

To disentangle *cis-* from *trans-* regulatory divergences in cavefish developmental evolution, we then searched for transcripts with fixed polymorphism(s) in parents and analysed their relative allelic expression in F1 hybrids. The proportion of parental alleles should be biased in hybrids in case a *cis*-regulatory change occurred in one morph, but not if the change occurred in *trans* (**Figure 1C**). Thereafter “surface hybrids” (HybSF) and “cave hybrids” (HybCF) will designate individuals resulting from a cross between a surface female and a cave male, and reciprocally.

Mapped reads were searched for variants, and only variants that were differently fixed in all parental replicates (3 SF *versus* 3 CF samples) were considered. We identified 26,390 fixed variants, contained in 5,573 genes, among which 900 were DEGs (=16%). The similar proportion of DEGs observed transcriptome-wide or among the subset of genes with fixed polymorphisms suggested that the 5573 genes set can provide a representative glimpse to the evolution of gene expression regulation in cavefish embryos. Many genes carried several associated variants (mean=4.7; median=4) and consistently for RNAseq data, the majority of fixed variants were annotated in exons and 3’-UTR (**Figure S1AB**).

*Cis*-regulation was then detected as a variation in allelic expression in F1 hybrids (i.e., the ratio of cave allele *vs* surface allele expression in <u>**H**</u>ybrids, termed **H**) (**Figure 1D**). *Trans*-regulation was detected by the difference between the variation in gene expression in <u>**P**</u>arents (i.e., cave/surface expression ratio at the SNP position, termed **P**) and *cis*-regulation. Therefore, ***trans*=P-H**. The regulatory category was attributed based on the statistical result of each 3 parameters, as described previously [20-22] and using a FDR threshold of 1%. For a given DEG in the parents, the regulatory change can be either in *cis*-(category 1), or in *trans*-(cat.2), or in both. In that case the effect can be enhancing (*cis* x *trans*, cat.3) or compensating (*cis* + *trans*, cat.4). No difference in parental expression with *cis*- and *trans*-regulatory differences in hybrids suggests a compensatory effect (cat.5). When no differences are detected in both parents and hybrids, regulation is conserved (cat.6). Any other combination was classified as ambiguous (cat.7). The results were similar whether parental expression counts were computed from reads exclusively encompassing the variant position, or from reads encompassing the entire transcripts (=DEG results) (compare **Figure 1EFG** and **Figure S1CD**). The majority of the 5573 genes analysed belonged to cat.6 in both HybSF and HybCF datasets (**Figure 1EF**, black dots). Allele-specific expression in hybrids revealed that 167 genes in HybSF and 228 genes in HybCF belonged to cat.1 (*cis*-regulatory divergence), respectively (**Figure 1EF**, red dots). Among these, 108 were shared (=2% of the 5573 genes studied; 547 variants on those 108 genes; **Figure 1G**). We considered these 108 genes as strong *cis*-regulatory divergent candidate genes between Pachón cave and surface *Astyanax*, and of major interest for further analyses (list in **Table S1**; full raw dataset in **Table S2**). Very few genes were found in cat.2 and cat.3 (0 gene) or cat.4 (*cis* + *trans*; 6 and 12 genes in reciprocal F1 hybrids, including 5 shared), suggesting that *trans*-regulatory evolution has limited impact on the evolution of cavefish development at tailbud stage (**Figure 1G)**. We concluded that *cis*-regulatory changes are the main drivers of variations in early developmental gene expression in the two *Astyanax* morphs at tailbud stage. This is consistent with the idea that changes in *trans* could have rather large pleiotropic effects whereas changes in *cis* fine-tunes gene expression.

### *Cis*-regulated genes in cavefish development

Gene Ontology (GO) analysis of the 108 *cis*-regulated genes revealed little or no over-representation of specific molecular function and biological process GO term (**Figure S2A**). Likewise, network analysis using String did not show a significant number of connections or clusters among the 108 genes (**Figure S2B**), suggesting that *cis*-regulatory changes affect a variety of seemingly unrelated developmental processes. Of note, 16 genes were located within less than 500kb of a previously described QTL, including seven close to an eye-related QTL (**Table S1**).

Some *cis*-regulated genes stood of particular interest with respect to the literature on the evolution of cavefish eye development and degeneration (**Table S1**). This included *Otx2*, up-regulated in CF tailbud, whereas it is downregulated at later stages and suspected as an important regulator of cave-associated phenotypes [12, 23]. The axial mesoderm expressed *admp* (antidorsalizing morphogenetic protein) and the Wnt pathway regulator *gsk3aa* (glycogene synthase kinase) were down-regulated in cavefish, potentially contributing to signalling variations affecting head development [13, 18, 24, 25]. The DNA-methyl transferase *dnmt3bb*.*3* was also under-expressed in cavefish and might contribute to early epigenetic regulation of eye development, as suggested for *dnmt3bb*.*1* at later stages [26]. Finally, *rx3*, a “master eye gene” which confers optic fate in the anterior neural plate [27] and whose loss of function leads to eyeless phenotypes in all vertebrates examined [28] including *Astyanax* [29], was also strongly down-regulated in *cis* (log2(FC)=-1.85; FDR=8.16e-19). In F1 hybrids, *rx3* expression level was intermediate between parental levels, with a biased allelic ratio (**Figure 2AB**). *Rx3* is located ∼320kb away from a “retina thickness” QTL [30]. Since *rx3* expression pattern is altered in many ways in 10hpf cavefish embryos [9], we next sought to disentangle the contribution of changes in *cis*-regulatory sequences to its dysregulation.

**Figure 2:**
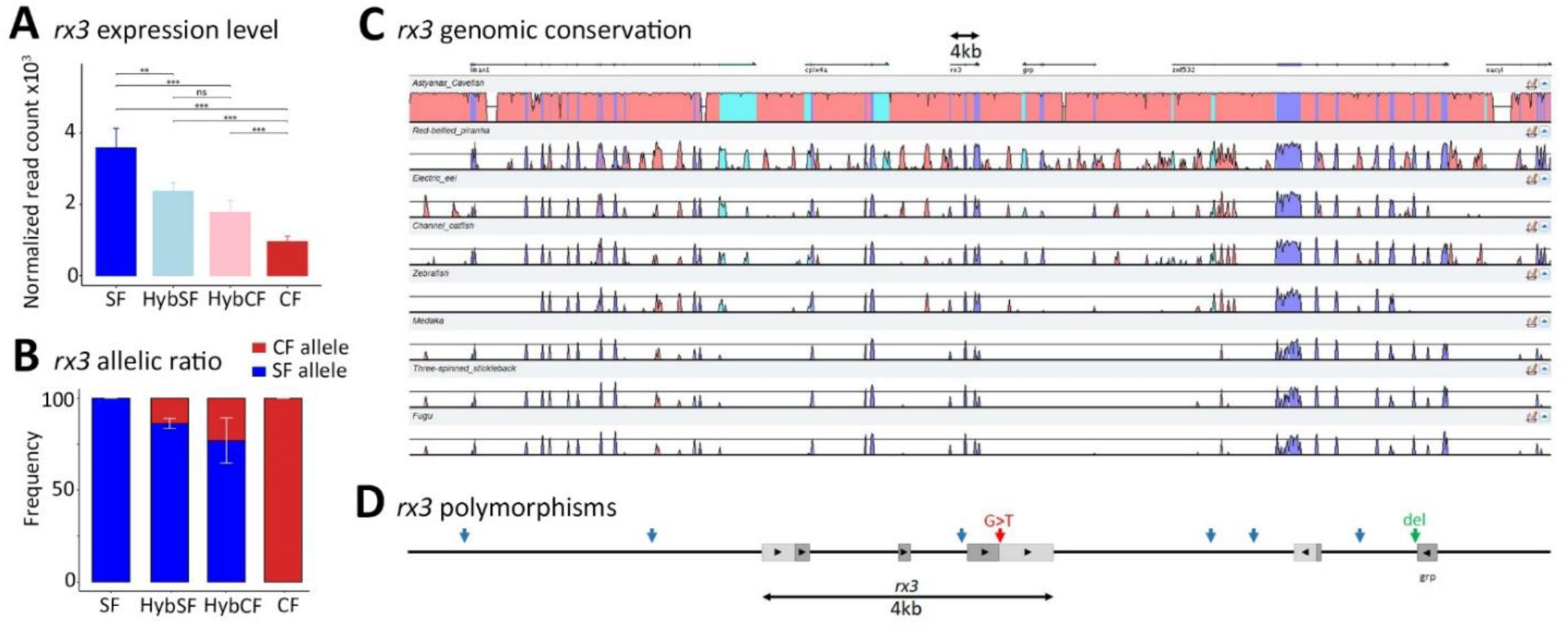
*rx3* regulatory features and landscape. **A, B**, Summary of *rx3* transcriptomic and allelic ratio calculation data. **C**, Vista plot showing alignments of the *rx3* genomic region (30kb) in the two *Astyanax* morphs and 8 additional teleost species. The baseline is the SF reference genome. **D**, Schema of the distribution of the few polymorphisms between SF and CF encountered around the *rx3* gene. See **DataS1** for complete alignments. Blue arrows denote the position of SNPs, the red arrow indicates the SNP used in the *cis/trans* analysis, and the green arrow indicates a 6bp deletion in cavefish.

### A change in *cis* affects *rx3* expression levels but not domain size in cavefish

We performed genomic alignments of *rx3* (5kb upstream of ATG to 5kb downstream of 3’UTR; total ∼15kb) from surface fish of two different origins (Rascon river, n=7 individuals and Choy river, n=9 individuals) against Pachón cavefish (n=9 individuals; all publicly available from [16]). The *rx3* coding sequence was identical in the two morphs. Eight fixed, morph-specific variants were found in non-coding sequences (**Data S1**). Along the scanned genomic region, seven variants corresponded to SNPs (including one in the *rx3* 3’UTR, used in *cis/trans* analysis above) and one corresponded to a small deletion of 6 nucleotides in a CT repeat region in an intron of the 3’ neighbouring gene. The non-coding landscape was extremely conserved in the two *Astyanax* morphs but not in other fish species on a larger genomic scale (30kb), preventing the inference of putative conserved non-coding regulatory elements (**Figure 2CD**). For each of the eight variants identified, +/-15 bp were scanned for putative transcription factor binding site (TFBS). No gain or loss of TFBS were detected, suggesting that the *cis*-regulatory change(s) responsible for the deregulation of *rx3* expression in cavefish must lie further away from the genomic region analysed.

To analyse *rx3* expression regulation at regional and cellular level, we turned to developmental biology approaches. The cavefish “eyefield”, i.e., the territory of the neural plate fated to become the retinas, showed low and heterogeneous *rx3* expression after cell-level quantifications of *rx3* fluorescent *in situ* hybridisation signals, and a smaller differently-shaped expression domain, as compared to SF at 10hpf (**Figure 3ABE**) [9]. In both HybSF and HybCF, eyefield cells presented high-to-intermediate, close to SF, homogeneous fluorescence levels (**Figure 3CDE**). The size of the *rx3* domain was similarly large in SF, HybSF and HybCF but markedly smaller in CF (**Figure 3A-D, F**). These results suggested a dominance of SF alleles on both *rx3*-related phenotypes, i.e., cellular expression levels and domain size. Moreover, as neither cell expression levels nor domain sizes were different between HybSF and HybCF, we concluded that *rx3* expression was not under maternal genetic control (see [13]). Together these results were compatible with a *cis*-regulatory change in the cavefish *rx3* gene affecting the cellular expression levels in the eyefield (HybSF and HybCF cells show lower expression than SF cells) but not the size of its expression domain (HybSF and HybCF domain sizes are as large as SF).

**Figure 3:**
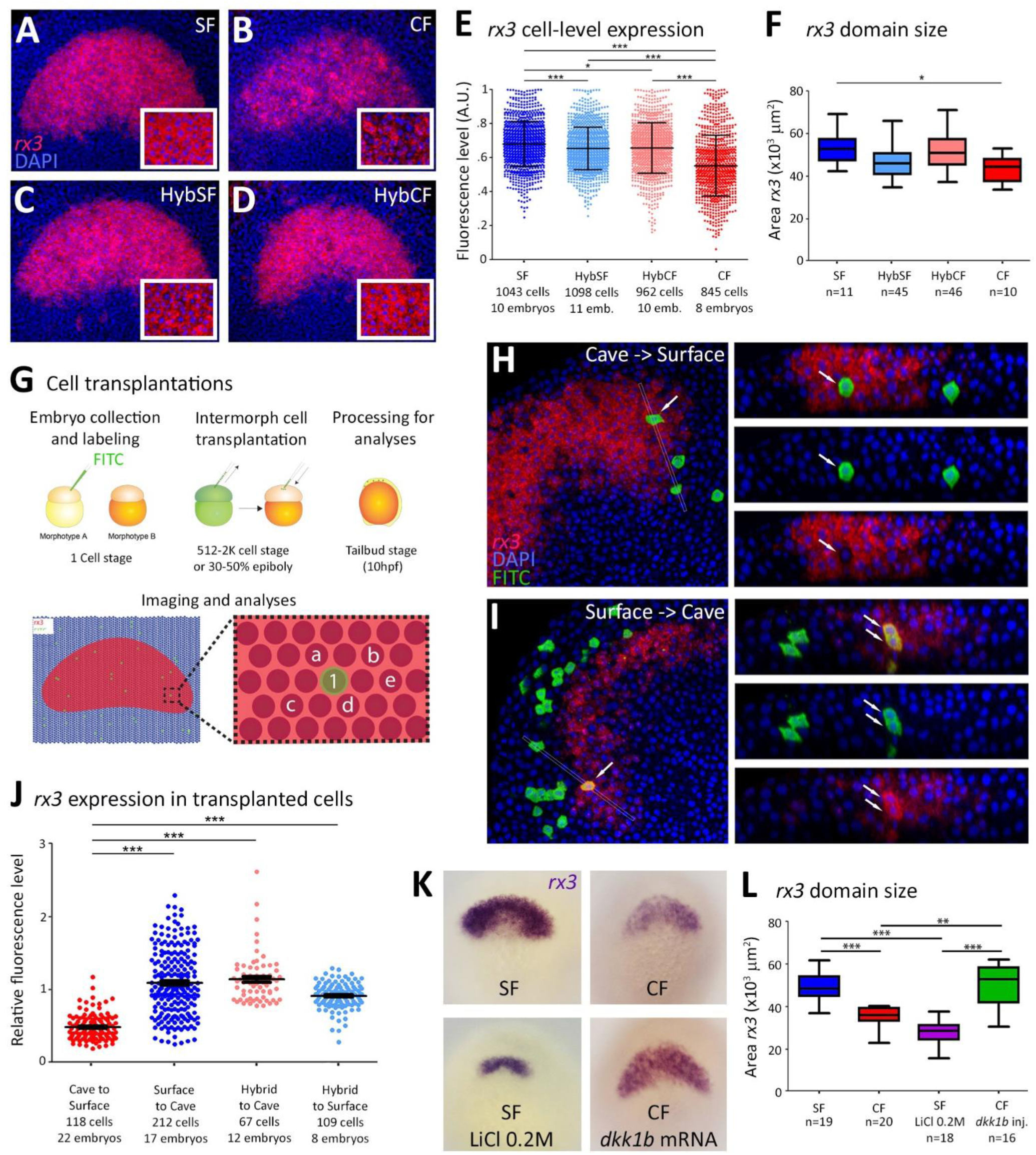
*rx3* regulation at cellular and regional level. **ABCD**, Confocal images of *rx3* fluorescent *in situ* hybridisation (red) at 10hpf, in dorsal views. DAPI nuclear staining is blue. Morphs are indicated. Anterior is up. **E**, Cell level quantification of *rx3* expression signals in SF, CF, HybSF and HybCF. Kruskal-Wallis tests. **F**, Box-plots showing the size area of the *rx3* expression domain in SF, CF, HybSF and HybCF. Kruskal-Wallis tests. **G**, Experimental procedure and principle for cell transplantations and quantification of the cell-level expression signals in chimeric embryos. **H-I**, Confocal images of transplantation results. *In situ* hybridisation for *rx3* is red, grafted cells are green, and cell nuclei are blue. Images on the right are reconstructed sections at the level of light grey lines in left images, showing fluorescence channels decompositions at high magnification. **J**, Cell level quantification of *rx3* expression signals in grafted cells relative to neighbouring host cells. The nature of the donor and host, and the number of cells and embryos analysed are indicated. Kruskal-Wallis tests. **K**, Colorimetric *in situ* hybridisation (purple) for *rx3* at 10hpf, in dorsal views, after the indicated manipulations of the Wnt signalling pathway. **L**, Box-plots showing the size area of the *rx3* expression domain after manipulations of the Wnt pathway shown in K. Kruskal-Wallis tests.

To explore *rx3* regulation at cell level, we performed transplantation experiments. Cells from one morph were transplanted isochronically into embryos of the other morph, either at blastula or gastrula stage [31]. At 10hpf, *rx3* expression levels in grafted cells incorporated in the eyefield were compared to those of the surrounding host cells (**Figure 3GHI**). Irrespective of the grafting stage, CF cells transplanted into SF hosts showed lower *rx3* levels than surrounding host cells, and *vice-versa*, SF cells grafted into a CF environment expressed higher *rx3* levels than neighbouring host cells (**Figure 3HIJ, Figure S3**). These results showed that *rx3* expression level in eyefield cells is controlled autonomously by the *rx3* alleles of the transplanted cells, and do not depend on the cellular environment. Moreover, F1 hybrid cells grafted into SF embryos showed *rx3* levels similar to adjacent SF host cells (**Figure 3J**, pale blue). Finally, F1 hybrid cells grafted into CF embryos behaved like transplanted SF cells (**Figure 3J**, pale red). This suggested that one copy of the SF allele(s) is sufficient to confer proper *rx3* level in eyefield cells. Together these experiments support the existence of a *cis*-regulatory mechanism controlling the quantitative aspect, i.e., the cellular expression level, of *rx3*.

Yet, above experiments could not explain the other aspect of the *rx3* phenotype, i.e., the reduced size of its expression domain in 10hpf cavefish embryos. We reasoned that *rx3* domain size might rather be controlled by signaling pathways that establish early embryonic patterning [32-34]. As we previously described an influence of the Wnt pathway on eye shape and size in *Astyanax* larvae [13], we manipulated Wnt signaling and assessed *rx3* expression at 10hpf (**Figure 3KL**). Overactivation of Wnt signaling in SF by pharmacological treatment (LiCl 0.2M) during a short window of time in late gastrulation strongly affected *rx3* domain size, but expression levels remained high and homogeneous, SF-like, in the tiny eyefield. Conversely, inhibition of Wnt signaling in CF embryos through injection of *dkk1b* mRNA (a Wnt signaling antagonist) at one-cell stage resulted in an expansion of the *rx3*-positive domain but expression level remained low and heterogeneous, CF-like, in the enlarged eyefield.

Together these results demonstrated that the evolution of the regulation of *rx3* domain size and expression level in cavefish are independent and uncoupled. Domain size, hence eyefield and future eye size, depends on non-autonomous signaling mechanisms while expression levels, hence the fate and behavior of optic cells, are cell-autonomously controlled and *cis*-regulated. Thus, cell fate and domain size depend on two distinct *rx3* regulatory modules.

### A contribution of coding mutations to cavefish developmental evolution?

Lastly, we sought to estimate the load of coding mutations during early developmental evolution in cavefish. Among fixed exonic variants identified above (11,336 total), the majority corresponded either to synonymous (n=6054) or missense mutations (n=5282), with a minority predicted to have a strong effect (n=20 in 19 genes; affecting the start/stop codon or generating a frameshift). A closer examination of these “severe” variants revealed that most of them should not change greatly the protein structure or function because they were in the close vicinity of other start or stop codons, respectively.

As any change in amino acid might alter protein function (e.g. [35, 36]), we systematically assessed putative effects of fixed missense variants. Among them, 76% (n=4003) changed either class, charge or polarity of amino acids (**Figure S4A**). To predict the impact of the missense variants on protein function, a pathogenic score was calculated using Mutpred2 [37], after ancestral state reconstruction to orient the mutations in the surface or the cavefish lineage [17]. Hence, 2539 mutations had occurred in CF lineage and 1464 had occurred in SF lineage. About 10% of the mutations displayed a g-score greater than 0.5 (Mutpred2 pathogenicity threshold) in both morphs (n=272/9.3% in CF and n=141/8.2% in SF, respectively). Hence, there was no enrichment for pathogenic mutations in CF compared to SF, suggesting that coding sequences remained mostly unaffected and had little contribution to the evolution cavefish development (**Figure 4A**). This result is in line with the description of only 3 pseudogenes related to loss of eyes and pigmentation in Pachón cavefish [17]. The CF and SF pathogenic gene sets were enriched in “metabolic process (RNA)” and “mRNA processing” GO terms, respectively, suggesting that RNA metabolism genes are globally prone to sequence evolution (**Figure S4B**). Of note, predicted pathogenic mutations were also found on genes involved in circadian rhythm (LOC103027946, C649S, g-score = 0.79, in CF lineage) and developmental pathway (*lrp5*, M698R, g-score = 0.75, in SF lineage). In sum, we found only minor differences on a functional level between SF and CF protein-coding sequences expressed at tailbud stage. This was consistent with the idea that coding mutations in early developmental genes are too deleterious to provide fuel to viable morphological evolution.

**Figure 4:**
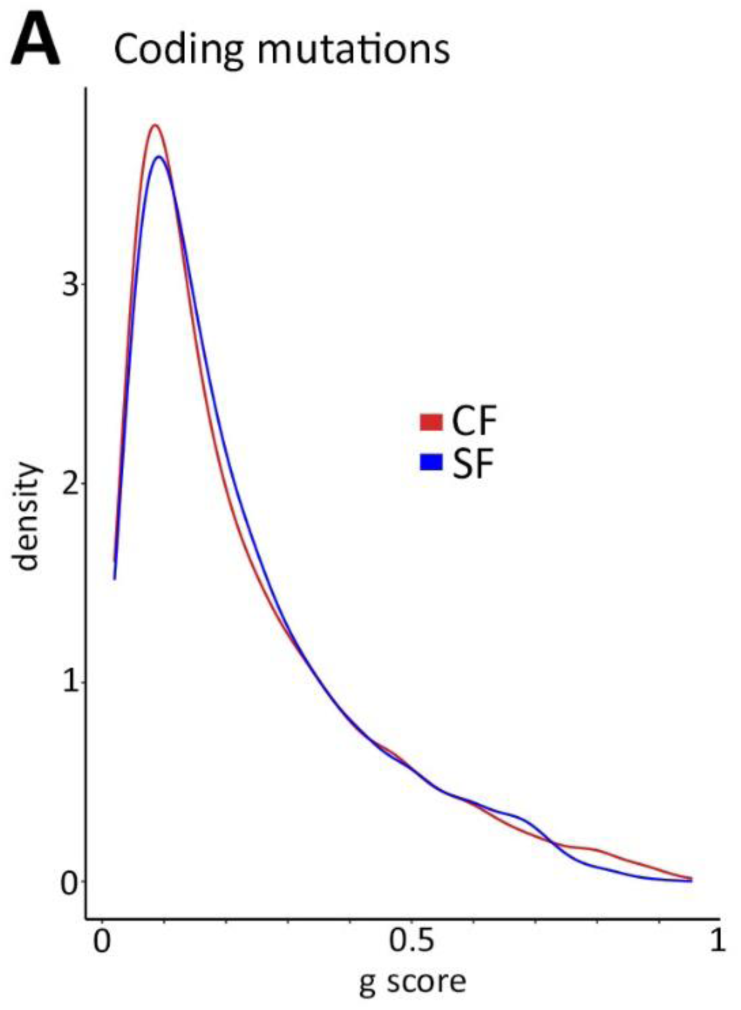
Contribution of coding changes to cavefish developmental evolution. **A**, Plot showing the distribution of predicted non-deleterious (g-score=0) to highly deleterious (g-score=1) effects among coding mutations identified in the SF and CF lineages

## DISCUSSION – CONCLUSION

Looking at changes in gene regulation is critical to understand how species evolve and adapt to their environment. The contribution of *cis*-versus *trans*-regulation to the process is increasingly studied in animals and plants [4, 5, 20-22, 38, 39]. To our knowledge, this work is the first to examine transcriptome-wide the regulatory changes occurring during early developmental evolution in a teleost. In cavefish, similar to domesticated cotton or diversifying drosophilae species, we found that *cis*-regulation is the main contributor to the evolution of gene expression during development. Moreover, the impact of coding mutations is probably marginal, in agreement with the idea that the modular nature of non-coding regulatory elements is more susceptible to give rise to viable changes [2, 40].

Among a list of 108 genes, we have identified *rx3* as an outstanding *cis*-regulated gene, with potential major consequences on the cavefish ocular phenotype [9, 29, 41]. As regional and cell-level information on gene expression is lost in bulk transcriptomics, we have combined insights from bioinformatics and developmental biology to decipher the “*rx3* regulatory puzzle”. We have discovered that its cellular expression level and the size of its expression domain are controlled independently and uncoupled, further highlighting the modularity and the complexity of *rx3* gene regulation evolution. There are famous examples of *cis*-regulatory changes in evolution in the literature, leading to spectacular or more discrete morphological changes e.g. [42-46]. Further genomic analyses of the 108 *cis*-regulated genes we have identified should reveal how cavefish lost their eyes.

## MATERIALS AND METHODS

### *In vitro* fertilization and RNA extraction

Texas surface fish and Pachón cavefish were maintained in an inverted light cycle (12/12) and induced to spawn via temperature change from 21°C to 26°C. Crossed were performed by *in vitro* fertilization and surface, Pachón cave and reciprocal F1 hybrid embryos were collected at 10 hours post fertilization, immersed in Trizol^®^ (Invitrogen) and stored at -80°C until RNA extraction. Hybrids from the cross of a male surface fish and a female cavefish is termed cave hybrid (HybCF). Reciprocal hybrid (cavefish male and surface fish female) is termed surface hybrid (HybSF). Each condition was obtained in triplicate, with each sample being a pool of ∼60 embryos from 2 distinct fertilization (1 female with 3 males each) to ensure a good representation of allele diversity among population. RNA were extracted following a phenol-chloroform procedure.

### RNA-sequencing and analysis

Library preparation and RNA-sequencing was done at the I2BC High-throughput sequencing platform (https://www.i2bc.paris-saclay.fr/spip.php?article399) using an Illumina NextSeq 500 sequencing instrument (paired-end). Analysis was performed on the European Galaxy server (usegalaxy.eu). Reads were mapped onto the Astyanax mexicanus 2.0 genome (NCBI) with HISAT2 (version 2.1.0+galaxy4), after filtering (paired-end single match and Q>20). Reads were counted with htseq-count (version 0.9.1) and differential gene expression was determined with Deseq2 ([47]; version 2.11.40.6). The full dataset is deposited as Sequence Read Archive (SRA) on NCBI under accession BioProject ID PRJNA848099. R scripts used in analyses below are available at https://github.com/LeclercqJ/Retaux-lab.

### Variant calling and filtering

Each RNA-seq experimental replicate originated from 2 *in vitro* fertilizations obtained from 1 female and 3 males. Therefore a maximum of 48 alleles were represented in each condition (4 individuals * 2 alleles * 2 FIV * 3 replicates). Variant calling was performed using FreeBayes (version 1.3.1, [48]). Genes with variant(s) were identified using SnpEff [49] with custom *Astyanax mexicanus* database from SnpEff build and the NCBI GTF file, then filtered on quality >30 with SnpSift (version 4.3). Reads containing fixed variants in parents were extracted using a custom R script and summed for a single gene in case of multiple variants. Parental expression (P) is the result of the combination of both *cis*- and *trans*-regulation. *Cis-*regulation (H) is estimated via allele specific expression in hybrid (i.e. the ratio of cave/surface alleles in hybrid). *Trans-*regulation is then estimated as the parental divergence that is not explained by *cis*-regulation (so Trans=P-H). Parental divergence and allele specific expression in hybrid were calculated using DESeq2 R package [47]. *Cis*- and *trans*-regulation was then estimated using an R script derived from Bao et al, 2019 [20]. Only genes with shared regulation category between surface hybrids and cave hybrids, and differentially-expressed in parents from the DESeq2 analysis on the whole transcript, were further considered. Unsurprisingly for RNAseq data, the majority of fixed variants were annotated in exons and 3’-UTR, with few variants found in 5’-UTR and intron (likely from unspliced mRNA, alternative isoform or error in intron/exon boundary annotation) (**Figure S1B**).

### Gene Ontology (GO) and network analysis

Genes falling under the “*cis*-only” category were analysed for Gene Ontology enrichment using GOEnrichment (Galaxy, version 2.0.1). Gene Product Annotation File for *Astyanax mexicanus* was generated in a previous study [13]. Putative gene interaction network between those genes was predicted using String (v11.5; [50]). Input gene names were manually adjusted (using Ensembl Ids or gene descriptions) to increase the number of entries retrieved. 94 out of 108 genes were retrieved and subjected to the analysis.

### *Rx3* gene alignment and TFBS prediction

The *rx3* genomic region (+/-50kb) and annotation from several fish species (Mexican tetra cave and surface fish, red-bellied piranha, electric eel, channel catfish, zebrafish, fugu, medaka, stickleback and tetraodon) were downloaded from Ensembl (v106) and aligned using mVista LAGAN alignment program (https://genome.lbl.gov/vista/mvista/submit.shtml; min cons width: 20 bp). The annotation of *Astyanax* surface fish was slightly modified to match NCBI exon boundaries. The Pachón cavefish *rx3* genomic region was extracted from the new cavefish genome assembly recently generated [51].

The reads encompassing the *rx3* region of several individuals from the Pachón cave as well as Choy and Rascon rivers (Mexico) were retrieved from Hermann et al. [16]. Pachón cave *rx3* sequences were aligned with each surface population independently with ClustalOmega (www.ebi.ac.uk/Tools/msa/clustalo/) and manually searched for fixed variants shared by both surface populations. We reasoned that a putative *cis-*regulatory candidate element (CRE) should be shared in both surface populations and different in the Pachón cave population. To identify a potential CRE, putative transcription factor binding sites (TFBS) were scanned around each fixed SNP detected +/-15 bp with both TFBSTools R package ([52]; min score > 0.9) and FIMO ([53], https://meme-suite.org/meme/tools/fimo, p-value of 10^−4^). Position Weight Matrix (PFM) from JASPAR2020 (Vertebrate, Core collection) were used. Predicted TFBS common with both method and unique to one morph were examined further.

### Fixed variant effect prediction

Fixed SNPs were extracted from the VCF file using VCF-Bed intersect on Galaxy and variant effects were predicted with SnpEff (canonical transcript only, custom database). Nonsense variants and premature start variants were manually checked (see Results). Missense variants were analysed with a R custom script. Protein sequences were retrieved from NCBI with the rentrez R package and corrected for discrepancy between sequences and annotations (150 variants/75 genes corrected; 2 genes/5 variants removed).

For each gene with a missense variant between surface and cave morphs, orthologous sequences from 9 other teleost species with annotated genomes (*Danio rerio*: GCF_000002035.6, *Pygocentrus nattereri*: GCF_015220715.1, *Electrophorus electricus*: GCF_013358815.1, *Anguilla anguilla*: GCF_013347855.1, *Tachysurus fulvidraco*: GCF_003724035.1, *Ameiurus melas*: GCA_012411365.1, *Triplophysa tibetana*: GCA_008369825.1, *Anabarilius grahami*: GCA_003731715.1, *Labeo rohita*: GCA_004120215.1) were retrieved using *BROCCOLI* (https://doi.org/10.1093/molbev/msaa159), which uses kmer clustering and phylogenetic approaches to infer orthologous groups.

Protein sequences of *A. mexicanus* surface and cave morphs were aligned with retrieved orthologous sequences using MAFFT v7.407 (https://academic.oup.com/mbe/article/30/4/772/1073398) and PAML v4.9h (https://doi.org/10.1093/molbev/msm088) was used for the reconstruction of ancestral sequences. Finally, for each gene, the ancestral sequence of either cave or surface morph was used as input in MutPred2 [33] to assign a deleterious score (g-score) to all missense mutations in these two lineages, which can vary between 0 (non-deleterious) and 1 (highly deleterious). Variants with a g-score >0.5 were analysed for Gene Ontology enrichment using GOEnrichment (Galaxy, version 2.0.1).

### In situ hybridization

Colorimetric and fluorescent *in situ* hybridization were performed as previously described [9]. Fluorescently labeled embryos were counterstained with DAPI, dissected and flat mounted in Vectashield antifade mounting media (Vector Laboratories) for imaging (SP8 confocal microscope, Leica). Non-fluorescent embryos were imaged in whole mount (Nikon macroscope).

### Cell transplantations and quantifications

Isochronic transplantations of FITC-labeled embryonic cells, and *in situ* hybridization followed by FITC revelation were performed as previously described [31]. Area measurements and quantification of fluorescence intensity in individual cells, semi-automatically segmented [9], were performed in Fiji.

### Manipulation of the Wnt pathway

To manipulate Wnt signaling levels, surface embryos were incubated in LiCl_2_ 0.2M in Embryo Medium from 90% of epiboly to tailbud stage, and cavefish embryos were injected at the one-cell stage with full-length *A. mexicanus dkk1b* mRNA [13]. Statistical tests were done in Graph pad Prism 5.

## Supplemental Information files

**Supplemental Figure 1:**
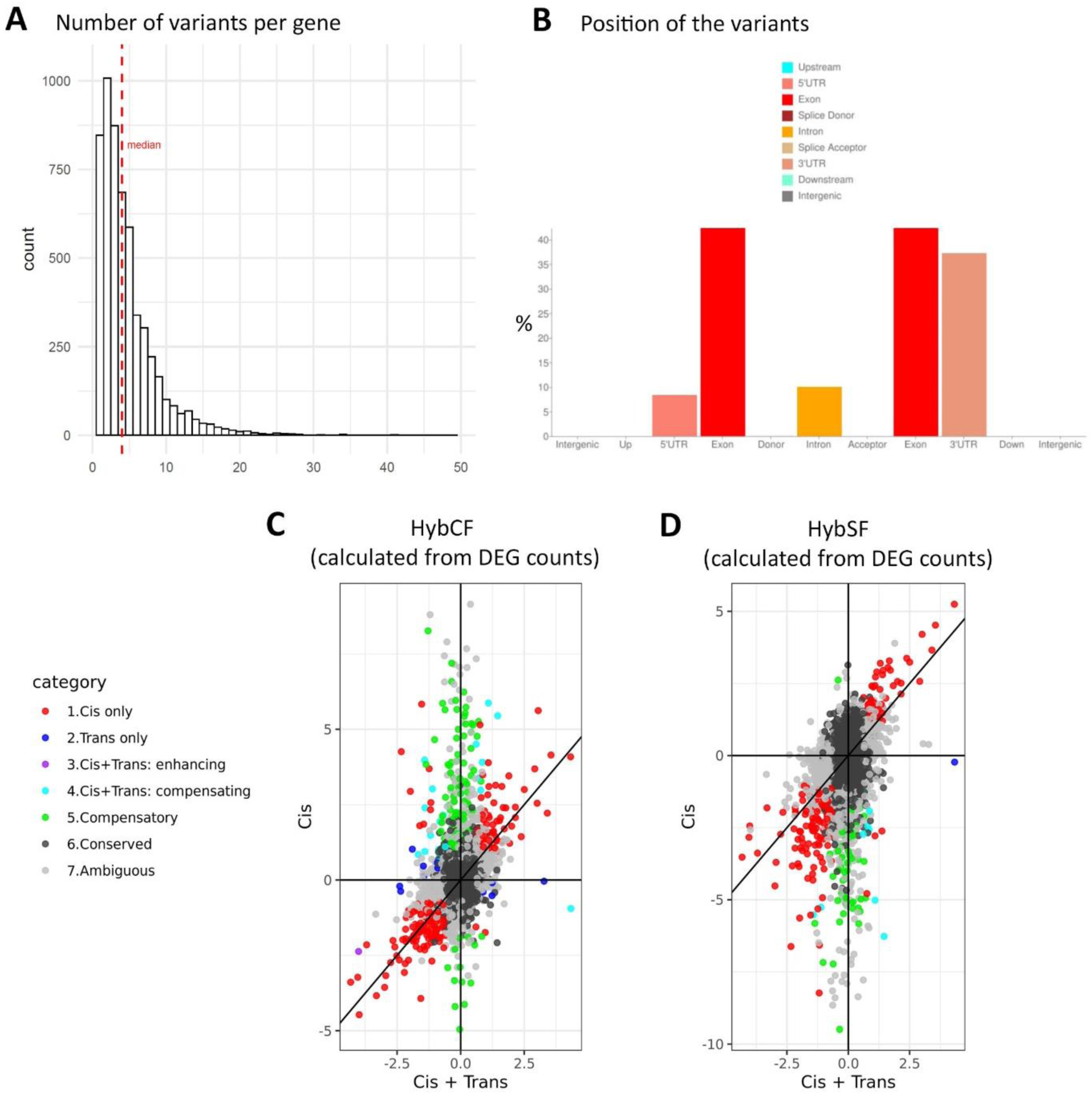
Using fixed variants for the analysis of *cis*-or *trans*-regulated genes. Related to Figure 1. **A**, Histogram of the distribution of the number of fixed variants per gene in the whole dataset. There were 26,390 fixed variants, corresponding to 5573 genes, among which 900 DEGs between SF and CF. A maximum of 49 variants was found on a single gene (LOC103024947, titin-like, a >200kb “giant” gene). **B**, Histogram showing the distribution of the position of the fixed variants analyzed. **C**,**D**, Scatter plots of *cis* regulatory divergence (y axis; from allelic expression ratios in hybrids) versus parental divergence (x-axis; from DEG in parents), where the different categories of genes are color-coded. These plots show results after computation and allelic expression counts from reads on the entire transcripts (i.,e., from the DEG dataset) whereas the main Figure 1 (panels EF) show the results after computation and allelic expression counts from reads exclusively encompassing the variant position. C is from HybSF analysis and D is from HybSF analysis.

**Supplemental Figure 2:**
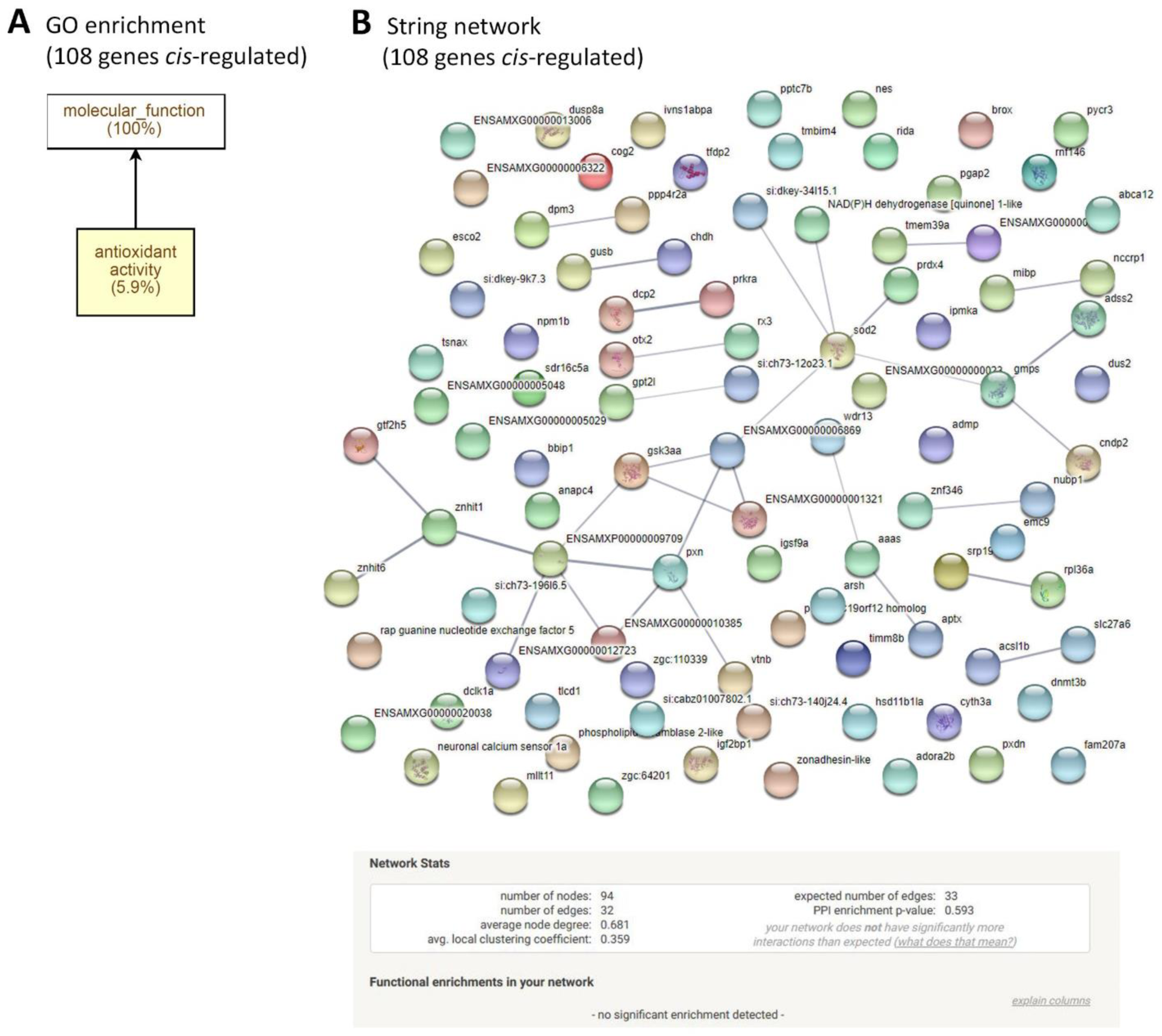
Gene Ontology and network analyses of the 108 genes de-regulated in *cis* identified at 10hpf in *Astyanax* embryos. **A**, GO analysis did not reveal specific enrichment of gene categories among the 108 genes list. **B**, Network analysis using String did not show a significant number of connections or clusters among the 108 genes list. Line thickness indicates the strength of data support.

**Supplemental Figure 3:**
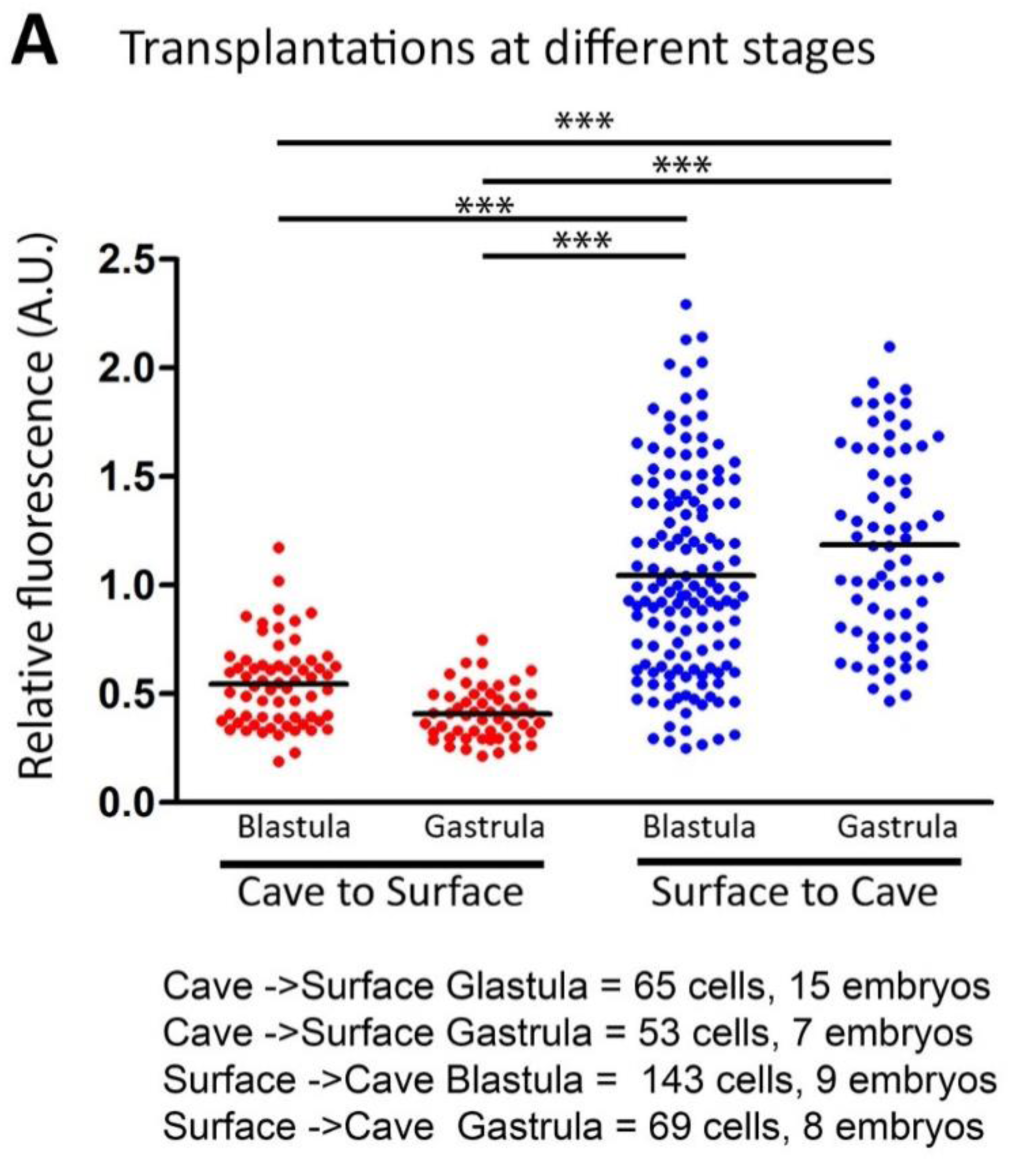
Results from inter-morph cell transplantations performed at blastula or gastrula stage. Related to Figure 3. **A**, Cell level quantification of *rx3* expression signals in grafted cells relative to neighbouring host cells. The stage of transplantation, the nature of the donor and host, and the number of cells and embryos analysed are indicated. Mann-Whitney tests. As the results were identical whether cells were transplanted at the blastula or the gastrula stage, in the main Figure 3 (panels J) the result are presented after pooling the two conditions.

**Supplemental Figure 4:**
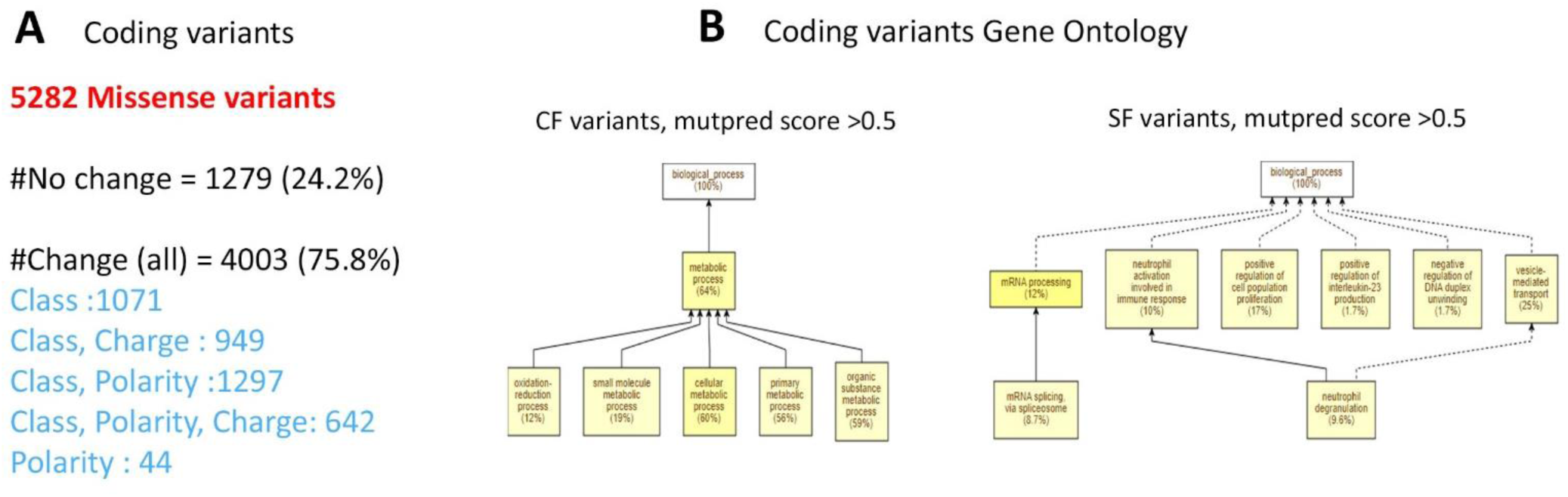
Features of the coding variants identified among transcripts expressed at 10hpf in *Astyanax* embryos. Related to Figure 4. **A**, The proportions of the categories of variants are indicated, including the types of amino acid changes involved. **B**, GO analysis on the coding variants identified in the CF lineage (left) and the SF lineage (right) that displayed g-score greater than 0.5 (Mutpred2 pathogenicity threshold).

