## Supplemental Data S1 for "Evolution of the regulation of developmental gene expression in blind Mexican cavefish"

```
Rascon surface fish (and reference) aligned with Pachon cavefish
Rx3 genomic region
Reference surface genome == Surface; NCBI annotation
CLUSTAL O(1.2.4) multiple sequence alignment (https://www.ebi.ac.uk/Tools/msa/clustalo/)

Rascon4         ACTGATAGAGACATAAGGAGGAAAGGATGGTCAACACCATAAAAGTTAAACACAGGCCTG        60
Surface         ACTGATAGAGACATAAGGAGGAAAGGATGGTCAACACCATAAAAGTTAAACACAGGCCTG        60
Rascon8         ACTGATAGAGACATAAGGAGGAAAGGATGGTCAACACCATAAAAGTTAAACACAGGCCTG        60
Rascon2         ACTGATAGAGACATAAGGAGGAAAGGATGGTCAACACCATAAAAGTTAAACACAGGCCTG        60
Rascon15        ACTGATAGAGACATAAGGAGGAAAGGATGGTCAACACCATAAAAGTTAAACACAGGCCTG        60
Rascon13        ACTGATAGAGACATAAGGAGGAAAGGATGGTCAACACCATAAAAGTTAAACACAGGCCTG        60
Rascon6         ACTGATAGAGACATAAGGAGGAAAGGATGGTCAACACCATAAAAGTTAAACACAGGCCTG        60
Pachon14        ACTGATAGAGACATAAGGAGGAAAGGATGGTCAACACCATAAAAGTTAAACACAGGCCTG        60
Pachon9         ACTGATAGAGACATAAGGAGGAAAGGATGGTCAACACCATAAAAGTTAAACACAGGCCTG        60
Pachon17        ACTGATAGAGACATAAGGAGGAAAGGATGGTCAACACCATAAAAGTTAAACACAGGCCTG        60
Pachon12        ACTGATAGAGACATAAGGAGGAAAGGATGGTCAACACCATAAAAGTTAAACACAGGCCTG        60
Pachon11        ACTGATAGAGACATAAGGAGGAAAGGATGGTCAACACCATAAAAGTTAAACACAGGCCTG        60
Pachon7         ACTGATAGAGACATAAGGAGGAAAGGATGGTCAACACCATAAAAGTTAAACACAGGCCTG        60
Pachon3         ACTGATAGAGACATAAGGAGGAAAGGATGGTCAACACCATAAAAGTTAAACACAGGCCTG        60
Pachon8         ACTGATAGAGACATAAGGAGGAAAGGATGGTCAACACCATAAAAGTTAAACACAGGCCTG        60
Pachon15        ACTGATAGAGACATAAGGAGGAAAGGATGGTCAACACCATAAAAGTTAAACACAGGCCTG        60
                ************************************************************

Rascon4         GACGTAAAATGCCAAATGAACTGATTACAAAGTGCTTTTTAAGTTACCTTGAGTTACCTC        120
Surface         GACGTAAAATGCCAAATGAACTGATTACAAAGTGCTTTTTAAGTTACCTTGAGTTACCTC        120
Rascon8         GACGTAAAATGCCAAATGAACTGATTACAAAGTGCTTTTTAAGTTACCTTGAGTTACCTC        120
Rascon2         GATGTAAAATGCCAAATGAACTGATTACAAAGTGCTTTTTAAGTTACCTTGAGTTACCTC        120
Rascon15        GACGTAAAATGCCAAATGAACTGATTACAAAGTGCTTTTTAAGTTACCTTGAGTTACCTC        120
Rascon13        GACGTAAAATGCCAAATGAACTGATTACAAAGTGCTTTTTAAGTTACCTTGAGTTACCTC        120
Rascon6         GACGTAAAATGCCAAATGAACTGATTACAAAGTGCTTTTTAAGTTACCTTGAGTTACCTC        120
Pachon14        GACGTAAAATGCCAAATGAACTGATTACAAAGTGCTTTTTAAGTTACCTTGAGTTACCTC        120
Pachon9         GACGTAAAATGCCAAATGAACTGATTACAAAGTGCTTTTTAAGTTACCTTGAGTTACCTC        120
Pachon17        GACGTAAAATGCCAAATGAACTGATTACAAAGTGCTTTTTAAGTTACCTTGAGTTACCTC        120
Pachon12        GACGTAAAATGCCAAATGAACTGATTACAAAGTGCTTTTTAAGTTACCTTGAGTTACCTC        120
Pachon11        GACGTAAAATGCCAAATGAACTGATTACAAAGTGCTTTTTAAGTTACCTTGAGTTACCTC        120
Pachon7         GACGTAAAATGCCAAATGAACTGATTACAAAGTGCTTTTTAAGTTACCTTGAGTTACCTC        120
Pachon3         GACGTAAAATGCCAAATGAACTGATTACAAAGTGCTTTTTAAGTTACCTTGAGTTACCTC        120
Pachon8         GACGTAAAATGCCAAATGAACTGATTACAAAGTGCTTTTTAAGTTACCTTGAGTTACCTC        120
Pachon15        GACGTAAAATGCCAAATGAACTGATTACAAAGTGCTTTTTAAGTTACCTTGAGTTACCTC        120
                ** *********************************************************

Rascon4         ATATTTTTTATGTTTGCATCTCTGGACATTAATAAATGAGCTGAATTTGAAATATATTTA        180
Surface         ATATTTTTTATGTTTGCATCTCTGGACATTAATAAATGAGCTGAATTTGAAATATATTTA        180
Rascon8         ATATTTTTTATGTTTGCATCTCTGGACATTAATAAATGAGCTGAATTTGAAATATATTTA        180
Rascon2         ATATTTTTTATGTTTGCA--TCTGGACATTAATAAATGAGCTGAATTTGAAATATATTTA        178
Rascon15        ATATTTTTTATGTTTGCATCTCTGGACATTAATAAATGAGCTGAATTTGAAATATATTTA        180
Rascon13        ATATTTTTTATGTTTGCATCTCTGGACATTAATAAATGAGCTGAATTTGAAATATATTTA        180
Rascon6         ATATTTGTTATGTTTGCATCTCTGGACATTAATAAATGAGCTGAATTTGAAATATATTTA        180
Pachon14        ATATTTTTTATGTTTGCATCTCTGGACATTAATAAATGAGCTGAATTTGAAATATATTTA        180
Pachon9         ATATTTTTTATGTTTGCATCTCTGGACATTAATAAATGAGCTGAATTTGAAATATATTTA        180
Pachon17        ATATTTTTTATGTTTGCATCTCTGGACATTAATAAATGAGCTGAATTTGAAATATATTTA        180
Pachon12        ATATTTTTTATGTTTGCATCTCTGGACATTAATAAATGAGCTGAATTTGAAATATATTTA        180
Pachon11        ATATTTTTTATGTTTGCATCTCTGGACATTAATAAATGAGCTGAATTTGAAATATATTTA        180
Pachon7         ATATTTTTTATGTTTGCATCTCTGGACATTAATAAATGAGCTGAATTTGAAATATATTTA        180
Pachon3         ATATTTTTTATGTTTGCATCTCTGGACATTAATAAATGAGCTGAATTTGAAATATATTTA        180
Pachon8         ATATTTTTTATGTTTGCATCTCTGGACATTAATAAATGAGCTGAATTTGAAATATATTTA        180
Pachon15        ATATTTTTTATGTTTGCATCTCTGGACATTAATAAATGAGCTGAATTTGAAATATATTTA        180
                ****** ***********  ****************************************

Rascon4         ATATTTTTGATTACCTCCATTTCTCCATTTACTGTTTAGTTCCATCTTTAGATTTGGACA        240
Surface         ATATTTTTGATTACCTCCATTTCTCCATTTACTGTTTAGTTCCATCTTTAGATTTGGACA        240
Rascon8         ATATTTTTGATTACCTCCATTTCTCCATTTACTGTTTAGTTCCATCTTTAGATTTGGACA        240
Rascon2         ATATTTTTGATTACCTCCATTTCTCCATTTACTGTTTAGTTCCATCTTTAGATTTGGACA        238
Rascon15        ATATTTTTGATTACCTCCATTTCTCCATTTACTGTTTAGTTCCATCTTTAGATTTGGACA        240
Rascon13        ATATTTTTGATTACCTCCATTTCTCCATTTACTGTTTAGTTCCATCTTTAGATTTGGACA        240
Rascon6         ATATTTTTGATTACCTCCATTTCTCCATTTACTGTTTAGTTCCATCTTTAGATTTGGACA        240
Pachon14        ATATTTTTGATTACCTCCATTTCTCCATTTACTGTTTAGTTCCATCTTTAGATTTGGACA        240
Pachon9         ATATTTTTGATTACCTCCATTTCTCCATTTACTGTTTAGTTCCATCTTTAGATTTGGACA        240
Pachon17        ATATTTTTGATTACCTCCATTTCTCCATTTACTGTTTAGTTCCATCTTTAGATTTGGACA        240
Pachon12        ATATTTTTGATTACCTCCATTTCTCCATTTACTGTTTAGTTCCATCTTTAGATTTGGACA        240
Pachon11        ATATTTTTGATTACCTCCATTTCTCCATTTACTGTTTAGTTCCATCTTTAGATTTGGACA        240
Pachon7         ATATTTTTGATTACCTCCATTTCTCCATTTACTGTTTAGTTCCATCTTTAGATTTGGACA        240
Pachon3         ATATTTTTGATTACCTCCATTTCTCCATTTACTGTTTAGTTCCATCTTTAGATTTGGACA        240
Pachon8         ATATTTTTGATTACCTCCATTTCTCCATTTACTGTTTAGTTCCATCTTTAGATTTGGACA        240
Pachon15        ATATTTTTGATTACCTCCATTTCTCCATTTACTGTTTAGTTCCATCTTTAGATTTGGACA        240
                ************************************************************
```

```
Rascon4      CCCATCTCACATTAAGTCCCCAGACTGGAAGGTCTATGACAAATGGAGTGCAGATCTCCG   300
Surface      CCCATCTCACATTAAGTCCCCAGACTGGAAGGTCTATGACAAATGGAGTGCAGATCTCCG   300
Rascon8      CCCATCTCACATTAAGTCCCCAGACTGGAAGGTCTATGACAAATGGAGTGCAGATCTCCG   300
Rascon2      CCCATCTCACATTAAGTCCCCAGACTGGAAGGTCTATGACAAATGGAGTGCAGATCTCCG   298
Rascon15     CCCATCTCACATTAAGTCCCCAGACTGGAAGGTCTATGACAAATGGAGTGCAGATCTCCG   300
Rascon13     CCCATCTCACATTAAGTCCCCAGACTGGAAGGTCTATGACAAATGGAGTGCAGATCTCTG   300
Rascon6      CCCATCTCACATTAAGTCCCCAGACTGGAAGGTCTATGACAAATGGAGTGCAGATCTCCG   300
Pachon14     CCCATCTCACATTAAGTCCCCAGACTGGAAGGTCTATGACAAATGGAGTGCAGATCTCCG   300
Pachon9      CCCATCTCACATTAAGTCCCCAGACTGGAAGGTCTATGACAAATGGAGTGCAGATCTCCG   300
Pachon17     CCCATCTCACATTAAGTCCCCAGACTGGAAGGTCTATGACAAATGGAGTGCAGATCTCCG   300
Pachon12     CCCATCTCACATTAAGTCCCCAGACTGGAAGGTCTATGACAAATGGAGTGCAGATCTCCG   300
Pachon11     CCCATCTCACATTAAGTCCCCAGACTGGAAGGTCTATGACAAATGGAGTGCAGATCTCCG   300
Pachon7      CCCATCTCACATTAAGTCCCCAGACTGGAAGGTCTATGACAAATGGAGTGCAGATCTCCG   300
Pachon3      CCCATCTCACATTAAGTCCCCAGACTGGAAGGTCTATGACAAATGGAGTGCAGATCTCCG   300
Pachon8      CCCATCTCACATTAAGTCCCCAGACTGGAAGGTCTATGACAAATGGAGTGCAGATCTCCG   300
Pachon15     CCCATCTCACATTAAGTCCCCAGACTGGAAGGTCTATGACAAATGGAGTGCAGATCTCCG   300
             ********************************************************** *

Rascon4      GCAAGTATATGAATTTATGGAATAAGCATAAAAAATAAGCAAGGAATAAGCATATCTTTA   360 SNP1 also
Surface      GCAAGTATATGAATTTATGGAATAAGCATAAAAAATAAGCAAGGAATAAGCATATCTTTA   360 Fixed in
Rascon8      GCAAGTATATGAATTTATGGAATAAGCATAAAAAATAAGCAAGGAATAAGCATATCTTTA   360 Choy SF
Rascon2      GCAAGTATATGAATTTATGGAATAAGCATAAAAAATAAGCAAGGAATAAGCATATCTTTA   358
Rascon15     GCAAGTATATGAATTTATGGAATAAGCATAAAAAATAAGCAAGGAATAAGCATATCTTTA   360
Rascon13     GCAAGTATATGAATTTATGGAATAAGCATAAAAAATAAGCAAGGAATAAGCATATCTTTA   360
Rascon6      GCAAGTATATGAATTTATGGAATAAGCATAAAAAATAAGCAAGGAATAAGCATATCTTTA   360
Pachon14     GCAAGTATATGAATTTATGGAATAAGCATAAAAGATAAGCAAGGAATAAGCATATCTTTA   360
Pachon9      GCAAGTATATGAATTTATGGAATAAGCATAAAAGATAAGCAAGGAATAAGCATATCTTTA   360
Pachon17     GCAAGTATATGAATTTATGGAATAAGCATAAAAGATAAGCAAGGAATAAGCATATCTTTA   360
Pachon12     GCAAGTATATGAATTTATGGAATAAGCATAAAAGATAAGCAAGGAATAAGCATATCTTTA   360
Pachon11     GCAAGTATATGAATTTATGGAATAAGCATAAAAGATAAGCAAGGAATAAGCATATCTTTA   360
Pachon7      GCAAGTATATGAATTTATGGAATAAGCATAAAAGATAAGCAAGGAATAAGCATATCTTTA   360
Pachon3      GCAAGTATATGAATTTATGGAATAAGCATAAAAGATAAGCAAGGAATAAGCATATCTTTA   360
Pachon8      GCAAGTATATGAATTTATGGAATAAGCATAAAAGATAAGCAAGGAATAAGCATATCTTTA   360
Pachon15     GCAAGTATATGAATTTATGGAATAAGCATAAAAGATAAGCAAGGAATAAGCATATCTTTA   360
             ********************************* **************************

Rascon4      TTACCAAAGCGTAGGCTGAAAGTGCAGCACAAGATGCTGTATCTGGAAACAGTATGAAAA   420
Surface      TTACCAAAGCGTAGGCTGAAAGTGCAGCACAAGATGCTGTATCTGGAAACAGTATGAAAA   420
Rascon8      TTACCAAAGCGTAGGCTGAAAGTGCAGCACAAGATGCTGTATCTGGAAACAGTATGAAAA   420
Rascon2      TTACCAAAGCGTAGGCTGAAAGTGCAGCACAAGATGCTGTATCTGGAAACAGTATGAAAA   418
Rascon15     TTACCAAAGGGTAGGCTGAAAGTGCAGCACAAGATGCTGTATCTGGAAACAGTATGAAAA   420
Rascon13     TTACCAAAGCGTAGGCTGAAAGTGCAGCACAAGATGCTGTATCTGGAAACAGTATGAAAA   420
Rascon6      TTACCAAAGGGTAGGCTGAAAGTGCAGCACAAGATGCTGTATCTGGAAACAGTATGAAAA   420
Pachon14     TTACCAAAGCGTAGGCTGAAAGTGCAGCACAAGATGCTGTATCTGGAAACAGTATGAAAA   420
Pachon9      TTACCAAAGCGTAGGCTGAAAGTGCAGCACAAGATGCTGTATCTGGAAACAGTATGAAAA   420
Pachon17     TTACCAAAGCGTAGGCTGAAAGTGCAGCACAAGATGCTGTATCTGGAAACAGTATGAAAA   420
Pachon12     TTACCAAAGCGTAGGCTGAAAGTGCAGCACAAGATGCTGTATCTGGAAACAGTATGAAAA   420
Pachon11     TTACCAAAGCGTAGGCTGAAAGTGCAGCACAAGATGCTGTATCTGGAAACAGTATGAAAA   420
Pachon7      TTACCAAAGCGTAGGCTGAAAGTGCAGCACAAGATGCTGTATCTGGAAACAGTATGAAAA   420
Pachon3      TTACCAAAGCGTAGGCTGAAAGTGCAGCACAAGATGCTGTATCTGGAAACAGTATGAAAA   420
Pachon8      TTACCAAAGCGTAGGCTGAAAGTGCAGCACAAGATGCTGTATCTGGAAACAGTATGAAAA   420
Pachon15     TTACCAAAGCGTAGGCTGAAAGTGCAGCACAAGATGCTGTATCTGGAAACAGTATGAAAA   420
             ********* **************************************************

Rascon4      CTCTAGAATTAAGCTCTTATTTAAAGATTAAGCAACTTAGCTTAGGCACTAAATATCTGA   480
Surface      CTCTAGAATTAAGCTCTTATTTAAAGATTAAGCAACTTAGCTTAGGCACTAAATATCTGA   480
Rascon8      CTCTAGAATTAAGCTCTTATTTAAAGATTAAGCAACTTAGCTTAGGCACTAAATATCTGA   480
Rascon2      CTCTAGAATTAAGCTCTTATTTAAAGATTAAGCAACTTAGCTTAGGCACTAAATATCTGA   478
Rascon15     CTCTAGAATTAAGCTCTTATTTAAAGATTAAGCAACTTAGCTTAGGCACTAAATATCTGA   480
Rascon13     CTCTAGAATTAAGCTCTTATTTAAAGATTAAGCAACTTAGCTTAGGCACTAAATATCTGA   480
Rascon6      CTCTAGAATTAAGCTCTTATTTAAAGATTAAGCAACTTAGCTTAGGCACTAAATATCTGA   480
Pachon14     CTCTAGAATTAAGCTCTTATTTAAAGATTAAGCAACTTAGCTTAGGCACTAAATATCTGA   480
Pachon9      CTCTAGAATTAAGCTCTTATTTAAAGATTAAGCAACTTAGCTTAGGCACTAAATATCTGA   480
Pachon17     CTCTAGAATTAAGCTCTTATTTAAAGATTAAGCAACTTAGCTTAGGCACTAAATATCTGA   480
Pachon12     CTCTAGAATTAAGCTCTTATTTAAAGATTAAGCAACTTAGCTTAGGCACTAAATATCTGA   480
Pachon11     CTCTAGAATTAAGCTCTTATTTAAAGATTAAGCAACTTAGCTTAGGCACTAAATATCTGA   480
Pachon7      CTCTAGAATTAAGCTCTTATTTAAAGATTAAGCAACTTAGCTTAGGCACTAAATATCTGA   480
Pachon3      CTCTAGAATTAAGCTCTTATTTAAAGATTAAGCAACTTAGCTTAGGCACTAAATATCTGA   480
Pachon8      CTCTAGAATTAAGCTCTTATTTAAAGATTAAGCAACTTAGCTTAGGCACTAAATATCTGA   480
Pachon15     CTCTAGAATTAAGCTCTTATTTAAAGATTAAGCAACTTAGCTTAGGCACTAAATATCTGA   480
             ************************************************************

Rascon4      AGTGCACCAACACGCTGTAATCTGCCAGATAAAGAAAGAAATGGGTTTTAATTTAATTGG   540
Surface      AGTGCACCAACACGCTGTAATCTGCCAGATAAAGAAAGAAATGGGTTTTAATTTAATTGG   540
Rascon8      AGTGCACCAACACGCTGTAATCTGCCAGATAAAGAAAGAAATGGGTTTTAATTTAATTGG   540
Rascon2      AGTGCACCAACACGCTGTAATCTGCCAGATAAAGAAAGAAATGGGTTTTAATTTAATTGG   538
Rascon15     AGTGCACCAACACGCTGTAATCTGCCAGATAAAGAAAGAAATGGGTTTTAATTTAATTGG   540
```

```
Rascon13     AGTGCACCAACACGCTGTAATCTGCCAGATAAAGAAAGAAATGGGTTTTAATTTAATTGG     540
Rascon6      AGTGCACCAACACGCTGTAATCTGCCAGATAAAGAAAGAAATGGGTTTTAATTTAATTGG     540
Pachon14     AGTGCACCAACACGCTGTAATCTGCCAGATAAAGAAAGAAATGGGTTTTAATTTAATTGG     540
Pachon9      AGTGCACCAACACGCTGTAATCTGCCAGATAAAGAAAGAAATGGGTTTTAATTTAATTGG     540
Pachon17     AGTGCACCAACACGCTGTAATCTGCCAGATAAAGAAAGAAATGGGTTTTAATTTAATTGG     540
Pachon12     AGTGCACCAACACGCTGTAATCTGCCAGATAAAGAAAGAAATGGGTTTTAATTTAATTGG     540
Pachon11     AGTGCACCAACACGCTGTAATCTGCCAGATAAAGAAAGAAATGGGTTTTAATTTAATTGG     540
Pachon7      AGTGCACCAACACGCTGTAATCTGCCAGATAAAGAAAGAAATGGGTTTTAATTTAATTGG     540
Pachon3      AGTGCACCAACACGCTGTAATCTGCCAGATAAAGAAAGAAATGGGTTTTAATTTAATTGG     540
Pachon8      AGTGCACCAACACGCTGTAATCTGCCAGATAAAGAAAGAAATGGGTTTTAATTTAATTGG     540
Pachon15     AGTGCACCAACACGCTGTAATCTGCCAGATAAAGAAAGAAATGGGTTTTAATTTAATTGG     540
             ************************************************************

Rascon4      AGATTATTTCCCAACTGTTGCATCCTTGATTAGTTAAAGGGAGGAGGAATTGGGCCTATT     600
Surface      AGATTATTTCCCAACTGTTGCATCCTTGATTAGTTAAAGGGAGGAGGAATTGGGCCTATT     600
Rascon8      AGATTATTTCCCAACTGTTGCATCCTTGATTAGTTAAAGGGAGGAGGAATTGGGCCTATT     600
Rascon2      AGATTATTTCCCAACTGTTGCATCCTTGATTAGTTAAAGGGAGGAGGAATTGGGCCTATT     598
Rascon15     AGATTATTTCCCAACTGTTGCATCCTTGATTAGTTAAAGGGAGGAGGAATTGGGCCTATT     600
Rascon13     AGATTATTTCCCAACTGTTGCATCCTTGATTAGTTAAAGGGAGGAGGAATTGGGCCTATT     600
Rascon6      AGATTATTTCCCAACTGTTGCATCCTTGATTAGTTAAAGGGAGGAGGAATTGGGCCTATT     600
Pachon14     AGATTATTTCCCAACTGTTGCATCCTTGATTAGTTAAAGGGAGGAGGAATTGGGCCTATT     600
Pachon9      AGATTATTTCCCAACTGTTGCATCCTTGATTAGTTAAAGGGAGGAGGAATTGGGCCTATT     600
Pachon17     AGATTATTTCCCAACTGTTGCATCCTTGATTAGTTAAAGGGAGGAGGAATTGGGCCTATT     600
Pachon12     AGATTATTTCCCAACTGTTGCATCCTTGATTAGTTAAAGGGAGGAGGAATTGGGCCTATT     600
Pachon11     AGATTATTTCCCAACTGTTGCATCCTTGATTAGTTAAAGGGAGGAGGAATTGGGCCTATT     600
Pachon7      AGATTATTTCCCAACTGTTGCATCCTTGATTAGTTAAAGGGAGGAGGAATTGGGCCTATT     600
Pachon3      AGATTATTTCCCAACTGTTGCATCCTTGATTAGTTAAAGGGAGGAGGAATTGGGCCTATT     600
Pachon8      AGATTATTTCCCAACTGTTGCATCCTTGATTAGTTAAAGGGAGGAGGAATTGGGCCTATT     600
Pachon15     AGATTATTTCCCAACTGTTGCATCCTTGATTAGTTAAAGGGAGGAGGAATTGGGCCTATT     600
             ************************************************************

Rascon4      AATGTGAATTTAATGTGTGTCCACGGTCAAAATGCTGTGGATTCTGAGTTCAAAAGATTA     660
Surface      AATGTGAATTTAATGTGTGTCCACGGTCAAAATGCTGTGGATTCTGAGTTCAAAAGATTA     660
Rascon8      AATGTGAATTTAATGTGTGTCCACGGTCAAAATGCTGTGGATTCTGAGTTCAAAAGATTA     660
Rascon2      AATGTGAATTTAATGTGTGTCCACGGTCAAAATGCTGTGGATTCTGAGTTCAAAAGATTA     658
Rascon15     AATGTGAATTTAATGTGTGTCCACGGTCAAAATGCTGTGGATTCTGAGTTCAAAAGATTA     660
Rascon13     AATGTGAATTTAATGTATGTCCACGGTCAAAATGCTGTGGATTCTGAGTTCAAAAGATTA     660
Rascon6      AATGTGAATTTAATGTGTGTCCACGGTCAAAATGCTGTGGATTCTGAGTTCAAAAGATTA     660
Pachon14     AATGTGAATTTAATGTGTGTCCACGGTCAAAATGCTGTGGATTCTGAGTTCAAAAGATTA     660
Pachon9      AATGTGAATTTAATGTGTGTCCACGGTCAAAATGCTGTGGATTCTGAGTTCAAAAGATTA     660
Pachon17     AATGTGAATTTAATGTGTGTCCACGGTCAAAATGCTGTGGATTCTGAGTTCAAAAGATTA     660
Pachon12     AATGTGAATTTAATGTGTGTCCACGGTCAAAATGCTGTGGATTCTGAGTTCAAAAGATTA     660
Pachon11     AATGTGAATTTAATGTGTGTCCACGGTCAAAATGCTGTGGATTCTGAGTTCAAAAGATTA     660
Pachon7      AATGTGAATTTAATGTGTGTCCACGGTCAAAATGCTGTGGATTCTGAGTTCAAAAGATTA     660
Pachon3      AATGTGAATTTAATGTGTGTCCACGGTCAAAATGCTGTGGATTCTGAGTTCAAAAGATTA     660
Pachon8      AATGTGAATTTAATGTGTGTCCACGGTCAAAATGCTGTGGATTCTGAGTTCAAAAGATTA     660
Pachon15     AATGTGAATTTAATGTGTGTCCACGGTCAAAATGCTGTGGATTCTGAGTTCAAAAGATTA     660
             **************** *******************************************

Rascon4      AGCACTAATAGGATCAGCTGAGCTGAGACGGTCATCCCATCAGGCTAAATGCCTCGGTCA     720
Surface      AGCACTAATAGGATCAGCTGAGCTGAGACGGTCATCCCATCAGGCTAAATGCCTCGGTCA     720
Rascon8      AGCACTAATAGGATCAGCTGAGCTGAGACGGTCATCCCATCAGGCTAAATGCCTCGGTCA     720
Rascon2      AGCACTAATAGGATCAGCTGAGCTGAGACGGTCATCCCATCAGGCTAAATGCCTCGGTCA     718
Rascon15     AGCACTAATAGGATCAGCTGAGCTGAGACGGTCATCCCATCAGGCTAAATGCCTCGGTCA     720
Rascon13     AGCACTAATAGGATCAGCTGAGCTGAGACGGTCATCCCATCAGGCTAAATGCCTCGGTCA     720
Rascon6      AGCACTAATAGGATCAGCTGAGCTGAGACGGTCATCCCATCAGGCTAAATGCCTCGGTCA     720
Pachon14     AGCACTAATAGGATCAGCTGAGCTGAGACGGTCATCCCATCAGGCTAAATGCCTCGGTCA     720
Pachon9      AGCACTAATAGGATCAGCTGAGCTGAGACGGTCATCCCATCAGGCTAAATGCCTCGGTCA     720
Pachon17     AGCACTAATAGGATCAGCTGAGCTGAGACGGTCATCCCATCAGGCTAAATGCCTCGGTCA     720
Pachon12     AGCACTAATAGGATCAGCTGAGCTGAGACGGTCATCCCATCAGGCTAAATGCCTCGGTCA     720
Pachon11     AGCACTAATAGGATCAGCTGAGCTGAGACGGTCATCCCATCAGGCTAAATGCCTCGGTCA     720
Pachon7      AGCACTAATAGGATCAGCTGAGCTGAGACGGTCATCCCATCAGGCTAAATGCCTCGGTCA     720
Pachon3      AGCACTAATAGGATCAGCTGAGCTGAGACGGTCATCCCATCAGGCTAAATGCCTCGGTCA     720
Pachon8      AGCACTAATAGGATCAGCTGAGCTGAGACGGTCATCCCATCAGGCTAAATGCCTCGGTCA     720
Pachon15     AGCACTAATAGGATCAGCTGAGCTGAGACGGTCATCCCATCAGGCTAAATGCCTCGGTCA     720
             ************************************************************

Rascon4      GACTTGTGTTGGTTGTGTAGTTGAGTTAGAAAACAGGACAGACCAAGATTCATGTGCTGA     780
Surface      GACTTGTGTTGGTTGTGTAGTTGAGTTAGAAAACAGGACAGACCAAGATTCATGTGCTGA     780
Rascon8      GACTTGTGTTGGTTGTGTAGTTGAGTTAGAAAACAGGACAGACCAAGATTCATGTGCTGA     780
Rascon2      GACTTGTGTTGGTTGTGTAGTTGAGTTAGAAAACAGGACAGACCAAGATTCATGTGCTGA     778
Rascon15     GACTTGTGTTGGTTGTGTAGTTGAGTTAGAAAACAGGACAGACCAAGATTCATGTGCTGA     780
Rascon13     GACTTGTGTTGGTTGTGTAGTTGAGTTAGAAAACAGGACAGACCAAGATTCATGTGCTGA     780
Rascon6      GACTTGTGTTGGTTGTGTAGTTGAGTTAGAAAACAGGACAGACCAAGATTCATGTGCTGA     780
Pachon14     GACTTGTGTTGGTTGTGTAGTTGAGTTAGAAAACAGGACAGACCAAGATTCATGTGCTGA     780
Pachon9      GACTTGTGTTGGTTGTGTAGTTGAGTTAGAAAACAGGACAGACCAAGATTCATGTGCTGA     780
Pachon17     GACTTGTGTTGGTTGTGTAGTTGAGTTAGAAAACAGGACAGACCAAGATTCATGTGCTGA     780
```

```
Pachon12     GACTTGTGTTGGTTGTGTAGTTGAGTTAGAAAACAGGACAGACCAAGATTCATGTGCTGA     780
Pachon11     GACTTGTGTTGGTTGTGTAGTTGAGTTAGAAAACAGGACAGACCAAGATTCATGTGCTGA     780
Pachon7      GACTTGTGTTGGTTGTGTAGTTGAGTTAGAAAACAGGACAGACCAAGATTCATGTGCTGA     780
Pachon3      GACTTGTGTTGGTTGTGTAGTTGAGTTAGAAAACAGGACAGACCAAGATTCATGTGCTGA     780
Pachon8      GACTTGTGTTGGTTGTGTAGTTGAGTTAGAAAACAGGACAGACCAAGATTCATGTGCTGA     780
Pachon15     GACTTGTGTTGGTTGTGTAGTTGAGTTAGAAAACAGGACAGACCAAGATTCATGTGCTGA     780
             ************************************************************

Rascon4      CATGATGAGTATGACCTGGTCTGCATATTATACGTTTCAGGACACCATTTATAATAATGG     840
Surface      CATGATGAGTATGACCTGGTCTGCATATTATACGTTTCAGGACACCATTTATAATAATGG     840
Rascon8      CATGATGAGTATGACCTGGTCTGCATATTATACGTTTCAGGACACCATTTATAATAATGG     840
Rascon2      CATGATGAGTATGACCTGGTCTGCATATTATACGTTTCAGGGCACCATTTATAATAATGG     838
Rascon15     CATGATGAGTATGACCTGGTCTGCATATTATACGTTTCAGGACACCATTTATAATAATGG     840
Rascon13     CATGATGAGTATGACCTGGTCTGCATATTATACGTTTCAGGGCACCATTTATAATAATGG     840
Rascon6      CATGATGAGTATGACCTGGTCTGCATATTATACGTTTCAGGACACCATTTATAATAATGG     840
Pachon14     CATGATGAGTATGACCTGGTCTGCATATTATACGTTTCAGGACACCATTTATAATAATGG     840
Pachon9      CATGATGAGTATGACCTGGTCTGCATATTATACGTTTCAGGACACCATTTATAATAATGG     840
Pachon17     CATGATGAGTATGACCTGGTCTGCATATTATACGTTTCAGGACACCATTTATAATAATGG     840
Pachon12     CATGATGAGTATGACCTGGTCTGCATATTATACGTTTCAGGACACCATTTATAATAATGG     840
Pachon11     CATGATGAGTATGACCTGGTCTGCATATTATACGTTTCAGGACACCATTTATAATAATGG     840
Pachon7      CATGATGAGTATGACCTGGTCTGCATATTATACGTTTCAGGACACCATTTATAATAATGG     840
Pachon3      CATGATGAGTATGACCTGGTCTGCATATTATACGTTTCAGGACACCATTTATAATAATGG     840
Pachon8      CATGATGAGTATGACCTGGTCTGCATATTATACGTTTCAGGACACCATTTATAATAATGG     840
Pachon15     CATGATGAGTATGACCTGGTCTGCATATTATACGTTTCAGGACACCATTTATAATAATGG     840
             ***************************************** ******************

Rascon4      ACCCATTGCTGATATAGGATGTGCAAATGTGCACATAAGCCCTATCTGGACCGGATTAGT     900
Surface      ACCCATTGCTGATATAGGATGTGCAAATGTGCACATAAGCCCTATCTGGACCGGATTAGT     900
Rascon8      ACCCATTGCTGATATAGGATGTGCAAATGTGCACATAAGCCCTATCTGGACCGGATTAGT     900
Rascon2      ACCCATTGCTGATATAGGATGTGCAAATGTGCACATAAGCCCTATCTGGACCGGATTAGT     898
Rascon15     ACCCATTGCTGATATAGGATGTGCAAATGTGCACATAAGCCCTATCTGGACCGGATTAGT     900
Rascon13     ACCCATTGCTGATATAGGATGTGCAAATGTGCACATAAGCCCTATCTGGACCGGATTAGT     900
Rascon6      ACCCATTGCTGATATAGGATGTGCAAATGTGCACATAAGCCCTATCTGGACCGGATTAGT     900
Pachon14     ACCCATTGCTGATATAGGATGTGCAAATGTGCACATAAGCCCTATCTGGACCGGATTAGT     900
Pachon9      ACCCATTGCTGATATAGGATGTGCAAATGTGCACATAAGCCCTATCTGGACCGGATTAGT     900
Pachon17     ACCCATTGCTGATATAGGATGTGCAAATGTGCACATAAGCCCTATCTGGACCGGATTAGT     900
Pachon12     ACCCATTGCTGATATAGGATGTGCAAATGTGCACATAAGCCCTATCTGGACCGGATTAGT     900
Pachon11     ACCCATTGCTGATATAGGATGTGCAAATGTGCACATAAGCCCTATCTGGACCGGATTAGT     900
Pachon7      ACCCATTGCTGATATAGGATGTGCAAATGTGCACATAAGCCCTATCTGGACCGGATTAGT     900
Pachon3      ACCCATTGCTGATATAGGATGTGCAAATGTGCACATAAGCCCTATCTGGACCGGATTAGT     900
Pachon8      ACCCATTGCTGATATAGGATGTGCAAATGTGCACATAAGCCCTATCTGGACCGGATTAGT     900
Pachon15     ACCCATTGCTGATATAGGATGTGCAAATGTGCACATAAGCCCTATCTGGACCGGATTAGT     900
             ************************************************************

Rascon4      TTCTC-AGGGGGACGTCTATGAAAAATGTTTACACTTCTACTCAGTGATAAAACTCCTGC     959
Surface      TTCTCAAGGGGGACGTCTATGAAAAATGTTTACACTTCTACTCAGTGATAAAACTCCTGC     960
Rascon8      TTCTC-AGGGGGACGTCTATGAAAAATGTTTACACTTCTACTCAGTGATAAAACTCCTGC     959
Rascon2      TTCTC-AGGGGGACGTCTATGAAAAATGTTTACACTTCTACTCAGTGATAAAACTCCTGC     957
Rascon15     TTCTC-AGGGGGACGTCTATGAAAAATGTTTACACTTCTACTCAGTGATAAAACTCCTGC     959
Rascon13     TTCTC-AGGGGGACGTCTATGAAAAATGTTTACACTTCTACTCAGTGATAAAACTCCTGC     959
Rascon6      TTCTC-AGGGGGACGTCTATGAAAAATGTTTACACTTCTACTCAGTGATAAAACTCCTGC     959
Pachon14     TTCTC-AGGGGGACGTCTATGAAAAATGTTTACACTTCTACTCAGTGATAAAACTCCTGC     959
Pachon9      TTCTC-AGGGGGACGTCTATGAAAAATGTTTACACTTCTACTCAGTGATAAAACTCCTGC     959
Pachon17     TTCTC-AGGGGGACGTCTATGAAAAATGTTTACACTTCTACTCAGTGATAAAACTCCTGC     959
Pachon12     TTCTC-AGGGGGACGTCTATGAAAAATGTTTACACTTCTACTCAGTGATAAAACTCCTGC     959
Pachon11     TTCTC-AGGGGGACGTCTATGAAAAATGTTTACACTTCTACTCAGTGATAAAACTCCTGC     959
Pachon7      TTCTC-AGGGGGACGTCTATGAAAAATGTTTACACTTCTACTCAGTGATAAAACTCCTGC     959
Pachon3      TTCTC-AGGGGGACGTCTATGAAAAATGTTTACACTTCTACTCAGTGATAAAACTCCTGC     959
Pachon8      TTCTC-AGGGGGACGTCTATGAAAAATGTTTACACTTCTACTCAGTGATAAAACTCCTGC     959
Pachon15     TTCTC-AGGGGGACGTCTATGAAAAATGTTTACACTTCTACTCAGTGATAAAACTCCTGC     959
             ***** ******************************************************

Rascon4      ATCTGGACTGCAGTTGAAAAAAACAGGAGGACCAGTGAGTTGTTTGGCTTTTTCAGTCAC     1019
Surface      ATCTGGACTGCAGTTGAAAAAAACAGGAGGACCAGTGAGTTGTTTGGCTTTTTCAGTCAC     1020
Rascon8      ATCTGGACTGCAGTTGAAAAAAACAGGAGGACCAGTGAGTTGTTTGGCTTTTTCAGTCAC     1019
Rascon2      ATCTGGACTGCAGTTGAAAAAAACAGGAGGACCAGTGAGTTGTTTGGCTTTTTCAGTCAC     1017
Rascon15     ATCTGGACTGCAGTTGAAAAAAACAGGAGGACCAGTGAGTTGTTTGGCTTTTTCAGTCAC     1019
Rascon13     ATCTGGACTGCAGTTGAAAAAAACAGGAGGACCAGTGAGTTGTTTGGCTTTTTCAGTCAC     1019
Rascon6      ATCTGGACTGCAGTTGAAAAAAACAGGAGGACCAGTGAGTTGTTTGGCTTTTTCAGTCAC     1019
Pachon14     ATCTGGACTGCAGTTGAAAAAAACAGGAGGACCAGTGAGTTGTTTGGCTTTTTCAGTCAC     1019
Pachon9      ATCTGGACTGCAGTTGAAAAAAACAGGAGGACCAGTGAGTTGTTTGGCTTTTTCAGTCAC     1019
Pachon17     ATCTGGACTGCAGTTGAAAAAAACAGGAGGACCAGTGAGTTGTTTGGCTTTTTCAGTCAC     1019
Pachon12     ATCTGGACTGCAGTTGAAAAAAACAGGAGGACCAGTGAGTTGTTTGGCTTTTTCAGTCAC     1019
Pachon11     ATCTGGACTGCAGTTGAAAAAAACAGGAGGACCAGTGAGTTGTTTGGCTTTTTCAGTCAC     1019
Pachon7      ATCTGGACTGCAGTTGAAAAAAACAGGAGGACCAGTGAGTTGTTTGGCTTTTTCAGTCAC     1019
Pachon3      ATCTGGACTGCAGTTGAAAAAAACAGGAGGACCAGTGAGTTGTTTGGCTTTTTCAGTCAC     1019
Pachon8      ATCTGGACTGCAGTTGAAAAAAACAGGAGGACCAGTGAGTTGTTTGGCTTTTTCAGTCAC     1019
```

```
Pachon15        ATCTGGACTGCAGTTGAAAAAAACAGGAGGACCAGTGAGTTGTTTGGCTTTTTCAGTCAC    1019
                ************************************************************

Rascon4         GACATCATGTTCAGTAGCTCCTCCATTTCCACTCGCTTTTGTGTTTACATGGATCTCAGC    1079
Surface         GACATCATGTTCAGTAGCTCCTCCATTTCCACTCGCTTTTGTGTTTACATGGATCTCAGC    1080
Rascon8         GACATCATGTTCAGTAGCTCCTCCATTTCCTCTCGCTTTTGTGTTTACATGGATCTCAGC    1079
Rascon2         GACATCATGTTCAGTAGCTCCTCCATTTCCACTCGCTTTTGTGTTTACATGGATCTCAGC    1077
Rascon15        GACATCATGTTCAGTAGCTCCTCCATTTCCACTCGCTTTTGTGTTTACATGGATCTCAGC    1079
Rascon13        GACATCATGTTCAGTAGCTCCTCCATTTCCACTCGCTTTTGTGTTTACATGGATCTCAGC    1079
Rascon6         GACATCATGTTCAGTAGCTCCTCCATTTCCACTCGCTTTTGTGTTTACATGGATCTCAGC    1079
Pachon14        GACATCATGTTCAGTAGCTCCTCCATTTCCACTCGCTTTTGTGTTTACATGGATCTCAGC    1079
Pachon9         GACATCATGTTCAGTAGCTCCTCCATTTCCACTCGCTTTTGTGTTTACATGGATCTCAGC    1079
Pachon17        GACATCATGTTCAGTAGCTCCTCCATTTCCACTCGCTTTTGTGTTTACATGGATCTCAGC    1079
Pachon12        GACATCATGTTCAGTAGCTCCTCCATTTCCACTCGCTTTTGTGTTTACATGGATCTCAGC    1079
Pachon11        GACATCATGTTCAGTAGCTCCTCCATTTCCACTCGCTTTTGTGTTTACATGGATCTCAGC    1079
Pachon7         GACATCATGTTCAGTAGCTCCTCCATTTCCACTCGCTTTTGTGTTTACATGGATCTCAGC    1079
Pachon3         GACATCATGTTCAGTAGCTCCTCCATTTCCACTCGCTTTTGTGTTTACATGGATCTCAGC    1079
Pachon8         GACATCATGTTCAGTAGCTCCTCCATTTCCACTCGCTTTTGTGTTTACATGGATCTCAGC    1079
Pachon15        GACATCATGTTCAGTAGCTCCTCCATTTCCACTCGCTTTTGTGTTTACATGGATCTCAGC    1079
                ****************************** *****************************

Rascon4         AGAAATGTGTAATTACGGTGTGTTGCAGTGGACAAATAGCTATTTTATTAAACACAAGGA    1139
Surface         AGAAATGTGTAATTACGGTGTGTTGCAGTGGACAAATAGCTATTTTATTAAACACAAGGA    1140
Rascon8         AGAAATGTGTAATTACGGTGTGTTGCAGTGGACAAATAGCTATTTTATTAAACACAAGGA    1139
Rascon2         AGAAATGTGTAATTACGGTGTGTTGCAGTGGACAAATAGCTATTTTATTAAACACAAGGA    1137
Rascon15        AGAAATGTGTAATTACGGTGTGTTGCAGTGGACAAATAGCTATTTTATTAAACACAAGGA    1139
Rascon13        AGAAATGTGTAATTACGGTGTGTTGCAGTGGACAAATAGCTATTTTATTAAACACAAGGA    1139
Rascon6         AAAAATGTGTAATTACGGTGTGTTGCAGTGGACAAATAGCTATTTTATTAAACACAAGGA    1139
Pachon14        AGAAATGTGTAATTACGGTGTGTTGCAGTGGACAAATAGCTATTTTATTAAACACAAGGA    1139
Pachon9         AGAAATGTGTAATTACGGTGTGTTGCAGTGGACAAATAGCTATTTTATTAAACACAAGGA    1139
Pachon17        AGAAATGTGTAATTACGGTGTGTTGCAGTGGACAAATAGCTATTTTATTAAACACAAGGA    1139
Pachon12        AGAAATGTGTAATTACGGTGTGTTGCAGTGGACAAATAGCTATTTTATTAAACACAAGGA    1139
Pachon11        AGAAATGTGTAATTACGGTGTGTTGCAGTGGACAAATAGCTATTTTATTAAACACAAGGA    1139
Pachon7         AGAAATGTGTAATTACGGTGTGTTGCAGTGGACAAATAGCTATTTTATTAAACACAAGGA    1139
Pachon3         AGAAATGTGTAATTACGGTGTGTTGCAGTGGACAAATAGCTATTTTATTAAACACAAGGA    1139
Pachon8         AGAAATGTGTAATTACGGTGTGTTGCAGTGGACAAATAGCTATTTTATTAAACACAAGGA    1139
Pachon15        AGAAATGTGTAATTACGGTGTGTTGCAGTGGACAAATAGCTATTTTATTAAACACAAGGA    1139
                * **********************************************************

Rascon4         GACGAAACTCAGAGTTGTGCATCCGGACAGGACTAAAATTACAGAAGGACTACAGAGTTT    1199
Surface         GACGAAACTCAGAGTTGTGCATCCGGACAGGACTAAAATTACAGAAGGACCACAGAGTTT    1200
Rascon8         GACGAAACTCAGAGTTGTGCATCCGGACAGGACTAAAATTACAGAAGGACCACAGAGTTT    1199
Rascon2         GACGAAACTCAGAGTTGTGCATCCGGACAGGACTAAAATTACAGAAGGACCACAGAGTTT    1197
Rascon15        GACGAAACTCAGAGTTGTGCATCCGGACAGGACTAAAATTACAGAAGGACCACAGAGTTT    1199
Rascon13        GACGAAACTCAGAGTTGTGCATCCGGACAGGACTAAAATTACAGAAGGACCACAGAGTTT    1199
Rascon6         GACGAAACTCAGAGTTGTGCATCCGGACAGGACTAAAATTACAGAAGGACCACAGAGTTT    1199
Pachon14        GACGAAACTCAGAGTTGTGCATCCGGACAGGACTAAAATTACAGAAGGACCACAGAGTTT    1199
Pachon9         GACGAAACTCAGAGTTGTGCATCCGGACAGGACTAAAATTACAGAAGGACCACAGAGTTT    1199
Pachon17        GACGAAACTCAGAGTTGTGCATCCGGACAGGACTAAAATTACAGAAGGACCACAGAGTTT    1199
Pachon12        GACGAAACTCAGAGTTGTGCATCCGGACAGGACTAAAATTACAGAAGGACCACAGAGTTT    1199
Pachon11        GACGAAACTCAGAGTTGTGCATCCGGACAGGACTAAAATTACAGAAGGACCACAGAGTTT    1199
Pachon7         GACGAAACTCAGAGTTGTGCATCCGGACAGGACTAAAATTACAGAAGGACCACAGAGTTT    1199
Pachon3         GACGAAACTCAGAGTTGTGCATCCGGACAGGACTAAAATTACAGAAGGACCACAGAGTTT    1199
Pachon8         GACGAAACTCAGAGTTGTGCATCCGGACAGGACTAAAATTACAGAAGGACCACAGAGTTT    1199
Pachon15        GACGAAACTCAGAGTTGTGCATCCGGACAGGACTAAAATTACAGAAGGACCACAGAGTTT    1199
                ************************************************** *********

Rascon4         TAGCGAAAAACAGTAGGTAATTCAGCCTGTAATTTTCTCAGAGGACGTCTAAGAAAAACA    1259
Surface         TAGCGAAAAACAGTAGGTAATTCAGCCTGTAATTTTCTCAGAGGACGTCTAAGAAAAACA    1260
Rascon8         TAGCGAAAAACAGTAGGTAATTCAGCCTGTAATTTTCTCAGAGGACGTCTAAGAAAAACA    1259
Rascon2         TAGCGAAAAACAGTAGGTAATTCAGCCTGTAATTTTCTCAGAGGACGTCTAAGAAAAACA    1257
Rascon15        TAGCGAAAAACAGTAGGTAATTCAGCCTGTAATTTTCTCAGAGGACGTCTAAGAAAAACA    1259
Rascon13        TAGCGAAAAACAGTAGGTAATTCAGCCTGTAATTTTCTCAGAGGACGTCTAAGAAAAACA    1259
Rascon6         TAGCGAAAAACAGTAGGTAATTCAGCCTGTAATTTTCTCAGAGGACGTCTAAAAAAAACA    1259
Pachon14        TAGCGAAAAACAGTAGGTAATTCAGCCTGTAATTTTCTCAGAGGACGTCTAAGAAAAACA    1259
Pachon9         TAGCGAAAAACAGTAGGTAATTCAGCCTGTAATTTTCTCAGAGGACGTCTAAGAAAAACA    1259
Pachon17        TAGCGAAAAACAGTAGGTAATTCAGCCTGTAATTTTCTCAGAGGACGTCTAAGAAAAACA    1259
Pachon12        TAGCGAAAAACAGTAGGTAATTCAGCCTGTAATTTTCTCAGAGGACGTCTAAGAAAAACA    1259
Pachon11        TAGCGAAAAACAGTAGGTAATTCAGCCTGTAATTTTCTCAGAGGACGTCTAAGAAAAACA    1259
Pachon7         TAGCGAAAAACAGTAGGTAATTCAGCCTGTAATTTTCTCAGAGGACGTCTAAGAAAAACA    1259
Pachon3         TAGCGAAAAACAGTAGGTAATTCAGCCTGTAATTTTCTCAGAGGACGTCTAAGAAAAACA    1259
Pachon8         TAGCGAAAAACAGTAGGTAATTCAGCCTGTAATTTTCTCAGAGGACGTCTAAGAAAAACA    1259
Pachon15        TAGCGAAAAACAGTAGGTAATTCAGCCTGTAATTTTCTCAGAGGACGTCTAAGAAAAACA    1259
                **************************************************** *******

Rascon4         TGTATATGGTCGTTCCGGATGAGAATAAAATCACAGAGGA-CCCCCCATAAAAGAGAACC    1318
Surface         TGTATATGGTCGTTCCGGATGAGAATAAAATCACAGAGGA-CCCCCCATAAAAGAGAACC    1319
```

```
Rascon8         TGTATATGGTCGTTCCGGATGAGAATAAAATCACAGAGGA-CCCCCCATAAAAGAGAACC        1318
Rascon2         TGTATATGGTCGTTCCGGATGAGAATAAAATCACAGAGGACCCCCCCATAAAAGAGAACC        1317
Rascon15        TGTATATGGTCGTTCCGGATGCGAATAAAATCACAGAAGA-CCCCCCATAAAAGAGAACC        1318
Rascon13        TGTATATGGTCGTTCCGGATGAGAATAAAATCACAGAGGA-CCCCCCATAAAAGAGAACC        1318
Rascon6         TGTATATGGTCGTTCCGGATGAGAATAAAATCACAGAGGA-CCCCCCATAAAAGAGAACC        1318
Pachon14        TGTATATGGTCGTTCCGGATGAGAATAAAATCACAGAGGA-CCCCCCATAAAAGAGAACC        1318
Pachon9         TGTATATGGTCGTTCCGGATGAGAATAAAATCACAGAGGA-CCCCCCATAAAAGAGAACC        1318
Pachon17        TGTATATGGTCGTTCCGGATGAGAATAAAATCACAGAGGA-CCCCCCATAAAAGAGAACC        1318
Pachon12        TGTATATGGTCGTTCCGGATGAGAATAAAATCACAGAGGA-CCCCCCATAAAAGAGAACC        1318
Pachon11        TGTATATGGTCGTTCCGGATGAGAATAAAATCACAGAGGA-CCCCCCATAAAAGAGAACC        1318
Pachon7         TGTATATGGTCGTTCCGGATGAGAATAAAATCACAGAGGA-CCCCCCATAAAAGAGAACC        1318
Pachon3         TGTATATGGTCGTTCCGGATGAGAATAAAATCACAGAGGA-CCCCCCATAAAAGAGAACC        1318
Pachon8         TGTATATGGTCGTTCCGGATGAGAATAAAATCACAGAGGA-CCCCCCATAAAAGAGAACC        1318
Pachon15        TGTATATGGTCGTTCCGGATGAGAATAAAATCACAGAGGA-CCCCCCATAAAAGAGAACC        1318
                ********************* *************** ** *******************

Rascon4         CAGAACCCAGGAGAAACTAATCCTGTCCAGATAGGGCTTAAATCTAATATATTTTCTCAT        1378
Surface         CAGAACCCAGGAGAAACTAATCCTGTCCAGATAGGGCTTAAATCTAATATATTTTCTCAT        1379
Rascon8         CAGAACCCAGGAGAAACTAATTCTGTCCAGATAGGGCTTAAATCTAATATATTTTCTCAT        1378
Rascon2         CAGAACCCAGGAGAAACTAATCCTGTCCAGATAGGGCTTAAATCTAATATATTTTCTCAT        1377
Rascon15        CAGAACCCAGGAGAAACTAATCCTGTCCAGATAGGGCTTAAATCTAATATATTTTCTCAT        1378
Rascon13        CAGAACCCAGGAGAAACTAATCCTGTCCAGATAGGGCTTAAATCTAATATATTTTCTCAT        1378
Rascon6         CAGAACCCAGGAGAAACTAATCCTGTCCAGATAGGGCTTAAATCTAATATATTTTCTCAT        1378
Pachon14        CAGAACCCAGGAGAAACTAATCCTGTCCAGATAGGGCTTAAATCTAATATATTTTCTCAT        1378
Pachon9         CAGAACCCAGGAGAAACTAATCCTGTCCAGATAGGGCTTAAATCTAATATATTTTCTCAT        1378
Pachon17        CAGAACCCAGGAGAAACTAATCCTGTCCAGATAGGGCTTAAATCTAATATATTTTCTCAT        1378
Pachon12        CAGAACCCAGGAGAAACTAATCCTGTCCAGATAGGGCTTAAATCTAATATATTTTCTCAT        1378
Pachon11        CAGAACCCAGGAGAAACTAATCCTGTCCAGATAGGGCTTAAATCTAATATATTTTCTCAT        1378
Pachon7         CAGAACCCAGGAGAAACTAATCCTGTCCAGATAGGGCTTAAATCTAATATATTTTCTCAT        1378
Pachon3         CAGAACCCAGGAGAAACTAATCCTGTCCAGATAGGGCTTAAATCTAATATATTTTCTCAT        1378
Pachon8         CAGAACCCAGGAGAAACTAATCCTGTCCAGATAGGGCTTAAATCTAATATATTTTCTCAT        1378
Pachon15        CAGAACCCAGGAGAAACTAATCCTGTCCAGATAGGGCTTAAATCTAATATATTTTCTCAT        1378
                ********************* **************************************

Rascon4         TTAACAAAAAGTATGTATTATGCTACAACTTGTTTTATTTCAGTTATAGATATTCCACCT        1438
Surface         TTAACAAAAAGTATGTATTATGCTACAACTTGTTTTATTTCAGTTATAGATATTCCACCT        1439
Rascon8         TTAACAAAAAGTATGTATTATGCTACAACTTGTTTTATTTCAGTTATAGATATTCCACCT        1438
Rascon2         TTAACAAAAAGTATGTATTATGCTACAACTTGTTTTATTTCAGTTATAGATATTCCACCT        1437
Rascon15        TTAACAAAAAGTATGTATTATGCTACAACTTGTTTTATTTCAGTTATAGATATTCCACCT        1438
Rascon13        TTAACAAAAAGTATGTATTATGCTACAACTTGTTTTATTTCAGTTATAGATATTCCACCT        1438
Rascon6         TTAACAAAAAGTATGTATTATGCTACAAATTGTTTTATTTCAGTTATAGATATTCCACCT        1438
Pachon14        TTAACAAAAAGTATGTATTATGCTACAACTTGTTTTATTTCAGTTATAGATATTCCACCT        1438
Pachon9         TTAACAAAAAGTATGTATTATGCTACAACTTGTTTTATTTCAGTTATAGATATTCCACCT        1438
Pachon17        TTAACAAAAAGTATGTATTATGCTACAACTTGTTTTATTTCAGTTATAGATATTCCACCT        1438
Pachon12        TTAACAAAAAGTATGTATTATGCTACAACTTGTTTTATTTCAGTTATAGATATTCCACCT        1438
Pachon11        TTAACAAAAAGTATGTATTATGCTACAACTTGTTTTATTTCAGTTATAGATATTCCACCT        1438
Pachon7         TTAACAAAAAGTATGTATTATGCTACAACTTGTTTTATTTCAGTTATAGATATTCCACCT        1438
Pachon3         TTAACAAAAAGTATGTATTATGCTACAACTTGTTTTATTTCAGTTATAGATATTCCACCT        1438
Pachon8         TTAACAAAAAGTATGTATTATGCTACAACTTGTTTTATTTCAGTTATAGATATTCCACCT        1438
Pachon15        TTAACAAAAAGTATGTATTATGCTACAACTTGTTTTATTTCAGTTATAGATATTCCACCT        1438
                **************************** *******************************

Rascon4         TAAATTCTTTATCTGCCTGTGAATTTCAAACATGTAATGAATCTTTGACACTTAATTCTA        1498
Surface         TAAATTCTTTATCTGCCTGTGAATTTCAAACATGTAATGAATCTTTGACACTTAATTCTA        1499
Rascon8         TAAATTCTTTATCTGCCTGTGAATTTCAAACATGTAATGAATCTTTGACACTTAATTCTA        1498
Rascon2         TAAATTCTTTATCTGCCTGTGAATTTCAAACATGTAATGAATCTCTGACACTTAATTCTA        1497
Rascon15        TAAATTCTTTATCTGCCTGTGAATTTCAAACATGTAATGAATCTTTGACACTTAATTCTA        1498
Rascon13        TAAATTCTTTATCTGCCTGTGAATTTCAAACAAGTAATGAATCTTTGACACTTAATTCTA        1498
Rascon6         TAAATTCTTTATCTGCCTGTGAATTTCAAACATGTAATGAATCTTTGACACTTAATTCTA        1498
Pachon14        TAAATTCTTTATCTGCCTGTGAATTTCAAACATGTAATGAATCTTTGACACTTAATTCTA        1498
Pachon9         TAAATTCTTTATCTGCCTGTGAATTTCAAACATGTAATGAATCTTTGACACTTAATTCTA        1498
Pachon17        TAAATTCTTTATCTGCCTGTGAATTTCAAACATGTAATGAATCTTTGACACTTAATTCTA        1498
Pachon12        TAAATTCTTTATCTGCCTGTGAATTTCAAACATGTAATGAATCTTTGACACTTAATTCTA        1498
Pachon11        TAAATTCTTTATCTGCCTGTGAATTTCAAACATGTAATGAATCTTTGACACTTAATTCTA        1498
Pachon7         TAAATTCTTTATCTGCCTGTGAATTTCAAACATGTAATGAATCTTTGACACTTAATTCTA        1498
Pachon3         TAAATTCTTTATCTGCCTGTGAATTTCAAACATGTAATGAATCTTTGACACTTAATTCTA        1498
Pachon8         TAAATTCTTTATCTGCCTGTGAATTTCAAACATGTAATGAATCTTTGACACTTAATTCTA        1498
Pachon15        TAAATTCTTTATCTGCCTGTGAATTTCAAACATGTAATGAATCTTTGACACTTAATTCTA        1498
                ******************************** *********** ***************

Rascon4         TAATGAGCAGTCTCGACACAGTATAAAGATGTAATGGGTGATTACACTGCTATCATTTTA        1558
Surface         TAATGAGCAGTCTCGACACAGTATAAAGATGTAATGGGTGATTACACTGCTATCATTTTA        1559
Rascon8         TAATGAGCAGTCTCGACACAGTATAAAGATGTAATGGGTGATTACACTGCTATCATTTTA        1558
Rascon2         TAATGAGCAGTCTCGACACAGTATAAAGATGTAATGGGTGATTACACTGCTATCATTTTA        1557
Rascon15        TAATGAGCAGTCTCGACACAGTATAAAGATGTAATGGGTGATTACACTGCTATCATTTTA        1558
Rascon13        TAATGAGCAGTCTCGACACAGTATAAAGATGTAATGGGTGATTACACTGCTATCATTTTA        1558
Rascon6         TAATGAGCAGTCTCGACACAGTATAAAGATGTAATGGGTGATTACACTGCTATCATTTTA        1558
```

```
Pachon14        TAATGAGCAGTCTCGACACAGTATAAAGATGTAATGGGTGATTACACTGCTATCATTTTA        1558
Pachon9         TAATGAGCAGTCTCGACACAGTATAAAGATGTAATGGGTGATTACACTGCTATCATTTTA        1558
Pachon17        TAATGAGCAGTCTCGACACAGTATAAAGATGTAATGGGTGATTACACTGCTATCATTTTA        1558
Pachon12        TAATGAGCAGTCTCGACACAGTATAAAGATGTAATGGGTGATTACACTGCTATCATTTTA        1558
Pachon11        TAATGAGCAGTCTCGACACAGTATAAAGATGTAATGGGTGATTACACTGCTATCATTTTA        1558
Pachon7         TAATGAGCAGTCTCGACACAGTATAAAGATGTAATGGGTGATTACACTGCTATCATTTTA        1558
Pachon3         TAATGAGCAGTCTCGACACAGTATAAAGATGTAATGGGTGATTACACTGCTATCATTTTA        1558
Pachon8         TAATGAGCAGTCTCGACACAGTATAAAGATGTAATGGGTGATTACACTGCTATCATTTTA        1558
Pachon15        TAATGAGCAGTCTCGACACAGTATAAAGATGTAATGGGTGATTACACTGCTATCATTTTA        1558
                ************************************************************

Rascon4         ACTACATTTCAACTAAATGCACATTATGTCTAAAGGTTTGGACACATCTCTTCTTATTTG        1618
Surface         ACTACATTTCAACTAAATGCACATTATGTCTAAAGGTTTGGACACATCTCTTCTTATTTG        1619
Rascon8         ACTACATTTCAACTAAATGCACATTATGTCTAAAGGTTTGGACACATCTCTTCTTATTTG        1618
Rascon2         ACTACATTTCAACTAAATGCACATTATGTCTAAAGGTTTGGACACATCTCTTCTTATTTG        1617
Rascon15        ACTACATTTCAACTAAATGCACATTATGTCTAAAGGTTTGGACACATCTCTTCTTATTTG        1618
Rascon13        ACTACATTTCAACTAAATGCACATTATGTCTAAAGGTTTGGACACATCTCTTCTTATTTG        1618
Rascon6         ACTACATTTCAACTAAATGCACATTATGTCTAAAGGTTTGGACACATCTCTTCTTATTTG        1618
Pachon14        ACTACATTTCAACTAAATGCACATTATGTCTAAAGGTTTGGACACATCTCTTCTTATTTG        1618
Pachon9         ACTACATTTCAACTAAATGCACATTATGTCTAAAGGTTTGGACACATCTCTTCTTATTTG        1618
Pachon17        ACTACATTTCAACTAAATGCACATTATGTCTAAAGGTTTGGACACATCTCTTCTTATTTG        1618
Pachon12        ACTACATTTCAACTAAATGCACATTATGTCTAAAGGTTTGGACACATCTCTTCTTATTTG        1618
Pachon11        ACTACATTTCAACTAAATGCACATTATGTCTAAAGGTTTGGACACATCTCTTCTTATTTG        1618
Pachon7         ACTACATTTCAACTAAATGCACATTATGTCTAAAGGTTTGGACACATCTCTTCTTATTTG        1618
Pachon3         ACTACATTTCAACTAAATGCACATTATGTCTAAAGGTTTGGACACATCTCTTCTTATTTG        1618
Pachon8         ACTACATTTCAACTAAATGCACATTATGTCTAAAGGTTTGGACACATCTCTTCTTATTTG        1618
Pachon15        ACTACATTTCAACTAAATGCACATTATGTCTAAAGGTTTGGACACATCTCTTCTTATTTG        1618
                ************************************************************

Rascon4         ATGTGTTTTCTTTCTTTTTTATAATTTTCTATATTGTACATTTAATTTTAGTAGCAAAAT        1678
Surface         ATGTGTTTTCTTTCTTTTTTATAATTTTCTATATTGTAAATTTAATTTTAGTAGCAAAAT        1679
Rascon8         ATGTGTTTTCTTTCTTTTTTATAATTTTCTATATTGTAAATTTAATTTTAGTAGCAAAAT        1678
Rascon2         ATGTGTTTTCTTTCTTTTTTATAATTTTCTATATTGTAAATTTAATTTTAGTAGCAAAAT        1677
Rascon15        ATGTGTTTTCTTTCTTTTTTATAATTTTCTATATTGTAAATTTAATTTTAGTAGCAAAAT        1678
Rascon13        ATGTGTTTTCTTTCTTTTTTATAATTTTCTATATTGTAAATTTAATTTTAGTAGCAAAAT        1678
Rascon6         ATGTGTTTTCTTTCTTTTTTATAATTTTCTATATTGTACATTTAATTTTAGTAGCAAAAT        1678
Pachon14        ATGTGTTTTCTTTCTTTTTTATAATTTTCTATATTGTAAATTTAATTTTAGTAGCAAAAT        1678
Pachon9         ATGTGTTTTCTTTCTTTTTTATAATTTTCTATATTGTAAATTTAATTTTAGTAGCAAAAT        1678
Pachon17        ATGTGTTTTCTTTCTTTTTTATAATTTTCTATATTGTAAATTTAATTTTAGTAGCAAAAT        1678
Pachon12        ATGTGTTTTCTTTCTTTTTTATAATTTTCTATATTGTAAATTTAATTTTAGTAGCAAAAT        1678
Pachon11        ATGTGTTTTCTTTCTTTTTTATAATTTTCTATATTGTAAATTTAATTTTAGTAGCAAAAT        1678
Pachon7         ATGTGTTTTCTTTCTTTTTTATAATTTTCTATATTGTAAATTTAATTTTAGTAGCAAAAT        1678
Pachon3         ATGTGTTTTCTTTCTTTTTTATAATTTTCTATATTGTAAATTTAATTTTAGTAGCAAAAT        1678
Pachon8         ATGTGTTTTCTTTCTTTTTTATAATTTTCTATATTGTAAATTTAATTTTAGTAGCAAAAT        1678
Pachon15        ATGTGTTTTCTTTCTTTTTTATAATTTTCTATATTGTAAATTTAATTTTAGTAGCAAAAT        1678
                ************************************** *********************

Rascon4         AAGGATAGCAGAATAATGACCTAATTAGCATTATTACTATACTATTATATCACTATAGTA        1738
Surface         AAGGATAGCAGAATAATGACCTAATTAGCATTATTACTATACTATTATATCACTATAGTA        1739
Rascon8         AAGGATAGCAGAATAATGACCTAATTAGCATTATTACTATACTATTATATCACTATAGTA        1738
Rascon2         AAGGATAGCAGAATAATGACCTAATTAGCATTATTACTATACTATTATATCACTATAATA        1737
Rascon15        AAGGATAGCAGAATAATGACCTAATTAGCATTATTACTATACTATTATATCACTATAGTA        1738
Rascon13        AAGGATAGCAGAATAATGACCTAATTAGCATTATTACTATACTATTATATCACTATAGTA        1738
Rascon6         AAGGATAGCAGAATAATGACCTAATTAGCATTATTACTATACTATTATATCACTATAGTA        1738
Pachon14        AAGGATAGCAGAATAATGACCTAATTAGCATTATTACTATACTATTATATCACTATAGTA        1738
Pachon9         AAGGATAGCAGAATAATGACCTAATTAGCATTATTACTATACTATTATATCACTATAGTA        1738
Pachon17        AAGGATAGCAGAATAATGACCTAATTAGCATTATTACTATACTATTATATCACTATAGTA        1738
Pachon12        AAGGATAGCAGAATAATGACCTAATTAGCATTATTACTATACTATTATATCACTATAGTA        1738
Pachon11        AAGGATAGCAGAATAATGACCTAATTAGCATTATTACTATACTATTATATCACTATAGTA        1738
Pachon7         AAGGATAGCAGAATAATGACCTAATTAGCATTATTACTATACTATTATATCACTATAGTA        1738
Pachon3         AAGGATAGCAGAATAATGACCTAATTAGCATTATTACTATACTATTATATCACTATAGTA        1738
Pachon8         AAGGATAGCAGAATAATGACCTAATTAGCATTATTACTATACTATTATATCACTATAGTA        1738
Pachon15        AAGGATAGCAGAATAATGACCTAATTAGCATTATTACTATACTATTATATCACTATAGTA        1738
                ********************************************************* **

Rascon4         TATTAGGTACCACTTTAAAATAAGACTACCTTTATAAAGAGTTTATAAATATTTACAAAA        1798
Surface         TATTAGGTACCACTTTAAAATAAGACTACCTTTATAAAGAGTTTATAAATATTTACAAAA        1799
Rascon8         TATTAGGTACCACTTTAAAATAAGACTACCTTTATAAAGAGTTTATAAATATTTACAAAA        1798
Rascon2         TATTAGGTACCACTTTAAAATAAGACTACCTTTATAAAGAGTTTATAAATATTTACAAAA        1797
Rascon15        TATTAGGTACCACTTTAAAATAAGACTACCTTTATAAAGAGTTTATAAATATTTACAAAA        1798
Rascon13        TATTAGGTACCACTTTAAAATAAGAGTACCTTTATAAAGAGTTTATAAATATTTACAAAA        1798
Rascon6         TATTAGGTACCACTTTAAAATAAGACTACCTTTATAAAGAGTTTATAAATATTTACAAAA        1798
Pachon14        TATTAGGTACCACTTTAAAATAAGACTACCTTTATAAAGAGTTTATAAATATTTACAAAA        1798
Pachon9         TATTAGGTACCACTTTAAAATAAGACTACCTTTATAAAGAGTTTATAAATATTTACAAAA        1798
Pachon17        TATTAGGTACCACTTTAAAATAAGACTACCTTTATAAAGAGTTTATAAATATTTACAAAA        1798
Pachon12        TATTAGGTACCACTTTAAAATAAGACTACCTTTATAAAGAGTTTATAAATATTTACAAAA        1798
Pachon11        TATTAGGTACCACTTTAAAATAAGACTACCTTTATAAAGAGTTTATAAATATTTACAAAA        1798
```

```
Pachon7         TATTAGGTACCACTTTAAAATAAGACTACCTTTATAAAGAGTTTATAAATATTTACAAAA      1798
Pachon3         TATTAGGTACCACTTTAAAATAAGACTACCTTTATAAAGAGTTTATAAATATTTACAAAA      1798
Pachon8         TATTAGGTACCACTTTAAAATAAGACTACCTTTATAAAGAGTTTATAAATATTTACAAAA      1798
Pachon15        TATTAGGTACCACTTTAAAATAAGACTACCTTTATAAAGAGTTTATAAATATTTACAAAA      1798
                ************************* **********************************

Rascon4         ACCATTAATAAACTCATTGTAAACCATTTATAAACCCTTTATAAAGGTAGTCTTATTTTA      1858
Surface         ACCATTAATAAACTCATTGTAAACCATTTATAAACCCTTTATAAAGGTAGACTTATTTTA      1859
Rascon8         ACCATTAATAAACTCATTGTAAACCATTTATAAACCCTTTATAAAGGTAGTCTTATTTTA      1858
Rascon2         ACCATTAATAAACTCATTGTAAACCATTTATAAACCCTTTATAAAGGTAGTCTTATTTTA      1857
Rascon15        ACCATTAATAAACTCATTGTAAACCATTTATAAACCCTTTATAAAGGTAGTCTTATTTTA      1858
Rascon13        ACCATTAATAAACTCATTGTAAACCATTTATAAACCCTTTATAAAGGTAGTCTTATTTTA      1858
Rascon6         ACCATTAATAAACTCATTGTAAACCATTTATAAACCCTTTATAAAGGTAGTCTTATTTTA      1858
Pachon14        ACCATTAATAAACTCATTGTAAACCATTTATAAACCCTTTATAAAGGTAGTCTTATTTTA      1858
Pachon9         ACCATTAATAAACTCATTGTAAACCATTTATAAACCCTTTATAAAGGTAGTCTTATTTTA      1858
Pachon17        ACCATTAATAAACTCATTGTAAACCATTTATAAACCCTTTATAAAGGTAGTCTTATTTTA      1858
Pachon12        ACCATTAATAAACTCATTGTAAACCATTTATAAACCCTTTATAAAGGTAGTCTTATTTTA      1858
Pachon11        ACCATTAATAAACTCATTGTAAACCATTTATAAACCCTTTATAAAGGTAGTCTTATTTTA      1858
Pachon7         ACCATTAATAAACTCATTGTAAACCATTTATAAACCCTTTATAAAGGTAGTCTTATTTTA      1858
Pachon3         ACCATTAATAAACTCATTGTAAACCATTTATAAACCCTTTATAAAGGTAGTCTTATTTTA      1858
Pachon8         ACCATTAATAAACTCATTGTAAACCATTTATAAACCCTTTATAAAGGTAGTCTTATTTTA      1858
Pachon15        ACCATTAATAAACTCATTGTAAACCATTTATAAACCCTTTATAAAGGTAGTCTTATTTTA      1858
                ************************************************** *********

Rascon4         AAGTGGTACCGTATATTATATATACACTATAGTATTATTTCATCTTATTTTCATTCTGGC      1918
Surface         AAGTGGTACCGTATATTATATATACACTATAGTATTATTTCATCTTATTTTCATTCTGGC      1919
Rascon8         AAGTGGTACCGTATATTATATATACACTATAGTATTATTTCATCTTATTTTCATTCTGGC      1918
Rascon2         AAGTGGTACCGTATATTATATATACACTATAGTATTATTTCATCTTATTTTCATTCTGGC      1917
Rascon15        AAGTGGTACCGTATATTATATATACACTATAGTATTATTTCATCTTATTTTCATTCTGGC      1918
Rascon13        AAGTGGTACCGTATATTATATATACACTATAGTATTATTTCATCTTATTTTCATTCTGGC      1918
Rascon6         AAGTGGTACCGTATATTATATATACACTATAGTATTATTTCATCTTATTTTCATTCTGGC      1918
Pachon14        AAGTGGTACCGTATATTATATATACACTATAGTATTATTTCATCTTATTTTCATTCTGGC      1918
Pachon9         AAGTGGTACCGTATATTATATATACACTATAGTATTATTTCATCTTATTTTCATTCTGGC      1918
Pachon17        AAGTGGTACCGTATATTATATATACACTATAGTATTATTTCATCTTATTTTCATTCTGGC      1918
Pachon12        AAGTGGTACCGTATATTATATATACACTATAGTATTATTTCATCTTATTTTCATTCTGGC      1918
Pachon11        AAGTGGTACCGTATATTATATATACACTATAGTATTATTTCATCTTATTTTCATTCTGGC      1918
Pachon7         AAGTGGTACCGTATATTATATATACACTATAGTATTATTTCATCTTATTTTCATTCTGGC      1918
Pachon3         AAGTGGTACCGTATATTATATATACACTATAGTATTATTTCATCTTATTTTCATTCTGGC      1918
Pachon8         AAGTGGTACCGTATATTATATATACACTATAGTATTATTTCATCTTATTTTCATTCTGGC      1918
Pachon15        AAGTGGTACCGTATATTATATATACACTATAGTATTATTTCATCTTATTTTCATTCTGGC      1918
                ************************************************************

Rascon4         CTCTTGTCATTCAATAAAACCCCTATCCAAATAAATCTCTCCATCCAGATAAGTATAGAA      1978
Surface         CTCTTGTCATTCAATAAAACCCCTATCCAAATAAATCTCTCCATCCAGATAAGTATAGAA      1979
Rascon8         CTCTTGTCATTCAATAAAACCCCTATCCAAATAAATCTCTCCATCCAGATAAGTATAGAA      1978
Rascon2         CTCTTGTCATTCAATAAAACCCCTATCCAAATAAATCTCTCCATCCAGATAAGTATAGAA      1977
Rascon15        CTCTTGTCATTCAATAAAACCCCTATCCAAATAAATCTCTCCATCCAGATAAGTATAGAA      1978
Rascon13        CTCTTGTCATTCAATAAAACCCCTATCCAAATAAATCTCTCCATCCAGATAAGTATAGAA      1978
Rascon6         CTCTTGTCATTCAATAAAACCCCTATCCAAATAAATCTCTCCATCCAGATAAGTATAGAA      1978
Pachon14        CTCTTGTCATTCAATAAAACCCCTATCCAAATAAATCTCTCCATCCAGATAAGTATAGAA      1978
Pachon9         CTCTTGTCATTCAATAAAACCCCTATCCAAATAAATCTCTCCATCCAGATAAGTATAGAA      1978
Pachon17        CTCTTGTCATTCAATAAAACCCCTATCCAAATAAATCTCTCCATCCAGATAAGTATAGAA      1978
Pachon12        CTCTTGTCATTCAATAAAACCCCTATCCAAATAAATCTCTCCATCCAGATAAGTATAGAA      1978
Pachon11        CTCTTGTCATTCAATAAAACCCCTATCCAAATAAATCTCTCCATCCAGATAAGTATAGAA      1978
Pachon7         CTCTTGTCATTCAATAAAACCCCTATCCAAATAAATCTCTCCATCCAGATAAGTATAGAA      1978
Pachon3         CTCTTGTCATTCAATAAAACCCCTATCCAAATAAATCTCTCCATCCAGATAAGTATAGAA      1978
Pachon8         CTCTTGTCATTCAATAAAACCCCTATCCAAATAAATCTCTCCATCCAGATAAGTATAGAA      1978
Pachon15        CTCTTGTCATTCAATAAAACCCCTATCCAAATAAATCTCTCCATCCAGATAAGTATAGAA      1978
                ************************************************************

Rascon4         GTAAATGTATATAAGTATAAACATATACAATTCTGACTCCCTATTGGTTTATACTGTGAT      2038
Surface         GTAAATGTATATAAGTATAAACATATACAATTCTGACTCCCTATTGGTTTATACTGTGAT      2039
Rascon8         GTAAATGTATATAAGTATAAACATATACAATTCTGACTCCCTATTGGTTTATACTGTGAT      2038
Rascon2         GTAAATGTATATAAGTATAAACATATACAATTCTGACTCCCTATTGGTTTATACTGTGAT      2037
Rascon15        GTAAATGTATATAAGTATAAACATATACAATTCTGACTCCCTATTGGTTTATACTGTGAT      2038
Rascon13        GTAAATGTATATAAGTATAAACATATACAATTCTGACTCCCTATTGGTTTATACTGTGAT      2038
Rascon6         GTAAATGTATATAAGTATAAACATATACAATTCTGACTCCCTATTGGTTTATACTGTGAT      2038
Pachon14        GTAAATGTATATAAGTATAAACATATACAATTCTGACTCCCTATTGGTTTATACTGTGAT      2038
Pachon9         GTAAATGTATATAAGTATAAACATATACAATTCTGACTCCCTATTGGTTTATACTGTGAT      2038
Pachon17        GTAAATGTATATAAGTATAAACATATACAATTCTGACTCCCTATTGGTTTATACTGTGAT      2038
Pachon12        GTAAATGTATATAAGTATAAACATATACAATTCTGACTCCCTATTGGTTTATACTGTGAT      2038
Pachon11        GTAAATGTATATAAGTATAAACATATACAATTCTGACTCCCTATTGGTTTATACTGTGAT      2038
Pachon7         GTAAATGTATATAAGTATAAACATATACAATTCTGACTCCCTATTGGTTTATACTGTGAT      2038
Pachon3         GTAAATGTATATAAGTATAAACATATACAATTCTGACTCCCTATTGGTTTATACTGTGAT      2038
Pachon8         GTAAATGTATATAAGTATAAACATATACAATTCTGACTCCCTATTGGTTTATACTGTGAT      2038
Pachon15        GTAAATGTATATAAGTATAAACATATACAATTCTGACTCCCTATTGGTTTATACTGTGAT      2038
                ************************************************************
```

```
Rascon4     CCATTACACCTGCACACATCAAAC-------------CCCCCCCACACACACACTCCAGC     2085
Surface     CCATTACACCTGCACACATCACACACCCCTCCCCCCCCCCCCCCACACACACACTCCAGC     2099
Rascon8     CCATTACACCTGCACACATCACAC------CCCCCT-CCCCCCCACACACACACTCCAGC     2091
Rascon2     CCATTACACCTGCACACATCACAC------ACCCCT-CCCCCCCACACACACACTCCAGC     2090
Rascon15    CCATTACACCTGCACACATCACA-------------CCCCCCCCACACACACACTCCAGC     2085
Rascon13    CCATTACACCTGCACACATCAAAC------------CCCCCCCCACACACACACTCCAGC     2086
Rascon6     CCATTACACCTGCACACATCACAC------CCCCCTCCCCCCCCACACACACACTCCAGC     2092
Pachon14    CCATTACACCTGCACACATCACAC------------CCCCCCCCACACACACACTCCAGC     2086
Pachon9     CCATTACACCTGCACACATCACAC------ACCCCTCCCCCCCCACACACACACTCCAGC     2092
Pachon17    CCATTACACCTGCACACATCACAC------CCCCCTCCCCCCCCACACACACACTCCAGC     2092
Pachon12    CCATTACACCTGCACACATCACAC------ACCCCTCCCCCCCCACACACACACTCCAGC     2092
Pachon11    CCATTACACCTGCACACATCACAC------CCCCCTCCCCCCCCACACACACACTCCAGC     2092
Pachon7     CCATTACACCTGCACACATCACAC------ACCCCTCCCCCCCCACACACACACTCCAGC     2092
Pachon3     CCATTACACCTGCACACATCACAC------CCCCCTCCCCCCCCACACACACACTCCAGC     2092
Pachon8     CCATTACACCTGCACACATCACAC------CCCCCTCCCCCCCCACACACACACTCCAGC     2092
Pachon15    CCATTACACCTGCACACATCACAC------CCCCCTCCCCCCCCACACACACACTCCAGC     2092
            ********************* *              ***********************

Rascon4     AAGAACATAGCAGACGTATGTAGCCTAGTGCATCCTAAATGTATCAACATTAACATTTTA     2145
Surface     AAGAACATAGCAGACGTATGTAGCCTAGTGCATCCTAAATGTATCAACATTAACATTTTA     2159
Rascon8     AAGAACATAGCAGACGTATGTAGCCTAGTGCATCCTAAATGTATCAACATTAACATTTTA     2151
Rascon2     AAGAACATAGCAGACGTATGTAGCCTAGTGCATCCTAAATGTATCAACATTAACATTTTA     2150
Rascon15    AAGAACATAGCAGACGTATGTAGCCTAGTGCATCCTAAATGTATCAACATTAACATTTTA     2145
Rascon13    AAGAACATAGCAGACGTATGTAGCCTAGTGCATCCTAAATGTATCAACATTAACATTTTA     2146
Rascon6     AAGAACATAGCAGACGTATGTAGCCTAGTGCATCCTAAATGCATCAACATTAACATTTTA     2152
Pachon14    AAGAACATAGCAGACGTATGTAGCCTAGTGCATCCTAAATGTATCAACATTAACATTTTA     2146
Pachon9     AAGAACATAGCAGACGTATGTAGCCTAGTGCATCCTAAATGTATCAACATTAACATTTTA     2152
Pachon17    AAGAACATAGCAGACGTATGTAGCCTAGTGCATCCTAAATGTATCAACATTAACATTTTA     2152
Pachon12    AAGAACATAGCAGACGTATGTAGCCTAGTGCATCCTAAATGTATCAACATTAACATTTTA     2152
Pachon11    AAGAACATAGCAGACGTATGTAGCCTAGTGCATCCTAAATGTATCAACATTAACATTTTA     2152
Pachon7     AAGAACATAGCAGACGTATGTAGCCTAGTGCATCCTAAATGTATCAACATTAACATTTTA     2152
Pachon3     AAGAACATAGCAGACGTATGTAGCCTAGTGCATCCTAAATGTATCAACATTAACATTTTA     2152
Pachon8     AAGAACATAGCAGACGTATGTAGCCTAGTGCATCCTAAATGTATCAACATTAACATTTTA     2152
Pachon15    AAGAACATAGCAGACGTATGTAGCCTAGTGCATCCTAAATGTATCAACATTAACATTTTA     2152
            ***************************************** ******************

Rascon4     ATAGAACATTATAGTCATTATGGCTTTTAGAAAAATGTGTTTTGGGAAGGGGGGGGGGTA     2205
Surface     ATAGAACATTATAGTCATTATGGCTTTTAGAAAAATGTGTTTTGGGAAGGGGGGGGGGTA     2219
Rascon8     ATAGAACATTATAGTCATTATGGCTTTTAGAAAAATGTGTTTTGGGAAGGGGGGGGGGTA     2211
Rascon2     ATAGAACATTATAGTCATTATGGCTTTTAGAAAAATGTGTTTTGGGAAGGGGGGGGGGTA     2210
Rascon15    ATAGAACATTATAGTCATTATGGCTTTTAGAAAAATGTGTTTTGGGAAGGGGGGGGGGTA     2205
Rascon13    ATAGAACATTATAGTCATTATGGCTTTTAGAAAAATGTGTTTTGGGAA-GGGGGGGGGTA     2205
Rascon6     ATAGAACATTATAGTCATTATGGCTTTTAGAAAAATGTGTTTTGGGAAGGGGGGGGGGTA     2212
Pachon14    ATAGAACATTATAGTCATTATGGCTTTTAGAAAAATGTGTTTTGGGAAGGGGGGGGGGTA     2206
Pachon9     ATAGAACATTATAGTCATTATGGCTTTTAGAAAAATGTGTTTTGGGAAGGGGGGGGGGTA     2212
Pachon17    ATAGAACATTATAGTCATTATGGCTTTTAGAAAAATGTGTTTTGGGAAGGGGGGGGGGTA     2212
Pachon12    ATAGAACATTATAGTCATTATGGCTTTTAGAAAAATGTGTTTTGGGAAGGGGGGGGGGTA     2212
Pachon11    ATAGAACATTATAGTCATTATGGCTTTTAGAAAAATGTGTTTTGGGAAGGGGGGGGGGTA     2212
Pachon7     ATAGAACATTATAGTCATTATGGCTTTTAGAAAAATGTGTTTTGGGAAGGGGGGGGGGTA     2212
Pachon3     ATAGAACATTATAGTCATTATGGCTTTTAGAAAAATGTGTTTTGGGAAGGGGGGGGGGTA     2212
Pachon8     ATAGAACATTATAGTCATTATGGCTTTTAGAAAAATGTGTTTTGGGAAGGGGGGGGGGTA     2212
Pachon15    ATAGAACATTATAGTCATTATGGCTTTTAGAAAAATGTGTTTTGGGAAGGGGGGGGGGTA     2212
            ************************************************ ***********

Rascon4     GTTCTCTATAGGACTCTCGGAGGACTCTGTGGTACTGCCTAAGCTTTTGTTCTTAATGCC     2265
Surface     GTTCTCTATAGGACTCTCGGAGGACTCTGTGGTGCTGCCTAAGCTTTTGTTCTTAATGCC     2279
Rascon8     GTTCTCTATAGGACTCTCGGAGGACTCTGTGGTGCTGCCTAAGCTTTTGTTCTTAATGCC     2271
Rascon2     GTTCTCTATAGGACTCTCGGAGGACTCTGTGGTGCTGCCTAAGCTTTTGTTCTTAATGCC     2270
Rascon15    GTTCTCTATAGGACTCTCGGAGGACTCTGTGGTGCTGCCTAAGCTTTTGTTCTTAATGCC     2265
Rascon13    GTTCTCTATAGGACTCTCGGAGGACTCTGTGGTGCTGCCTAAGCTTTTGTTCTTAATGCC     2265
Rascon6     GTTCTCTATAGGACTCTCGGAGGACTCTGTGGTGCTGCCTAAGCTTTTGTTCTTAATGCC     2272
Pachon14    GTTCTCTATAGGACTCTCGGAGGACTCTGTGGTGCTGCCTAAGCTTTTGTTCTTAATGCC     2266
Pachon9     GTTCTCTATAGGACTCTCGGAGGACTCTGTGGTGCTGCCTAAGCTTTTGTTCTTAATGCC     2272
Pachon17    GTTCTCTATAGGACTCTCGGAGGACTCTGTGGTGCTGCCTAAGCTTTTGTTCTTAATGCC     2272
Pachon12    GTTCTCTATAGGACTCTCGGAGGACTCTGTGGTGCTGCCTAAGCTTTTGTTCTTAATGCC     2272
Pachon11    GTTCTCTATAGGACTCTCGGAGGACTCTGTGGTGCTGCCTAAGCTTTTGTTCTTAATGCC     2272
Pachon7     GTTCTCTATAGGACTCTCGGAGGACTCTGTGGTGCTGCCTAAGCTTTTGTTCTTAATGCC     2272
Pachon3     GTTCTCTATAGGACTCTCGGAGGACTCTGTGGTGCTGCCTAAGCTTTTGTTCTTAATGCC     2272
Pachon8     GTTCTCTATAGGACTCTCGGAGGACTCTGTGGTGCTGCCTAAGCTTTTGTTCTTAATGCC     2272
Pachon15    GTTCTCTATAGGACTCTCGGAGGACTCTGTGGTGCTGCCTAAGCTTTTGTTCTTAATGCC     2272
            ********************************* **************************

Rascon4     ATTTTTTCCCCTTACACTGCTTTTACTTTTACTCTTTAAGTAGTTTTAAAACCAGTACTT     2325
Surface     ATTTTTTCCCCTCACACTGCTTTTACTTTTACTCTTTAAGTAGTTTTAAAACCAGTACTT     2339
Rascon8     ATTTTTTCCCCTTACACTGCTTTTACTTTTACTCTTTAAGTAGTTTTAAAACCAGTACTT     2331
Rascon2     ATTTTTTCCCCTTACACTGCTTTTACTTTTACTCTTTAAGTAGTTTTAAAACCAGTACTT     2330
```

```
Rascon15      ATTTTTTCCCCTTACACTGCTTTTACTTTTACTCTTTAAGTAGTTTTAAAACCAGTACTT      2325
Rascon13      ATTTTTTCCCCTTACACTGCTTTTACTTTTACTCTTTAAGTAGTTTTAAAACCAGTACTT      2325
Rascon6       ATTTTTTCCCCTTACACTGCTTTTACTTTTACTCTTTAAGTAGTTTTAAAACCAGTACTT      2332
Pachon14      ATTTTTTCCCCTTACACTGCTTTTACTTTTACTCTTTAAGTAGTTTTAAAACCAGTACTT      2326
Pachon9       ATTTTTTCCCCTTACACTGCTTTTACTTTTACTCTTTAAGTAGTTTTAAAACCAGTACTT      2332
Pachon17      ATTTTTTCCCCTTACACTGCTTTTACTTTTACTCTTTAAGTAGTTTTAAAACCAGTACTT      2332
Pachon12      ATTTTTTCCCCTTACACTGCTTTTACTTTTACTCTTTAAGTAGTTTTAAAACCAGTACTT      2332
Pachon11      ATTTTTTCCCCTTACACTGCTTTTACTTTTACTCTTTAAGTAGTTTTAAAACCAGTACTT      2332
Pachon7       ATTTTTTCCCCTTACACTGCTTTTACTTTTACTCTTTAAGTAGTTTTAAAACCAGTACTT      2332
Pachon3       ATTTTTTCCCCTTACACTGCTTTTACTTTTACTCTTTAAGTAGTTTTAAAACCAGTACTT      2332
Pachon8       ATTTTTTCCCCTTACACTGCTTTTACTTTTACTCTTTAAGTAGTTTTAAAACCAGTACTT      2332
Pachon15      ATTTTTTCCCCTTACACTGCTTTTACTTTTACTCTTTAAGTAGTTTTAAAACCAGTACTT      2332
              ************ ***********************************************

Rascon4       TTACTTAAAGAGTAAAAGTGTTGAGCTTGAGTTGATACTTCAACTTCTACAGAAGTATTT      2385
Surface       TTACTTAAAGAGTAAAAGTGTTGAGCTTGAGTTGATACTTCAACTTCTACAGAAGTATTT      2399
Rascon8       TTACTTAAAGAGTAAAAGTGTTGAGCTTGAGTTGATACTTCAACTTCTACAGAAGTATTT      2391
Rascon2       TTACTTAAAGAGTAAAAGTGTTGAGCTTGAGTTGATACTTCAACTTCTACAGAAGTATTT      2390
Rascon15      TTACTTAAAGAGTAAAAGTGTTGAGCTTGAGTTGATACTTCAACTTCTACAGAAGTATTT      2385
Rascon13      TTACTTAAAGAGTAAAAGTGTTGAGCTTGAGTTGATACTTCAACTTCTACAGAAGTATTT      2385
Rascon6       TTACTTAAAGAGTAAAAGTGTTGAGCTTGAGTTGATACTTCAACTTCTACAGAAGTATTT      2392
Pachon14      TTACTTAAAGAGTAAAAGTGTTGAGCTTGAGTTGATACTTCAACTTCTACAGAAGTATTT      2386
Pachon9       TTACTTAAAGAGTAAAAGTGTTGAGCTTGAGTTGATACTTCAACTTCTACAGAAGTATTT      2392
Pachon17      TTACTTAAAGAGTAAAAGTGTTGAGCTTGAGTTGATACTTCAACTTCTACAGAAGTATTT      2392
Pachon12      TTACTTAAAGAGTAAAAGTGTTGAGCTTGAGTTGATACTTCAACTTCTACAGAAGTATTT      2392
Pachon11      TTACTTAAAGAGTAAAAGTGTTGAGCTTGAGTTGATACTTCAACTTCTACAGAAGTATTT      2392
Pachon7       TTACTTAAAGAGTAAAAGTGTTGAGCTTGAGTTGATACTTCAACTTCTACAGAAGTATTT      2392
Pachon3       TTACTTAAAGAGTAAAAGTGTTGAGCTTGAGTTGATACTTCAACTTCTACAGAAGTATTT      2392
Pachon8       TTACTTAAAGAGTAAAAGTGTTGAGCTTGAGTTGATACTTCAACTTCTACAGAAGTATTT      2392
Pachon15      TTACTTAAAGAGTAAAAGTGTTGAGCTTGAGTTGATACTTCAACTTCTACAGAAGTATTT      2392
              ************************************************************

Rascon4       TAAAGTCTAGTATCGATATTTCTTCCTACAGAGTAATTTATGGAGATACTTTTGACACCT      2445
Surface       TAAAGTCTAGTATCGATATTTCTTCCTACAGAGTAATTTATGGAGATACTTTTGACACCT      2459
Rascon8       TAAAGTCTAGTATCGATATTTCTTCCTACAGAGTAATTTATGGAGATACTTTTGACACCT      2451
Rascon2       TAAAGTCTAGTATCGATATTTCTTCCTACAGAGTAATTTATGGAGATACTTTTGACACCT      2450
Rascon15      TAAAGTCTAGTATCGATATTTCTTCCTACAGAGTAATTTATGGAGATACTTTTGACACCT      2445
Rascon13      TAAATTCTAGTATCGATATTTCTTCCTACAGAGTAATTTATGGAGATACTTTTGACACCT      2445
Rascon6       TAAAGTCTAGTATCGATATTTCTTCCTACAGAGTAATTTATGGAGATACTTTTGACACCT      2452
Pachon14      TAAAGTCTAGTATCGATATTTCTTCCTACAGAGTAATTTATGGAGATACTTTTGACACCT      2446
Pachon9       TAAAGTCTAGTATCGATATTTCTTCCTACAGAGTAATTTATGGAGATACTTTTGACACCT      2452
Pachon17      TAAAGTCTAGTATCGATATTTCTTCCTACAGAGTAATTTATGGAGATACTTTTGACACCT      2452
Pachon12      TAAAGTCTAGTATCGATATTTCTTCCTACAGAGTAATTTATGGAGATACTTTTGACACCT      2452
Pachon11      TAAAGTCTAGTATCGATATTTCTTCCTACAGAGTAATTTATGGAGATACTTTTGACACCT      2452
Pachon7       TAAAGTCTAGTATCGATATTTCTTCCTACAGAGTAATTTATGGAGATACTTTTGACACCT      2452
Pachon3       TAAAGTCTAGTATCGATATTTCTTCCTACAGAGTAATTTATGGAGATACTTTTGACACCT      2452
Pachon8       TAAAGTCTAGTATCGATATTTCTTCCTACAGAGTAATTTATGGAGATACTTTTGACACCT      2452
Pachon15      TAAAGTCTAGTATCGATATTTCTTCCTACAGAGTAATTTATGGAGATACTTTTGACACCT      2452
              **** *******************************************************

Rascon4       TTGTTGATTATTTAATATTTTAGAATAATATTATAGAAACTGGAATAAAAGTTGGAATAT      2505
Surface       TTGTTGATTATTTAATATTTTAGAATAATATTATAGAAACTGGAATAAAAGTTGGAATAT      2519
Rascon8       TTGTTGATTATTTAATATTTTAGAATAATATTATAGAAACTGGAATAAAAGTTGGAATAT      2511
Rascon2       TTGTTGATTATTTAATATTTTAGAATAATATTATAGAAACTGGAATAAAAGTTGGAATAT      2510
Rascon15      TTGTTGATTATTTAATATTTTAGAATAATATTATAGAAACTGGAATAAAAGTTGGAATAT      2505
Rascon13      TTGTTGATTATTTAATATTTTAGAATAATATTATAGAAACTGGAATAAAAGTTGGAATAT      2505
Rascon6       TTGTTGATTATTTAATATTTTAGAATAATATTATAGAAACTGGAATAAAAGTTGGAATAT      2512
Pachon14      TTGTTGATTATTTAATATTTTAGAATAATATTATAGAAACTGGAATAAAAGTTGGAATAT      2506
Pachon9       TTGTTGATTATTTAATATTTTAGAATAATATTATAGAAACTGGAATAAAAGTTGGAATAT      2512
Pachon17      TTGTTGATTATTTAATATTTTAGAATAATATTATAGAAACTGGAATAAAAGTTGGAATAT      2512
Pachon12      TTGTTGATTATTTAATATTTTAGAATAATATTATAGAAACTGGAATAAAAGTTGGAATAT      2512
Pachon11      TTGTTGATTATTTAATATTTTAGAATAATATTATAGAAACTGGAATAAAAGTTGGAATAT      2512
Pachon7       TTGTTGATTATTTAATATTTTAGAATAATATTATAGAAACTGGAATAAAAGTTGGAATAT      2512
Pachon3       TTGTTGATTATTTAATATTTTAGAATAATATTATAGAAACTGGAATAAAAGTTGGAATAT      2512
Pachon8       TTGTTGATTATTTAATATTTTAGAATAATATTATAGAAACTGGAATAAAAGTTGGAATAT      2512
Pachon15      TTGTTGATTATTTAATATTTTAGAATAATATTATAGAAACTGGAATAAAAGTTGGAATAT      2512
              ************************************************************

Rascon4       AATATAATTATTAATAATTATTACTAGCAATTTCTAATGGTTTTGTTACTGTTGCAGTTC      2565
Surface       GATATAATTATTAATAATTATTACTAGCAATTTCTAATGGTTTTGTTACTGTTGCAGTTC      2579
Rascon8       AATATAATTATTAATAATTATTACTAGCAATTTCTAATGGTTTTGTTACTGTTGCAGTTC      2571
Rascon2       AATATAATTATTAATAATTATTACTAGCAATTTCTAATGGTTTTGTTACTGTTGCAGTTC      2570
Rascon15      AATATAATTATTAATAATTATTACTAGCAATTTCTAATGGTTTTGTTACTGTTGCAGTTC      2565
Rascon13      AATATAATTATTAATAATTATTACTAGCAATTTCTAATGGTTTTGTTACTGTTGCAGTTC      2565
Rascon6       AATATAATTATTAATAATTATTACTAGCAATTTCTAATGGTTTTGTTACTGTTGCAGTTC      2572
Pachon14      AATATAATTATTAATAATTATTACTAGCAATTTCTAATGGTTTTGTTACTGTTGCAGTTC      2566
Pachon9       AATATAATTATTAATAATTATTACTAGCAATTTCTAATGGTTTTGTTACTGTTGCAGTTC      2572
```

```
Pachon17      AATATAATTATTAATAATTATTACTAGCAATTTCTAATGGTTTTGTTACTGTTGCAGTTC      2572
Pachon12      AATATAATTATTAATAATTATTACTAGCAATTTCTAATGGTTTTGTTACTGTTGCAGTTC      2572
Pachon11      AATATAATTATTAATAATTATTACTAGCAATTTCTAATGGTTTTGTTACTGTTGCAGTTC      2572
Pachon7       AATATAATTATTAATAATTATTACTAGCAATTTCTAATGGTTTTGTTACTGTTGCAGTTC      2572
Pachon3       AATATAATTATTAATAATTATTACTAGCAATTTCTAATGGTTTTGTTACTGTTGCAGTTC      2572
Pachon8       AATATAATTATTAATAATTATTACTAGCAATTTCTAATGGTTTTGTTACTGTTGCAGTTC      2572
Pachon15      AATATAATTATTAATAATTATTACTAGCAATTTCTAATGGTTTTGTTACTGTTGCAGTTC      2572
               ********************************** ************************

Rascon4       AGTTCCTTTCAGAACAGCCTGCCCCTAGAGTTCCCCTTCCACACTTAATTTCACTATTGT      2625
Surface       AGTTCCTTTCAGAACAGCCTGCCCCTAGAGTTCCCCTTCCACACTTAATTTCACTATTGT      2639
Rascon8       AGTTCCTTTCAGAACAGCCTGCCCCTAGAGTTCCCCTTCCACACTTAATTTCACTATTGT      2631
Rascon2       AGTTCCTTTCAGAACAGCCTGCCCCTAGAGTTCCCCTTCCACACTTAATTTCACTATTGT      2630
Rascon15      AGTTCCTTTCAGAACAGCCTGCCCCTAGAGTTCCCCTTCCACACTTAATTTCACTATTGT      2625
Rascon13      AGTTCCTTTCAGAACAGCCTGCCCCTAGAGTTCCCCTTCCACACTTAATTTCACTATTGT      2625
Rascon6       AGTTCCTTTCAGAACAGCCTGCCCCTAGAGTTCCCCTTCCACACTTAATTTCACTATTGT      2632
Pachon14      AGTTCCTTTCAGAACAGCCTGCCCCTAGAGTTCCCCTTCCACACTTAATTTCACTATTGT      2626
Pachon9       AGTTCCTTTCAGAACAGCCTGCCCCTAGAGTTCCCCTTCCACACTTAATTTCACTATTGT      2632
Pachon17      AGTTCCTTTCAGAACAGCCTGCCCCTAGAGTTCCCCTTCCACACTTAATTTCACTATTGT      2632
Pachon12      AGTTCCTTTCAGAACAGCCTGCCCCTAGAGTTCCCCTTCCACACTTAATTTCACTATTGT      2632
Pachon11      AGTTCCTTTCAGAACAGCCTGCCCCTAGAGTTCCCCTTCCACACTTAATTTCACTATTGT      2632
Pachon7       AGTTCCTTTCAGAACAGCCTGCCCCTAGAGTTCCCCTTCCACACTTAATTTCACTATTGT      2632
Pachon3       AGTTCCTTTCAGAACAGCCTGCCCCTAGAGTTCCCCTTCCACACTTAATTTCACTATTGT      2632
Pachon8       AGTTCCTTTCAGAACAGCCTGCCCCTAGAGTTCCCCTTCCACACTTAATTTCACTATTGT      2632
Pachon15      AGTTCCTTTCAGAACAGCCTGCCCCTAGAGTTCCCCTTCCACACTTAATTTCACTATTGT      2632
              ************************************************************

Rascon4       AATTTTACGCTGCTAAATGTATCTAATAATCTA-TTTTTTGTCTTTGTGTGTTTTCACTT      2684
Surface       AATTTTACGCTGCTAAATGTATCTAATAATCTATTTTTTTGTCTTTGTGTGTTTTCACTT      2699
Rascon8       AATTTTACGCTGCTAAATGTATCTAATAATCTA-TTTTTTGTCTTTGTGTGTTTTCACTT      2690
Rascon2       AATTTTACGCTGCTAAATGTATCTAATAATCTA-TTTTTTGTCTTTGTGTGTTTTCACTT      2689
Rascon15      AATTTTACGCTGCTAAATGTATCTAATAATCTA-TTTTTTGTCTTTGTGTGTTTTCACTT      2684
Rascon13      AATTTTACGCTGCTAAATGTATCTAATAATCTATTTTTTTGTCTTTGTGTGTTTTCACTT      2685
Rascon6       AATTTTACGCTGCTAAATGTATCTAATAATCTA-TTTTTTGTCTTTGTGTGTTTTCACTT      2691
Pachon14      AATTTTACGCTGCTAAATGTATCTAATAATCTA-TTTTTTGTCTTTGTGTGTTTTCACTT      2685
Pachon9       AATTTTACGCTGCTAAATGTATCTAATAATCTATTTTTTTGTCTTTGTGTGTTTTCACTT      2692
Pachon17      AATTTTACGCTGCTAAATGTATCTAATAATCTA-TTTTTTGTCTTTGTGTGTTTTCACTT      2691
Pachon12      AATTTTACGCTGCTAAATGTATCTAATAATCTA-TTTTTTGTCTTTGTGTGTTTTCACTT      2691
Pachon11      AATTTTACGCTGCTAAATGTATCTAATAATCTA-TTTTTTGTCTTTGTGTGTTTTCACTT      2691
Pachon7       AATTTTACGCTGCTAAATGTATCTAATAATCTA-TTTTTTGTCTTTGTGTGTTTTCACTT      2691
Pachon3       AATTTTACGCTGCTAAATGTATCTAATAATCTA-TTTTTTGTCTTTGTGTGTTTTCACTT      2691
Pachon8       AATTTTACGCTGCTAAATGTATCTAATAATCTA-TTTTTTGTCTTTGTGTGTTTTCACTT      2691
Pachon15      AATTTTACGCTGCTAAATGTATCTAATAATCTA-TTTTTTGTCTTTGTGTGTTTTCACTT      2691
              ********************************* **************************

Rascon4       TTCCTCTATTGAGAGGTCATGAATAAAAACAGCTTAGAGAGACTAATAGACTGTTATCTG      2744
Surface       TTCCTCTATTGAGAGGTCATGAAT-AAAACAGCTTAGAGAGACTAATAGACTGTTATCTG      2758
Rascon8       TTCCTCTATTGAGAGGTCATGAATAAAAACAGCTTAGAGAGACTAATAGACTGTTATCTG      2750
Rascon2       TTCCTCTATTGAGAGGTCATGAATAAAAACAGCTTAGAGAGACTAATAGACTGTTATCTG      2749
Rascon15      TTCCTCTATTGAGAGGTCATGAATAAAAACAGCTTAGAGAGACTAATAGACTGTTATCTG      2744
Rascon13      TTCCTCTATTGAGAGGTCATGAATAAAAACAGCTTAGAGAGACTAATAGACTGTTATCTG      2745
Rascon6       TTCCTCTATTGAGACGTCATGAATAAAAACAGCTTAGAGAGACTAATAGACTGTTATCTG      2751
Pachon14      TTCCTCTATTGAGAGGTCATGAAT-AAAACAGCTTAGAGAGACTAATAGACTGTTATCTG      2744
Pachon9       TTCCTCTATTGAGAGGTCATGAATAAAAACAGCTTAGAGAGACTAATAGACTGTTATCTG      2752
Pachon17      TTCCTCTATTGAGAGGTCATGAATAAAAACAGCTTAGAGAGACTAATAGACTGTTATCTG      2751
Pachon12      TTCCTCTATTGAGAGGTCATGAATAAAAACAGCTTAGAGAGACTAATAGACTGTTATCTG      2751
Pachon11      TTCCTCTATTGAGAGGTCATGAATAAAAACAGCTTAGAGAGACTAATAGACTGTTATCTG      2751
Pachon7       TTCCTCTATTGAGAGGTCATGAATAAAAACAGCTTAGAGAGACTAATAGACTGTTATCTG      2751
Pachon3       TTCCTCTATTGAGAGGTCATGAATAAAAACAGCTTAGAGAGACTAATAGACTGTTATCTG      2751
Pachon8       TTCCTCTATTGAGAGGTCATGAATAAAAACAGCTTAGAGAGACTAATAGACTGTTATCTG      2751
Pachon15      TTCCTCTATTGAGAGGTCATGAATAAAAACAGCTTAGAGAGACTAATAGACTGTTATCTG      2751
              ************** ********* ***********************************

Rascon4       TTGTAATAACACACAATCTAAACAGTGGGAATAAAGGGGGGGCTTTGTTTTAGTGTTTTG      2804
Surface       TTGTAATAACACACAATCTAAACAGTGGGAATAAAGGGGGGGCTTTGTTTTAGTGTTTTG      2818
Rascon8       TTGTAATAACACACAATCTAAACAGTGGGAATAAAGGGGGGGCTTTGTTTTAGTGTTTTG      2810
Rascon2       TTGTAATAACACACAATCTAAACAGTGGGAATAAAGGGGGGGCTTTGTTTTAGTGTTTTG      2809
Rascon15      TTGTAATAACACACAATCTAAACAGTGGGAATAAAGGGGGGGCTTTGTTTTAGTGTTTTG      2804
Rascon13      TTGTAATAACACACAATCTAAACAGTGGGAATAAAGGGGGGGCTTTGTTTTAGTGTTTTG      2805
Rascon6       TTGTAATAACACACAATCTAAACAGTGGGAATAAAGGGGGGGCTTTGTTTTAGTGTTTTG      2811
Pachon14      TTGTAATAACACACAATCTAAACAGTGGGAATAAAGGGGGGGCTTTGTTTTAGTGTTTTG      2804
Pachon9       TTGTAATAACACACAATCTAAACAGTGGGAATAAAGGGGGGGCTTTGTTTTAGTGTTTTG      2812
Pachon17      TTGTAATAACACACAATCTAAACAGTGGGAATAAAGGGGGGGCTTTGTTTTAGTGTTTTG      2811
Pachon12      TTGTAATAACACACAATCTAAACAGTGGGAATAAAGGGGGGGCTTTGTTTTAGTGTTTTG      2811
Pachon11      TTGTAATAACACACAATCTAAACAGTGGGAATAAAGGGGGGGCTTTGTTTTAGTGTTTTG      2811
Pachon7       TTGTAATAACACACAATCTAAACAGTGGGAATAAAGGGGGGGCTTTGTTTTAGTGTTTTG      2811
Pachon3       TTGTAATAACACACAATCTAAACAGTGGGAATAAAGGGGGGGCTTTGTTTTAGTGTTTTG      2811
```

```
Pachon8      TTGTAATAACACACAATCTAAACAGTGGGAATAAAGGGGGGGCTTTGTTTTAGTGTTTTG   2811
Pachon15     TTGTAATAACACACAATCTAAACAGTGGGAATAAAGGGGGGGCTTTGTTTTAGTGTTTTG   2811
             ************************************************************

Rascon4      AGGGGAGAAAAGCGTAATCCCCTTAAACAGCGTGAGGCCGGCGGATTACCAGCCTGCTGT   2864 Not in
Surface      AGGGGAGAAAAGCGTAATCCCCTTAAACAGCGTGAGGCCGGCGGATTACCAGCCTGCTGT   2878 Choy SF
Rascon8      AGGGGAGAAAAGCGTAATCCCCTTAAACAGCGTGAGGCCGGCGGATTACCAGCCTGCTGT   2870 Mix A/G
Rascon2      AGGGGAGAAAAGCGTAATCCCCTTAAACAGCGTGAGGCCGGCGGATTACCAGCCTGCTGT   2869
Rascon15     AGGGGAGAAAAGCGTAATCCCCTTAAACAGCGTGAGGCCGGCGGATTACCAGCCTGCTGT   2864
Rascon13     AGGGGAGAAAAGCGTAATCCCCTTAAACAGCGTGAGGCCGGCGGATTACCAGCCTGCTGT   2865
Rascon6      AGGGGAGAAAAGCGTAATCCCCTTAAACAGTGTGAGGCCGGCAGATTACCAGCCTGCTGT   2871
Pachon14     AGGGGAGAAAAGCGTAATCCCCTTAAACAGCGTGAGGCCGGCAGATTACCAGCCTGCTGT   2864
Pachon9      AGGGGAGAAAAGCGTAATCCCCTTAAACAGCGTGAGGCCGGCAGATTACCAGCCTGCTGT   2872
Pachon17     AGGGGAGAAAAGCGTAATCCCCTTAAACAGCGTGAGGCCGGCAGATTACCAGCCTGCTGT   2871
Pachon12     AGGGGAGAAAAGCGTAATCCCCTTAAACAGCGTGAGGCCGGCAGATTACCAGCCTGCTGT   2871
Pachon11     AGGGGAGAAAAGCGTAATCCCCTTAAACAGCGTGAGGCCGGCAGATTACCAGCCTGCTGT   2871
Pachon7      AGGGGAGAAAAGCGTAATCCCCTTAAACAGCGTGAGGCCGGCAGATTACCAGCCTGCTGT   2871
Pachon3      AGGGGAGAAAAGCGTAATCCCCTTAAACAGCGTGAGGCCGGCAGATTACCAGCCTGCTGT   2871
Pachon8      AGGGGAGAAAAGCGTAATCCCCTTAAACAGCGTGAGGCCGGCAGATTACCAGCCTGCTGT   2871
Pachon15     AGGGGAGAAAAGCGTAATCCCCTTAAACAGCGTGAGGCCGGCAGATTACCAGCCTGCTGT   2871
             ****************************** *********** *****************

Rascon4      CAGTCTTACTTCACCACTAATTACACATATCCATTCAGCACTGAGGAAGTTGACAAATTC   2924 SNP2
Surface      CAGTCTTACTTCACCACTAATTACACATATCCATTCAGCACTGAGGAAGTTGACAAATTC   2938 also
Rascon8      CAGTCTTACTTCACCACTAATTACACATATCCATTCAGCACTGAGGAAGTTGACAAATTC   2930 fixed in
Rascon2      CAGTCTTACTTCACCACTAATTACACATATCCATTCAGCACTGAGGAAGTTGACAAATTC   2929 Choy SF
Rascon15     CAGTCTTACTTCACCACTAATTACACATATCCATTCAGCACTGAGGAAGTTGACAAATTC   2924
Rascon13     CAGTCTTACTTCACCACTAATTACACATATCCATTCAGCACTGAGGAAGTTGACAAATTC   2925
Rascon6      CAGTCTTACTTCACCACTAATTACACATATCCATTCAGCACTGAGGAAGTTGACAAATTC   2931
Pachon14     CAGTCTTACTTCACCACTAATTACACATATCCATTCAGCACTGAGGAAGTTGACAAATTA   2924
Pachon9      CAGTCTTACTTCACCACTAATTACACATATCCATTCAGCACTGAGGAAGTTGACAAATTA   2932
Pachon17     CAGTCTTACTTCACCACTAATTACACATATCCATTCAGCACTGAGGAAGTTGACAAATTA   2931
Pachon12     CAGTCTTACTTCACCACTAATTACACATATCCATTCAGCACTGAGGAAGTTGACAAATTA   2931
Pachon11     CAGTCTTACTTCACCACTAATTACACATATCCATTCAGCACTGAGGAAGTTGACAAATTA   2931
Pachon7      CAGTCTTACTTCACCACTAATTACACATATCCATTCAGCACTGAGGAAGTTGACAAATTA   2931
Pachon3      CAGTCTTACTTCACCACTAATTACACATATCCATTCAGCACTGAGGAAGTTGACAAATTA   2931
Pachon8      CAGTCTTACTTCACCACTAATTACACATATCCATTCAGCACTGAGGAAGTTGACAAATTA   2931
Pachon15     CAGTCTTACTTCACCACTAATTACACATATCCATTCAGCACTGAGGAAGTTGACAAATTA   2931
             *********************************************************** 

Rascon4      TTCCAATTAAGTTCACCTCAGTCACT----------------------------------   2950
Surface      TTCCAATTAAGTTCACCTCAGTCACTCTCACACACACACACACACACACACACACACACA   2998
Rascon8      TTCCAATTAAGTTCACCTCAGTCACTCTCACACACACACACACACACACACACACACACA   2990
Rascon2      TTCCAATTAAGTTCACCTCAGTCACTCTCACACACACACACACACACACACACACACACA   2989
Rascon15     TTCCAATTAAGTTCACCTCAGTCACTCTCACACACACACACACACACACACACACACACA   2984
Rascon13     TTCCAATTAAGTTCACCTCAGTCACTCTCACACACACACACACACACACACACACACACA   2985
Rascon6      TTCCAATTAAGTTCACCTCAGTCACTCTCACACACACACACACACACACACACACACACA   2991
Pachon14     TTCCAATTAAGTTCACCTCAGTCACTCTCACACACACACACACACACACACACACACACA   2984
Pachon9      TTCCAATTAAGTTCACCTCAGTCACTCTCACACACACACACACACACACACACACACACA   2992
Pachon17     TTCCAATTAAGTTCACCTCAGTCACTCTCACACACACACACACACACACACACACACACA   2991
Pachon12     TTCCAATTAAGTTCACCTCAGTCACTCTCACACACACACACACACACACACACACACACA   2991
Pachon11     TTCCAATTAAGTTCACCTCAGTCACTCTCACACACACACACACACACACACACACACACA   2991
Pachon7      TTCCAATTAAGTTCACCTCAGTCACTCTCACACACACACACACACACACACACACACACA   2991
Pachon3      TTCCAATTAAGTTCACCTCAGTCACTCTCACACACACACACACACACACACACACACACA   2991
Pachon8      TTCCAATTAAGTTCACCTCAGTCACTCTCACACACACACACACACACACACACACACACA   2991
Pachon15     TTCCAATTAAGTTCACCTCAGTCACTCTCACACACACACACACACACACACACACACACA   2991
             **************************

Rascon4      --CACACACACACACACACACACACACCTCTTCAGCTCATCTCAGGTCTGGGCAAGTG   3008
Surface      CACACACACACACACACACACACACACACCTCTTCAGCTCATCTCAGGTCTGGGCAAGTG   3058
Rascon8      CACACACACACACACACACACACACACACCTCTTCAGCTCATCTCAGGTCTGGGCAAGTG   3050
Rascon2      CACACACACACACACACACACACACACACCTCTTCAGCTCATCTCAGGTCTGGGCAAGTG   3049
Rascon15     CACACACACACACACACACACACACACACCTCTTCAGCTCATCTCAGGTCTGGGCAAGTG   3044
Rascon13     CACACACACACACACACACACACACACACCTCTTCAGCTCATCTCAGGTCTGGGCAAGTG   3045
Rascon6      CACACACACACACACACACACACACACACCTCTTCAGCTCATCTCAGGTCTGGGCAAGTG   3051
Pachon14     CACACACACACACACACACACACACACACCTCTTCAGCTCATCTCAGGTCTGGGCAAGTG   3044
Pachon9      CACACACACACACACACACACACACACACCTCTTCAGCTCATCTCAGGTCTGGGCAAGTG   3052
Pachon17     CACACACACACACACACACACACACACACCTCTTCAGCTCATCTCAGGTCTGGGCAAGTG   3051
Pachon12     CACACACACACACACACACACACACACACCTCTTCAGCTCATCTCAGGTCTGGGCAAGTG   3051
Pachon11     CACACACACACACACACACACACACACACCTCTTCAGCTCATCTCAGGTCTGGGCAAGTG   3051
Pachon7      CACACACACACACACACACACACACACACCTCTTCAGCTCATCTCAGGTCTGGGCAAGTG   3051
Pachon3      CACACACACACACACACACACACACACACCTCTTCAGCTCATCTCAGGTCTGGGCAAGTG   3051
Pachon8      CACACACACACACACACACACACACACACCTCTTCAGCTCATCTCAGGTCTGGGCAAGTG   3051
Pachon15     CACACACACACACACACACACACACACACCTCTTCAGCTCATCTCAGGTCTGGGCAAGTG   3051
               **********************************************************

Rascon4      AGTGTGAAGGAGTTTGTAAGAGAGATTAGTGGCTCTGCAGCCGCCTCCATTAACTCTGCT   3068
```

```
Surface       AGTGTGAAGGAGTTTGTAAGAGAGATTAGTGGCTCTGCAGCCGCCTCCATTAACTCTGCT    3118
Rascon8       AGTGTGAAGGAGTTTGTAAGAGAGATTAGTGGCTCTGCAGCCGCCTCCATTAACTCTGCT    3110
Rascon2       AGTGTGAAGGAGTTTGTAAGAGAGATTAGTGGCTCTGCAGCCGCCTCCATTAACTCTGCT    3109
Rascon15      AGTGTGAAGGAGTTTGTAAGAGAGATTAGTGGCTCTGCAGCCGCCTCCATTAACTCTGCT    3104
Rascon13      AGTATGAAGGAGTTTGTAAGAGAGATTAGTGGCTCTGCAGCCGCCTCCATTAACTCTGCT    3105
Rascon6       AGTGTGAAGGAGTTTGTAAGAGAGATTAGTGGCTCTGCAGCCGCCTCCATTAACTCTGCT    3111
Pachon14      AGTGTGAAGGAGTTTGTAAGAGAGATTAGTGGCTCTGCAGCCGCCTCCATTAACTCTGCT    3104
Pachon9       AGTGTGAAGGAGTTTGTAAGAGAGATTAGTGGCTCTGCAGCCGCCTCCATTAACTCTGCT    3112
Pachon17      AGTGTGAAGGAGTTTGTAAGAGAGATTAGTGGCTCTGCAGCCGCCTCCATTAACTCTGCT    3111
Pachon12      AGTGTGAAGGAGTTTGTAAGAGAGATTAGTGGCTCTGCAGCCGCCTCCATTAACTCTGCT    3111
Pachon11      AGTGTGAAGGAGTTTGTAAGAGAGATTAGTGGCTCTGCAGCCGCCTCCATTAACTCTGCT    3111
Pachon7       AGTGTGAAGGAGTTTGTAAGAGAGATTAGTGGCTCTGCAGCCGCCTCCATTAACTCTGCT    3111
Pachon3       AGTGTGAAGGAGTTTGTAAGAGAGATTAGTGGCTCTGCAGCCGCCTCCATTAACTCTGCT    3111
Pachon8       AGTGTGAAGGAGTTTGTAAGAGAGATTAGTGGCTCTGCAGCCGCCTCCATTAACTCTGCT    3111
Pachon15      AGTGTGAAGGAGTTTGTAAGAGAGATTAGTGGCTCTGCAGCCGCCTCCATTAACTCTGCT    3111
              *** ********************************************************

Rascon4       TTATAATGGTGTTTTACAATTCCAGCCCAGTCCAGCACCCCGCTAACATCTGCTCTACAG    3128
Surface       TTATAATGGTGTTTTACAATTCCAGCCCAGTCCAGCACCCCGCTAACATCTGCTCTACAG    3178
Rascon8       TTATAATGGTGTTTTACAATTCCAGCCCAGTCCAGCACCCCGCTAACATCTGCTCTACAG    3170
Rascon2       TTATAATGGTGTTTTACAATTCCAGCCCAGTCCAGCACCCCGCTAACATCTGCTCTACAG    3169
Rascon15      TTATAATGGTGTTTTACAATTCCAGCCCAGTCCAGCACCCCGCTAACATCTGCTCTACAG    3164
Rascon13      TTATAATGGTGTTTTACAATTCCAGCCCAGTCCAGCACCCCGCTAACATCTGCTCTACAG    3165
Rascon6       TTATAATGGTGTTTTACAATTCCAGCCCAGTCCAGCACCCCGCTAACATCTGCTCTACAG    3171
Pachon14      TTATAATGGTGTTTTACAATTCCAGCCCAGTCCAGCACCCCGCTAACATCTGCTCTACAG    3164
Pachon9       TTATAATGGTGTTTTACAATTCCAGCCCAGTCCAGCACCCCGCTAACATCTGCTCTACAG    3172
Pachon17      TTATAATGGTGTTTTACAATTCCAGCCCAGTCCAGCACCCCGCTAACATCTGCTCTACAG    3171
Pachon12      TTATAATGGTGTTTTACAATTCCAGCCCAGTCCAGCACCCCGCTAACATCTGCTCTACAG    3171
Pachon11      TTATAATGGTGTTTTACAATTCCAGCCCAGTCCAGCACCCCGCTAACATCTGCTCTACAG    3171
Pachon7       TTATAATGGTGTTTTACAATTCCAGCCCAGTCCAGCACCCCGCTAACATCTGCTCTACAG    3171
Pachon3       TTATAATGGTGTTTTACAATTCCAGCCCAGTCCAGCACCCCGCTAACATCTGCTCTACAG    3171
Pachon8       TTATAATGGTGTTTTACAATTCCAGCCCAGTCCAGCACCCCGCTAACATCTGCTCTACAG    3171
Pachon15      TTATAATGGTGTTTTACAATTCCAGCCCAGTCCAGCACCCCGCTAACATCTGCTCTACAG    3171
              ************************************************************

Rascon4       ACGCCCTGCGGTCACAAGCTGACCTGCGGTGAATGCAGTGATGCAGAGCTGATAAACTTT    3188
Surface       ACGCCCTGCGGTCACAAGCTGACCTGCGGTGAATGCAGTGATGCAGAGCTGATAAACTTT    3238
Rascon8       ACGCCCTGCGGTCACAAGCTGACCTGCGGTGAATGCAGTGATGCAGAGCTGATAAACTTT    3230
Rascon2       ACGCCCTGCGGTCACAAGCTGACCTGCGGTGAATGCAGTGATGCAGAGCTGATAAACTTT    3229
Rascon15      ACGCCCTGCGGTCACAAGCTGACCTGCGGTGAATGCAGTGATGCAGAGCTGATAAACTTT    3224
Rascon13      ACGCCCTGCGGTCACAAGCTGACCTGCGGTGAATGCAGTGATGCAGAGCTGATAAACTTT    3225
Rascon6       ACGCCCTGCGGTCACAAGCTGACCTGCGGTGAATGCAGTGATGCAGAGCTGATAAACTTT    3231
Pachon14      ACGCCCTGCGGTCACAAGCTGACCTGCGGTGAATGCAGTGATGCAGAGCTGATAAACTTT    3224
Pachon9       ACGCCCTGCGGTCACAAGCTGACCTGCGGTGAATGCAGTGATGCAGAGCTGATAAACTTT    3232
Pachon17      ACGCCCTGCGGTCACAAGCTGACCTGCGGTGAATGCAGTGATGCAGAGCTGATAAACTTT    3231
Pachon12      ACGCCCTGCGGTCACAAGCTGACCTGCGGTGAATGCAGTGATGCAGAGCTGATAAACTTT    3231
Pachon11      ACGCCCTGCGGTCACAAGCTGACCTGCGGTGAATGCAGTGATGCAGAGCTGATAAACTTT    3231
Pachon7       ACGCCCTGCGGTCACAAGCTGACCTGCGGTGAATGCAGTGATGCAGAGCTGATAAACTTT    3231
Pachon3       ACGCCCTGCGGTCACAAGCTGACCTGCGGTGAATGCAGTGATGCAGAGCTGATAAACTTT    3231
Pachon8       ACGCCCTGCGGTCACAAGCTGACCTGCGGTGAATGCAGTGATGCAGAGCTGATAAACTTT    3231
Pachon15      ACGCCCTGCGGTCACAAGCTGACCTGCGGTGAATGCAGTGATGCAGAGCTGATAAACTTT    3231
              ************************************************************

Rascon4       ACAACAGCAGATCAGCTGCAGTCTGGAACAGTATATTTGCAGAGTGGAGCTGTATAGTTG    3248
Surface       ACAACAGCAGATCAGCTGCAGTCTGGAACTGTATATTTGCAGAGTGGAGCTGTATAGTTG    3298
Rascon8       ACAACAGCAGATCAGCTGCAGTCTGGAACTGTATATTTGCAGAGTGGAGCTGTATAGTTG    3290
Rascon2       ACAACAGCAGATCAGCTGCAGTCTGGAACTGTATATTTGCAGAGTGGAGCTGTATAGTTG    3289
Rascon15      ACAACAGCAGATCAGCTGCAGTCTGGAACTGTATATTTGCAGAGTGGAGCTGTATAGTTG    3284
Rascon13      ACAACAGCAGATCAGCTGCAGTCTGGAACTGTATATTTGCAGAGTGGAGCTGTATAGTTG    3285
Rascon6       ACAACAGCAGATCAGCTGCAGTCTGGAACTGTATATTTGCAGAGTGGAGCTGTATAGTTG    3291
Pachon14      ACAACAGCAGATCAGCTGCAGTCTGGAACTGTATATTTGCAGAGTGGAGCTGTATAGTTG    3284
Pachon9       ACAACAGCAGATCAGCTGCAGTCTGGAACTGTATATTTGCAGAGTGGAGCTGTATAGTTG    3292
Pachon17      ACAACAGCAGATCAGCTGCAGTCTGGAACTGTATATTTGCAGAGTGGAGCTGTATAGTTG    3291
Pachon12      ACAACAGCAGATCAGCTGCAGTCTGGAACTGTATATTTGCAGAGTGGAGCTGTATAGTTG    3291
Pachon11      ACAACAGCAGATCAGCTGCAGTCTGGAACTGTATATTTGCAGAGTGGAGCTGTATAGTTG    3291
Pachon7       ACAACAGCAGATCAGCTGCAGTCTGGAACTGTATATTTGCAGAGTGGAGCTGTATAGTTG    3291
Pachon3       ACAACAGCAGATCAGCTGCAGTCTGGAACTGTATATTTGCAGAGTGGAGCTGTATAGTTG    3291
Pachon8       ACAACAGCAGATCAGCTGCAGTCTGGAACTGTATATTTGCAGAGTGGAGCTGTATAGTTG    3291
Pachon15      ACAACAGCAGATCAGCTGCAGTCTGGAACTGTATATTTGCAGAGTGGAGCTGTATAGTTG    3291
              ***************************** ******************************

Rascon4       TAAAGTATAGGTATAGCTTGTCCACAGGTGTATGCAGTAAGGTATAGATGTTAGATGTTT    3308
Surface       TAAAGTATAGGTATAGCTTGTCCACAGGTGTATGCAGTAAGGTATAGATGTTAGATGTTT    3358
Rascon8       TAAAGTATAGGTATAGCTTGTCCACAGGTGTATGCAGTAAGGTATAGATGTTAGATGTTT    3350
Rascon2       TAAAGTATAGGTATAGCTTGTCCACAGGTGTATGCAGTAAGGTATAGATGTTAGATGTTT    3349
Rascon15      TAAAGTATAGGTATAGCTTGTCCACAGGTGTATGCAGTAAGGTATAGATGTTAGATGTTT    3344
Rascon13      TAAAGTATAGGTATAGCTTGTCCACAGGTGTATGCAGTAAGGTATAGATGTTAAATGTTT    3345
```

```
Rascon6      TAAAGTATAGGTATAGCTTGTCCACAGGTGTATGCAGTAAGGTATAGATGTTAGATGTTT     3351
Pachon14     TAAAGTATAGGTATAGCTTGTCCACAGGTGTATGCAGTAAGGTATAGATGTTAGATGTTT     3344
Pachon9      TAAAGTATAGGTATAGCTTGTCCACAGGTGTATGCAGTAAGGTATAGATGTTAGATGTTT     3352
Pachon17     TAAAGTATAGGTATAGCTTGTCCACAGGTGTATGCAGTAAGGTATAGATGTTAGATGTTT     3351
Pachon12     TAAAGTATAGGTATAGCTTGTCCACAGGTGTATGCAGTAAGGTATAGATGTTAGATGTTT     3351
Pachon11     TAAAGTATAGGTATAGCTTGTCCACAGGTGTATGCAGTAAGGTATAGATGTTAGATGTTT     3351
Pachon7      TAAAGTATAGGTATAGCTTGTCCACAGGTGTATGCAGTAAGGTATAGATGTTAGATGTTT     3351
Pachon3      TAAAGTATAGGTATAGCTTGTCCACAGGTGTATGCAGTAAGGTATAGATGTTAGATGTTT     3351
Pachon8      TAAAGTATAGGTATAGCTTGTCCACAGGTGTATGCAGTAAGGTATAGATGTTAGATGTTT     3351
Pachon15     TAAAGTATAGGTATAGCTTGTCCACAGGTGTATGCAGTAAGGTATAGATGTTAGATGTTT     3351
             ***************************************************** ******

Rascon4      CTAATAAAGT-------TAGTGTGTTTTATTCATTTGTATTGTAATTTGTACAGTAATTA     3361
Surface      CTAATAAAGTGGACTGATAGTGTGTTTTATTCATTTGTATTGTAATTTGTACAGTAATTA     3418
Rascon8      CTAATAAAGT-------TAGTGTGTTTTATTCATTTGTATTGTAATTTGTACAGTAATTA     3403
Rascon2      CTAATAAAGTGGACTGATAGTGTGTTTTATTCATTTGTATTGTAATCTGTACAGTAATTA     3409
Rascon15     CTAATAAAGT-------TAGTGTGTTTTATTCATTTGTATTGTAATTTGTACAGTAATTA     3397
Rascon13     CTAATAAAGTGGACTGATAGTGTGTTTTATTCATTTGTATTGTAATTTGTACAGTAATTA     3405
Rascon6      CTAATAAAGT-------TAGTGTGTTTTATTCATTTGTATTGTAATTTGTACAGTAATTA     3404
Pachon14     CTAATAAAGTGGACTGATAGTGTGTTTTATTCATTTGTATTGTAATTTGTACAGTAATTA     3404
Pachon9      CTAATAAAGTGGACTGATAGTGTGTTTTATTCATTTGTATTGTAATTTGTACAGTAATTA     3412
Pachon17     CTAATAAAGTGGACTGATAGTGTGTTTTATTCATTTGTATTGTAATTTGTACAGTAATTA     3411
Pachon12     CTAATAAAGTGGACTGATAGTGTGTTTTATTCATTTGTATTGTAATTTGTACAGTAATTA     3411
Pachon11     CTAATAAAGTGGACTGATAGTGTGTTTTATTCATTTGTATTGTAATTTGTACAGTAATTA     3411
Pachon7      CTAATAAAGTGGACTGATAGTGTGTTTTATTCATTTGTATTGTAATTTGTACAGTAATTA     3411
Pachon3      CTAATAAAGTGGACTGATAGTGTGTTTTATTCATTTGTATTGTAATTTGTACAGTAATTA     3411
Pachon8      CTAATAAAGTGGACTGATAGTGTGTTTTATTCATTTGTATTGTAATTTGTACAGTAATTA     3411
Pachon15     CTAATAAAGTGGACTGATAGTGTGTTTTATTCATTTGTATTGTAATTTGTACAGTAATTA     3411
             **********       ***************************** *************

Rascon4      CTTTGAAAATGCAACTAAATACTATTAATTAATACTATATATAAGATTAACAATTGATCT     3421
Surface      CTTTGAAAATGCAACTAAATACTATTAATTAATACTATATATAAGATTAACAATTGATCT     3478
Rascon8      CTTTGAAAATGCAACTAAATACTATTAATTAATACTATATATAAGATTAACAATTGATCT     3463
Rascon2      CTTTGAAAATGCAACTAAATACTATTAATTAATACTATATATAAGATTAACAATTGATCT     3469
Rascon15     CTTTGAAAATGCAACTAAATACTATTAATTAATACTATATATAAGATTAACAATTGATCT     3457
Rascon13     CTTTGAAAATGCAACTAAATACTATTAATTAATACTATATATAAGATTAACAATTGATCT     3465
Rascon6      CTTTGAAAATGCAACTAAATACTATTAATTAATACTATATATAAGATTAACAATTGATCT     3464
Pachon14     CTTTGAAAATGCAACTAAATACTATTAATTAATACTATATATAAGATTAACAATTGATCT     3464
Pachon9      CTTTGAAAATGCAACTAAATACTATTAATTAATACTATATATAAGATTAACAATTGATCT     3472
Pachon17     CTTTGAAAATGCAACTAAATACTATTAATTAATACTATATATAAGATTAACAATTGATCT     3471
Pachon12     CTTTGAAAATGCAACTAAATACTATTAATTAATACTATATATAAGATTAACAATTGATCT     3471
Pachon11     CTTTGAAAATGCAACTAAATACTATTAATTAATACTATATATAAGATTAACAATTGATCT     3471
Pachon7      CTTTGAAAATGCAACTAAATACTATTAATTAATACTATATATAAGATTAACAATTGATCT     3471
Pachon3      CTTTGAAAATGCAACTAAATACTATTAATTAATACTATATATAAGATTAACAATTGATCT     3471
Pachon8      CTTTGAAAATGCAACTAAATACTATTAATTAATACTATATATAAGATTAACAATTGATCT     3471
Pachon15     CTTTGAAAATGCAACTAAATACTATTAATTAATACTATATATAAGATTAACAATTGATCT     3471
             ************************************************************

Rascon4      TGGGACAGATTTCTGTTTCATGTACACAACTCGCATATAAACCTTATAGTAGCACCTGAC     3481
Surface      TGGGACAGATTTCTGTTTCATGTACACAACTCGCATATAAACCTTATAGTAGCACCTGAC     3538
Rascon8      TGGGACAGATTTCTGTTTCATGTACACAACTCGCATATAAACCTTATAGTAGCACCTGAC     3523
Rascon2      TGGGACAGATTTCTGTTTCATGTACACAACTCGCATATAAACCTTATAGTAGCACCTGAC     3529
Rascon15     TGGGACAGATTTCTGTTTCATGTACACAACTCGCATATAAACCTTATAGTAGCACCTGAC     3517
Rascon13     TGGGACAGATTTCTGTTTCATGTACACAACTCGCATATAAACCTTATAGTAGCACCTGAC     3525
Rascon6      TGGGACAGATTTCTGTTTCATGTACACAACTCGCATATAAACCTTATAGTAGCACCTGAC     3524
Pachon14     TGGGACAGATTTCTGTTTCATGTACACAACTCGCATATAAACCTTATAGTAGCACCTGAC     3524
Pachon9      TGGGACAGATTTCTGTTTCATGTACACAACTCGCATATAAACCTTATAGTAGCACCTGAC     3532
Pachon17     TGGGACAGATTTCTGTTTCATGTACACAACTCGCATATAAACCTTATAGTAGCACCTGAC     3531
Pachon12     TGGGACAGATTTCTGTTTCATGTACACAACTCGCATATAAACCTTATAGTAGCACCTGAC     3531
Pachon11     TGGGACAGATTTCTGTTTCATGTACACAACTCGCATATAAACCTTATAGTAGCACCTGAC     3531
Pachon7      TGGGACAGATTTCTGTTTCATGTACACAACTCGCATATAAACCTTATAGTAGCACCTGAC     3531
Pachon3      TGGGACAGATTTCTGTTTCATGTACACAACTCGCATATAAACCTTATAGTAGCACCTGAC     3531
Pachon8      TGGGACAGATTTCTGTTTCATGTACACAACTCGCATATAAACCTTATAGTAGCACCTGAC     3531
Pachon15     TGGGACAGATTTCTGTTTCATGTACACAACTCGCATATAAACCTTATAGTAGCACCTGAC     3531
             ************************************************************

Rascon4      ATAACTGAATATGTTACTGTGGTGTACTATTATTTATAGTGGATACTGGAGAGCATTTAC     3541
Surface      ATAACTGAATATGTTACTGTGGTGTACTATTATTTATAGCGGATACTGGAGAGCATTTAC     3598
Rascon8      ATAACTGAATATGTTACTGTGGTGTACTATTATTTATAGTGGATACTGGAGAGCATTTAC     3583
Rascon2      ATAACTGAATATGTTACTGTGGTGTACTATTATTTATAGTGGATACTGGAGAGCATTTAC     3589
Rascon15     ATAACTGAATATGTTACTGTGGTGTACTATTATTTATAGTGGATACTGGAGAGCATTTAC     3577
Rascon13     ATAACTGAATATGTTACTGTGGTGTACTATTATTTATAGTGGATACTGGAGAGCATTTAC     3585
Rascon6      ATAACTGAATATGTTACTGTGGTGTACTATTATTTATAGTGGATACTGGAGAGCATTTAC     3584
Pachon14     ATAACTGAATATGTTACTGTGGTGTACTATTATTTATAGTGGATACTGGAGAGCATTTAC     3584
Pachon9      ATAACTGAATATGTTACTGTGGTGTACTATTATTTATAGTGGATACTGGAGAGCATTTAC     3592
Pachon17     ATAACTGAATATGTTACTGTGGTGTACTATTATTTATAGTGGATACTGGAGAGCATTTAC     3591
Pachon12     ATAACTGAATATGTTACTGTGGTGTACTATTATTTATAGTGGATACTGGAGAGCATTTAC     3591
```

```
Pachon11        ATAACTGAATATGTTACTGTGGTGTACTATTATTTATAGTGGATACTGGAGAGCATTTAC      3591
Pachon7         ATAACTGAATATGTTACTGTGGTGTACTATTATTTATAGTGGATACTGGAGAGCATTTAC      3591
Pachon3         ATAACTGAATATGTTACTGTGGTGTACTATTATTTATAGTGGATACTGGAGAGCATTTAC      3591
Pachon8         ATAACTGAATATGTTACTGTGGTGTACTATTATTTATAGTGGATACTGGAGAGCATTTAC      3591
Pachon15        ATAACTGAATATGTTACTGTGGTGTACTATTATTTATAGTGGATACTGGAGAGCATTTAC      3591
                *************************************** ********************

Rascon4         TGAATCATTTCTTAGACTATAATTCATTCACAGTAAATACCTTATACTTTTTCATGGCGA      3601
Surface         TGAATCATTTCTTAGACTATAATTCATTCACAGTAAATACCTTATACTTTTTCATGGCGA      3658
Rascon8         TGAATCATTTCTTAGACTATAATTCATTCACAGTAAATACCTTATACTTTTTCATGGCGA      3643
Rascon2         TGAATCATTTCTTAGACTATAATTCATTCACAGTAAATACCTTATACTTTTTCATGGCGA      3649
Rascon15        TGAATCATTTCTTAGACTATAATTCATTCACAGTAAATACCTTATACTTTTTCATGGCGA      3637
Rascon13        TGAATCATTTCTTAGACTATAATTCATTCACAGTAAATACCTTATACTTTTTCATGGCGA      3645
Rascon6         TGAATCATTTCTTAGACTATAATTCATTCACAGTAAATACCTTATACTTTTTCATGGCGA      3644
Pachon14        TGAATCATTTCTTAGACTATAATTCATTCACAGTAAATACCTTATACTTTTTCATGGCGA      3644
Pachon9         TGAATCATTTCTTAGACTATAATTCATTCACAGTAAATACCTTATACTTTTTCATGGCGA      3652
Pachon17        TGAATCATTTCTTAGACTATAATTCATTCACAGTAAATACCTTATACTTTTTCATGGCGA      3651
Pachon12        TGAATCATTTCTTAGACTATAATTCATTCACAGTAAATACCTTATACTTTTTCATGGCGA      3651
Pachon11        TGAATCATTTCTTAGACTATAATTCATTCACAGTAAATACCTTATACTTTTTCATGGCGA      3651
Pachon7         TGAATCATTTCTTAGACTATAATTCATTCACAGTAAATACCTTATACTTTTTCATGGCGA      3651
Pachon3         TGAATCATTTCTTAGACTATAATTCATTCACAGTAAATACCTTATACTTTTTCATGGCGA      3651
Pachon8         TGAATCATTTCTTAGACTATAATTCATTCACAGTAAATACCTTATACTTTTTCATGGCGA      3651
Pachon15        TGAATCATTTCTTAGACTATAATTCATTCACAGTAAATACCTTATACTTTTTCATGGCGA      3651
                ************************************************************

Rascon4         CTACAGATTCATTTACAGGGGTTACTGTTTCGACCAACATATGTACTGAATAAATTATTA      3661
Surface         CTACAGATTCATTTACAGGGGTTACTGTTTCGACCAACATATGTACTGAATAAATGATTA      3718
Rascon8         CTACAGATTCATTTACAGGGGTTACTGTTTCGACCAACATATGTACTGAATAAATTATTA      3703
Rascon2         CTACAGATTCATTTACAGGGGTTACTGTTTCGACCAACATATGTACTGAATAAATTATTA      3709
Rascon15        CTACAGATTCATTTACAGGGGTTACTGTTTCGACCAACATATGTACTGAATAAATTATTA      3697
Rascon13        CTACAGATTCATTTACAGGGGTTACTGTTTCGACCAACATATGTACTGAATAAATTATTA      3705
Rascon6         CTACAGATTCATTTACAGGGGTTACTGTTTCGACCAACATATGTACTGAATAAATTATTA      3704
Pachon14        CTACAGATTCATTTACAGGGGTTACTGTTTCGACCAACATATGTACTGAATAAATTATTA      3704
Pachon9         CTACAGATTCATTTACAGGGGTTACTGTTTCGACCAACATATGTACTGAATAAATTATTA      3712
Pachon17        CTACAGATTCATTTACAGGGGTTACTGTTTCGACCAACATATGTACTGAATAAATTATTA      3711
Pachon12        CTACAGATTCATTTACAGGGGTTACTGTTTCGACCAACATATGTACTGAATAAATTATTA      3711
Pachon11        CTACAGATTCATTTACAGGGGTTACTGTTTCGACCAACATATGTACTGAATAAATTATTA      3711
Pachon7         CTACAGATTCATTTACAGGGGTTACTGTTTCGACCAACATATGTACTGAATAAATTATTA      3711
Pachon3         CTACAGATTCATTTACAGGGGTTACTGTTTCGACCAACATATGTACTGAATAAATTATTA      3711
Pachon8         CTACAGATTCATTTACAGGGGTTACTGTTTCGACCAACATATGTACTGAATAAATTATTA      3711
Pachon15        CTACAGATTCATTTACAGGGGTTACTGTTTCGACCAACATATGTACTGAATAAATTATTA      3711
                ******************************************************* ****

Rascon4         TAGACCCTACATTCTGAATAACTCATATTAGACACTGAATAAATTCTAGTAAATACTGAA      3721
Surface         TAGACACTACATTCTGAATAACTCATATTAGACACTGAATAAATTCTAGTAAATACTGAA      3778
Rascon8         TAGACCCTACATTCTGAATAACTCATATTAGACACTGAATAAATTCTAGTAAATACTGAA      3763
Rascon2         TAGACACTACATTCTGAATAACTCATATTAGACACTGAATAAATTCTAGTAAATACTGAA      3769
Rascon15        TAGACACTACATTCTGAATAACTCATATTAGACACTGAATAAATTCTAGTAAATACTGAA      3757
Rascon13        TAGACACTACATTCTGAATAACTCATATTAGACACTGAATAAATTCTAGTAAATACTGAA      3765
Rascon6         TAGACCCTTCATTCTGAATAACTCATATTAGACACTGAATAAATTCTAGTAAATACTGAA      3764
Pachon14        TAGACCCTACATTCTGAATAACTCATATTAGACACTGAATAAATTCTAGTAAATACTGAA      3764
Pachon9         TAGACCCTACATTCTGAATAACTCATATTAGACACTGAATAAATTCTAGTAAATACTGAA      3772
Pachon17        TAGACCCTACATTCTGAATAACTCATATTAGACACTGAATAAATTCTAGTAAATACTGAA      3771
Pachon12        TAGACCCTACATTCTGAATAACTCATATTAGACACTGAATAAATTCTAGTAAATACTGAA      3771
Pachon11        TAGACCCTACATTCTGAATAACTCATATTAGACACTGAATAAATTCTAGTAAATACTGAA      3771
Pachon7         TAGACCCTACATTCTGAATAACTCATATTAGACACTGAATAAATTCTAGTAAATACTGAA      3771
Pachon3         TAGACCCTACATTCTGAATAACTCATATTAGACACTGAATAAATTCTAGTAAATACTGAA      3771
Pachon8         TAGACCCTACATTCTGAATAACTCATATTAGACACTGAATAAATTCTAGTAAATACTGAA      3771
Pachon15        TAGACCCTACATTCTGAATAACTCATATTAGACACTGAATAAATTCTAGTAAATACTGAA      3771
                ***** ** ***************************************************

Rascon4         TAATTCATAATAGATACTGAATAATTGATTGTAGATACTGA-------------------      3762
Surface         TAATTCATAATAGATACTGAATAATTGATTGTAGATACTGAGTAATTAATTGTAGATACT      3838
Rascon8         TAATTCATAATAGATACTGAATAATTGATTGTAGATACTGAGTAATTAATTGTAGATACT      3823
Rascon2         TAATTCATAATAGATACTGAATAATTGATTGTAGATACTGAGTAATTAATTGTAGATACT      3829
Rascon15        TAATTCATCATGGATACTGAATAATTGATTGTAGATACTGAGTAATTAATTGTAGATACT      3817
Rascon13        TAATTCATAATAGATACTGAATAATTGATTGTAGATACTGAGTAATTAATTGTAGATACT      3825
Rascon6         TAATTCATAATAGATACTGAATAATTGATTGTAGATACTGAGTAATTAATTGTAGATACT      3824
Pachon14        TAATTCATAATAGATACTGAATAATTGATTGTAGATACTGAGTAATTAATTGTAGATACT      3824
Pachon9         TAATTCATCATGGATACTGAATAATTGATTGTAGATACTGAGTAATTAATTGTAGATACT      3832
Pachon17        TAATTCATCATGGATACTGAATAATTGATTGTAGATACTGAGTAATTAATTGTAGATACT      3831
Pachon12        TAATTCATCATGGATACTGAATAATTGATTGTAGATACTGAGTAATTAATTGTAGATACT      3831
Pachon11        TAATTCATAATAGATACTGAATAATTGATTGTAGATACTGAGTAATTAATTGTAGATACT      3831
Pachon7         TAATTCATCATGGATACTGAATAATTGATTGTAGATACTGAGTAATTAATTGTAGATACT      3831
Pachon3         TAATTCATAATAGATACTGAATAATTGATTGTAGATACTGAGTAATTAATTGTAGATACT      3831
Pachon8         TAATTCATCATAGATACTGAATAATTGATTGTAGATACTGAGTAATTAATTGTAGATACT      3831
Pachon15        TAATTCATCATGGATACTGAATAATTGATTGTAGATACTGAGTAATTAATTGTAGATACT      3831
```

```
                    ******** ** *****************************

Rascon4             --ATAAATGACAGTAGATACTGAAAACTAACGAAAGGTACAAGGAATATTTTTTTGGTGC     3820
Surface             ACATAAATGACAGTAGATACTGAAAACTAACGAAAGGTACAAGGAATATTTTTTTGGTGC     3898
Rascon8             AAATAAATGACAGTAGATACTGAAAACTAACGAAAGGTACAAGGAATATTTTTTTGGTGC     3883
Rascon2             AAATAAATGACAGTAGATACTGAAAACTAACGAAAGGTACAAGGAATATTTTTTTGGTGC     3889
Rascon15            AAATAAATGACAGTAGATACTGAAAACTAACGAAAGGTACAAGGAATATTTTTTTGGTGC     3877
Rascon13            AAATAAATGACAGTAGATACTGAAAACTAACGAAAGGTACAAGGAATATTTTTTTGGTGC     3885
Rascon6             AAATAAATGACAGTAGATACTGAACACTAACGAAAAGTACAAGGAATATTTTTTTGGTGC     3884
Pachon14            AAATAAATGACAGTAGATACTGAAAACTAACGAAAGGTACAAGGAATATTTTTTTGGTGC     3884
Pachon9             AAATAAATGACAGTAGATACTGAAAACTAACGAAAGGTACAAGGAATATTTTTTTGGTGC     3892
Pachon17            AAATAAATGACAGTAGATACTGAAAACTAACGAAAGGTACAAGGAATATTTTTTTGGTGC     3891
Pachon12            AAATAAATGACAGTAGATACTGAAAACTAACGAAAGGTACAAGGAATATTTTTTTGGTGC     3891
Pachon11            AAATAAATGACAGTAGATACTGAAAACTAACGAAAGGTACAAGGAATATTTTTTTGGTGC     3891
Pachon7             AAATAAATGACAGTAGATACTGAAAACTAACGAAAGGTACAAGGAATATTTTTTTGGTGC     3891
Pachon3             AAATAAATGACAGTAGATACTGAAAACTAACGAAAGGTACAAGGAATATTTTTTTGGTGC     3891
Pachon8             AAATAAATGACAGTAGATACTGAAAACTAACGAAAGGTACAAGGAATATTTTTTTGGTGC     3891
Pachon15            AAATAAATGACAGTAGATACTGAAAACTAACGAAAGGTACAAGGAATATTTTTTTGGTGC     3891
                      ********************** ********** ************************

Rascon4             AGGTATCAACATTTAAATAGAATAGAATATATTAGTAGATGGATAGAAAGAAAAAAAATC     3880
Surface             AGGTATCAACATTTAAATAGAATAGAATATATTAGTAGATGGATAGAAAGAAAAAAAATC     3958
Rascon8             AGGTATCAACATTTAAATAGAATAGAATATATTAGTAGATGGATAGAAAGAAAAAAAATC     3943
Rascon2             AGGTATCAACATTTAAATAGAATAGAATATATTAGTAGATGGATAGAAAGAAAAAAAATC     3949
Rascon15            AGGTATCAACATTTAAATAGAATATAATATATTAGTAGATGGATAGAAAGAAAAAAAATC     3937
Rascon13            AGGTATCAACATTTAAATAGAATAGAATATATTAGTAGATGGATAGAAAGAAAAAAAATC     3945
Rascon6             AGGTATCAACATTTAAATAGAATAGAATATATTAGTAGATGGATAGAAAGAAAAAAAATC     3944
Pachon14            AGGTATCAACATTTAAATAGAATAGAATATATTAGTAGATGGATAGAAAGAAAAAAAATC     3944
Pachon9             AGGTATCAACATTTAAATAGAATAGAATATATTAGTAGATGGATAGAAAGAAAAAAAATC     3952
Pachon17            AGGTATCAACATTTAAATAGAATAGAATATATTAGTAGATGGATAGAAAGAAAAAAAATC     3951
Pachon12            AGGTATCAACATTTAAATAGAATAGAATATATTAGTAGATGGATAGAAAGAAAAAAAATC     3951
Pachon11            AGGTATCAACATTTAAATAGAATAGAATATATTAGTAGATGGATAGAAAGAAAAAAAATC     3951
Pachon7             AGGTATCAACATTTAAATAGAATAGAATATATTAGTAGATGGATAGAAAGAAAAAAAATC     3951
Pachon3             AGGTATCAACATTTAAATAGAATAGAATATATTAGTAGATGGATAGAAAGAAAAAAAATC     3951
Pachon8             AGGTATCAACATTTAAATAGAATAGAATATATTAGTAGATGGATAGAAAGAAAAAAAATC     3951
Pachon15            AGGTATCAACATTTAAATAGAATAGAATATATTAGTAGATGGATAGAAAGAAAAAAAATC     3951
                    ************************ ***********************************

Rascon4             TGAATTATATAAACATGAATTAAAGTACACACACAAAAGTAAATGTATTATGTAAAATGT     3940
Surface             TGAATTATATAAACATGAATTAAAGTACACACACAAAAGTAAATGTATTATGTAAAATGT     4018
Rascon8             TGAATTATATAAACATGAATTAAAGTACACACACAAAAGTAAATGTATTATGTAAAATGT     4003
Rascon2             TGAATTATATAAACATGAATTAAAGTACACACACAAAAGTAAATGTATTATGTAAAATGT     4009
Rascon15            TGAATTATATAAACATGAATTAAAGTACACACACAAAAGTAAATGTATTATGTAAAATGT     3997
Rascon13            TGAATTATATAAACATGAATTAAAGTACACACACAAAAGTAAATGTATTATGTAAAATGT     4005
Rascon6             TGAATTATATAAACATGAATTAAAGTACACACACAAAAGTAAATGTATTATGTAAAATGT     4004
Pachon14            TGAATTATATAAACATGAATTAAAGTACACACACAAAAGTAAATGTATTATGTAAAATGT     4004
Pachon9             TGAATTATATAAACATGAATTAAAGTACACACACAAAAGTAAATGTATTATGTAAAATGT     4012
Pachon17            TGAATTATATAAACATGAATTAAAGTACACACACAAAAGTAAATGTATTATGTAAAATGT     4011
Pachon12            TGAATTATATAAACATGAATTAAAGTACACACACAAAAGTAAATGTATTATGTAAAATGT     4011
Pachon11            TGAATTATATAAACATGAATTAAAGTACACACACAAAAGTAAATGTATTATGTAAAATGT     4011
Pachon7             TGAATTATATAAACATGAATTAAAGTACACACACAAAAGTAAATGTATTATGTAAAATGT     4011
Pachon3             TGAATTATATAAACATGAATTAAAGTACACACACAAAAGTAAATGTATTATGTAAAATGT     4011
Pachon8             TGAATTATATAAACATGAATTAAAGTACACACACAAAAGTAAATGTATTATGTAAAATGT     4011
Pachon15            TGAATTATATAAACATGAATTAAAGTACACACACAAAAGTAAATGTATTATGTAAAATGT     4011
                    ************************************************************

Rascon4             ATTATAAGTATTGTATACTTTACTGCAGTTCTAAAGAACTGATTGTAGATACTGAATAAA     4000
Surface             ATTATAAGTATTGTATACTTTACTGCAGTTCTAAAGAACTGATTGTAGATACTGAATAAA     4078
Rascon8             ATTATAAGTATTGTATACTTTACTGCAGTTCTAAAGAACTGATTGTAGATACTGAATAAA     4063
Rascon2             ATTATAAGTATTGTATACTTTACTGCAGTTCTAAAGAACTGATTGTAGATACTGAATAAA     4069
Rascon15            ATTATAAGTATTGTATACTTTACTGCAGTTCTAAAGAACTGATTGTAGATACTGAATAAA     4057
Rascon13            ATTATAAGTATTGTATACTTTACTGCAGTTCTAAAGAACTGATTGTAGATACTGAATAAA     4065
Rascon6             ATTATAAGTATTGTATACTTTACTGCAGTTCTAAAGAACTGATTGTAGATACTGAATAAA     4064
Pachon14            ATTATAAGTATTGTATACTTTACTGCAGTTCTAAAGAACTGATTGTAGATACTGAATAAA     4064
Pachon9             ATTATAAGTATTGTATACTTTACTGCAGTTCTAAAGAACTGATTGTAGATACTGAATAAA     4072
Pachon17            ATTATAAGTATTGTATACTTTACTGCAGTTCTAAAGAACTGATTGTAGATACTGAATAAA     4071
Pachon12            ATTATAAGTATTGTATACTTTACTGCAGTTCTAAAGAACTGATTGTAGATACTGAATAAA     4071
Pachon11            ATTATAAGTATTGTATACTTTACTGCAGTTCTAAAGAACTGATTGTAGATACTGAATAAA     4071
Pachon7             ATTATAAGTATTGTATACTTTACTGCAGTTCTAAAGAACTGATTGTAGATACTGAATAAA     4071
Pachon3             ATTATAAGTATTGTATACTTTACTGCAGTTCTAAAGAACTGATTGTAGATACTGAATAAA     4071
Pachon8             ATTATAAGTATTGTATACTTTACTGCAGTTCTAAAGAACTGATTGTAGATACTGAATAAA     4071
Pachon15            ATTATAAGTATTGTATACTTTACTGCAGTTCTAAAGAACTGATTGTAGATACTGAATAAA     4071
                    ************************************************************

Rascon4             TGTTTGTCCTAAATGTGCTAAAATTCTCTTTTAAAAATGTTACATCCACAGAGTTAATGT     4060
Surface             TGTTTGTCCTAAATGTGCTAAAATTCTCTTTTAAAAATGTTACATCCACACAGTTAATGT     4138
Rascon8             TGTTTGTCCTAAATGTGCTAAAATTCTCTTTTAAAAATGTTACATCCACACAGTTAATGT     4123
```

```
Rascon2         TGTTTGTCCTAAATGTGCTAAAATTCTCTTTTAAAAATGTTACATCCACACAGTTAATGT        4129
Rascon15        TGTTTGTCCTAAATGTGCTAAAATTCTCTTTTAAAAATGTTACATCCACAGAGTTAATGT        4117
Rascon13        TGTTTGTCCTAAATGTGCTAAAATTCTCTTTTAAAAATGTTACATCCACAGAGTTAATGT        4125
Rascon6         TGTTTGTCCTAAATGTGCTAAAATTCTCTTTTAAAAATGTTACATCCACAGAGTTAATGT        4124
Pachon14        TGTTTGTCCTAAATGTGCTAAAATTCTCTTTTAAAAATGTTACATCCACACAGTTAATGT        4124
Pachon9         TGTTTGTCCTAAATGTGCTAAAATTCTCTTTTAAAAATGTTACATCCACACAGTTAATGT        4132
Pachon17        TGTTTGTCCTAAATGTGCTAAAATTCTCTTTTAAAAATGTTACATCCACACAGTTAATGT        4131
Pachon12        TGTTTGTCCTAAATGTGCTAAAATTCTCTTTTAAAAATGTTACATCCACACAGTTAATGT        4131
Pachon11        TGTTTGTCCTAAATGTGCTAAAATTCTCTTTTAAAAATGTTACATCCACACAGTTAATGT        4131
Pachon7         TGTTTGTCCTAAATGTGCTAAAATTCTCTTTTAAAAATGTTACATCCACACAGTTAATGT        4131
Pachon3         TGTTTGTCCTAAATGTGCTAAAATTCTCTTTTAAAAATGTTACATCCACACAGTTAATGT        4131
Pachon8         TGTTTGTCCTAAATGTGCTAAAATTCTCTTTTAAAAATGTTACATCCACACAGTTAATGT        4131
Pachon15        TGTTTGTCCTAAATGTGCTAAAATTCTCTTTTAAAAATGTTACATCCACACAGTTAATGT        4131
                ************************************************** *********

Rascon4         CTCTCAGTTTACTCACATGTGAATAAAACTAATG---TTTTTTTAGTCTAATTCCTAATC        4117
Surface         CTCTCAGTTTACTCACATGTGAATAAAACTAATGTTTTTTTTTTAGTCTAATTCCTAATC        4198
Rascon8         CTCTCAGTTTACTCACATGTGAATAAAACTAATG---TTTTTTTAGTCTAATTCCTAATC        4180
Rascon2         CTCTCAGTTTACTCACATGTGAATAAAACTAATG---TTTTTTTAGTCTAATTCCTAATC        4186
Rascon15        CTCTCAGTTTACTCACATGTGAATAAAACTAATG---TTTTTTTAGTCTAATTCCTAATC        4174
Rascon13        CTCTCAGTTTACTCACATGTGAATAAAACTAATG---TTTTTTTAGTCTAATTCCTAATC        4182
Rascon6         CTCTCAGTTTACTCACATGTGAATAAAACTAATG---TTTTTTTAGTCTAATTCCTAATC        4181
Pachon14        CTCTCAGTTTACTCACATGTGAATAAAACTAATG---TTTTTTTAGTCTAATTCCTAATC        4181
Pachon9         CTCTCAGTTTACTCACATGTGAATAAAACTAATG---TTTTTTTAGTCTAATTCCTAATC        4189
Pachon17        CTCTCAGTTTACTCACATGTGAATAAAACTAATG---TTTTTTTAGTCTAATTCCTAATC        4188
Pachon12        CTCTCAGTTTACTCACATGTGAATAAAACTAATG---TTTTTTTAGTCTAATTCCTAATC        4188
Pachon11        CTCTCAGTTTACTCACATGTGAATAAAACTAATG---TTTTTTTAGTCTAATTCCTAATC        4188
Pachon7         CTCTCAGTTTACTCACATGTGAATAAAACTAATG---TTTTTTTAGTCTAATTCCTAATC        4188
Pachon3         CTCTCAGTTTACTCACATGTGAATAAAACTAATG---TTTTTTTAGTCTAATTCCTAATC        4188
Pachon8         CTCTCAGTTTACTCACATGTGAATAAAACTAATG---TTTTTTTAGTCTAATTCCTAATC        4188
Pachon15        CTCTCAGTTTACTCACATGTGAATAAAACTAATG---TTTTTTTAGTCTAATTCCTAATC        4188
                **********************************   ***********************

Rascon4         TGGAATATCTGAAGAGCAGCTCATGGTTTGGGGGGTTAATGCGCAGTAATCTGAGATTGG        4177
Surface         TGGAATATCTGAAGAGCAGCTCATGGTTTGGGGGGTTAATGCGCAGTAATCTGAGATTGG        4258
Rascon8         TGGAATATCTGAAGAGCAGCTCATGGTTTGGGGGGTTAATGCGCAGTAATCTGAGATTGG        4240
Rascon2         TGGAATATCTGAAGAGCAGCTCATGGTTTGGGGGGTTAATGCGCAGTAATCTGAGATTGG        4246
Rascon15        TGGAATATCTGAAGAGCAGCTCATGGTTTGGGGGGTTAATGCGCAGTAATCTGAGATTGG        4234
Rascon13        TGGAATATCTGAAGAGCAGCTCATGGTTTGGGGGGTTAATGCGCAGTAATCTGAGATTGG        4242
Rascon6         TGGAATATCTGAAGAGCAGCTCATGGTTTGGGGGGTTAATGCGCAGTAATCTGAGATTGG        4241
Pachon14        TGGAATATCTGAAGAGCAGCTCATGGTTTGGGGGGTTAATGCGCAGTAATCTGAGATTGG        4241
Pachon9         TGGAATATCTGAAGAGCAGCTCATGGTTTGGGGGGTTAATGCGCAGTAATCTGAGATTGG        4249
Pachon17        TGGAATATCTGAAGAGCAGCTCATGGTTTGGGGGGTTAATGCGCAGTAATCTGAGATTGG        4248
Pachon12        TGGAATATCTGAAGAGCAGCTCATGGTTTGGGGGGTTAATGCGCAGTAATCTGAGATTGG        4248
Pachon11        TGGAATATCTGAAGAGCAGCTCATGGTTTGGGGGGTTAATGCGCAGTAATCTGAGATTGG        4248
Pachon7         TGGAATATCTGAAGAGCAGCTCATGGTTTGGGGGGTTAATGCGCAGTAATCTGAGATTGG        4248
Pachon3         TGGAATATCTGAAGAGCAGCTCATGGTTTGGGGGGTTAATGCGCAGTAATCTGAGATTGG        4248
Pachon8         TGGAATATCTGAAGAGCAGCTCATGGTTTGGGGGGTTAATGCGCAGTAATCTGAGATTGG        4248
Pachon15        TGGAATATCTGAAGAGCAGCTCATGGTTTGGGGGGTTAATGCGCAGTAATCTGAGATTGG        4248
                ************************************************************

Rascon4         AGAAGTGTGGTTTTACTCCACCTTCAGCGAGATATGGAGATTGAAGAAGTGATTTAATAT        4237
Surface         AGAAGTGTGGTTTTACTCCACCTTCAGCGAGATATGGAGATTGAAGAAGTGATTTAATAT        4318
Rascon8         AGAAGTGTGGTTTTACTCCACCTTCAGCGAGATATGGAGATTGAAGAAGTGATTTAATAT        4300
Rascon2         AGAAGTGTGGTTTTACTCCACCTTCAGCAAGATATGGAGATTGAAGAAGTGATTTAATAT        4306
Rascon15        AGAAGTGTGGTTTTACTCCACCTTCAGCGAGATATGGAGATTGAAGAAGTGATTTAATAT        4294
Rascon13        AGAAGTGTGGTTTTACTCCACCTTCAGCGAGATATGGAGATTGAAGAAGTGATTTAATAT        4302
Rascon6         AGAAGTGTGGTTTTACTCCACCTTCAGCGAGATATGGAGATTGAAGAAGTGATTTAATAT        4301
Pachon14        AGAAGTGTGGTTTTACTCCACCTTCAGCGAGATATGGAGATTGAAGAAGTGATTTAATAT        4301
Pachon9         AGAAGTGTGGTTTTACTCCACCTTCAGCGAGATATGGAGATTGAAGAAGTGATTTAATAT        4309
Pachon17        AGAAGTGTGGTTTTACTCCACCTTCAGCGAGATATGGAGATTGAAGAAGTGATTTAATAT        4308
Pachon12        AGAAGTGTGGTTTTACTCCACCTTCAGCGAGATATGGAGATTGAAGAAGTGATTTAATAT        4308
Pachon11        AGAAGTGTGGTTTTACTCCACCTTCAGCGAGATATGGAGATTGAAGAAGTGATTTAATAT        4308
Pachon7         AGAAGTGTGGTTTTACTCCACCTTCAGCGAGATATGGAGATTGAAGAAGTGATTTAATAT        4308
Pachon3         AGAAGTGTGGTTTTACTCCACCTTCAGCGAGATATGGAGATTGAAGAAGTGATTTAATAT        4308
Pachon8         AGAAGTGTGGTTTTACTCCACCTTCAGCGAGATATGGAGATTGAAGAAGTGATTTAATAT        4308
Pachon15        AGAAGTGTGGTTTTACTCCACCTTCAGCGAGATATGGAGATTGAAGAAGTGATTTAATAT        4308
                **************************** *******************************

Rascon4         TGCTGTAAGGCTGTTTAAGGGTGTTAAGGGTGTTGGGGAGGAGTCTCGAACCCTTCCAGC        4297
Surface         TGCTGTAAGGCTGTTTAAGGGTGTTAAGGGTGTTGGGGAGGAGTCTCGAACCCTTCCAGC        4378
Rascon8         TGCTGTAAGGCTGTTTAAGGGTGTTAAGGGTGTTGGGGAGGAGTCTCGAACCCTTCCAGC        4360
Rascon2         TGCTGTAAGGCTGTTTAAGGGTGTTAAGGGTGTTGGGGAGGAGTCTCGAACCCTTCCAGC        4366
Rascon15        TGCTGTAAGGCTGTTTAAGGGTGTTAAGGGTGTTGGGGAGGAGTCTCGAACCCTTCCAGC        4354
Rascon13        TGCTGTAAGGCTGTTTAAGGGTGTTAAGGGTGTTGGGGAGGAGTCTCGAACCCTTCCAGC        4362
Rascon6         TGCTGTAAGGCTGTTTAAGGGTGTTAAGGGTGTTGGGGAGGAGTCTTGAACTCTTCCAGC        4361
Pachon14        TGCTGTAAGGCTGTTTAAGGGTGTTAAGGGTGTTGGGGAGGAGTCTCGAACCCTTCCAGC        4361
```

```
Pachon9      TGCTGTAAGGCTGTTTAAGGGTGTTAAGGGTGTTGGGGAGGAGTCTCGAACCCTTCCAGC   4369
Pachon17     TGCTGTAAGGCTGTTTAAGGGTGTTAAGGGTGTTGGGGAGGAGTCTCGAACCCTTCCAGC   4368
Pachon12     TGCTGTAAGGCTGTTTAAGGGTGTTAAGGGTGTTGGGGAGGAGTCTCGAACCCTTCCAGC   4368
Pachon11     TGCTGTAAGGCTGTTTAAGGGTGTTAAGGGTGTTGGGGAGGAGTCTCGAACCCTTCCAGC   4368
Pachon7      TGCTGTAAGGCTGTTTAAGGGTGTTAAGGGTGTTGGGGAGGAGTCTCGAACCCTTCCAGC   4368
Pachon3      TGCTGTAAGGCTGTTTAAGGGTGTTAAGGGTGTTGGGGAGGAGTCTCGAACCCTTCCAGC   4368
Pachon8      TGCTGTAAGGCTGTTTAAGGGTGTTAAGGGTGTTGGGGAGGAGTCTCGAACCCTTCCAGC   4368
Pachon15     TGCTGTAAGGCTGTTTAAGGGTGTTAAGGGTGTTGGGGAGGAGTCTCGAACCCTTCCAGC   4368
             ********************************************** **** ********

Rascon4      CAATCTGAATCCTCGCTGAGGCCCCAGGGGGGATGAGATACATACAAAAGCCCCCTGGCT   4357
Surface      CAATCTGAATCCTCGCTGAGGCCCCAGGGGGGATGAGATACACACAAAAGCCCCCTGGCT   4438
Rascon8      CAATCTGAATCCTCGCTGAGGCCCCAGGGGGGATGAGATACACACAAAAGCCCCCTGGCT   4420
Rascon2      CAATCTGAATCCTCGCTGAGGCCCCAGGGGGGATGAGATACACACAAAAGCCCCCTGGCT   4426
Rascon15     CAATCTGAATCCTCGCTGAGGCCCCAGGGGGGATGAGATACACACAAAAGCCCCCTGGCT   4414
Rascon13     CAATCTGAATCCTCGCTGAGGCCCCAGGGGGGATGAGATACACACAAAAGCCCCCTGGCT   4422
Rascon6      CAATCTGAATCCTCGCTGAGGCCCCAGGGGGGATGAGATACATACAAAAGCCCCCTGGCT   4421
Pachon14     CAATCTGAATCCTCGCTGAGGCCCCAGGGGGGATGAGATACACACAAAAGCCCCCTGGCT   4421
Pachon9      CAATCTGAATCCTCGCTGAGGCCCCAGGGGGGATGAGATACACACAAAAGCCCCCTGGCT   4429
Pachon17     CAATCTGAATCCTCGCTGAGGCCCCAGGGGGGATGAGATACACACAAAAGCCCCCTGGCT   4428
Pachon12     CAATCTGAATCCTCGCTGAGGCCCCAGGGGGGATGAGATACACACAAAAGCCCCCTGGCT   4428
Pachon11     CAATCTGAATCCTCGCTGAGGCCCCAGGGGGGATGAGATACACACAAAAGCCCCCTGGCT   4428
Pachon7      CAATCTGAATCCTCGCTGAGGCCCCAGGGGGGATGAGATACACACAAAAGCCCCCTGGCT   4428
Pachon3      CAATCTGAATCCTCGCTGAGGCCCCAGGGGGGATGAGATACACACAAAAGCCCCCTGGCT   4428
Pachon8      CAATCTGAATCCTCGCTGAGGCCCCAGGGGGGATGAGATACACACAAAAGCCCCCTGGCT   4428
Pachon15     CAATCTGAATCCTCGCTGAGGCCCCAGGGGGGATGAGATACACACAAAAGCCCCCTGGCT   4428
             ****************************************** *****************

Rascon4      GAGGGGTCTGTATAAAAGCCCCCAGAGGACACTCAATCGCCACTGCGATCAGCAGCTGGA   4417 Rx3
Surface      GAGGGGTCTGTATAAAAGCCCCCAGAGGACACTCAATCGCCACTGCGATCAGCAGCTGGA   4498 5′-UTR
Rascon8      GAGGGGTCTGTATAAAAGCCCCCAGAGGACACTCAATCGCCACTGCGATCAGCAGCTGGA   4480
Rascon2      GAGGGGTCTGTATAAAAGCCCCCAGAGGACACTCAATCGCCACTGCGATCAGCAGCTGGA   4486
Rascon15     GAGGGGTCTGTATAAAAGCCCCCAGAGGACACTCAATCGCCACTGCGATCAGCAGCTGGA   4474
Rascon13     GAGGGGTCTGTATAAAAGCCCCCAGAGGACACTCAATCGCCACTGCGATCAGCAGCTGGA   4482
Rascon6      GAGGGGTCTGTATAAAAGCCCCCAGAGGACACTCAATCGCCACTGCGATCAGCAGCTGGA   4481
Pachon14     GAGGGGTCTGTATAAAAGCCCCCAGAGGACACTCAATCGCCACTGCGATCAGCAGCTGGA   4481
Pachon9      GAGGGGTCTGTATAAAAGCCCCCAGAGGACACTCAATCGCCACTGCGATCAGCAGCTGGA   4489
Pachon17     GAGGGGTCTGTATAAAAGCCCCCAGAGGACACTCAATCGCCACTGCGATCAGCAGCTGGA   4488
Pachon12     GAGGGGTCTGTATAAAAGCCCCCAGAGGACACTCAATCGCCACTGCGATCAGCAGCTGGA   4488
Pachon11     GAGGGGTCTGTATAAAAGCCCCCAGAGGACACTCAATCGCCACTGCGATCAGCAGCTGGA   4488
Pachon7      GAGGGGTCTGTATAAAAGCCCCCAGAGGACACTCAATCGCCACTGCGATCAGCAGCTGGA   4488
Pachon3      GAGGGGTCTGTATAAAAGCCCCCAGAGGACACTCAATCGCCACTGCGATCAGCAGCTGGA   4488
Pachon8      GAGGGGTCTGTATAAAAGCCCCCAGAGGACACTCAATCGCCACTGCGATCAGCAGCTGGA   4488
Pachon15     GAGGGGTCTGTATAAAAGCCCCCAGAGGACACTCAATCGCCACTGCGATCAGCAGCTGGA   4488
             ************************************************************

Rascon4      CCTAGGTGATGCCTTCAGTCTCTGGAGTGACCAACAGTTTTCTACAGCAGTGGAGCGACT   4477
Surface      CCTAGGTGATGCCTTCAGTCTCTGGAGTGACCAACAGTTTTCTACAGCAGTGGAGCGACT   4558
Rascon8      CCTAGGTGATGCCTTCAGTCTCTGGAGTGACCAACAGTTTTCTACAGCAGTGGAGCGACT   4540
Rascon2      CCTAGGTGATGCCTTCAGTCTCTGGAGTGACCAACAGTTTTCTACAGCAGTGGAGCGACT   4546
Rascon15     CCTAGGTGATGCCTTCAGTCTCTGGAGTGACCAACAGTTTTCTACAGCAGTGGAGCGACT   4534
Rascon13     CCTAGGTGATGCCTTCAGTCTCTGGAGTGACCAACAGTTTTCTACAGCAGTGGAGCGACT   4542
Rascon6      CCTAGGTGATGCCTTCAGTCTCTGGAGTGACCAACAGTTTTCTACAGCAGTGGAGCGACT   4541
Pachon14     CCTAGGTGATGCCTTCAGTCTCTGGAGTGACCAACAGTTTTCTACAGCAGTGGAGCGACT   4541
Pachon9      CCTAGGTGATGCCTTCAGTCTCTGGAGTGACCAACAGTTTTCTACAGCAGTGGAGCGACT   4549
Pachon17     CCTAGGTGATGCCTTCAGTCTCTGGAGTGACCAACAGTTTTCTACAGCAGTGGAGCGACT   4548
Pachon12     CCTAGGTGATGCCTTCAGTCTCTGGAGTGACCAACAGTTTTCTACAGCAGTGGAGCGACT   4548
Pachon11     CCTAGGTGATGCCTTCAGTCTCTGGAGTGACCAACAGTTTTCTACAGCAGTGGAGCGACT   4548
Pachon7      CCTAGGTGATGCCTTCAGTCTCTGGAGTGACCAACAGTTTTCTACAGCAGTGGAGCGACT   4548
Pachon3      CCTAGGTGATGCCTTCAGTCTCTGGAGTGACCAACAGTTTTCTACAGCAGTGGAGCGACT   4548
Pachon8      CCTAGGTGATGCCTTCAGTCTCTGGAGTGACCAACAGTTTTCTACAGCAGTGGAGCGACT   4548
Pachon15     CCTAGGTGATGCCTTCAGTCTCTGGAGTGACCAACAGTTTTCTACAGCAGTGGAGCGACT   4548
             ************************************************************

Rascon4      TTTGTCAATTTTTTGTTTTTAATTTTGTCAAGAAATTTTGATCTCTTACGAAATTCGGAA   4537
Surface      TTTGTCAATTTTTTGTTTTTAATTTTGTCAAGAAATTTTGATCTCTCACGAAATTCGGAA   4618
Rascon8      TTTGTCAATTTTTTGTTTTTAATTTTGTCAAGAAATTTTGATCTCTTACGAAATTCGGAA   4600
Rascon2      TTTGTCAATTTTTTGTTTTTAATTTTGTCAAGAAATTTTGATCTCTTACGAAATTCGGAA   4606
Rascon15     TTTGTCAATTTTTTGTTTTTAATTTTGTCAAGAAATTTTGATCTCTTACGAAATTCGGAA   4594
Rascon13     TTTGTAAATTTTTTGTTTTTAATTTTGTCAAGAAATTTTGATCTCTTACAAAATTCGGAA   4602
Rascon6      TTTGTCAATTTTTTGTTTTTAATTTTGTCAAGAAATTTTGATCTCTTACGAAATTCGGAA   4601
Pachon14     TTTGTCAATTTTTTGTTTTTAATTTTGTCAAGAAATTTTGATCTCTTACGAAATTCGGAA   4601
Pachon9      TTTGTCAATTTTTTGTTTTTAATTTTGTCAAGAAATTTTGATCTCTTACGAAATTCGGAA   4609
Pachon17     TTTGTCAATTTTTTGTTTTTAATTTTGTCAAGAAATTTTGATCTCTTACGAAATTCGGAA   4608
Pachon12     TTTGTCAATTTTTTGTTTTTAATTTTGTCAAGAAATTTTGATCTCTTACGAAATTCGGAA   4608
Pachon11     TTTGTCAATTTTTTGTTTTTAATTTTGTCAAGAAATTTTGATCTCTTACGAAATTCGGAA   4608
Pachon7      TTTGTCAATTTTTTGTTTTTAATTTTGTCAAGAAATTTTGATCTCTTACGAAATTCGGAA   4608
```

```
Pachon3      TTTGTCAATTTTTTGTTTTTAATTTTGTCAAGAAATTTTGATCTCTTACGAAATTCGGAA   4608
Pachon8      TTTGTCAATTTTTTGTTTTTAATTTTGTCAAGAAATTTTGATCTCTTACGAAATTCGGAA   4608
Pachon15     TTTGTCAATTTTTTGTTTTTAATTTTGTCAAGAAATTTTGATCTCTTACGAAATTCGGAA   4608
             ***** **************************************** ** **********

Rascon4      GAACACTTTTAAACAAAACTGTCACTGTTAATCAGTGGGAGCAACTTGGACTTGTGTTTG   4597
Surface      GAACACTTTTAAACAAAACTGTCACTGTTAATCAGTGGGAGCAACTTGGACTTGTGTTTG   4678
Rascon8      GAACACTTTTAAACAAAACTGTCACTGTTAATCAGTGGGAGCAACTTGGACTTGTGTTTG   4660
Rascon2      GAACACTTTTAAACAAAACTGTCACTGTTAATCAGTGGGAGCAACTTGGACTTGTGTTTG   4666
Rascon15     GAACACTTTTAAACAAAACTGTCACTGTTAATCAGTGGGAGCAACTTGGACTTGTGTTTG   4654
Rascon13     GAACACTTTTAAACAAAACTGTCACTGTTAATCAGTGGGAGCAACTTGGACTTGTGTTTG   4662
Rascon6      GAACACTTTTAAACAAAACTGTCACTGTTAATCAGTGGGAGCAACTTGGACTTGTGTTTG   4661
Pachon14     GAACACTTTTAAACAAAACTGTCACTGTTAATCAGTGGGAGCAACTTGGACTTGTGTTTG   4661
Pachon9      GAACACTTTTAAACAAAACTGTCACTGTTAATCAGTGGGAGCAACTTGGACTTGTGTTTG   4669
Pachon17     GAACACTTTTAAACAAAACTGTCACTGTTAATCAGTGGGAGCAACTTGGACTTGTGTTTG   4668
Pachon12     GAACACTTTTAAACAAAACTGTCACTGTTAATCAGTGGGAGCAACTTGGACTTGTGTTTG   4668
Pachon11     GAACACTTTTAAACAAAACTGTCACTGTTAATCAGTGGGAGCAACTTGGACTTGTGTTTG   4668
Pachon7      GAACACTTTTAAACAAAACTGTCACTGTTAATCAGTGGGAGCAACTTGGACTTGTGTTTG   4668
Pachon3      GAACACTTTTAAACAAAACTGTCACTGTTAATCAGTGGGAGCAACTTGGACTTGTGTTTG   4668
Pachon8      GAACACTTTTAAACAAAACTGTCACTGTTAATCAGTGGGAGCAACTTGGACTTGTGTTTG   4668
Pachon15     GAACACTTTTAAACAAAACTGTCACTGTTAATCAGTGGGAGCAACTTGGACTTGTGTTTG   4668
             ************************************************************

Rascon4      TTATTTGTTTTTTATTATATTTCTTTATTTTTTAATTTATTTGTTGTTAAAC--TTTTTT   4655
Surface      TTATTTGTTTTTTATTATATTTCTTTATTTTTTAATTTATTTGTTGTTAAAC--TTTTTT   4736
Rascon8      TTATTTGTTTTTTATTATATTTCTTTATTTTTTAATTTATTTGTTGTTAAAC--TTTTTT   4718
Rascon2      TTATTTGTTTTTTATTATATTTCTTTATTTTTTAATTTATTTGTTGTTAAAC--TTTTTT   4724
Rascon15     TTATTTGTTTTTTATTACATTTCTTTATTTTTTAATTTATTTGTTGTTAAAC--TTTTTT   4712
Rascon13     TTATTTGTTTTTTATTATATTTCTTTATTTTTTAATTTATTTGTTGTTAAACTTTTTTTT   4722
Rascon6      TTATTTGTTTTTTATTATATTTCTTTATTTTTTAATTTATTTGTTGTTAAAC--TTTTTT   4719
Pachon14     TTATTTGTTTTTTATTATATTTCTTTATTTTTTAATTTATTTGTTGTTAAAC-TTTTTTT   4720
Pachon9      TTATTTGTTTTTTATTATATTTCTTTATTTTTTAATTTATTTGTTGTTAAAC-TTTTTTT   4728
Pachon17     TTATTTGTTTTTTATTATATTTCTTTATTTTTTAATTTATTTGTTGTTAAAC--TTTTTT   4726
Pachon12     TTATTTGTTTTTTATTATATTTCTTTATTTTTTAATTTATTTGTTGTTAAAC-TTTTTTT   4727
Pachon11     TTATTTGTTTTTTATTATATTTCTTTATTTTTTAATTTATTTGTTGTTAAAC--TTTTTT   4726
Pachon7      TTATTTGTTTTTTATTATATTTCTTTATTTTTTAATTTATTTGTTGTTAAAC-TTTTTTT   4727
Pachon3      TTATTTGTTTTTTATTATATTTCTTTATTTTTTAATTTATTTGTTGTTAAAC-TTTTTTT   4727
Pachon8      TTATTTGTTTTTTATTATATTTCTTTATTTTTTAATTTATTTGTTGTTAAAC-TTTTTTT   4727
Pachon15     TTATTTGTTTTTTATTATATTTCTTTATTTTTTAATTTATTTGTTGTTAAAC-TTTTTTT   4727
             ***************** **********************************  ******

Rascon4      TTTATTGTACTACCATCACTCTCTCAAAGTAACTTTTTTACATACAAAACTTTACATTTA   4715
Surface      TTTATTGTACTACCATCACTCTCTCAAAGTAACTTTTTTACATACAAAACTTTACATTTA   4796
Rascon8      TTTATTGTACTACCATCACTCTCTCAAAGTAACTTTTTTACATACAAAACTTTACATTTA   4778
Rascon2      TTTATTGTGCTACCATCACTCTCTCAAAGTAACTTTTTTACATACAAAACTTTACATTTA   4784
Rascon15     TTTATTGTACTACCATCACTCTCTCAAAGTAACTTTTTTACATACAAAACTTTACATTTA   4772
Rascon13     TTTATTGTACTACCATCACTCTCTCAAAGTAACTTTTTTACATACAAAACTTTACATTTA   4782
Rascon6      TTTATTGTACTACCATCACTCTCTCAAAGTAACTTTTTTACATACAAAACTTTACATTTA   4779
Pachon14     TTTATTGTACTACCATCACTCTCTCAAAGTAACTTTTTTACATACAAAACTTTACATTTA   4780
Pachon9      TTTATTGTACTACCATCACTCTCTCAAAGTAACTTTTTTACATACAAAACTTTACATTTA   4788
Pachon17     TTTATTGTACTACCATCACTCTCTCAAAGTAACTTTTTTACATACAAAACTTTACATTTA   4786
Pachon12     TTTATTGTACTACCATCACTCTCTCAAAGTAACTTTTTTACATACAAAACTTTACATTTA   4787
Pachon11     TTTATTGTACTACCATCACTCTCTCAAAGTAACTTTTTTACATACAAAACTTTACATTTA   4786
Pachon7      TTTATTGTACTACCATCACTCTCTCAAAGTAACTTTTTTACATACAAAACTTTACATTTA   4787
Pachon3      TTTATTGTACTACCATCACTCTCTCAAAGTAACTTTTTTACATACAAAACTTTACATTTA   4787
Pachon8      TTTATTGTACTACCATCACTCTCTCAAAGTAACTTTTTTACATACAAAACTTTACATTTA   4787
Pachon15     TTTATTGTACTACCATCACTCTCTCAAAGTAACTTTTTTACATACAAAACTTTACATTTA   4787
             ******** ***************************************************

Rascon4      AAACTTTTTTTCATATTATTTTTCTACCAACTGACCTTTACTTTGTAGTAAAAGTAAAGT   4775
Surface      AAACTTTTTTTCATATTATTTTTCTACCAACTGACCTTTACTTTGTAGTAAAAGTAAAGT   4856
Rascon8      AAACTTTTTTTCATATTATTTTTCTACCAACTGACCTTTACTTTGTAGTAAAAGTAAAGT   4838
Rascon2      AAACTTTTTTTCATATTATTTTTCTACCAACTGACCTTTACTTTGTAGTAAAAGTAAAGT   4844
Rascon15     AAACTTTTTTTCATATTATTTTTCTACCAACTGACCTTTACTTTGTAGTAAAAGTAAAGT   4832
Rascon13     AAACTTTTTTTCATATTATTTTTCTACCAACTGACCTTTACTTTGTAGTAAAAGTAAAGT   4842
Rascon6      AAACTTTTTTTCATATTATTTTTCTACCAACTGACCTTTACTTTGTAGTAAAAGTAAAGT   4839
Pachon14     AAACTTTTTTTCATATTATTTTTCTACCAACTGACCTTTACTTTGTAGTAAAAGTAAAGT   4840
Pachon9      AAACTTTTTTTCATATTATTTTTCTACCAACTGACCTTTACTTTGTAGTAAAAGTAAAGT   4848
Pachon17     AAACTTTTTTTCATATTATTTTTCTACCAACTGACCTTTACTTTGTAGTAAAAGTAAAGT   4846
Pachon12     AAACTTTTTTTCATATTATTTTTCTACCAACTGACCTTTACTTTGTAGTAAAAGTAAAGT   4847
Pachon11     AAACTTTTTTTCATATTATTTTTCTACCAACTGACCTTTACTTTGTAGTAAAAGTAAAGT   4846
Pachon7      AAACTTTTTTTCATATTATTTTTCTACCAACTGACCTTTACTTTGTAGTAAAAGTAAAGT   4847
Pachon3      AAACTTTTTTTCATATTATTTTTCTACCAACTGACCTTTACTTTGTAGTAAAAGTAAAGT   4847
Pachon8      AAACTTTTTTTCATATTATTTTTCTACCAACTGACCTTTACTTTGTAGTAAAAGTAAAGT   4847
Pachon15     AAACTTTTTTTCATATTATTTTTCTACCAACTGACCTTTACTTTGTAGTAAAAGTAAAGT   4847
             ************************************************************
```

```
Rascon4     CACTAACTGGAAGTAACTGGCCCAAGTTTTCTTATTTTTAAATTTTTTAAATTACGTAAT   4835
Surface     CACTAACTGGAAGTAACTGG---AAGTTTTCTTATTTTTAAATTTTTTAAATTACGTAAT   4913

Rascon8     CACTAACTGGAAGTAACTGGCCCAAGTTTTCTCATTTTTAAATTTTTTAAATTACGTAAT   4898
Rascon2     CACTAACTGGAAGTAACTGGCCCAAGTTTTCTTATTTTTAAATTTTTTAAATTACGTAAT   4904
Rascon15    CACTAACTGGAAGTAACTGGCCCAAGTTTTCTTATTTTTAAATTTTTTAAATTACGTAAT   4892
Rascon13    CACTAACTGGAAGTAACTGGCCCAAGTTTTCTTATTTTTAAATTTTTTAAATTACGTAAT   4902
Rascon6     CACTAACTGGAAGTAACTGG---AAGTTTTCTTATTTTTAAATTTTTTAAATTACGTAAT   4896
Pachon14    CACTAACTGGAAGTAACTGGCCCAAGTTTTCTTATTTTTAAATTTTTTAAATTACGTAAT   4900
Pachon9     CACTAACTGGAAGTAACTGGCCCAAGTTTTCTTATTTTTAAATTTTTTAAATTACGTAAT   4908
Pachon17    CACTAACTGGAAGTAACTGGCCCAAGTTTTCTTATTTTTAAATTTTTTAAATTACGTAAT   4906
Pachon12    CACTAACTGGAAGTAACTGGCCCAAGTTTTCTTATTTTTAAATTTTTTAAATTACGTAAT   4907
Pachon11    CACTAACTGGAAGTAACTGGCCCAAGTTTTCTTATTTTTAAATTTTTTAAATTACGTAAT   4906
Pachon7     CACTAACTGGAAGTAACTGGCCCAAGTTTTCTTATTTTTAAATTTTTTAAATTACGTAAT   4907
Pachon3     CACTAACTGGAAGTAACTGGCCCAAGTTTTCTTATTTTTAAATTTTTTAAATTACGTAAT   4907
Pachon8     CACTAACTGGAAGTAACTGGCCCAAGTTTTCTTATTTTTAAATTTTTTAAATTACGTAAT   4907
Pachon15    CACTAACTGGAAGTAACTGGCCCAAGTTTTCTTATTTTTAAATTTTTTAAATTACGTAAT   4907
            ********************   ********* ***************************

Rascon4     TTTTTGTTCTACAAATTAAATGCGTCTTGTCGGGGCTCAGTATCAGGGCATGGAGGAGCA   4895 Rx3
Surface     TTTTTGTTCTACAAATTAAATGCGTCTTGTCGGGGCTCAGTATCAGGGCATGGAGGAGCA   4973 exon 1
Rascon8     TTTTTGTTCTACAAATTAAATGCGTCTTGTCGGGGCTCAGTATCAGGGCATGGAGGAGCA   4958 start
Rascon2     TTTTTGTTCTACAAATTAAATGCGTCTTGTCGGGGCTCAGTATCAGGGCATGGAGGAGCA   4964
Rascon15    TTTTTGTTCTACAAATTAAATGCGTCTTGTCGGGGCTCAGTATCAGGGCATGGAGGAGCA   4952
Rascon13    TTTTTGTTCTACAAATTAAATGCGTCTTGTCGGGGCTCAGTATCAGGGCATGGAGGAGCA   4962
Rascon6     TTTTTGTTCTACAAATTAAATGCGTCTTGTCGGGGCTCAGTATCAGGGCATGGAGGAGCA   4956
Pachon14    TTTTTGTTCTACAAATTAAATGCGTCTTGTCGGGGCTCAGTATCAGGGCATGGAGGAGCA   4960
Pachon9     TTTTTGTTCTACAAATTAAATGCGTCTTGTCGGGGCTCAGTATCAGGGCATGGAGGAGCA   4968
Pachon17    TTTTTGTTCTACAAATTAAATGCGTCTTGTCGGGGCTCAGTATCAGGGCATGGAGGAGCA   4966
Pachon12    TTTTTGTTCTACAAATTAAATGCGTCTTGTCGGGGCTCAGTATCAGGGCATGGAGGAGCA   4967
Pachon11    TTTTTGTTCTACAAATTAAATGCGTCTTGTCGGGGCTCAGTATCAGGGCATGGAGGAGCA   4966
Pachon7     TTTTTGTTCTACAAATTAAATGCGTCTTGTCGGGGCTCAGTATCAGGGCATGGAGGAGCA   4967
Pachon3     TTTTTGTTCTACAAATTAAATGCGTCTTGTCGGGGCTCAGTATCAGGGCATGGAGGAGCA   4967
Pachon8     TTTTTGTTCTACAAATTAAATGCGTCTTGTCGGGGCTCAGTATCAGGGCATGGAGGAGCA   4967
Pachon15    TTTTTGTTCTACAAATTAAATGCGTCTTGTCGGGGCTCAGTATCAGGGCATGGAGGAGCA   4967
            ************************************************************

Rascon4     GCTGTCCCCGTCGGCTCGTTTGGCTCGTGCTGCCGGCCCCCAGTCCAGAGTCCACAGCAT   4955
Surface     GCTGTCCCCGTCGGCTCGTTTGGCTCGTGCTGCCGGCCCCCAGTCCAGAGTCCACAGCAT   5033
Rascon8     GCTGTCCCCGTCGGCTCGTTTGGCTCGTGCTGCCGGCCCCCAGTCCAGAGTCCACAGCAT   5018
Rascon2     GCTGTCCCCGTCGGCTCGTTTGGCTCGTGCTGCCGGCCCCCAGTCCAGAGTCCACAGCAT   5024
Rascon15    GCTGTCCCCGTCGGCTCGTTTGGCTCGTGCTGCCGGCCCCCAGTCCAGAGTCCACAGCAT   5012
Rascon13    GCTGTCCCCGTCGGCTCGTTTGGCTCGTGCTGCCGGCCCCCAGTCCAGAGTCCACAGCAT   5022
Rascon6     GCTGTCCCCGTCGGCTCGTTTGGCTCGTGCTGCCGGCCCCCAGTCCAGAGTCCACAGCAT   5016
Pachon14    GCTGTCCCCGTCGGCTCGTTTGGCTCGTGCTGCCGGCCCCCAGTCCAGAGTCCACAGCAT   5020
Pachon9     GCTGTCCCCGTCGGCTCGTTTGGCTCGTGCTGCCGGCCCCCAGTCCAGAGTCCACAGCAT   5028
Pachon17    GCTGTCCCCGTCGGCTCGTTTGGCTCGTGCTGCCGGCCCCCAGTCCAGAGTCCACAGCAT   5026
Pachon12    GCTGTCCCCGTCGGCTCGTTTGGCTCGTGCTGCCGGCCCCCAGTCCAGAGTCCACAGCAT   5027
Pachon11    GCTGTCCCCGTCGGCTCGTTTGGCTCGTGCTGCCGGCCCCCAGTCCAGAGTCCACAGCAT   5026
Pachon7     GCTGTCCCCGTCGGCTCGTTTGGCTCGTGCTGCCGGCCCCCAGTCCAGAGTCCACAGCAT   5027
Pachon3     GCTGTCCCCGTCGGCTCGTTTGGCTCGTGCTGCCGGCCCCCAGTCCAGAGTCCACAGCAT   5027
Pachon8     GCTGTCCCCGTCGGCTCGTTTGGCTCGTGCTGCCGGCCCCCAGTCCAGAGTCCACAGCAT   5027
Pachon15    GCTGTCCCCGTCGGCTCGTTTGGCTCGTGCTGCCGGCCCCCAGTCCAGAGTCCACAGCAT   5027
            ************************************************************

Rascon4     AGAGGCCATTCTGGGCTTCAGAGAGGAGAAGATGCTGCAGGTGGAATCATGCGGCACCAA   5015
Surface     AGAGGCCATTCTGGGCTTCAGAGAGGAGAAGATGCTGCAGGTGGAATCATGCGGCACCAA   5093
Rascon8     AGAGGCCATTCTGGGCTTCAGAGAGGAGAAGATGCTGCAGGTGGAATCATGCGGCACCAA   5078
Rascon2     AGAGGCCATTCTGGGCTTCAGAGAGGAGAAGATGCTGCAGGTGGAATCATGCGGCACCAA   5084
Rascon15    AGAGGCCATTCTGGGCTTCAGAGAGGAGAAGATGCTGCAGGTGGAATCATGCGGCACCAA   5072
Rascon13    AGAGGCCATTCTGGGCTTCAGAGAGGAGAAGATGCTGCAGGTGGAATCATGCGGCACCAA   5082
Rascon6     AGAGGCCATTCTGGGCTTCAGAGAGGAGAAGATGCTGCAGGTGGAATCATGCGGCACCAA   5076
Pachon14    AGAGGCCATTCTGGGCTTCAGAGAGGAGAAGATGCTGCAGGTGGAATCATGCGGCACCAA   5080
Pachon9     AGAGGCCATTCTGGGCTTCAGAGAGGAGAAGATGCTGCAGGTGGAATCATGCGGCACCAA   5088
Pachon17    AGAGGCCATTCTGGGCTTCAGAGAGGAGAAGATGCTGCAGGTGGAATCATGCGGCACCAA   5086
Pachon12    AGAGGCCATTCTGGGCTTCAGAGAGGAGAAGATGCTGCAGGTGGAATCATGCGGCACCAA   5087
Pachon11    AGAGGCCATTCTGGGCTTCAGAGAGGAGAAGATGCTGCAGGTGGAATCATGCGGCACCAA   5086
Pachon7     AGAGGCCATTCTGGGCTTCAGAGAGGAGAAGATGCTGCAGGTGGAATCATGCGGCACCAA   5087
Pachon3     AGAGGCCATTCTGGGCTTCAGAGAGGAGAAGATGCTGCAGGTGGAATCATGCGGCACCAA   5087
Pachon8     AGAGGCCATTCTGGGCTTCAGAGAGGAGAAGATGCTGCAGGTGGAATCATGCGGCACCAA   5087
Pachon15    AGAGGCCATTCTGGGCTTCAGAGAGGAGAAGATGCTGCAGGTGGAATCATGCGGCACCAA   5087
            ************************************************************

Rascon4     AGGGGCCACAAAAACAGACACGACGACACCCGACAACTTCCAGAGTAAGTCTAATACTAA   5075 Exon 1
Surface     AGGGGCCACAAAAACAGACACGACGACACCCGACAACTTCCAGAGTAAGTCTAATACTAA   5153 end
Rascon8     AGGGGCCACAAAAACAGACACGACGACACCCGACAACTTCCAGAGTAAGTCTAATACTAA   5138
Rascon2     AGGGGCCACAAAAACAGACACGACGACACCCGACAACTTCCAGAGTAAGTCTAATACTAA   5144
```

```
Rascon15     AGGGGCCACAAAAACAGACACGACGACACCCGACAACTTCCAGAGTAAGTCTAATACTAA     5132
Rascon13     AGGGGCCACAAAAACAGACACGACGACACCCGACAACTTCCAGAGTAAGTCTAATACTAA     5142
Rascon6      AGGGGCCACAAAAACAGACACGACGACACCCGACAACTTCCAGAGTAAGTCTAATACTAA     5136
Pachon14     AGGGGCCACAAAAACAGACACGACGACACCCGACAACTTCCAGAGTAAGTCTAATACTAA     5140
Pachon9      AGGGGCCACAAAAACAGACACGACGACACCCGACAACTTCCAGAGTAAGTCTAATACTAA     5148
Pachon17     AGGGGCCACAAAAACAGACACGACGACACCCGACAACTTCCAGAGTAAGTCTAATACTAA     5146
Pachon12     AGGGGCCACAAAAACAGACACGACGACACCCGACAACTTCCAGAGTAAGTCTAATACTAA     5147
Pachon11     AGGGGCCACAAAAACAGACACGACGACACCCGACAACTTCCAGAGTAAGTCTAATACTAA     5146
Pachon7      AGGGGCCACAAAAACAGACACGACGACACCCGACAACTTCCAGAGTAAGTCTAATACTAA     5147
Pachon3      AGGGGCCACAAAAACAGACACGACGACACCCGACAACTTCCAGAGTAAGTCTAATACTAA     5147
Pachon8      AGGGGCCACAAAAACAGACACGACGACACCCGACAACTTCCAGAGTAAGTCTAATACTAA     5147
Pachon15     AGGGGCCACAAAAACAGACACGACGACACCCGACAACTTCCAGAGTAAGTCTAATACTAA     5147
             ************************************************************

Rascon4      TACTCCACACTTTTTTTTTTTACTGTAAAATAATCTCCTGTACTATACTATACTGGCATA     5135 Intron 1
Surface      TACTCCACAC--TTTTTTTTTACTGTAAAATAATCTCCTGTACTATACTATACTGGCATA     5211
Rascon8      TACTCCACAC--TTTTTTTTTACTGTAAAATAATCTCCTGTACTATACTATACTGGCATA     5196
Rascon2      TACTCCACAC--TTTTTTTTTACTGTAAAATAATCTCCTGTACTATACTATACTGGCATA     5202
Rascon15     TACTCCACAC--TTTTTTTTAATTGTAAAATAATCTCCTGTACTATACTATACTGGCATA     5190
Rascon13     TACTCCACAC--TTTTTTTTAACTGTAAAATAATCTCCTGTACTATACTATACTGGCATA     5200
Rascon6      TACTCCACACTTTTTTTTTTTACTGTAAAATAATCTCCTGTACTATACTATACTGGCATA     5196
Pachon14     TACTCCACAC--TTTTTTTTAACTGTAAAATAATCTCCTGTACTATACTATACTGGCATA     5198
Pachon9      TACTCCACAC--TTTTTTTTAACTGTAAAATAATCTCCTGTACTATACTATACTGGCATA     5206
Pachon17     TACTCCACAC--TTTTTTTTAACTGTAAAATAATCTCCTGTACTATACTATACTGGCATA     5204
Pachon12     TACTCCACAC--TTTTTTTTAACTGTAAAATAATCTCCTGTACTATACTATACTGGCATA     5205
Pachon11     TACTCCACAC--TTTTTTTTAACTGTAAAATAATCTCCTGTACTATACTATACTGGCATA     5204
Pachon7      TACTCCACAC--TTTTTTTTAACTGTAAAATAATCTCCTGTACTATACTATACTGGCATA     5205
Pachon3      TACTCCACAC--TTTTTTTTAACTGTAAAATAATCTCCTGTACTATACTATACTGGCATA     5205
Pachon8      TACTCCACAC--TTTTTTTTAACTGTAAAATAATCTCCTGTACTATACTATACTGGCATA     5205
Pachon15     TACTCCACAC--TTTTTTTTAACTGTAAAATAATCTCCTGTACTATACTATACTGGCATA     5205
             **********  ******** * *************************************

Rascon4      GTATATATTGGAGGGATATACTAAAATGCAAATTTAGGTGAAAACTTAAA--ATATATAT     5193
Surface      GTATATATTGGAGGGATATACTAAAATGCAAATTTTGGTGAAAACTTAAAATATATATAT     5271
Rascon8      GTATATATTGGAGGGATATACTAAAATGCAAATTTTGGTGAAAACTTAAAATATATATAT     5256
Rascon2      GTATATATTGGAGGGATATACTAAAATGCAAATTTTGGTGAAAACTAAAA--ATATATAT     5260
Rascon15     GTATATATTGGAGGGATATACTAAAATGCAAATTTTGGTGAAAACTTAAAATATATATAT     5250
Rascon13     GTATATATTGGAGGGATATACTAAAATGCAAATTTTGGTGAAAACTTAAAATATATATAT     5260
Rascon6      GTATATATTGGAGGGATATACTAAAATGCAAATTTAGGTGAAAACTTAAA--ATATATAT     5254
Pachon14     GTATATATTGGAGGGATATACTAAAATGCAAATTTTGGTGAAAACTTAAAATATATATAT     5258
Pachon9      GTATATATTGGAGGGATATACTAAAATGCAAATTTTGGTGAAAACTTAAAATATATATAT     5266
Pachon17     GTATATATTGGAGGGATATACTAAAATGCAAATTTTGGTGAAAACTTAAAATATATATAT     5264
Pachon12     GTATATATTGGAGGGATATACTAAAATGCAAATTTTGGTGAAAACTTAAAATATATATAT     5265
Pachon11     GTATATATTGGAGGGATATACTAAAATGCAAATTTTGGTGAAAACTTAAAATATATATAT     5264
Pachon7      GTATATATTGGAGGGATATACTAAAATGCAAATTTTGGTGAAAACTTAAAATATATATAT     5265
Pachon3      GTATATATTGGAGGGATATACTAAAATGCAAATTTTGGTGAAAACTTAAAATATATATAT     5265
Pachon8      GTATATATTGGAGGGATATACTAAAATGCAAATTTTGGTGAAAACTTAAAATATATATAT     5265
Pachon15     GTATATATTGGAGGGATATACTAAAATGCAAATTTTGGTGAAAACTTAAAATATATATAT     5265
             *********************************** ********** ***  ********

Rascon4      ATATTTTTTATTCTGCCACTTTTGCCACAATTGCATAAACAAAATACAATTAAATGTATT     5253
Surface      ATATTTTTTATTCTGCAACTTTTGCCACAATTGCATAAACAAAATACAATTAAATGTATT     5331
Rascon8      ATATTTTTTATTCTGCCACTTTTGCCACAATTGCATAAACAAAATACAATTAAATGTATT     5316
Rascon2      ATATTTTTTATTCTGCCACTTTTGCCACAATTGCATAAACAAAATACAATTAAATGTATT     5320
Rascon15     ATATTTTTTGTTCTGCCACTTTTGCCACAATTGCATAAACAAAATACAATTAAATGTATT     5310
Rascon13     ATATTTTTTGTTCTGCCACTTTTGCCACAATTGCATAAACAAAATACAATTAAATGTATT     5320
Rascon6      ATATTTTTTATTCTGCCACTTTTGCCACAATTGCATAAACAAAATACAATTAAATGTATT     5314
Pachon14     ATATTTTTTGTTCTGCCACTTTTGCCACAATTGCATAAACAAAATACAATTAAATGTATT     5318
Pachon9      ATATTTTTTGTTCTGCCACTTTTGCCACAATTGCATAAACAAAATACAATTAAATGTATT     5326
Pachon17     ATATTTTTTGTTCTGCCACTTTTGCCACAATTGCATAAACAAAATACAATTAAATGTATT     5324
Pachon12     ATATTTTTTGTTCTGCCACTTTTGCCACAATTGCATAAACAAAATACAATTAAATGTATT     5325
Pachon11     ATATTTTTTGTTCTGCCACTTTTGCCACAATTGCATAAACAAAATACAATTAAATGTATT     5324
Pachon7      ATATTTTTTGTTCTGCCACTTTTGCCACAATTGCATAAACAAAATACAATTAAATGTATT     5325
Pachon3      ATATTTTTTGTTCTGCCACTTTTGCCACAATTGCATAAACAAAATACAATTAAATGTATT     5325
Pachon8      ATATTTTTTGTTCTGCCACTTTTGCCACAATTGCATAAACAAAATACAATTAAATGTATT     5325
Pachon15     ATATTTTTTGTTCTGCCACTTTTGCCACAATTGCATAAACAAAATACAATTAAATGTATT     5325
             ********* ****** *******************************************

Rascon4      TTTGTCTTGTTTAAAAAAAA--ATAATAATAATAATAATTTGGTATAAATTATTCTGTAT     5311
Surface      TTTGTCTTGTTTAAAAAAAATAATAATAATAATAATAATTTGGTATAAATTATTCTGTAT     5391
Rascon8      TTTGTCTTGTTTAAAAAAAAA--TAATAATAATAATAATTTGGTATAAATTATTCTGTAT     5374
Rascon2      TTTGTCTTGTTTAAAAAAAAA--TAATAATAATAATAATTTGGTATAAATTATTCTGTAT     5378
Rascon15     TTTGTCTTGTTTAAAAAAA-----AATAATAATAATAATTTGGTATAAATTATTCTGTAT     5365
Rascon13     TTTGTCTTGTTTAAAAAAAAT--AATAAATAATAATAATTTGGTATAAATTATTCTGTAT     5378
Rascon6      TTTGTCTTGTTTAAAAAAAAT--AAATAATAATAATAATTTGGTATAAATTATTCTGTAT     5372
Pachon14     TTTGTCTTGTTTAAAAAAAAA--TAATAATAATAATAATTTGGTATAAATTATTCTGTAT     5376
Pachon9      TTTGTCTTGTTTAAAAAAAAA--TAATAATAATAATAATTTGGTATAAATTATTCTGTAT     5384
```

```
Pachon17      TTTGTCTTGTTTAAAAAAAAA--TAATAATAATAATAATTTGGTATAAATTATTCTGTAT      5382
Pachon12      TTTGTCTTGTTTAAAAAAAAA--TAATAATAATAATAATTTGGTATAAATTATTCTGTAT      5383
Pachon11      TTTGTCTTGTTTAAAAAAAAA--TAATAATAATAATAATTTGGTATAAATTATTCTGTAT      5382
Pachon7       TTTGTCTTGTTTAAAAAAAAA--TAATAATAATAATAATTTGGTATAAATTATTCTGTAT      5383
Pachon3       TTTGTCTTGTTTAAAAAAAAA--TAATAATAATAATAATTTGGTATAAATTATTCTGTAT      5383
Pachon8       TTTGTCTTGTTTAAAAAAAAA--TAATAATAATAATAATTTGGTATAAATTATTCTGTAT      5383
Pachon15      TTTGTCTTGTTTAAAAAAAAA--TAATAATAATAATAATTTGGTATAAATTATTCTGTAT      5383
              ******************      *  *********************************

Rascon4       TGTCAATAATAATTTAGGTTGCCAAATAAGTATTTCAGTTGCAAAATATTTAATATTCAA      5371 Rx3
Surface       TTTCAATAATAATTTAGGTTGCCAAATAAGTATTTCAGTTGCAAAATATTTAATATTCAA      5451 Intron 1
Rascon8       TTTCAATAATAATTTAGGTTGCCAAATAAGTATTTCAGTTGCAAAATATTTAATATTCAA      5434
Rascon2       TTTCAATAATAATTTAGGTTGCCAAATAAGTATTTCAGTTGCAAAATATTTAATATTCAA      5438
Rascon15      TTTCAATAATAATTTAGGTTGCCAAATAAGTATTTCAGTTGCAAAATATTTAATATTCAA      5425
Rascon13      TTTCAATAATAATTTAGGTTGCCAAATAAGTATTTCAGTTGCAAAATATTTAATATTCAA      5438
Rascon6       TTTCAATAATAATTTGGGTTGCCAAATAAGTATTTCAGTTGCAAAATATTTAATATTCAA      5432
Pachon14      TTTCAATAATAATTTAGGTTGCCAAATAAGTATTTCAGTTGCAAAATATTTAATATTCAA      5436
Pachon9       TTTCAATAATAATTTAGGTTGCCAAATAAGTATTTCAGTTGCAAAATATTTAATATTCAA      5444
Pachon17      TTTCAATAATAATTTAGGTTGCCAAATAAGTATTTCAGTTGCAAAATATTTAATATTCAA      5442
Pachon12      TTTCAATAATAATTTAGGTTGCCAAATAAGTATTTCAGTTGCAAAATATTTAATATTCAA      5443
Pachon11      TTTCAATAATAATTTAGGTTGCCAAATAAGTATTTCAGTTGCAAAATATTTAATATTCAA      5442
Pachon7       TTTCAATAATAATTTAGGTTGCCAAATAAGTATTTCAGTTGCAAAATATTTAATATTCAA      5443
Pachon3       TTTCAATAATAATTTAGGTTGCCAAATAAGTATTTCAGTTGCAAAATATTTAATATTCAA      5443
Pachon8       TTTCAATAATAATTTAGGTTGCCAAATAAGTATTTCAGTTGCAAAATATTTAATATTCAA      5443
Pachon15      TTTCAATAATAATTTAGGTTGCCAAATAAGTATTTCAGTTGCAAAATATTTAATATTCAA      5443
              * ************* ********************************************

Rascon4       GCATGCTTAATGCCAGTATACTGTG--TTTTTTTTTACATTTTTGGTCAAATTTTGGTTG      5429
Surface       GCATGCTTAATGCCAGTATACTGTGTTTTTTTTTTTACATTTTTGGTCAAATTTTGGTTG      5511
Rascon8       GCATGCTTAATGCCAGTATACTGTG--TTTTTTTTTACATTTTTGGTCAAATTTTGGTTG      5492
Rascon2       GCATGCTTAATGCCAGTATACTGTG--TTTTTTTTTACATTTTTGGTCAAATTTTGGTTG      5496
Rascon15      GCATGCTTAATGCCAGTATACTGTG--TTTTTTTTTACATTTTTGGTCAAATTTTGGTTG      5483
Rascon13      GCATGCTTAATGCCAGTATACTGTGTTTTTTTTTTTACATTTTTGGTCAAATTTTGGTTG      5498
Rascon6       GCATGCTTAATGCCAGTATACTGTG--TTTTTTTTTACATTTTTGGTCAAATTTTGGTTG      5490
Pachon14      GCATGCTTAATGCCAGTATACTGTG--TTTTTTTTTACATTTTTGGTCAAATTTTGGTTG      5494
Pachon9       GCATGCTTAATGCCAGTATACTGTG--TTTTTTTTTACATTTTTGGTCAAATTTTGGTTG      5502
Pachon17      GCATGCTTAATGCCAGTATACTGTG--TTTTTTTTTACATTTTTGGTCAAATTTTGGTTG      5500
Pachon12      GCATGCTTAATGCCAGTATACTGTG--TTTTTTTTTACATTTTTGGTCAAATTTTGGTTG      5501
Pachon11      GCATGCTTAATGCCAGTATACTGTG--TTTTTTTTTACATTTTTGGTCAAATTTTGGTTG      5500
Pachon7       GCATGCTTAATGCCAGTATACTGTG--TTTTTTTTTACATTTTTGGTCAAATTTTGGTTG      5501
Pachon3       GCATGCTTAATGCCAGTATACTGTG--TTTTTTTTTACATTTTTGGTCAAATTTTGGTTG      5501
Pachon8       GCATGCTTAATGCCAGTATACTGTG--TTTTTTTTTACATTTTTGGTCAAATTTTGGTTG      5501
Pachon15      GCATGCTTAATGCCAGTATACTGTG--TTTTTTTTTACATTTTTGGTCAAATTTTGGTTG      5501
              *************************  *********************************

Rascon4       CCAAATAATTTAATTTGCTGAATGTTTGATGTTTTTTCAAATAATTTGTAAATGTACTGT      5489
Surface       CCAAATAATTTAATTTGCTGAATGTTTGATGTTTTTTCAAATAATTTGTAAATGTACTGT      5571
Rascon8       CCAAATAATTTAATTTGCTGAATGTTTGATGTTTTTTCAAATAATTTGTAAATGTACTGT      5552
Rascon2       CCAAATAATTTAATTTGCTGAATGTTTGATGTTTTTTCAAATAATTTGTAAATGTACTGT      5556
Rascon15      CCAAATAATTTAATTTGCTGAATGTTTGATGTTTTTTCAAATAATTTGTAAATGTACTGT      5543
Rascon13      CCAAATAATTTAATTTGCTGAATGTTTGATGTTTTTTCAAATAATTTGTAAATGTACTGT      5558
Rascon6       CCAAATAATTTAATTTGCTGAATGTTTGATGTTTTTTCAAATAATTTGTAAATGTACTGT      5550
Pachon14      CCAAATAATTTAATTTGCTGAATGTTTGATGTTTTTTCAAATAATTTGTAAATGTACTGT      5554
Pachon9       CCAAATAATTTAATTTGCTGAATGTTTGATGTTTTTTCAAATAATTTGTAAATGTACTGT      5562
Pachon17      CCAAATAATTTAATTTGCTGAATGTTTGATGTTTTTTCAAATAATTTGTAAATGTACTGT      5560
Pachon12      CCAAATAATTTAATTTGCTGAATGTTTGATGTTTTTTCAAATAATTTGTAAATGTACTGT      5561
Pachon11      CCAAATAATTTAATTTGCTGAATGTTTGATGTTTTTTCAAATAATTTGTAAATGTACTGT      5560
Pachon7       CCAAATAATTTAATTTGCTGAATGTTTGATGTTTTTTCAAATAATTTGTAAATGTACTGT      5561
Pachon3       CCAAATAATTTAATTTGCTGAATGTTTGATGTTTTTTCAAATAATTTGTAAATGTACTGT      5561
Pachon8       CCAAATAATTTAATTTGCTGAATGTTTGATGTTTTTTCAAATAATTTGTAAATGTACTGT      5561
Pachon15      CCAAATAATTTAATTTGCTGAATGTTTGATGTTTTTTCAAATAATTTGTAAATGTACTGT      5561
              ************************************************************

Rascon4       ACTTTTTTTTGCCATTTCCTGACGAATAATTTTAGTTGAAAAGCATTCAGTGCATTCCTA      5549
Surface       ACTTTTTTTTGCCATTTCCTGACGAATAATTTTAGTTGAAAAGCATTCAGTGCATTCCTA      5631
Rascon8       ACTTTTTTTTGCCATTTCCTGATGAATAATTTTAGTTGAAAAGCATTCAGTGCATTCCTA      5612
Rascon2       ACTTTTTTTTGCCATTTCCTGACGAATAATTTTAGTTGAAAAGCATTCAGTGCATTCCTA      5616
Rascon15      ACTTTTTTTTGCCATTTCCTGACGAATAATTTTAGTTGAAAAGCATTCAGTGCATTCCTA      5603
Rascon13      ACTTTTGTTTGCCATTTCCTGACGAATAATTTTAGTTGAAAAGCATTCAGTGCATTCCTA      5618
Rascon6       ACTTTTTTTTGCCATTTCCTGACGAATAATTTTAGTTGAAAAGCATTCAGTGCATTCCTA      5610
Pachon14      ACTTTTGTTTGCCATTTCCTGACGAATAATTTTAGTTGAAAAGCATTCAGTGCATTCCTA      5614
Pachon9       ACTTTTGTTTGCCATTTCCTGACGAATAATTTTAGTTGAAAAGCATTCAGTGCATTCCTA      5622
Pachon17      ACTTTTGTTTGCCATTTCCTGACGAATAATTTTAGTTGAAAAGCATTCAGTGCATTCCTA      5620
Pachon12      ACTTTTGTTTGCCATTTCCTGACGAATAATTTTAGTTGAAAAGCATTCAGTGCATTCCTA      5621
Pachon11      ACTTTTGTTTGCCATTTCCTGACGAATAATTTTAGTTGAAAAGCATTCAGTGCATTCCTA      5620
Pachon7       ACTTTTGTTTGCCATTTCCTGACGAATAATTTTAGTTGAAAAGCATTCAGTGCATTCCTA      5621
Pachon3       ACTTTTGTTTGCCATTTCCTGACGAATAATTTTAGTTGAAAAGCATTCAGTGCATTCCTA      5621
```

```
Pachon8      ACTTTTGTTTGCCATTTCCTGACGAATAATTTTAGTTGAAAAGCATTCAGTGCATTCCTA    5621
Pachon15     ACTTTTGTTTGCCATTTCCTGACGAATAATTTTAGTTGAAAAGCATTCAGTGCATTCCTA    5621
             ****** *************** *************************************

Rascon4      ATAACTTAAGTTTTAGTTTTAATTAAATTAAAAAAAAATTATATCTTTATTGCAATTAAA    5609 Rx3
Surface      ATAACTTAAGTTTTAGTTTTAATTAAATTAAACATTTTTTATATCTTTATTGCAATTAAA    5691 Intron 1
Rascon8      ATAACTTAAGTTTTAGTTTTAATTAAATTAAACATTTTTTATATCTTTATTGCAATTAAA    5672
Rascon2      ATAACTTAAGTTTTAGTTTTAATTAAATTAAACATTTTTTATATCTTTATTGCAATTAAA    5676
Rascon15     ATAACTTAAGTTTTAGTTTTAATTAAATTAAACATTTTTTATATCTTTATTGCAATTAAA    5663
Rascon13     ATAACTTAAGTTTTAGTTTTAATTAAATTAAACATTTTTTATATCTTTATTGCAATTAAA    5678
Rascon6      ATAACTTAAGTTTTAGTTTTAATTAAATTAAAAAAATTTTATATCTTTATTGCAATTAAA    5670
Pachon14     ATAACTTAAGTTTTAGTTTTAATTAAATTAAACATTTTTTATATCTTTATTGCAATTAAA    5674
Pachon9      ATAACTTAAGTTTTAGTTTTAATTAAATTAAACATTTTTTATATCTTTATTGCAATTAAA    5682
Pachon17     ATAACTTAAGTTTTAGTTTTAATTAAATTAAACATTTTTTATATCTTTATTGCAATTAAA    5680
Pachon12     ATAACTTAAGTTTTAGTTTTAATTAAATTAAACATTTTTTATATCTTTATTGCAATTAAA    5681
Pachon11     ATAACTTAAGTTTTAGTTTTAATTAAATTAAACATTTTTTATATCTTTATTGCAATTAAA    5680
Pachon7      ATAACTTAAGTTTTAGTTTTAATTAAATTAAACATTTTTTATATCTTTATTGCAATTAAA    5681
Pachon3      ATAACTTAAGTTTTAGTTTTAATTAAATTAAACATTTTTTATATCTTTATTGCAATTAAA    5681
Pachon8      ATAACTTAAGTTTTAGTTTTAATTAAATTAAACATTTTTTATATCTTTATTGCAATTAAA    5681
Pachon15     ATAACTTAAGTTTTAGTTTTAATTAAATTAAACATTTTTTATATCTTTATTGCAATTAAA    5681
             ******************************** *    **********************

Rascon4      CTGAATCAACAGAACTGTATCTG--AAAAAAAAAAAATAGCACAAACAAATTGATATCTG    5667
Surface      CTGAATCAACAGAACTGTATCTG----AAAAAAAAAATAGCACAAACAAATTGATATCTG    5747
Rascon8      CTGAATCAACAGAACTGTATCTG--AAAAAAAAAAAATAGCACAAACAAATTGATATCTG    5730
Rascon2      CTGAATCAACAGAACTGTATCTG--AAAAAAAAAAAATAGCACAAACAAATTGATATCTG    5734
Rascon15     CTGAATCAACAGAACTGTATCTGAAAAAAAAAAAAAATAGCACAAACAAATTGATATCTG    5723
Rascon13     CTGAATCAACAGAACTGTATCTG----AAAAAAAAAATAGCACAAACAAATTGATATCTG    5734
Rascon6      CTGAATCAACAGAACTGTATCTG-AAAAAAAAAAAAATAGCACAAACAAATTGATATCTG    5729
Pachon14     CTGAATCAACAGAACTGTATCTG-----AAAAAAAAATAGCACAAACAAATTGATATCTG    5729
Pachon9      CTGAATCAACAGAACTGTATCTG----AAAAAAAAAATAGCACAAACAAATTGATATCTG    5738
Pachon17     CTGAATCAACAGAACTGTATCTG----AAAAAAAAAATAGCACAAACAAATTGATATCTG    5736
Pachon12     CTGAATCAACAGAACTGTATCTG-----AAAAAAAAATAGCACAAACAAATTGATATCTG    5736
Pachon11     CTGAATCAACAGAACTGTATCTG-----AAAAAAAAATAGCACAAACAAATTGATATCTG    5735
Pachon7      CTGAATCAACAGAACTGTATCTG-----AAAAAAAAATAGCACAAACAAATTGATATCTG    5736
Pachon3      CTGAATCAACAGAACTGTATCTG-----AAAAAAAAATAGCACAAACAAATTGATATCTG    5736
Pachon8      CTGAATCAACAGAACTGTATCTG-----AAAAAAAAATAGCACAAACAAATTGATATCTG    5736
Pachon15     CTGAATCAACAGAACTGTATCTG-----AAAAAAAAATAGCACAAACAAATTGATATCTG    5736
             ***********************     ********************************

Rascon4      CAAATTTAGAGAGCTAAGGGTGATCCTGTAAGATTTGTAGCAGTCACAGTCCATCAGTTG    5727
Surface      CAAATTTAGAGAGCTAAGGGTGATCCTGTAAGATTTGTAGCAGTCACAGTCCATCAGTTG    5807
Rascon8      CAAATTTAGAGAGCTAAGGGTGATCCTGTAAGATTTGTAGCAGTCACAGTCCATCAGTTG    5790
Rascon2      CAAATTTAGAGAGCTAAGGGTGATCCTGTAAGATTTGTAGCAGTCACAGTCCATCAGTTG    5794
Rascon15     CAAATTTAGAGAGCTAAGGGTGATCCTGTAAGATTTGTAGCAGTCACAGTCCATCAGTTG    5783
Rascon13     CAAATTTAGAGAGCTAAGGGTGATCCTGTAAGATTTGTAGCAGTCACAGTCCATCAGTTG    5794
Rascon6      CAAATTTAGAGAGCTAAGGGTGATCCTGTAAGATTTGTAGCAGTCACAGTCCATCAGTTG    5789
Pachon14     CAAATTTAGAGAGCTAAGGGTGATCCTGTAAGATTTGTAGCAGTCACAGTCCATCAGTTG    5789
Pachon9      CAAATTTAGAGAGCTAAGGGTGATCCTGTAAGATTTGTAGCAGTCACAGTCCATCAGTTG    5798
Pachon17     CAAATTTAGAGAGCTAAGGGTGATCCTGTAAGATTTGTAGCAGTCACAGTCCATCAGTTG    5796
Pachon12     CAAATTTAGAGAGCTAAGGGTGATCCTGTAAGATTTGTAGCAGTCACAGTCCATCAGTTG    5796
Pachon11     CAAATTTAGAGAGCTAAGGGTGATCCTGTAAGATTTGTAGCAGTCACAGTCCATCAGTTG    5795
Pachon7      CAAATTTAGAGAGCTAAGGGTGATCCTGTAAGATTTGTAGCAGTCACAGTCCATCAGTTG    5796
Pachon3      CAAATTTAGAGAGCTAAGGGTGATCCTGTAAGATTTGTAGCAGTCACAGTCCATCAGTTG    5796
Pachon8      CAAATTTAGAGAGCTAAGGGTGATCCTGTAAGATTTGTAGCAGTCACAGTCCATCAGTTG    5796
Pachon15     CAAATTTAGAGAGCTAAGGGTGATCCTGTAAGATTTGTAGCAGTCACAGTCCATCAGTTG    5796
             ************************************************************

Rascon4      ACCCAATCAAGCGTTTATGCTGGTGGTGCTGAATTTATACATTTTGAGCATATTGTTGAG    5787
Surface      ACCCAATCAAGCGTTTATGCTGGTGGTGCTGAATTTATACATTTTGAGCATATTGTTGAG    5867
Rascon8      ACCCAATCAAGCGTTTATGCTGGTGGTGCTGAATTTATACATTTTGAGCATATTGTTGAG    5850
Rascon2      ACCCAATCAAGCGTTTATGCTGGTGGTGCTGAATTTATACATTTTGAGCATATTGTTGAG    5854
Rascon15     ACCCAATCAAGCGTTTATGCTGGTGGTGCTGAAT-------TTTTGAGCATATTGTTGAG    5836
Rascon13     ACCCAATCAAGCGTTTATGCTGGTGGTGCTGAAT-------TTTTGAGCATATTGTTGAG    5847
Rascon6      ACCCAATCAAGCGTTTATGCTGGTGGTGCTGAATTTATACATTTTGAGCATATTGTTGAG    5849
Pachon14     ACCCAATCAAGCGTTTATGCTGGTGGTGCTGAAT-------TTTTGAGCATATTGTTGAG    5842
Pachon9      ACCCAATCAAGCGTTTATGCTGGTGGTGCTGAAT-------TTTTGAGCATATTGTTGAG    5851
Pachon17     ACCCAATCAAGCGTTTATGCTGGTGGTGCTGAAT-------TTTTGAGCATATTGTTGAG    5849
Pachon12     ACCCAATCAAGCGTTTATGCTGGTGGTGCTGAAT-------TTTTGAGCATATTGTTGAG    5849
Pachon11     ACCCAATCAAGCGTTTATGCTGGTGGTGCTGAAT-------TTTTGAGCATATTGTTGAG    5848
Pachon7      ACCCAATCAAGCGTTTATGCTGGTGGTGCTGAAT-------TTTTGAGCATATTGTTGAG    5849
Pachon3      ACCCAATCAAGCGTTTATGCTGGTGGTGCTGAAT-------TTTTGAGCATATTGTTGAG    5849
Pachon8      ACCCAATCAAGCGTTTATGCTGGTGGTGCTGAAT-------TTTTGAGCATATTGTTGAG    5849
Pachon15     ACCCAATCAAGCGTTTATGCTGGTGGTGCTGAAT-------TTTTGAGCATATTGTTGAG    5849
             **********************************       *******************

Rascon4      ACAAAAACTAAAAAAAAAATGTATAGAAGCAGCAAAATAGTCAGCATAACTTTTGTGTTG    5847
```

```
Surface        ACAAAAACTAAAAAATAAAATGTATAGAAGCAGCAAAATAGTCAGCATAACTTTTGTGTTG    5927
Rascon8        ACAAAAACTAAAAAAAAAATGTATAGAAGCAGCAAAATAGTCAGCATAACTTTTGTGTTG    5910
Rascon2        ACAAAAACT-AAAAAAAAATGTATAGAAGCAGCAAAATAGTCAGCATAACTTTTGTGTTG    5913
Rascon15       ACAAAAACTAAAAAAAAAATGTATAGAAGCAGCAAAATAGTCAGCATAACTTTTGTGTTG    5896
Rascon13       ACAAAAACTAAAAAAAAAATGTATAGAAGCAGCAAAATAGTCAGCATAACTTTTGTGTTG    5907
Rascon6        ACAAAAACTAAAAAAAAAATGTATAGAAGCAGCAAAATAGTCAGCATAACTTTTGTGTTG    5909
Pachon14       ACAAAAACTAAAAAAAAAATGTATAGAAGCAGCAAAATAGTCAGCATAACTTTTGTGTTG    5902
Pachon9        ACAAAAACTAAAAAAAAAATGTATAGAAGCAGCAAAATAGTCAGCATAACTTTTGTGTTG    5911
Pachon17       ACAAAAACTAAAAAAAAAATGTATAGAAGCAGCAAAATAGTCAGCATAACTTTTGTGTTG    5909
Pachon12       ACAAAAACTAAAAAAAAAATGTATAGAAGCAGCAAAATAGTCAGCATAACTTTTGTGTTG    5909
Pachon11       ACAAAAACTAAAAAAAAAATGTATAGAAGCAGCAAAATAGTCAGCATAACTTTTGTGTTG    5908
Pachon7        ACAAAAACTAAAAAAAAAATGTATAGAAGCAGCAAAATAGTCAGCATAACTTTTGTGTTG    5909
Pachon3        ACAAAAACTAAAAAAAAAATGTATAGAAGCAGCAAAATAGTCAGCATAACTTTTGTGTTG    5909
Pachon8        ACAAAAACTAAAAAAAAAATGTATAGAAGCAGCAAAATAGTCAGCATAACTTTTGTGTTG    5909
Pachon15       ACAAAAACTAAAAAAAAAATGTATAGAAGCAGCAAAATAGTCAGCATAACTTTTGTGTTG    5909
               ********* ***** ********************************************

Rascon4        TTTTAAGATGACCCACCACCATGTTTAACAGCTAATAGAAAAACCTACATTACCTTTATA    5907 Rx3
Surface        TTTTAAGATGACCCACCACCATGTTTAACAGCTAATAGAAAAACCTACATTACCTTTATA    5987 Intron 1
Rascon8        TTTTAAGATGACCCACCACCATGTTTAACAGCTAATAGAAAAACCTACATTACCTTTATA    5970
Rascon2        TTTTAAGATGACCCACCACCATGTTTAACAGCTAATAGAAAAACCTACATTACCTTTATA    5973
Rascon15       TTTTAAGATGACCCACCACCATGTTTAACAGCTAATAGAAAAACCTACATTACCTTTATA    5956
Rascon13       TTTTAAGATGACCCCCCACCATGTTTAACAGCTAATAGAAAAACCTACATTACCTTTATA    5967
Rascon6        TTTTAAGATGACCCACCACCATGTTTAACAGCTAATAGAAAAACCTACATTACCTTTATA    5969
Pachon14       TTTTAAGATGACCCACCACCATGTTTAACAGCTAATAGAAAAACCTACATTACCTTTATA    5962
Pachon9        TTTTAAGATGACCCACCACCATGTTTAACAGCTAATAGAAAAACCTACATTACCTTTATA    5971
Pachon17       TTTTAAGATGACCCACCACCATGTTTAACAGCTAATAGAAAAACCTACATTACCTTTATA    5969
Pachon12       TTTTAAGATGACCCACCACCATGTTTAACAGCTAATAGAAAAACCTACATTACCTTTATA    5969
Pachon11       TTTTAAGATGACCCACCACCATGTTTAACAGCTAATAGAAAAACCTACATTACCTTTATA    5968
Pachon7        TTTTAAGATGACCCACCACCATGTTTAACAGCTAATAGAAAAACCTACATTACCTTTATA    5969
Pachon3        TTTTAAGATGACCCACCACCATGTTTAACAGCTAATAGAAAAACCTACATTACCTTTATA    5969
Pachon8        TTTTAAGATGACCCACCACCATGTTTAACAGCTAATAGAAAAACCTACATTACCTTTATA    5969
Pachon15       TTTTAAGATGACCCACCACCATGTTTAACAGCTAATAGAAAAACCTACATTACCTTTATA    5969
               ************** *********************************************

Rascon4        AGCTTGAAATGTAATACATTTTCCAAGACAAATCCATTTTCTTTATAGCTGCTTTGATCA    5967
Surface        AGCTTGAAATGTAATACATTTTCCAAGACAAATCCATTTTCTTTATAGCTGCTTTGATCA    6047
Rascon8        AGCTTGAAATGTAATACATTTTCCAAGACAAATCCATTTTCTTTATAGCTGCTTTGATCA    6030
Rascon2        AGCTTGAAATGTAATACATTTTCCAAGACAAATCCATTTTCTTTATAGCTGCTTTGATCA    6033
Rascon15       AGCTTGAAATGTAATACATTTTCCAAGACAAATCCATTTTCTTTATAGCTGCTTTGATCA    6016
Rascon13       AGCTTGAAATGTAATACATTTTCCAAGACAAATCCATTTTCTTTATAGCTGCTTTGATCA    6027
Rascon6        AGCTTGAAATGTAATACATTTTCCAAGACAAATCCATTTTCTTTATAGCTGCTTTGATCA    6029
Pachon14       AGCTTGAAATGTAATACATTTTCCAAGACAAATCCATTTTCTTTATAGCTGCTTTGATCA    6022
Pachon9        AGCTTGAAATGTAATACATTTTCCAAGACAAATCCATTTTCTTTATAGCTGCTTTGATCA    6031
Pachon17       AGCTTGAAATGTAATACATTTTCCAAGACAAATCCATTTTCTTTATAGCTGCTTTGATCA    6029
Pachon12       AGCTTGAAATGTAATACATTTTCCAAGACAAATCCATTTTCTTTATAGCTGCTTTGATCA    6029
Pachon11       AGCTTGAAATGTAATACATTTTCCAAGACAAATCCATTTTCTTTATAGCTGCTTTGATCA    6028
Pachon7        AGCTTGAAATGTAATACATTTTCCAAGACAAATCCATTTTCTTTATAGCTGCTTTGATCA    6029
Pachon3        AGCTTGAAATGTAATACATTTTCCAAGACAAATCCATTTTCTTTATAGCTGCTTTGATCA    6029
Pachon8        AGCTTGAAATGTAATACATTTTCCAAGACAAATCCATTTTCTTTATAGCTGCTTTGATCA    6029
Pachon15       AGCTTGAAATGTAATACATTTTCCAAGACAAATCCATTTTCTTTATAGCTGCTTTGATCA    6029
               ************************************************************

Rascon4        CGTTTGTAAAAGCTCTAAAAAAACAGGGTCCAGAGGCCCTGATCTTTAGAACAACAATGA    6027
Surface        CTTTTGTAAAAGCTCTAAAAAAACAGGGTCCAGAGGCCCTGATCTTTAGAACAACAATGA    6107
Rascon8        CTTTTGTAAAAGCTCTAAAAAAACAGGGTCCAGAGGCCCTGATCTTTAGAACAACAATGA    6090
Rascon2        CTTTTGTAAAAGCTCTAAAAAAACAGGGTCCAGAGGCCCTGATCTTTAGAACAACAATGA    6093
Rascon15       CTTTTGTAAAAGCTCTAAAAAAACAGGGTCCAGAGGCCCTGATCTTTAGAACAACAATGA    6076
Rascon13       CTTTTGTAAAAGCTCTAAAAAAACAGGGTCCAGAGGCCCTGATCTTTAGAACAACAATGA    6087
Rascon6        CTTTTGTAAAAGCTCTAAAAAAACAGGGTCCAGAGGCCCTGATCTTTAGAACAACAATGA    6089
Pachon14       CTTTTGTAAAAGCTCTAAAAAAACAGGGTCCAGAGGCCCTGATCTTTAGAACAACAATGA    6082
Pachon9        CTTTTGTAAAAGCTCTAAAAAAACAGGGTCCAGAGGCCCTGATCTTTAGAACAACAATGA    6091
Pachon17       CTTTTGTAAAAGCTCTAAAAAAACAGGGTCCAGAGGCCCTGATCTTTAGAACAACAATGA    6089
Pachon12       CTTTTGTAAAAGCTCTAAAAAAACAGGGTCCAGAGGCCCTGATCTTTAGAACAACAATGA    6089
Pachon11       CTTTTGTAAAAGCTCTAAAAAAACAGGGTCCAGAGGCCCTGATCTTTAGAACAACAATGA    6088
Pachon7        CTTTTGTAAAAGCTCTAAAAAAACAGGGTCCAGAGGCCCTGATCTTTAGAACAACAATGA    6089
Pachon3        CTTTTGTAAAAGCTCTAAAAAAACAGGGTCCAGAGGCCCTGATCTTTAGAACAACAATGA    6089
Pachon8        CTTTTGTAAAAGCTCTAAAAAAACAGGGTCCAGAGGCCCTGATCTTTAGAACAACAATGA    6089
Pachon15       CTTTTGTAAAAGCTCTAAAAAAACAGGGTCCAGAGGCCCTGATCTTTAGAACAACAATGA    6089
               * **********************************************************

Rascon4        TATTAACCCCAGGCAATAATATTTTATTATAATAAAAAATGAGTAGAAATAAATTACCTT    6087
Surface        TATTAACCCCAGGCAATAATATTTTATTATAATAAAAAATGAGTAGAAATAAATTACCTT    6167
Rascon8        TATTAACCCCAGGCAATAATATTTTATTATAATAAAAAATGAGTAGAAATAAATTACCTT    6150
Rascon2        TATTAACCCCAGGCAATAATATTTTATTATAATAAAAAATGAGTAGAAATAAATTACCTT    6153
Rascon15       TATTAACCCCAGGCAATAATATTTTATTATAATAAAAAATGAGTAGAAATAAATTACCTT    6136
Rascon13       TATTAACCCCAGGCAATAATATTTTATTATAATAAAAAATGAGTAGAAATAAATTACCTT    6147
```

```
Rascon6     TATTAACCCCAGGCAATAATATTTTATTATAATAAAAAATGAGTAGAAATAAATTACCTT     6149
Pachon14    TATTAACCCCAGGCAATAATATTTTATTATAATAAAAAATGAGTAGAAATAAATTACCTT     6142
Pachon9     TATTAACCCCAGGCAATAATATTTTATTATAATAAAAAATGAGTAGAAATAAATTACCTT     6151
Pachon17    TATTAACCCCAGGCAATAATATTTTATTATAATAAAAAATGAGTAGAAATAAATTACCTT     6149
Pachon12    TATTAACCCCAGGCAATAATATTTTATTATAATAAAAAATGAGTAGAAATAAATTACCTT     6149
Pachon11    TATTAACCCCAGGCAATAATATTTTATTATAATAAAAAATGAGTAGAAATAAATTACCTT     6148
Pachon7     TATTAACCCCAGGCAATAATATTTTATTATAATAAAAAATGAGTAGAAATAAATTACCTT     6149
Pachon3     TATTAACCCCAGGCAATAATATTTTATTATAATAAAAAATGAGTAGAAATAAATTACCTT     6149
Pachon8     TATTAACCCCAGGCAATAATATTTTATTATAATAAAAAATGAGTAGAAATAAATTACCTT     6149
Pachon15    TATTAACCCCAGGCAATAATATTTTATTATAATAAAAAATGAGTAGAAATAAATTACCTT     6149
            ************************************************************

Rascon4     ATTATATTTTACTATTTTGTTGTTTAAATAATATTTAATCATGTTTATACTAACGGGGAA     6147
Surface     ATTATATTTTACTATTTTGTTGTTTAAATAATATTTAATCATTTTTATACTAACGGGGAA     6227
Rascon8     ATTATATTTTACTATTTTGTTGTTTAAATAATATTTAATTATGTTTATACTAACGGGGAA     6210
Rascon2     ATTATATTTTACTATTTTGTTGTTTAAATAATATTTAATCATGTTTATACTAACAGGGAA     6213
Rascon15    ATTATATTTTACTATTTTGTTGTTTAAATAATATTTAATCATGTTTATACTAACAGGGAA     6196
Rascon13    ATTATATTTTACTATTTTGTTGTTTAAATAATATTTAATTATGTTTATACTAACGGGGAA     6207
Rascon6     ATTATATTTTACTATTTTGTTGTTTAAATAATATTTAATCATGTTTATACTAACGGGGAA     6209
Pachon14    ATTATATTTTACTATTTTGTTGTTTAAATAATATTTAATCATGTTTATACTAACGGGGAA     6202
Pachon9     ATTATATTTTACTATTTTGTTGTTTAAATAATATTTAATCATGTTTATACTAACGGGGAA     6211
Pachon17    ATTATATTTTACTATTTTGTTGTTTAAATAATATTTAATCATGTTTATACTAACGGGGAA     6209
Pachon12    ATTATATTTTACTATTTTGTTGTTTAAATAATATTTAATCATGTTTATACTAACGGGGAA     6209
Pachon11    ATTATATTTTACTATTTTGTTGTTTAAATAATATTTAATCATGTTTATACTAACGGGGAA     6208
Pachon7     ATTATATTTTACTATTTTGTTGTTTAAATAATATTTAATCATGTTTATACTAACGGGGAA     6209
Pachon3     ATTATATTTTACTATTTTGTTGTTTAAATAATATTTAATCATGTTTATACTAACGGGGAA     6209
Pachon8     ATTATATTTTACTATTTTGTTGTTTAAATAATATTTAATCATGTTTATACTAACGGGGAA     6209
Pachon15    ATTATATTTTACTATTTTGTTGTTTAAATAATATTTAATCATGTTTATACTAACGGGGAA     6209
            *************************************** ** *********** *****

Rascon4     TTTTAAGACTCTTTGAATCCGAATTGAAGAATCTGAAGAATCACACTCAATATAAACATT     6207
Surface     TTTTAAGACTCTTTGAATCCGAATTGAA----CTGAAGAATCACACTCAATATAAACACT     6283
Rascon8     TTTTAAGACTCTTTGAATCCGAATTGAA----CTGAAGAATCACACTCAATATAAACATT     6266
Rascon2     TTTTAAGACTCTTTGAATCCGAATTGAA----CTGAAGAATCACACTCAATATAAACATT     6269
Rascon15    TTTTAAGACTCTTTGAATCCGAATTGAA----CTGAAGAATCACACTCAATATAAACATT     6252
Rascon13    TTTTAAGACTCTTTGAATCCGAATTGAA----CTGAAGAATCACACTCAATATAAACATT     6263
Rascon6     TTTTAAGACTCTTTGAATCCGAATTGAA----CTGAAGAATCACACTCAATATAAACATT     6265
Pachon14    TTTTAAGACTCTTTGAATCCGAATTGAA----CTGAAGAATCACACTCAATATAAACACT     6258
Pachon9     TTTTAAGACTCTTTGAATCCGAATTGAA----CTGAAGAATCACACTCAATATAAACACT     6267
Pachon17    TTTTAAGACTCTTTGAATCCGAATTGAA----CTGAAGAATCACACTCAATATAAACACT     6265
Pachon12    TTTTAAGACTCTTTGAATCCGAATTGAA----CTGAAGAATCACACTCAATATAAACACT     6265
Pachon11    TTTTAAGACTCTTTGAATCCGAATTGAA----CTGAAGAATCACACTCAATATAAACACT     6264
Pachon7     TTTTAAGACTCTTTGAATCCGAATTGAA----CTGAAGAATCACACTCAATATAAACACT     6265
Pachon3     TTTTAAGACTCTTTGAATCCGAATTGAA----CTGAAGAATCACACTCAATATAAACACT     6265
Pachon8     TTTTAAGACTCTTTGAATCCGAATTGAA----CTGAAGAATCACACTCAATATAAACACT     6265
Pachon15    TTTTAAGACTCTTTGAATCCGAATTGAA----CTGAAGAATCACACTCAATATAAACACT     6265
            ****************************    ************************** *

Rascon4     TAAATAAGAAAAATTATAAGAAACAACGTTAATAATAAAGTGTCTGTGTGTCTACAGGTG     6267 Rx3
Surface     TAAATAAGAAAAATTATAAGAAACAACGTTAATAATAAAGTGTCTGTGTGTCTACAGGTG     6343 Exon 2
Rascon8     TAAATAAGAAAAATTATAAGAAACAACGTTAATAATAAAGTGTCTGTGTGTCTACAGGTG     6326 start
Rascon2     TAAATAAGAAAAATTATAAGAAACAACGTTAATAATAAAGTGTCTGTGTGTCTACAGGTG     6329
Rascon15    TAAATAAGAAAAATTATAAGAAACAACGTTAATAATAAAGTGTCTGTGTGTCTACAGGTG     6312
Rascon13    TAAATAAGAAAAATTATAAGAAACAACGTTAATAATAAAGTGTCTGTGTGTCTACAGGTG     6323
Rascon6     TAAATAAGAAAAATTATAAGAAACAATGTTAATAATAAAGTGTCTGTGTGTCTACAGGTG     6325
Pachon14    TAAATAAGAAAAATTATAAGAAACAACGTTAATAATAAAGTGTCTGTGTGTCTACAGGTG     6318
Pachon9     TAAATAAGAAAAATTATAAGAAACAACGTTAATAATAAAGTGTCTGTGTGTCTACAGGTG     6327
Pachon17    TAAATAAGAAAAATTATAAGAAACAACGTTAATAATAAAGTGTCTGTGTGTCTACAGGTG     6325
Pachon12    TAAATAAGAAAAATTATAAGAAACAACGTTAATAATAAAGTGTCTGTGTGTCTACAGGTG     6325
Pachon11    TAAATAAGAAAAATTATAAGAAACAACGTTAATAATAAAGTGTCTGTGTGTCTACAGGTG     6324
Pachon7     TAAATAAGAAAAATTATAAGAAACAACGTTAATAATAAAGTGTCTGTGTGTCTACAGGTG     6325
Pachon3     TAAATAAGAAAAATTATAAGAAACAACGTTAATAATAAAGTGTCTGTGTGTCTACAGGTG     6325
Pachon8     TAAATAAGAAAAATTATAAGAAACAACGTTAATAATAAAGTGTCTGTGTGTCTACAGGTG     6325
Pachon15    TAAATAAGAAAAATTATAAGAAACAACGTTAATAATAAAGTGTCTGTGTGTCTACAGGTG     6325
            ************************** *********************************

Rascon4     ACCCTTCCAGCCCGGACCGTAAGAAGCACCGGCGGAACAGAACCACCTTCACCACGTTTC     6327
Surface     ACCCTTCCAGCCCGGACCGTAAGAAGCACCGGCGGAACAGAACCACCTTCACCACGTTTC     6403
Rascon8     ACCCTTCCAGCCCGGACCGTAAGAAGCACCGGCGGAACAGAACCACCTTCACCACGTTTC     6386
Rascon2     ACCCTTCCAGCCCGGACCGTAAGAAGCACCGGCGGAACAGAACCACCTTCACCACGTTTC     6389
Rascon15    ACCCTTCCAGCCCGGACCGTAAGAAGCACCGGCGGAACAGAACCACCTTCACCACGTTTC     6372
Rascon13    ACCCTTCCAGCCCGGACCGTAAGAAGCACCGGCGGAACAGAACCACCTTCACCACGTTTC     6383
Rascon6     ACCCTTCCAGCCCGGACCGTAAGAAGCACCGGCGGAACAGAACCACCTTCACCACGTTTC     6385
Pachon14    ACCCTTCCAGCCCGGACCGTAAGAAGCACCGGCGGAACAGAACCACCTTCACCACGTTTC     6378
Pachon9     ACCCTTCCAGCCCGGACCGTAAGAAGCACCGGCGGAACAGAACCACCTTCACCACGTTTC     6387
Pachon17    ACCCTTCCAGCCCGGACCGTAAGAAGCACCGGCGGAACAGAACCACCTTCACCACGTTTC     6385
Pachon12    ACCCTTCCAGCCCGGACCGTAAGAAGCACCGGCGGAACAGAACCACCTTCACCACGTTTC     6385
```

```
Pachon11      ACCCTTCCAGCCCGGACCGTAAGAAGCACCGGCGGAACAGAACCACCTTCACCACGTTTC      6384
Pachon7       ACCCTTCCAGCCCGGACCGTAAGAAGCACCGGCGGAACAGAACCACCTTCACCACGTTTC      6385
Pachon3       ACCCTTCCAGCCCGGACCGTAAGAAGCACCGGCGGAACAGAACCACCTTCACCACGTTTC      6385
Pachon8       ACCCTTCCAGCCCGGACCGTAAGAAGCACCGGCGGAACAGAACCACCTTCACCACGTTTC      6385
Pachon15      ACCCTTCCAGCCCGGACCGTAAGAAGCACCGGCGGAACAGAACCACCTTCACCACGTTTC      6385
              ************************************************************

Rascon4       AGCTACACGAGCTGGAGAGAGCGTTCGAGAGGTCGCACTACCCCGACGTTTACAGCAGAG      6387
Surface       AGCTACACGAGCTGGAGAGAGCGTTCGAGAGGTCGCACTACCCCGACGTTTACAGCAGAG      6463
Rascon8       AGCTACACGAGTTGGAGAGAGCGTTCGAGAGGTCGCACTACCCCGACGTTTACAGCAGAG      6446
Rascon2       AGCTACACGAGCTGGAGAGAGCGTTCGAGAGGTCGCACTACCCCGACGTTTACAGCAGAG      6449
Rascon15      AGCTACACGAGCTGGAGAGAGCGTTCGAGAGGTCGCACTACCCCGACGTTTACAGCAGAG      6432
Rascon13      AGCTACACGAGCTGGAGAGAGCGTTCGAGAGGTCGCACTACCCCGACGTTTACAGCAGAG      6443
Rascon6       AGCTACACGAGCTGGAGAGAGCGTTCGAGAGGTCGCACTACCCCGACGTTTACAGCAGAG      6445
Pachon14      AGCTACACGAGCTGGAGAGAGCGTTCGAGAGGTCGCACTACCCCGACGTTTACAGCAGAG      6438
Pachon9       AGCTACACGAGCTGGAGAGAGCGTTCGAGAGGTCGCACTACCCCGACGTTTACAGCAGAG      6447
Pachon17      AGCTACACGAGCTGGAGAGAGCGTTCGAGAGGTCGCACTACCCCGACGTTTACAGCAGAG      6445
Pachon12      AGCTACACGAGCTGGAGAGAGCGTTCGAGAGGTCGCACTACCCCGACGTTTACAGCAGAG      6445
Pachon11      AGCTACACGAGCTGGAGAGAGCGTTCGAGAGGTCGCACTACCCCGACGTTTACAGCAGAG      6444
Pachon7       AGCTACACGAGCTGGAGAGAGCGTTCGAGAGGTCGCACTACCCCGACGTTTACAGCAGAG      6445
Pachon3       AGCTACACGAGCTGGAGAGAGCGTTCGAGAGGTCGCACTACCCCGACGTTTACAGCAGAG      6445
Pachon8       AGCTACACGAGCTGGAGAGAGCGTTCGAGAGGTCGCACTACCCCGACGTTTACAGCAGAG      6445
Pachon15      AGCTACACGAGCTGGAGAGAGCGTTCGAGAGGTCGCACTACCCCGACGTTTACAGCAGAG      6445
              *********** ************************************************

Rascon4       AAGAACTGGCGCTAAAGGTCAACCTGCCGGAAGTGCGCGTCCAGGTATGAAGCGGATTTA      6447 Rx3
Surface       AAGAACTGGCGCTAAAGGTCAACCTGCCGGAAGTGCGCGTCCAGGTATGAAGCGGATTTA      6523 exon 2
Rascon8       AAGAACTGGCGCTAAAGGTCAACCTGCCGGAAGTGCGCGTCCAGGTATGAAGCGGATTTA      6506 end
Rascon2       AAGAACTGGCGCTAAAGGTCAACCTGCCGGAAGTGCGCGTCCAGGTATGAAGCGGATTTA      6509
Rascon15      AAGAACTGGCGCTAAAGGTCAACCTGCCGGAAGTGCGCGTCCAGGTATGAAGCGGATTTA      6492
Rascon13      AAGAACTGGCGCTAAAGGTCAACCTGCCGGAAGTGCGCGTCCAGGTATGAAGCGGATTTA      6503
Rascon6       AAGAACTGGCGCTAAAGGTCAACCTGCCGGAAGTGCGCGTCCAGGTATGAAGCGGATTTA      6505
Pachon14      AAGAACTGGCGCTAAAGGTCAACCTGCCGGAAGTGCGCGTCCAGGTATGAAGCGGATTTA      6498
Pachon9       AAGAACTGGCGCTAAAGGTCAACCTGCCGGAAGTGCGCGTCCAGGTATGAAGCGGATTTA      6507
Pachon17      AAGAACTGGCGCTAAAGGTCAACCTGCCGGAAGTGCGCGTCCAGGTATGAAGCGGATTTA      6505
Pachon12      AAGAACTGGCGCTAAAGGTCAACCTGCCGGAAGTGCGCGTCCAGGTATGAAGCGGATTTA      6505
Pachon11      AAGAACTGGCGCTAAAGGTCAACCTGCCGGAAGTGCGCGTCCAGGTATGAAGCGGATTTA      6504
Pachon7       AAGAACTGGCGCTAAAGGTCAACCTGCCGGAAGTGCGCGTCCAGGTATGAAGCGGATTTA      6505
Pachon3       AAGAACTGGCGCTAAAGGTCAACCTGCCGGAAGTGCGCGTCCAGGTATGAAGCGGATTTA      6505
Pachon8       AAGAACTGGCGCTAAAGGTCAACCTGCCGGAAGTGCGCGTCCAGGTATGAAGCGGATTTA      6505
Pachon15      AAGAACTGGCGCTAAAGGTCAACCTGCCGGAAGTGCGCGTCCAGGTATGAAGCGGATTTA      6505
              ************************************************************

Rascon4       TATTCAGGATTTTTTATTAGTTA----GTTTTATCAAATTAATTTTTAAAGAAACCTTAA      6503 Rx3
Surface       TATTCAGGATTTTTTATTAGTTAGTTTGTTTTATCAAATTAATTTTTAAAGAAACCTTAA      6583 Intron 2
Rascon8       TATTCAGGATTTTTTATTAGTTAGTTTGTTTTATCAAATTAATTTTTAAAGAAACCTTAA      6566
Rascon2       TATTCAGGATTTTTTATTAGTTAGTTTGTTTTATCAAATTAATTTTTAAAGAAACCTTAA      6569
Rascon15      TATTCAGGATTTTTTATTAGTTAGTTTGTTTTATCAAATTGATTTTTAAAGAAACCTTAA      6552
Rascon13      TATTCAGGATTTTTTATTAGTTAGTTTGTTTTATCAAATTAATTTTTAAAGAAACCTTAA      6563
Rascon6       TATTCAGGATTTTTTATTAGTTAGTTTGTTTTATCAAATTAATTTTTAAAGAAACCTTAA      6565
Pachon14      TATTCAGGATTTTTTATTAGTTAGTTTGTTTTATCAAATTAATTTTTAAAGAAACCTTAA      6558
Pachon9       TATTCAGGATTTTTTATTAGTTAGTTTGTTTTATCAAATTAATTTTTAAAGAAACCTTAA      6567
Pachon17      TATTCAGGATTTTTTATTAGTTAGTTTGTTTTATCAAATTAATTTTTAAAGAAACCTTAA      6565
Pachon12      TATTCAGGATTTTTTATTAGTTAGTTTGTTTTATCAAATTAATTTTTAAAGAAACCTTAA      6565
Pachon11      TATTCAGGATTTTTTATTAGTTAGTTTGTTTTATCAAATTAATTTTTAAAGAAACCTTAA      6564
Pachon7       TATTCAGGATTTTTTATTAGTTAGTTTGTTTTATCAAATTAATTTTTAAAGAAACCTTAA      6565
Pachon3       TATTCAGGATTTTTTATTAGTTAGTTTGTTTTATCAAATTAATTTTTAAAGAAACCTTAA      6565
Pachon8       TATTCAGGATTTTTTATTAGTTAGTTTGTTTTATCAAATTAATTTTTAAAGAAACCTTAA      6565
Pachon15      TATTCAGGATTTTTTATTAGTTAGTTTGTTTTATCAAATTAATTTTTAAAGAAACCTTAA      6565
              ***********************    ************* *******************

Rascon4       TTAAATGCATGTCATTTGTCCTCTGCAGTCAGTGTTTAACATAGCATGTAATAGCTAGTA      6563
Surface       TTAAATGCATGTCATTTGTCCTCTGCAGTCAGTGTTTAACATAGCATGTAATAGCTAGTA      6643
Rascon8       TTAAATGCATGTCATTTGTCCTCTGCAATCAGTGTTTAACATAGCATGTAATAGCTAGTA      6626
Rascon2       TTAAATGCATGTCATTTGTCCTCTGCAGTCAGTGTTTAACATAGCATGTAATAGCTAGTA      6629
Rascon15      TTAAATGCATGTCATTTGTCCTCTGCAGTCAATGTTTAACATAGCATGTAATAGCTAGTA      6612
Rascon13      TTAAATGCATGTCATTTGTCCTCTGCAGTCAGTGTTTAACATAGCATGTAATAGCTAGTA      6623
Rascon6       TTAAATGCATGTCATTTGTCCTCTGCAATCAGTGTTTAACATAGCATGTAATAGCTAGTA      6625
Pachon14      TTAAATGCATGTCATTTGTCCTCTGCAGTCAGTGTTTAACATAGCATGTAATAGCTAGTA      6618
Pachon9       TTAAATGCATGTCATTTGTCCTCTGCAGTCAGTGTTTAACATAGCATGTAATAGCTAGTA      6627
Pachon17      TTAAATGCATGTCATTTGTCCTCTGCAGTCAGTGTTTAACATAGCATGTAATAGCTAGTA      6625
Pachon12      TTAAATGCATGTCATTTGTCCTCTGCAGTCAGTGTTTAACATAGCATGTAATAGCTAGTA      6625
Pachon11      TTAAATGCATGTCATTTGTCCTCTGCAGTCAGTGTTTAACATAGCATGTAATAGCTAGTA      6624
Pachon7       TTAAATGCATGTCATTTGTCCTCTGCAGTCAGTGTTTAACATAGCATGTAATAGCTAGTA      6625
Pachon3       TTAAATGCATGTCATTTGTCCTCTGCAGTCAGTGTTTAACATAGCATGTAATAGCTAGTA      6625
Pachon8       TTAAATGCATGTCATTTGTCCTCTGCAGTCAGTGTTTAACATAGCATGTAATAGCTAGTA      6625
Pachon15      TTAAATGCATGTCATTTGTCCTCTGCAGTCAGTGTTTAACATAGCATGTAATAGCTAGTA      6625
```

```
                    *************************** *** ****************************

Rascon4             ATACCT--AAAAAAACTAATTTTATTTATTTATTAAAAATGTATAACTCATTTAAAAACG          6621 Rx3
Surface             ATACCT--AAAAAAACTAATTTTATTTATTTATTAAAAATGTATAACTCATTTAAAAACG          6701 Intron 2
Rascon8             ATACCTAAAAAAAAACTAATTTTATTTATTTATTAAAAATGTATAACTCATTTAAAAACG          6686
Rascon2             ATACCT-AAAAAAAACTAATTTTATTTATTTATTAAAAATGTATAACTCATTTAAAAATG          6688
Rascon15            ATACCT-AAAAAAAACTAATTTTATTTATTTATTAAAAATGTATAACTCATTTAAAAATG          6671
Rascon13            ATACCTAAAAAAAAACTAATTTTATTTATTTATTAAAAATGTATAACTCATTTAAAAATG          6683
Rascon6             ATACCTAAAAAAAAACTAATTTTATTTATTTATTAAAAATGTATAACTCATTTAAAAATG          6685
Pachon14            ATACCT--AAAAAAACTAATTTTATTTATTTATTAAAAATGTATAACTCATTTAAAAACG          6676
Pachon9             ATACCT--AAAAAAACTAATTTTATTTATTTATTAAAAATGTATAACTCATTTAAAAACG          6685
Pachon17            ATACCT--AAAAAAACTAATTTTATTTATTTATTAAAAATGTATAACTCATTTAAAAACG          6683
Pachon12            ATACCT--AAAAAAACTAATTTTATTTATTTATTAAAAATGTATAACTCATTTAAAAACG          6683
Pachon11            ATACCT--AAAAAAACTAATTTTATTTATTTATTAAAAATGTATAACTCATTTAAAAACG          6682
Pachon7             ATACCT--AAAAAAACTAATTTTATTTATTTATTAAAAATGTATAACTCATTTAAAAACG          6683
Pachon3             ATACCT--AAAAAAACTAATTTTATTTATTTATTAAAAATGTATAACTCATTTAAAAACG          6683
Pachon8             ATACCT--AAAAAAACTAATTTTATTTATTTATTAAAAATGTATAACTCATTTAAAAACG          6683
Pachon15            ATACCT--AAAAAAACTAATTTTATTTATTTATTAAAAATGTATAACTCATTTAAAAACG          6683
                    ******  ************************************************** *

Rascon4             CTTATATGATAATTTATTAGTTTATTAAGAATTCTTCAGTTGAAGAATAAAAATTCATTT          6681
Surface             CTTATATGATAATTTATTAGTTTATTAAGAATTCTTCAGTTGAAGAATAAAAATTCATTT          6761
Rascon8             CTTATATGATAATTTATTAGTTTATTAAGAATTCTTCAGTTGAAGAATAAAAATTCATTT          6746
Rascon2             CTTATATGATAATTTATTAGTTTATTAAGAATTCTTCAGTTGAAGAATAAAAATTCATTT          6748
Rascon15            CTTATATGATAATTTATTAGTTTATTAAGAATTCTTCAGTTGAAGAATAAAAATTCATTT          6731
Rascon13            CTTATATGATAATTTATTAGTTTATTAAGAATTCTTCAGTTGAAGAATAAAAATTCATTT          6743
Rascon6             CTTATATGATAATTTATTAGTTTATTAAGAATTCTTCAGTTGAAGAATAAAAATTCATTT          6745
Pachon14            CTTATATGATAATTTATTAGTTTATTAAGAATTCTTCAGTTGAAGAATAAAAATTCATTT          6736
Pachon9             CTTATATGATAATTTATTAGTTTATTAAGAATTCTTCAGTTGAAGAATAAAAATTCATTT          6745
Pachon17            CTTATATGATAATTTATTAGTTTATTAAGAATTCTTCAGTTGAAGAATAAAAATTCATTT          6743
Pachon12            CTTATATGATAATTTATTAGTTTATTAAGAATTCTTCAGTTGAAGAATAAAAATTCATTT          6743
Pachon11            CTTATATGATAATTTATTAGTTTATTAAGAATTCTTCAGTTGAAGAATAAAAATTCATTT          6742
Pachon7             CTTATATGATAATTTATTAGTTTATTAAGAATTCTTCAGTTGAAGAATAAAAATTCATTT          6743
Pachon3             CTTATATGATAATTTATTAGTTTATTAAGAATTCTTCAGTTGAAGAATAAAAATTCATTT          6743
Pachon8             CTTATATGATAATTTATTAGTTTATTAAGAATTCTTCAGTTGAAGAATAAAAATTCATTT          6743
Pachon15            CTTATATGATAATTTATTAGTTTATTAAGAATTCTTCAGTTGAAGAATAAAAATTCATTT          6743
                    ************************************************************

Rascon4             CTGGCGGTTTATGGGGTTAATAAAGTGGGTAAACAGTACACTGATTTAATTATTAGATTT          6741
Surface             ATGGCGGTTTATGGGGTTAATAAAGTGGGTAAACAGTACACTGATTTAATTATTAGATTT          6821
Rascon8             ATGGCGGTTTATGGGGTTAATAAAGTGGGTAAACAGTACACTGATTTAATTATTAGATTT          6806
Rascon2             ATGGCGGTTTATGGGGTTAATAAAGTGGGTAAACAGTACACTGATTTAATTATTAGATTT          6808
Rascon15            ATGGCGGTTTATGGGGTTAATAAAGTGGGTAAACAGTACACTGATTTAATTATTAGATTT          6791
Rascon13            ATGGCGGTTTATGGGGTTAATAAAGTTGGTAAACAGTACACTGATTTAATTATTAGATTT          6803
Rascon6             ATGGCGGTTTATGGGGTTAATAAAGTGGGTAAACAGTACACTGATTTAATTATTAGATTT          6805
Pachon14            ATGGCGGTTTATGGGGTTAATAAAGTGGGTAAACAGTACACTGATTTAATTATTAGATTT          6796
Pachon9             ATGGCGGTTTATGGGGTTAATAAAGTGGGTAAACAGTACACTGATTTAATTATTAGATTT          6805
Pachon17            ATGGCGGTTTATGGGGTTAATAAAGTGGGTAAACAGTACACTGATTTAATTATTAGATTT          6803
Pachon12            ATGGCGGTTTATGGGGTTAATAAAGTGGGTAAACAGTACACTGATTTAATTATTAGATTT          6803
Pachon11            ATGGCGGTTTATGGGGTTAATAAAGTGGGTAAACAGTACACTGATTTAATTATTAGATTT          6802
Pachon7             ATGGCGGTTTATGGGGTTAATAAAGTGGGTAAACAGTACACTGATTTAATTATTAGATTT          6803
Pachon3             ATGGCGGTTTATGGGGTTAATAAAGTGGGTAAACAGTACACTGATTTAATTATTAGATTT          6803
Pachon8             ATGGCGGTTTATGGGGTTAATAAAGTGGGTAAACAGTACACTGATTTAATTATTAGATTT          6803
Pachon15            ATGGCGGTTTATGGGGTTAATAAAGTGGGTAAACAGTACACTGATTTAATTATTAGATTT          6803
                     ************************* *********************************

Rascon4             TAGCCTTATTTAATTGTATGACTAATGACTTCTCTTAATTTCTTTTTTATTATTATTATT          6801
Surface             TAGCCTTATTTAATTGTATGACTAATGACTTCTCTTAATTTCTTTTTTATTATTATTATT          6881
Rascon8             TAGCCTTATTTAATTGTATGACTAATGACTTCTCTTAATTTCTTTTTTATTATTATTATT          6866
Rascon2             TAGCCTTATTTAATTGTATGACTAATGACTTCTCTTAATTTCTTTTTTATTATTATTATT          6868
Rascon15            TAGCCTTATTTAATTGTATGACTAATGACTTCTCTTAATTTCTTTTTTATTATTATTATT          6851
Rascon13            TAGCCTTATTTAATTGTATGACTAATGACTTCTCTTAATTTCTTTTTTATTATTATTATT          6863
Rascon6             TAGCCTTATTTAATTGTATGACTAATGACTTCTCTTAATTTCTTTTTTATTATTATTATT          6865
Pachon14            TAGCCTTATTTAATTGTATGACTAATGACTTCTCTTAATTTCTTTTTTATTATTATTATT          6856
Pachon9             TAGCCTTATTTAATTGTATGACTAATGACTTCTCTTAATTTCTTTTTTATTATTATTATT          6865
Pachon17            TAGCCTTATTTAATTGTATGACTAATGACTTCTCTTAATTTCTTTTTTATTATTATTATT          6863
Pachon12            TAGCCTTATTTAATTGTATGACTAATGACTTCTCTTAATTTCTTTTTTATTATTATTATT          6863
Pachon11            TAGCCTTATTTAATTGTATGACTAATGACTTCTCTTAATTTCTTTTTTATTATTATTATT          6862
Pachon7             TAGCCTTATTTAATTGTATGACTAATGACTTCTCTTAATTTCTTTTTTATTATTATTATT          6863
Pachon3             TAGCCTTATTTAATTGTATGACTAATGACTTCTCTTAATTTCTTTTTTATTATTATTATT          6863
Pachon8             TAGCCTTATTTAATTGTATGACTAATGACTTCTCTTAATTTCTTTTTTATTATTATTATT          6863
Pachon15            TAGCCTTATTTAATTGTATGACTAATGACTTCTCTTAATTTCTTTTTTATTATTATTATT          6863
                    ************************************************************

Rascon4             CAGTAATTACTGGTTATTCGCAATATAACTACTAAATTACCTTTTTAACTTTCATTAAAT          6861
Surface             CAGTAATTACTGGTTATTCGCAATATAACTACTAAATTACCTTTTTAACTTTCATTAAAT          6941
Rascon8             CAGTAATTACTGGTTATTCGCAATATAACTACTAAATTACCTTTTTAACTTTCATTAAAT          6926
```

```
Rascon2      CAGTAATTACTGGTTATTCGCAATATAACTACTAAATTACCTTTTTAACTTTCATTAAAT   6928
Rascon15     CAGTAATTACTGGTTATTCGCAATATAACTACTAAATTACCTTTTTAACTTTCATTAAAT   6911
Rascon13     CAGTAATTACTGGTTATTCGCAATATAACTACTAAATTACCTTTTTAACTTTCATTAAAT   6923
Rascon6      CAGTAATTACTGGTTATTCGCAATATAACTACTAAATTACCTTTTTAACTTTCATTAAAT   6925
Pachon14     CAGTAATTACTGGTTATTCGCAATATAACTACTAAATTACCTTTTTAACTTTCATTAAAT   6916
Pachon9      CAGTAATTACTGGTTATTCGCAATATAACTACTAAATTACCTTTTTAACTTTCATTAAAT   6925
Pachon17     CAGTAATTACTGGTTATTCGCAATATAACTACTAAATTACCTTTTTAACTTTCATTAAAT   6923
Pachon12     CAGTAATTACTGGTTATTCGCAATATAACTACTAAATTACCTTTTTAACTTTCATTAAAT   6923
Pachon11     CAGTAATTACTGGTTATTCGCAATATAACTACTAAATTACCTTTTTAACTTTCATTAAAT   6922
Pachon7      CAGTAATTACTGGTTATTCGCAATATAACTACTAAATTACCTTTTTAACTTTCATTAAAT   6923
Pachon3      CAGTAATTACTGGTTATTCGCAATATAACTACTAAATTACCTTTTTAACTTTCATTAAAT   6923
Pachon8      CAGTAATTACTGGTTATTCGCAATATAACTACTAAATTACCTTTTTAACTTTCATTAAAT   6923
Pachon15     CAGTAATTACTGGTTATTCGCAATATAACTACTAAATTACCTTTTTAACTTTCATTAAAT   6923
             ************************************************************

Rascon4      ATTTAATTCCAAGTAATTTCGACACAATGTAAAAACAACTACCTGGTTCACATTATGTCA   6921 Rx3
Surface      ATTTAATTCCAAGTCATTTCGACACAATGTAAAAACAACTACCTGGTTCACATTATGTCA   7001 Intron 2
Rascon8      ATTTAATTCCAAGTAATTTCGACACAATGTAAAAACAACTACCTGGTTCACATTATGTCA   6986
Rascon2      ATTTAATTCCAAGTAATTTCGACACAATGTAAAAACAACTACCTGGTTCACATTATGTCA   6988
Rascon15     ATTTAATTCCAAGTAATTTCGACACAATGTAAAAACAACTACCTGGTTCACATTATGTCA   6971
Rascon13     ATTTAATTCCAAGTAATTTCGACACAATGTAAAAACAACTACCTGGTTCACATTATGTCA   6983
Rascon6      ATTTAATTCCAAGTAATTTCGACACAATGTAAAAACAACTACCTGGTTCACATTATGTCA   6985
Pachon14     ATTTAATTCCAAGTAATTTCGACACAATGTAAAAACAACTACCTGGTTCACATTATGTCA   6976
Pachon9      ATTTAATTCCAAGTAATTTCGACACAATGTAAAAACAACTACCTGGTTCACATTATGTCA   6985
Pachon17     ATTTAATTCCAAGTAATTTCGACACAATGTAAAAACAACTACCTGGTTCACATTATGTCA   6983
Pachon12     ATTTAATTCCAAGTAATTTCGACACAATGTAAAAACAACTACCTGGTTCACATTATGTCA   6983
Pachon11     ATTTAATTCCAAGTAATTTCGACACAATGTAAAAACAACTACCTGGTTCACATTATGTCA   6982
Pachon7      ATTTAATTCCAAGTAATTTCGACACAATGTAAAAACAACTACCTGGTTCACATTATGTCA   6983
Pachon3      ATTTAATTCCAAGTAATTTCGACACAATGTAAAAACAACTACCTGGTTCACATTATGTCA   6983
Pachon8      ATTTAATTCCAAGTAATTTCGACACAATGTAAAAACAACTACCTGGTTCACATTATGTCA   6983
Pachon15     ATTTAATTCCAAGTAATTTCGACACAATGTAAAAACAACTACCTGGTTCACATTATGTCA   6983
             ************** *********************************************

Rascon4      AAAACGGAAAAAATACAAAATTGAGATAAAATATTGATATTTGCTCTGTAGCTATAAAAT   6981
Surface      AAAACGGAAAAAATACAAAATTGAGATAAAATATTGATATTTGCTCTGTAGCTATAAAAT   7061
Rascon8      AAAACGGAAAAAATACAAAATTGAGATAAAATATTGATATTTGCTCTGTAGCTATAAAAT   7046
Rascon2      AAAACGGAAAAAATACAAAATTGAGATACAATATTGATATTTGCTCTGTAGCTATAAAAT   7048
Rascon15     AAAACGGAAAAAATACAAAATTGAGATAAAATATTGATATTTGCTCTGTAGCTATAAAAT   7031
Rascon13     AAAACGGAAAAAATACAAAATTGAGATAAAATATTGATATTTGCTCTGTAGCTATAAAAT   7043
Rascon6      AAAACGGAAAAAATACAAAATTGAGATAAAATATTGATATTTGCTCTGTAGCTATAAAAT   7045
Pachon14     AAAACGGAAAAAATACAAAATTGAGATAAAATATTGATATTTGCTCTGTAGCTATAAAAT   7036
Pachon9      AAAACGGAAAAAATACAAAATTGAGATAAAATATTGATATTTGCTCTGTAGCTATAAAAT   7045
Pachon17     AAAACGGAAAAAATACAAAATTGAGATAAAATATTGATATTTGCTCTGTAGCTATAAAAT   7043
Pachon12     AAAACGGAAAAAATACAAAATTGAGATAAAATATTGATATTTGCTCTGTAGCTATAAAAT   7043
Pachon11     AAAACGGAAAAAATACAAAATTGAGATAAAATATTGATATTTGCTCTGTAGCTATAAAAT   7042
Pachon7      AAAACGGAAAAAATACAAAATTGAGATAAAATATTGATATTTGCTCTGTAGCTATAAAAT   7043
Pachon3      AAAACGGAAAAAATACAAAATTGAGATAAAATATTGATATTTGCTCTGTAGCTATAAAAT   7043
Pachon8      AAAACGGAAAAAATACAAAATTGAGATAAAATATTGATATTTGCTCTGTAGCTATAAAAT   7043
Pachon15     AAAACGGAAAAAATACAAAATTGAGATAAAATATTGATATTTGCTCTGTAGCTATAAAAT   7043
             **************************** *******************************

Rascon4      GGGGTTTTGTAGAAATATGGAAACATCTGTACTGCATGAAAAGACACTCAATTAATGTGA   7041
Surface      GGGGTTTTGTAGAAATATGGAAGCATCTGTACTGCATGAAAAGACACTCAATTAATGTGA   7121
Rascon8      GGGGTTTTGTAGAAATATGGAAGCATCTGTACTGCATGAAAAGACACTCAATTAATGTGA   7106
Rascon2      GGGGTTTTGTAGAAATATGGAAGCATCTGTACTGCATGAAAAGACACTCAATTAATGTGA   7108
Rascon15     GGGGTTTTGTAGAAATATGGAAGCATCTGTACTGCATGAAAAGACACTCAATTAATGTGA   7091
Rascon13     GGGGTTTTGTAGAAATATGGAAGCATCTGTACTGCATGAAAAGACACTCAATTAATGTGA   7103
Rascon6      GGGGTTTTGTAGAAATATGGAAGCATCTGTACTGCATGAAAAGACACTCAATTAATGTGA   7105
Pachon14     GGGGTTTTGTAGAAATATGGAAGCATCTGTACTGCATGAAAAGACACTCAATTAATGTGA   7096
Pachon9      GGGGTTTTGTAGAAATATGGAAGCATCTGTACTGCATGAAAAGACACTCAATTAATGTGA   7105
Pachon17     GGGGTTTTGTAGAAATATGGAAGCATCTGTACTGCATGAAAAGACACTCAATTAATGTGA   7103
Pachon12     GGGGTTTTGTAGAAATATGGAAGCATCTGTACTGCATGAAAAGACACTCAATTAATGTGA   7103
Pachon11     GGGGTTTTGTAGAAATATGGAAGCATCTGTACTGCATGAAAAGACACTCAATTAATGTGA   7102
Pachon7      GGGGTTTTGTAGAAATATGGAAGCATCTGTACTGCATGAAAAGACACTCAATTAATGTGA   7103
Pachon3      GGGGTTTTGTAGAAATATGGAAGCATCTGTACTGCATGAAAAGACACTCAATTAATGTGA   7103
Pachon8      GGGGTTTTGTAGAAATATGGAAGCATCTGTACTGCATGAAAAGACACTCAATTAATGTGA   7103
Pachon15     GGGGTTTTGTAGAAATATGGAAGCATCTGTACTGCATGAAAAGACACTCAATTAATGTGA   7103
             ********************** *************************************

Rascon4      GTATTCCTTCCTTAAATAGGTTAAATAAAATCCTGTCTTTGTCAAACTCTTTATTAACCC   7101
Surface      GTATTCCTTCCTTAAATAGGTTAAATAAAATCCTGTCTTTGTCAAACTCTTTATTAACCC   7181
Rascon8      GTATTCCTTCCTTAAATAGGTTAAATAAAATCCTGTCTTTGTCAAACTCTTTATTAACCC   7166
Rascon2      GTATTCCTTCCTTAAATAGGTTAAATAAAATCCTGTCTTTGTCAAACTCTTTATTAACCC   7168
Rascon15     GTATTCCTTCCTTAAATAGGTTAAATAAAATCCTGTCTTTGTCAAACTCTTTATTAACCC   7151
Rascon13     GTATTCCTTCCTTAAATAGGTTAAATAAAATCCTGTCTTTGTCAAACTCTTTATTAACCC   7163
Rascon6      GTATTCCTTCCTTAAATAGGTTAAATAAAATCCTGTCTTTGTCAAACTCTTTATTAACCC   7165
Pachon14     GTATTCCTTCCTTAAATAGGTTAAATAAAATCCTGTCTTTGTCAAACTCTTTATTAACCC   7156
```

```
Pachon9      GTATTCCTTCCTTAAATAGGTTAAATAAAATCCTGTCTTTGTCAAACTCTTTATTAACCC      7165
Pachon17     GTATTCCTTCCTTAAATAGGTTAAATAAAATCCTGTCTTTGTCAAACTCTTTATTAACCC      7163 Rx3
Pachon12     GTATTCCTTCCTTAAATAGGTTAAATAAAATCCTGTCTTTGTCAAACTCTTTATTAACCC      7163 Intron 2
Pachon11     GTATTCCTTCCTTAAATAGGTTAAATAAAATCCTGTCTTTGTCAAACTCTTTATTAACCC      7162
Pachon7      GTATTCCTTCCTTAAATAGGTTAAATAAAATCCTGTCTTTGTCAAACTCTTTATTAACCC      7163
Pachon3      GTATTCCTTCCTTAAATAGGTTAAATAAAATCCTGTCTTTGTCAAACTCTTTATTAACCC      7163
Pachon8      GTATTCCTTCCTTAAATAGGTTAAATAAAATCCTGTCTTTGTCAAACTCTTTATTAACCC      7163
Pachon15     GTATTCCTTCCTTAAATAGGTTAAATAAAATCCTGTCTTTGTCAAACTCTTTATTAACCC      7163
             ************************************************************

Rascon4      CAGATTTTTTTTTTTTAAATCAATATGCTCTGTGGTTAGAGAATCAGGAATTATGATATA      7161 SNP3
Surface      CAGATTTTTTTAAAAAAAATCAATATGCTCTGTGGTTAGAGAATC-------ATGATATA      7234 Also
Rascon8      CAGATTTTTTTTTTAAAAATCAATATGCTCTGTGGTTAGAGAATCAGGAATTATGATATA      7226 fixed in
Rascon2      CAGATTTTTTTTTTAAAAATCAATATGCTCTGTGGTTAGAGAATCAGGAATTATGATATA      7228 Choy SF
Rascon15     CAGATTTTTTTTTTAAAAATCAATATGCTCTGTGGTTAGAGAATCAGGAATTATGATATA      7211
Rascon13     CAGATTTTTTTTTTAAAAATCAATATGCTCTGTGGTTAGAGAATCAGGAATTATGATATA      7223
Rascon6      CAGATTTTTTTTTTAAAAATCAATATGCTCTGTGGTTAGAGAATTAGGAATTATGATATA      7225
Pachon14     CAGATTTTATTTTAAAAAATCAATATGCTCTGTGGTTAGAGAATGAGGAATCATGATATA      7216
Pachon9      CAGATTTTATTTTAAAAAATCAATATGCTCTGTGGTTAGAGAATGAGGAATCATGATATA      7225
Pachon17     CAGATTTTATTTTAAAAAATCAATATGCTCTGTGGTTAGAGAATGAGGAATCATGATATA      7223
Pachon12     CAGATTTTATTTTAAAAAATCAATATGCTCTGTGGTTAGAGAATGAGGAATCATGATATA      7223
Pachon11     CAGATTTTATTTTAAAAAATCAATATGCTCTGTGGTTAGAGAATGAGGAATCATGATATA      7222
Pachon7      CAGATTTTATTTTAAAAAATCAATATGCTCTGTGGTTAGAGAATGAGGAATCATGATATA      7223
Pachon3      CAGATTTTATTTTAAAAAATCAATATGCTCTGTGGTTAGAGAATGAGGAATCATGATATA      7223
Pachon8      CAGATTTTATTTTAAAAAATCAATATGCTCTGTGGTTAGAGAATGAGGAATCATGATATA      7223
Pachon15     CAGATTTTATTTTAAAAAATCAATATGCTCTGTGGTTAGAGAATGAGGAATCATGATATA      7223
             ******** **     ****************************        ********

Rascon4      ATGGAACTCTGTATTAACCCTTCATCACTTTGTGCTTCTACAGGTGTGGTTCCAGAACCG      7221 Rx3
Surface      ATGGAACTCTGTATTAACCCTTCATCACTTTGTGCTTCTACAGGTGTGGTTCCAGAACCG      7294 exon 3
Rascon8      ATGGAACTCTGTATTAACCCTTCATCACTTTGTGCTTCTACAGGTGTGGTTCCAGAACCG      7286 start
Rascon2      ATGGAACTCTGTATTAACCCTTCATCACTTTGTGCTTCTACAGGTGTGGTTCCAGAACCG      7288
Rascon15     ATGGAACTCTGTATTAACCCTTCATCACTTTGTGCTTCTACAGGTGTGGTTCCAGAACCG      7271
Rascon13     ATGGAACTCTGTATTAACCCTTCATCACTTTGTGCTTCTACAGGTGTGGTTCCAGAACCG      7283
Rascon6      ATGGAACTCTGTATTAACCCTTCATCACTTTGTGCTTCTACAGGTGTGGTTCCAGAACCG      7285
Pachon14     ATGGAACTCTGTATTAACCCTTCATCACTTTGTGCTTCTACAGGTGTGGTTCCAGAACCG      7276
Pachon9      ATGGAACTCTGTATTAACCCTTCATCACTTTGTGCTTCTACAGGTGTGGTTCCAGAACCG      7285
Pachon17     ATGGAACTCTGTATTAACCCTTCATCACTTTGTGCTTCTACAGGTGTGGTTCCAGAACCG      7283
Pachon12     ATGGAACTCTGTATTAACCCTTCATCACTTTGTGCTTCTACAGGTGTGGTTCCAGAACCG      7283
Pachon11     ATGGAACTCTGTATTAACCCTTCATCACTTTGTGCTTCTACAGGTGTGGTTCCAGAACCG      7282
Pachon7      ATGGAACTCTGTATTAACCCTTCATCACTTTGTGCTTCTACAGGTGTGGTTCCAGAACCG      7283
Pachon3      ATGGAACTCTGTATTAACCCTTCATCACTTTGTGCTTCTACAGGTGTGGTTCCAGAACCG      7283
Pachon8      ATGGAACTCTGTATTAACCCTTCATCACTTTGTGCTTCTACAGGTGTGGTTCCAGAACCG      7283
Pachon15     ATGGAACTCTGTATTAACCCTTCATCACTTTGTGCTTCTACAGGTGTGGTTCCAGAACCG      7283
             ************************************************************

Rascon4      TCGTGCTAAGTGGCGTCGGCAGGAGAAGCTGGAGGTCAGCTCCATTAAACTCCAGGACTC      7281
Surface      TCGTGCTAAGTGGCGTCGGCAGGAGAAGCTGGAGGTCAGCTCCATTAAACTCCAGGACTC      7354
Rascon8      TCGTGCTAAGTGGCGTCGGCAGGAGAAGCTGGAGGTCAGCTCCATTAAACTCCAGGACTC      7346
Rascon2      TCGTGCTAAGTGGCGTCGGCAGGAGAAGCTGGAGGTCAGCTCCATTAAACTCCAGGACTC      7348
Rascon15     TCGTGCTAAGTGGCGTCGGCAGGAGAAGCTGGAGGTCAGCTCCATTAAACTCCAGGACTC      7331
Rascon13     TCGTGCTAAGTGGCGTCGGCAGGAGAAGCTGGAGGTCAGCTCCATTAAACTCCAGGACTC      7343
Rascon6      TCGTGCTAAGTGGCGTCGGCAGGAGAAGCTGGAGGTCAGCTCCATTAAACTCCAGGACTC      7345
Pachon14     TCGTGCTAAGTGGCGTCGGCAGGAGAAGCTGGAGGTCAGCTCCATTAAACTCCAGGACTC      7336
Pachon9      TCGTGCTAAGTGGCGTCGGCAGGAGAAGCTGGAGGTCAGCTCCATTAAACTCCAGGACTC      7345
Pachon17     TCGTGCTAAGTGGCGTCGGCAGGAGAAGCTGGAGGTCAGCTCCATTAAACTCCAGGACTC      7343
Pachon12     TCGTGCTAAGTGGCGTCGGCAGGAGAAGCTGGAGGTCAGCTCCATTAAACTCCAGGACTC      7343
Pachon11     TCGTGCTAAGTGGCGTCGGCAGGAGAAGCTGGAGGTCAGCTCCATTAAACTCCAGGACTC      7342
Pachon7      TCGTGCTAAGTGGCGTCGGCAGGAGAAGCTGGAGGTCAGCTCCATTAAACTCCAGGACTC      7343
Pachon3      TCGTGCTAAGTGGCGTCGGCAGGAGAAGCTGGAGGTCAGCTCCATTAAACTCCAGGACTC      7343
Pachon8      TCGTGCTAAGTGGCGTCGGCAGGAGAAGCTGGAGGTCAGCTCCATTAAACTCCAGGACTC      7343
Pachon15     TCGTGCTAAGTGGCGTCGGCAGGAGAAGCTGGAGGTCAGCTCCATTAAACTCCAGGACTC      7343
             ************************************************************

Rascon4      CTCCATCCTGTCTTTCTCCAGACCGGCAGGTCCTGGCCTGGGCAGCGGAGGCCTGCCACT      7341
Surface      CTCCATCCTGTCTTTCTCCAGACCGGCAGGTCCTGGCCTGGGCAGCGGAGGCCTGCCACT      7414
Rascon8      CTCCATCCTGTCTTTCTCCAGACCTGCAGGTCCTGGCCTGGGCAGCGGAGGCCTGCCACT      7406
Rascon2      CTCCATCCTGTCTTTCTCCAGACCGGCAGGTCCTGGCCTGGGCAGCGGAGGCCTGCCACT      7408
Rascon15     CTCCATCCTGTCTTTCTCCAGACCGGCAGGTCCTGGCCTGGGCAGCGGAGGCCTGCCACT      7391
Rascon13     CTCCATCCTGTCTTTCTCCAGACCGGCAGGTCCTGGCCTGGGCAGCGGAGGCCTGCCACT      7403
Rascon6      CTCCATCCTGTCTTTCTCCAGACCGGCAGGTCCTGGCCTGGGCAGCGGAGGCCTGCCACT      7405
Pachon14     CTCCATCCTGTCTTTCTCCAGACCGGCAGGTCCTGGCCTGGGCAGCGGAGGCCTGCCACT      7396
Pachon9      CTCCATCCTGTCTTTCTCCAGACCGGCAGGTCCTGGCCTGGGCAGCGGAGGCCTGCCACT      7405
Pachon17     CTCCATCCTGTCTTTCTCCAGACCGGCAGGTCCTGGCCTGGGCAGCGGAGGCCTGCCACT      7403
Pachon12     CTCCATCCTGTCTTTCTCCAGACCGGCAGGTCCTGGCCTGGGCAGCGGAGGCCTGCCACT      7403
Pachon11     CTCCATCCTGTCTTTCTCCAGACCGGCAGGTCCTGGCCTGGGCAGCGGAGGCCTGCCACT      7402
Pachon7      CTCCATCCTGTCTTTCTCCAGACCGGCAGGTCCTGGCCTGGGCAGCGGAGGCCTGCCACT      7403
```

```
Pachon3      CTCCATCCTGTCTTTCTCCAGACCGGCAGGTCCTGGCCTGGGCAGCGGAGGCCTGCCACT    7403
Pachon8      CTCCATCCTGTCTTTCTCCAGACCGGCAGGTCCTGGCCTGGGCAGCGGAGGCCTGCCACT    7403
Pachon15     CTCCATCCTGTCTTTCTCCAGACCGGCAGGTCCTGGCCTGGGCAGCGGAGGCCTGCCACT    7403
             ************************ ***********************************

Rascon4      GGATCCCTGGCTGACTGGGCCCATTTCAGGTCCTATCTCATCCCCTAGCTCACCCTTACA    7401 Rx3
Surface      GGATCCCTGGCTGACTGGGCCCATTTCAGGTCCTATCTCATCCCCTAGCTCACCCTTACA    7474 exon 3
Rascon8      GGATCCCTGGCTGACTGGGCCCATTTCAGGTCCTATCTCATCCCCTAGCTCACCCTTACA    7466
Rascon2      GGATCCCTGGCTGACTGGGCCCATTTCAGGTCCTATCTCATCCCCTAGCTCACCCTTACA    7468
Rascon15     GGATCCCTGGCTGACTGGGCCCATTTCAGGTCCTATCTCATCCCCTAGCTCACCCTTACA    7451
Rascon13     GGATCCCTGGCTGACTGGGCCCATTTCAGGTCCTATCTCATCCCCTAGCTCACCCTTACA    7463
Rascon6      GGATCCCTGGCTGACTGGGCCCATTTCAGGTCCTATCTCATCCCCTAGCTCACCCTTACA    7465
Pachon14     GGATCCCTGGCTGACTGGGCCCATTTCAGGTCCTATCTCATCCCCTAGCTCACCCTTACA    7456
Pachon9      GGATCCCTGGCTGACTGGGCCCATTTCAGGTCCTATCTCATCCCCTAGCTCACCCTTACA    7465
Pachon17     GGATCCCTGGCTGACTGGGCCCATTTCAGGTCCTATCTCATCCCCTAGCTCACCCTTACA    7463
Pachon12     GGATCCCTGGCTGACTGGGCCCATTTCAGGTCCTATCTCATCCCCTAGCTCACCCTTACA    7463
Pachon11     GGATCCCTGGCTGACTGGGCCCATTTCAGGTCCTATCTCATCCCCTAGCTCACCCTTACA    7462
Pachon7      GGATCCCTGGCTGACTGGGCCCATTTCAGGTCCTATCTCATCCCCTAGCTCACCCTTACA    7463
Pachon3      GGATCCCTGGCTGACTGGGCCCATTTCAGGTCCTATCTCATCCCCTAGCTCACCCTTACA    7463
Pachon8      GGATCCCTGGCTGACTGGGCCCATTTCAGGTCCTATCTCATCCCCTAGCTCACCCTTACA    7463
Pachon15     GGATCCCTGGCTGACTGGGCCCATTTCAGGTCCTATCTCATCCCCTAGCTCACCCTTACA    7463
             ************************************************************

Rascon4      GTCCCTGCCAGGCTTTATGGCCTCTCAGCCTGCCAGCTACACCCCTCCACCTTTCCTCAC    7461
Surface      GTCCCTGCCAGGCTTTATGGCCTCTCAGCCTGCCAGCTACACCCCTCCACCTTTCCTCAC    7534
Rascon8      GTCCCTGCCAGGCTTTATGGCCTCTCAGCCTGCCAGCTACACCCCTCCACCTTTCCTCAC    7526
Rascon2      GTCCCTGCCAGGCTTTATGGCCTCTCAGCCTGCCAGCTACACCCCTCCACCTTTCCTCAC    7528
Rascon15     GTCCCTGCCAGGCTTTATGGCCTCTCAGCCTGCCAGCTACACCCCTCCACCTTTCCTCAC    7511
Rascon13     GTCCCTGCCAGGCTTTATGGCCTCTCAGCCTGCCAGCTACACCCCTCCACCTTTCCTCAC    7523
Rascon6      GTCCCTGCCAGGCTTTATGGCCTCTCAGCCTGCCAGCTACACCCCTCCACCTTTCCTCAC    7525
Pachon14     GTCCCTGCCAGGCTTTATGGCCTCTCAGCCTGCCAGCTACACCCCTCCACCTTTCCTCAC    7516
Pachon9      GTCCCTGCCAGGCTTTATGGCCTCTCAGCCTGCCAGCTACACCCCTCCACCTTTCCTCAC    7525
Pachon17     GTCCCTGCCAGGCTTTATGGCCTCTCAGCCTGCCAGCTACACCCCTCCACCTTTCCTCAC    7523
Pachon12     GTCCCTGCCAGGCTTTATGGCCTCTCAGCCTGCCAGCTACACCCCTCCACCTTTCCTCAC    7523
Pachon11     GTCCCTGCCAGGCTTTATGGCCTCTCAGCCTGCCAGCTACACCCCTCCACCTTTCCTCAC    7522
Pachon7      GTCCCTGCCAGGCTTTATGGCCTCTCAGCCTGCCAGCTACACCCCTCCACCTTTCCTCAC    7523
Pachon3      GTCCCTGCCAGGCTTTATGGCCTCTCAGCCTGCCAGCTACACCCCTCCACCTTTCCTCAC    7523
Pachon8      GTCCCTGCCAGGCTTTATGGCCTCTCAGCCTGCCAGCTACACCCCTCCACCTTTCCTCAC    7523
Pachon15     GTCCCTGCCAGGCTTTATGGCCTCTCAGCCTGCCAGCTACACCCCTCCACCTTTCCTCAC    7523
             ************************************************************

Rascon4      CCACTCTCTACCTCACCTCGGTCCGGTGTGCCACCCTCCTCCTCCACCGCCGGCCTACCA    7521
Surface      CCACTCTCTACCTCACCTCGGTCCGGTGTGCCACCCTCCTCCTCCACCGCCGGCCTACCA    7594
Rascon8      CCACTCTCTACCTCACCTCGGTCCGGTGTGCCACCCTCCTCCTCCACCGCCGGCCTACCA    7586
Rascon2      CCACTCTCTACCTCACCTCGGTCCGGTGTGCCACCCTCCTCCTCCACCGCCGGCCTACCA    7588
Rascon15     CCACTCTCTACCTCACCTCGGTCCGGTGTGCCACCCTCCTCCTCCACCGCCGGCCTACCA    7571
Rascon13     CCACTCTCTACCTCACCTCGGTCCAGTGTGCCACCCTCCTCCTCCACCGCCGGCCTACCA    7583
Rascon6      CCACTCTCTACCTCACCTCGGTCCGGTGTGCCACCCTCCTCCTCCACCGCCGGCCTACCA    7585
Pachon14     CCACTCTCTACCTCACCTCGGTCCGGTGTGCCACCCTCCTCCTCCACCGCCGGCCTACCA    7576
Pachon9      CCACTCTCTACCTCACCTCGGTCCGGTGTGCCACCCTCCTCCTCCACCGCCGGCCTACCA    7585
Pachon17     CCACTCTCTACCTCACCTCGGTCCGGTGTGCCACCCTCCTCCTCCACCGCCGGCCTACCA    7583
Pachon12     CCACTCTCTACCTCACCTCGGTCCGGTGTGCCACCCTCCTCCTCCACCGCCGGCCTACCA    7583
Pachon11     CCACTCTCTACCTCACCTCGGTCCGGTGTGCCACCCTCCTCCTCCACCGCCGGCCTACCA    7582
Pachon7      CCACTCTCTACCTCACCTCGGTCCGGTGTGCCACCCTCCTCCTCCACCGCCGGCCTACCA    7583
Pachon3      CCACTCTCTACCTCACCTCGGTCCGGTGTGCCACCCTCCTCCTCCACCGCCGGCCTACCA    7583
Pachon8      CCACTCTCTACCTCACCTCGGTCCGGTGTGCCACCCTCCTCCTCCACCGCCGGCCTACCA    7583
Pachon15     CCACTCTCTACCTCACCTCGGTCCGGTGTGCCACCCTCCTCCTCCACCGCCGGCCTACCA    7583
             ************************ ***********************************

Rascon4      GTGCTCAGGCTTCATGGACAAACTCAGCCTGGAGGAGACGGACCCACGCAACTCCAGCAT    7581
Surface      GTGCTCAGGCTTCATGGACAAACTCAGCCTGGAGGAGACGGACCCACGCAACTCCAGCAT    7654
Rascon8      GTGCTCAGGCTTCATGGACAAACTCAGCCTGGAGGAGACGGACCCACGCAACTCCAGCAT    7646
Rascon2      GTGCTCAGGCTTCATGGACAAACTCAGCCTGGAGGAGACGGACCCACGCAACTCCAGCAT    7648
Rascon15     GTGCTCAGGCTTCATGGACAAACTCAGCCTGGAGGAGACGGACCCACGCAACTCCAGCAT    7631
Rascon13     GTGCTCAGGCTTCATGGACAAACTCAGCCTGGAGGAGACGGACCCACGCAACTCCAGCAT    7643
Rascon6      GTGCTCAGGCTTCATGGACAAACTCAGCCTGGAGGAGACGGACCCACGCAACTCCAGCAT    7645
Pachon14     GTGCTCAGGCTTCATGGACAAACTCAGCCTGGAGGAGACGGACCCACGCAACTCCAGCAT    7636
Pachon9      GTGCTCAGGCTTCATGGACAAACTCAGCCTGGAGGAGACGGACCCACGCAACTCCAGCAT    7645
Pachon17     GTGCTCAGGCTTCATGGACAAACTCAGCCTGGAGGAGACGGACCCACGCAACTCCAGCAT    7643
Pachon12     GTGCTCAGGCTTCATGGACAAACTCAGCCTGGAGGAGACGGACCCACGCAACTCCAGCAT    7643
Pachon11     GTGCTCAGGCTTCATGGACAAACTCAGCCTGGAGGAGACGGACCCACGCAACTCCAGCAT    7642
Pachon7      GTGCTCAGGCTTCATGGACAAACTCAGCCTGGAGGAGACGGACCCACGCAACTCCAGCAT    7643
Pachon3      GTGCTCAGGCTTCATGGACAAACTCAGCCTGGAGGAGACGGACCCACGCAACTCCAGCAT    7643
Pachon8      GTGCTCAGGCTTCATGGACAAACTCAGCCTGGAGGAGACGGACCCACGCAACTCCAGCAT    7643
Pachon15     GTGCTCAGGCTTCATGGACAAACTCAGCCTGGAGGAGACGGACCCACGCAACTCCAGCAT    7643
             ************************************************************
```

```
Rascon4      CGCATCGCTGCGGATGAAGGCAAAGGAGCACATTCAGTCTATAGGGAAGACATGGTAAAC   7641 Rx3
Surface      CGCATCGCTGCGGATGAAGGCAAAGGAGCACATTCAGTCTATAGGGAAGACATGGTAAAC   7714 exon 3
Rascon8      CGCATCGCTGCGGATGAAGGCAAAGGAGCACATTCAGTCTATAGGGAAGACATGGTAAAC   7706 end
Rascon2      CGCATCGCTGCGGATGAAGGCAAAGGAGCACATTCAGTCTATAGGGAAGACATGGTAAAC   7708
Rascon15     CGCATCGCTGCGGATGAAGGCAAAGGAGCACATTCAGTCTATAGGGAAGACATGGTAAAC   7691
Rascon13     CGCATCGCTGCGGATGAAGGCAAAGGAGCACATTCAGTCTATAGGGAAGACATGGTAAAC   7703
Rascon6      CGCATCGCTGCGGATGAAGGCAAAGGAGCACATTCAGTCTATAGGGAAGACATGGTAAAC   7705
Pachon14     CGCATCGCTGCGGATGAAGGCAAAGGAGCACATTCAGTCTATAGGGAAGACATGGTAAAC   7696
Pachon9      CGCATCGCTGCGGATGAAGGCAAAGGAGCACATTCAGTCTATAGGGAAGACATGGTAAAC   7705
Pachon17     CGCATCGCTGCGGATGAAGGCAAAGGAGCACATTCAGTCTATAGGGAAGACATGGTAAAC   7703
Pachon12     CGCATCGCTGCGGATGAAGGCAAAGGAGCACATTCAGTCTATAGGGAAGACATGGTAAAC   7703
Pachon11     CGCATCGCTGCGGATGAAGGCAAAGGAGCACATTCAGTCTATAGGGAAGACATGGTAAAC   7702
Pachon7      CGCATCGCTGCGGATGAAGGCAAAGGAGCACATTCAGTCTATAGGGAAGACATGGTAAAC   7703
Pachon3      CGCATCGCTGCGGATGAAGGCAAAGGAGCACATTCAGTCTATAGGGAAGACATGGTAAAC   7703
Pachon8      CGCATCGCTGCGGATGAAGGCAAAGGAGCACATTCAGTCTATAGGGAAGACATGGTAAAC   7703
Pachon15     CGCATCGCTGCGGATGAAGGCAAAGGAGCACATTCAGTCTATAGGGAAGACATGGTAAAC   7703
             ************************************************************

Rascon4      ATCAGAACTTCAGAGACTGTGACGTTGCACTGTGACAGACTGGGAGCACTGGCCTATTGG   7701 Rx3
Surface      ATCAGAACTTCAGAGACTGTGACGTTGCACTGTGACAGACTGGGAGCACTGGCCTATTGG   7774 3'-UTR
Rascon8      ATCAGAACTTCAGAGACTGTGACGTTGCACTGTGACAGACTGGGAGCACTGGCCTATTGG   7766
Rascon2      ATCAGAACTTCAGAGACTGTGACGTTGCACTGTGACAGACTGGGAGCACTGGCCTATTGG   7768
Rascon15     ATCAGAACTTCAGAGACTGTGACGTTGCACTGTGACAGACTGGGAGCACTGGCCTATTGG   7751
Rascon13     ATCAGAACTTCAGAGACTGTGACGTTGCACTGTGACAGACTGGGAGCACTGGCCTATTGG   7763 SNP4
Rascon6      ATCAGAACTTCAGAGACTGTGACGTTGCACTGTGACAGACTGGGAGCACTGGCCTATTGG   7765 used in
Pachon14     ATCAGAACTTCAGAGACTGTGACTTTGCACTGTGACAGACTGGGAGCACTGGCCTATTGG   7756 FreeBayes
Pachon9      ATCAGAACTTCAGAGACTGTGACTTTGCACTGTGACAGACTGGGAGCACTGGCCTATTGG   7765
Pachon17     ATCAGAACTTCAGAGACTGTGACTTTGCACTGTGACAGACTGGGAGCACTGGCCTATTGG   7763
Pachon12     ATCAGAACTTCAGAGACTGTGACTTTGCACTGTGACAGACTGGGAGCACTGGCCTATTGG   7763
Pachon11     ATCAGAACTTCAGAGACTGTGACTTTGCACTGTGACAGACTGGGAGCACTGGCCTATTGG   7762
Pachon7      ATCAGAACTTCAGAGACTGTGACTTTGCACTGTGACAGACTGGGAGCACTGGCCTATTGG   7763
Pachon3      ATCAGAACTTCAGAGACTGTGACTTTGCACTGTGACAGACTGGGAGCACTGGCCTATTGG   7763
Pachon8      ATCAGAACTTCAGAGACTGTGACTTTGCACTGTGACAGACTGGGAGCACTGGCCTATTGG   7763
Pachon15     ATCAGAACTTCAGAGACTGTGACTTTGCACTGTGACAGACTGGGAGCACTGGCCTATTGG   7763
             *********************** ************************************

Rascon4      ACTCCAAAAGGGTCATTCAGCAGAGCTGGACCTATTGAACAGTGTGTATTTTGTTATAAA   7761
Surface      ACTCCAAAAGGGTCATTCAGCAGAGCTGGACCTATTGAACAGTGTGTATTTTGTTATAAA   7834
Rascon8      ACTCCAAAAGGGTCATTCAGCAGAGCTGGACCTATTGAACAGTGTGTATTTTGTTATAAA   7826
Rascon2      ACTCCAAAAGGGTCATTCAGCAGAGCTGGACCTATTGAACAGTGTGTATTTTGTTATAAA   7828
Rascon15     ACTCCAAAAGGGTCATTCAGCAGAGCTGGACCTATTGAACAGTGTGTATTTTGTTATAAA   7811
Rascon13     ACTCCAAAAGGGTCATTCAGCAGAGCTGGACCTATTGAACAGTGTGTATTTTGTTATAAA   7823
Rascon6      ACTCCAAAAGGGTCATTCAGCAGAGCTGGACCTATTGAACAGTGTGTATTTTGTTATAAA   7825
Pachon14     ACTCCAAAAGGGTCATTCAGCAGAGCTGGACCTATTGAACAGTGTGTATTTTGTTATAAA   7816
Pachon9      ACTCCAAAAGGGTCATTCAGCAGAGCTGGACCTATTGAACAGTGTGTATTTTGTTATAAA   7825
Pachon17     ACTCCAAAAGGGTCATTCAGCAGAGCTGGACCTATTGAACAGTGTGTATTTTGTTATAAA   7823
Pachon12     ACTCCAAAAGGGTCATTCAGCAGAGCTGGACCTATTGAACAGTGTGTATTTTGTTATAAA   7823
Pachon11     ACTCCAAAAGGGTCATTCAGCAGAGCTGGACCTATTGAACAGTGTGTATTTTGTTATAAA   7822
Pachon7      ACTCCAAAAGGGTCATTCAGCAGAGCTGGACCTATTGAACAGTGTGTATTTTGTTATAAA   7823
Pachon3      ACTCCAAAAGGGTCATTCAGCAGAGCTGGACCTATTGAACAGTGTGTATTTTGTTATAAA   7823
Pachon8      ACTCCAAAAGGGTCATTCAGCAGAGCTGGACCTATTGAACAGTGTGTATTTTGTTATAAA   7823
Pachon15     ACTCCAAAAGGGTCATTCAGCAGAGCTGGACCTATTGAACAGTGTGTATTTTGTTATAAA   7823
             ************************************************************

Rascon4      ACCCATCAACAAGGCTGGACTTTGCTGAGCGAGGCCTGAATTTGAACCCTACTTGATTTT   7821
Surface      ACCCATCAACAAGGCTGGACTTTGCTGAGCGAGGCCTGAATTTGAACCCTACTTGATTTT   7894
Rascon8      ACCCATCAACAAGGCTGGACTTTGCTGAGCGAGGCCTGAATTTGAACCCTACTTGATTTT   7886
Rascon2      ACCCATCAACAAGGCTGGACTTTGCTGAGCGAGGCCTGAATTTGAACCCTACTTGATTTT   7888
Rascon15     ACCCATCAACAAGGCTGGACTTTGCTGAGCGAGGCCTGAATTTGAACCCTACTTGATTTT   7871
Rascon13     ACCCATCAACAAGGCTGGACTTTGCTGAGCGAGGCCTGAATTTGAACCCTACTTGATTTT   7883
Rascon6      ACCCATCAACAAGGCTGGACTTTGCTGAGCGAGGCCTGAATTTGAACCCTACTTGATTTT   7885
Pachon14     ACCCATCAACAAGGCTGGACTTTGCTGAGCGAGGCCTGAATTTGAACCCTACTTGATTTT   7876
Pachon9      ACCCATCAACAAGGCTGGACTTTGCTGAGCGAGGCCTGAATTTGAACCCTACTTGATTTT   7885
Pachon17     ACCCATCAACAAGGCTGGACTTTGCTGAGCGAGGCCTGAATTTGAACCCTACTTGATTTT   7883
Pachon12     ACCCATCAACAAGGCTGGACTTTGCTGAGCGAGGCCTGAATTTGAACCCTACTTGATTTT   7883
Pachon11     ACCCATCAACAAGGCTGGACTTTGCTGAGCGAGGCCTGAATTTGAACCCTACTTGATTTT   7882
Pachon7      ACCCATCAACAAGGCTGGACTTTGCTGAGCGAGGCCTGAATTTGAACCCTACTTGATTTT   7883
Pachon3      ACCCATCAACAAGGCTGGACTTTGCTGAGCGAGGCCTGAATTTGAACCCTACTTGATTTT   7883
Pachon8      ACCCATCAACAAGGCTGGACTTTGCTGAGCGAGGCCTGAATTTGAACCCTACTTGATTTT   7883
Pachon15     ACCCATCAACAAGGCTGGACTTTGCTGAGCGAGGCCTGAATTTGAACCCTACTTGATTTT   7883
             ************************************************************

Rascon4      TCTAATAGAATTTTGAACAACATTCATATTGAATAACCAAAATGAAGCAAAAAGGGACAT   7881
Surface      TCTTATAGAATTTTGAACAACAATCATATTGAATAACCAAAATGAAGCAAAAAGGGACAT   7954
Rascon8      TCTTATAGAATTTTGAACAACAATCATATTGAAGAACCAAAATGAAGCAAAAAGGGACAT   7946
Rascon2      TCTTATAGAATTTTGAACAACAATCATATTGAAGAACCAAAATGAAGCAAAAAGGGACAT   7948
Rascon15     TCTTATAGAATTTTGAACAACAATCATATTGAAGAACCAAAATGAAGCAAAAAGGGACAT   7931
```

```
Rascon13     TCTTATAGAATTTTGAACAACAATCATATTGAAGAACCAAAATGAAGCAAAAAGGGACAT     7943
Rascon6      TCTTATAGAATTTTGAACAACAATCATATTGAAGAACCAAAATGAAGCAAAAAGGGACAT     7945
Pachon14     TCTTATAGAATTTTGAACAACAATCATATTGAAGAACCAAAATGAAGCAAAAAGGGACAT     7936
Pachon9      TCTTATAGAATTTTGAACAACAATCATATTGAAGAACCAAAATGAAGCAAAAAGGGACAT     7945
Pachon17     TCTTATAGAATTTTGAACAACAATCATATTGAAGAACCAAAATGAAGCAAAAAGGGACAT     7943
Pachon12     TCTTATAGAATTTTGAACAACAATCATATTGAAGAACCAAAATGAAGCAAAAAGGGACAT     7943
Pachon11     TCTTATAGAATTTTGAACAACAATCATATTGAAGAACCAAAATGAAGCAAAAAGGGACAT     7942
Pachon7      TCTTATAGAATTTTGAACAACAATCATATTGAAGAACCAAAATGAAGCAAAAAGGGACAT     7943
Pachon3      TCTTATAGAATTTTGAACAACAATCATATTGAAGAACCAAAATGAAGCAAAAAGGGACAT     7943
Pachon8      TCTTATAGAATTTTGAACAACAATCATATTGAAGAACCAAAATGAAGCAAAAAGGGACAT     7943
Pachon15     TCTTATAGAATTTTGAACAACAATCATATTGAAGAACCAAAATGAAGCAAAAAGGGACAT     7943
             *** ****************** ********** ***************************

Rascon4      TCAGAGACTTTATTTGATATGCTTTGCTCTATTTTTGATAACAGATCGTATTTTATTCAC     7941 Rx3
Surface      TCAGAGACTTTATTTGATATGCTTTGCTCTATTTTTGATAACAGATCGTATTTTATTCAC     8014 3’-UTR
Rascon8      TCAGAGACTTTATTTGATATGCTTTGCTCTATTTTTGATAACAGATCGTATTTTATTCAC     8006
Rascon2      TCAGAGACTTTATTTGATATGCTTTGCTCTATTTTTGATAACAGATCGTATTTTATTCAC     8008
Rascon15     TCAGAGACTTTATTTGATATGCTTTGCTCTATTTTTGATAACAGATCGTATTTTATTCAC     7991
Rascon13     TCAGAGACTTTATTTGATATGCTTTGCTCTATTTTTGATAACAGATCGTATTTTATTCAC     8003
Rascon6      TCAGAGACTTTATTTGATATGCTTTGCTCTATTTTTGATAACAGATCGTATTTTATTCAC     8005
Pachon14     TCAGAGACTTTATTTGATATGCTTTGCTCTATTTTTGATAACAGATCGTATTTTATTCAC     7996
Pachon9      TCAGAGACTTTATTTGATATGCTTTGCTCTATTTTTGATAACAGATCGTATTTTATTCAC     8005
Pachon17     TCAGAGACTTTATTTGATATGCTTTGCTCTATTTTTGATAACAGATCGTATTTTATTCAC     8003
Pachon12     TCAGAGACTTTATTTGATATGCTTTGCTCTATTTTTGATAACAGATCGTATTTTATTCAC     8003
Pachon11     TCAGAGACTTTATTTGATATGCTTTGCTCTATTTTTGATAACAGATCGTATTTTATTCAC     8002
Pachon7      TCAGAGACTTTATTTGATATGCTTTGCTCTATTTTTGATAACAGATCGTATTTTATTCAC     8003
Pachon3      TCAGAGACTTTATTTGATATGCTTTGCTCTATTTTTGATAACAGATCGTATTTTATTCAC     8003
Pachon8      TCAGAGACTTTATTTGATATGCTTTGCTCTATTTTTGATAACAGATCGTATTTTATTCAC     8003
Pachon15     TCAGAGACTTTATTTGATATGCTTTGCTCTATTTTTGATAACAGATCGTATTTTATTCAC     8003
             ************************************************************

Rascon4      TAATAAAAGCAATGTAAGTATGCCGTCTACAATCACAAAATCAAAATACACAAAACAAAA     8001
Surface      TAATAAAAGCAATGTAAGTATGCCGTCTACAATCACAAAATCAAAATACACAAAACAAAA     8074
Rascon8      TAATAAAAGCAATGTAAGTATGCCGTCTACAATCACAAAATCAAAATACACAAAACAAAA     8066
Rascon2      TAATAAAAGCAATGTAAGTATGCCGTCTACAATCACAAAATCAAAATACACAAAACAAAA     8068
Rascon15     TAATAAAAGCAATGTAAGTATGCCGTCTACAATCACAAAATCAAAATACACAAAACAAAA     8051
Rascon13     TAATAAAAGCAATGTAAGTATGCCGTCTACAATCACAAAATCAAAATACACAAAACAAAA     8063
Rascon6      TAATAAAAGCAATGTAAGTATGCCGTCTACAATCACAAAATCAAAATACACAAAACAAAA     8065
Pachon14     TAATAAAAGCAATGTAAGTATGCCGTCTACAATCACAAAATCAAAATACACAAAACAAAA     8056
Pachon9      TAATAAAAGCAATGTAAGTATGCCGTCTACAATCACAAAATCAAAATACACAAAACAAAA     8065
Pachon17     TAATAAAAGCAATGTAAGTATGCCGTCTACAATCACAAAATCAAAATACACAAAACAAAA     8063
Pachon12     TAATAAAAGCAATGTAAGTATGCCGTCTACAATCACAAAATCAAAATACACAAAACAAAA     8063
Pachon11     TAATAAAAGCAATGTAAGTATGCCGTCTACAATCACAAAATCAAAATACACAAAACAAAA     8062
Pachon7      TAATAAAAGCAATGTAAGTATGCCGTCTACAATCACAAAATCAAAATACACAAAACAAAA     8063
Pachon3      TAATAAAAGCAATGTAAGTATGCCGTCTACAATCACAAAATCAAAATACACAAAACAAAA     8063
Pachon8      TAATAAAAGCAATGTAAGTATGCCGTCTACAATCACAAAATCAAAATACACAAAACAAAA     8063
Pachon15     TAATAAAAGCAATGTAAGTATGCCGTCTACAATCACAAAATCAAAATACACAAAACAAAA     8063
             ************************************************************

Rascon4      AAATTAAGTTGACAGTGTTACAGTTCCCCTTCTGAGAAGAATTCAAAAACTATATTTGAG     8061
Surface      AAATTAAGTTGACAGTGTTACAGTTCCCCTTCTGAGAAGAATTCAAAAACTATATTTGAG     8134
Rascon8      AAATTAAGTTGACAGTGTTACAGTTCCCCTTCTGAGAAGAATTCAAAAACTATATTTGAG     8126
Rascon2      AAATTAAGTTGACAGTGTTACAGTTCCCCTTCTGAGAAGAATTCAAAAACTATATTTGAG     8128
Rascon15     AAATTAAGTTGACAGTGTTACAGTTCCCCTTCTGAGAAGAATTCAAAAACTATATTTGAG     8111
Rascon13     AAATTAAGTTGACAGTGTTACAGTTCCCCTTCTGAGAAGAATTCAAAAACTATATTTGAG     8123
Rascon6      AAATTAAGTTGACAGTGTTACAGTTCCCCTTCTGAGAAGAATTCAAAAACTATATTTGAG     8125
Pachon14     AAATTAAGTTGACAGTGTTACAGTTCCCCTTCTGAGAAGAATTCAAAAACTATATTTGAG     8116
Pachon9      AAATTAAGTTGACAGTGTTACAGTTCCCCTTCTGAGAAGAATTCAAAAACTATATTTGAG     8125
Pachon17     AAATTAAGTTGACAGTGTTACAGTTCCCCTTCTGAGAAGAATTCAAAAACTATATTTGAG     8123
Pachon12     AAATTAAGTTGACAGTGTTACAGTTCCCCTTCTGAGAAGAATTCAAAAACTATATTTGAG     8123
Pachon11     AAATTAAGTTGACAGTGTTACAGTTCCCCTTCTGAGAAGAATTCAAAAACTATATTTGAG     8122
Pachon7      AAATTAAGTTGACAGTGTTACAGTTCCCCTTCTGAGAAGAATTCAAAAACTATATTTGAG     8123
Pachon3      AAATTAAGTTGACAGTGTTACAGTTCCCCTTCTGAGAAGAATTCAAAAACTATATTTGAG     8123
Pachon8      AAATTAAGTTGACAGTGTTACAGTTCCCCTTCTGAGAAGAATTCAAAAACTATATTTGAG     8123
Pachon15     AAATTAAGTTGACAGTGTTACAGTTCCCCTTCTGAGAAGAATTCAAAAACTATATTTGAG     8123
             ************************************************************

Rascon4      AAAAAATTGCTCTATTTTTATATTAATTTGGAAATCATGGAATCATGGAAGCATGGAAAT     8121
Surface      AAAAAATTGCTCTATTTTTATATTAATTTGGAAATCCTGGAATCATGGAAGCATGGAAAT     8194
Rascon8      AAAAAATTGCTCTATTTTTATATTAATTTGGAAATCCTGGAATCATGGAAGCATGGAAAT     8186
Rascon2      AAAAAATTGCTCTATTTTTATATTAATTTGGAAATCCTGGAATCATGGAAGCATGGAAAT     8188
Rascon15     AAAAAATTGCTCTATTTTTATATTAATTTGGAAATCCTGGAATCATGGAAGCATGGAAAT     8171
Rascon13     AAAAAATTGCTCTATTTTTATATTAATTTGGAAATCCTGGAATCATGGAAGCATGGAAAT     8183
Rascon6      AAAAAATTGCTCTATTTTTATATTAATTTGGAAATCCTGGAATCATGGAAGCATGGAAAT     8185
Pachon14     AAAAAATTGCTCTATTTTTATATTAATTTGGAAATCCTGGAATCATGGAAGCATGGAAAT     8176
Pachon9      AAAAAATTGCTCTATTTTTATATTAATTTGGAAATCCTGGAATCATGGAAGCATGGAAAT     8185
Pachon17     AAAAAATTGCTCTATTTTTATATTAATTTGGAAATCCTGGAATCATGGAAGCATGGAAAT     8183
```

```
Pachon12     AAAAAATTGCTCTATTTTTATATTAATTTGGAAATCCTGGAATCATGGAAGCATGGAAAT     8183
Pachon11     AAAAAATTGCTCTATTTTTATATTAATTTGGAAATCCTGGAATCATGGAAGCATGGAAAT     8182
Pachon7      AAAAAATTGCTCTATTTTTATATTAATTTGGAAATCCTGGAATCATGGAAGCATGGAAAT     8183
Pachon3      AAAAAATTGCTCTATTTTTATATTAATTTGGAAATCCTGGAATCATGGAAGCATGGAAAT     8183
Pachon8      AAAAAATTGCTCTATTTTTATATTAATTTGGAAATCCTGGAATCATGGAAGCATGGAAAT     8183
Pachon15     AAAAAATTGCTCTATTTTTATATTAATTTGGAAATCCTGGAATCATGGAAGCATGGAAAT     8183
             ************************************ ***********************

Rascon4      CTTAATGGCAATGGCTTTGAACTATCAAAACCAGCAGTAATTTTACTCCACTCTCTGAGA     8181 Rx3
Surface      CTTAATGGCAATGGCTTTGAACTATCAAAACCAGCAGTAATTTTACTCCACTCTCTGAGA     8254 3’-UTR
Rascon8      CTTAATGGCAATGGCTTTGAACTATCAAAACCAGCAGTAATTTTACTCCACTCTCTGAGA     8246
Rascon2      CTTAATGGCAATGGCTTTGAACTATCAAAACCAGCAGTAATTTTACTCCACTCTCTGAGA     8248
Rascon15     CTTAATGGCAATGGCTTTGAACTATCAAAACCAGCAGTAATTTTACTCCACTCTCTGAGA     8231
Rascon13     CTTAATGGCAATGGCTTTGAACTATCAAAACCAGCAGTAATTTTACTCCACTCTCTGAGA     8243
Rascon6      CTTAATGGCAATGGCTTTGAACTATCAAAACCAGCAGTAATTTTACTCCACTCTCTGAGA     8245
Pachon14     CTTAATGGCAATGGCTTTGAACTATCAAAACCAGCAGTAATTTTACTCCACTCTCTGAGA     8236
Pachon9      CTTAATGGCAATGGCTTTGAACTATCAAAACCAGCAGTAATTTTACTCCACTCTCTGAGA     8245
Pachon17     CTTAATGGCAATGGCTTTGAACTATCAAAACCAGCAGTAATTTTACTCCACTCTCTGAGA     8243
Pachon12     CTTAATGGCAATGGCTTTGAACTATCAAAACCAGCAGTAATTTTACTCCACTCTCTGAGA     8243
Pachon11     CTTAATGGCAATGGCTTTGAACTATCAAAACCAGCAGTAATTTTACTCCACTCTCTGAGA     8242
Pachon7      CTTAATGGCAATGGCTTTGAACTATCAAAACCAGCAGTAATTTTACTCCACTCTCTGAGA     8243
Pachon3      CTTAATGGCAATGGCTTTGAACTATCAAAACCAGCAGTAATTTTACTCCACTCTCTGAGA     8243
Pachon8      CTTAATGGCAATGGCTTTGAACTATCAAAACCAGCAGTAATTTTACTCCACTCTCTGAGA     8243
Pachon15     CTTAATGGCAATGGCTTTGAACTATCAAAACCAGCAGTAATTTTACTCCACTCTCTGAGA     8243
             ************************************************************

Rascon4      TGGTTCCTTACTTTCCCCTGACCCCTATGATGCCACTACAATCTATTTATTCTAATTTCA     8241
Surface      TGGTTCCTTACTTTCCCCTGACCCCTATGATGCCACTACAATCTATTTATTCTAATTTCA     8314
Rascon8      TGGTTCCTTACTTTCCCCTGACCCCTATGATGCCACTACAATCTATTTATTCTAATTTCA     8306
Rascon2      TGGTTCCTTACTTTCCCCTGACCCCTATGATGCCACTACAATCTATTTATTCTAATTTCA     8308
Rascon15     TGGTTCCTTACTTTCCCCTGACCCCTATGATGCCACTACAATCTATTTATTCTAATTTCA     8291
Rascon13     TGGTTCCTTACTTTCCCCTGACCCCTATGATGCCACTACAATCTATTTATTCTAATTTCA     8303
Rascon6      TGGTTCCTTACTTTCCCCTGACCCCTATGATGCCACTACAATCTATTTATTCTAATTTCA     8305
Pachon14     TGGTTCCTTACTTTCCCCTGACCCCTATGATGCCACTACAATCTATTTATTCTAATTTCA     8296
Pachon9      TGGTTCCTTACTTTCCCCTGACCCCTATGATGCCACTACAATCTATTTATTCTAATTTCA     8305
Pachon17     TGGTTCCTTACTTTCCCCTGACCCCTATGATGCCACTACAATCTATTTATTCTAATTTCA     8303
Pachon12     TGGTTCCTTACTTTCCCCTGACCCCTATGATGCCACTACAATCTATTTATTCTAATTTCA     8303
Pachon11     TGGTTCCTTACTTTCCCCTGACCCCTATGATGCCACTACAATCTATTTATTCTAATTTCA     8302
Pachon7      TGGTTCCTTACTTTCCCCTGACCCCTATGATGCCACTACAATCTATTTATTCTAATTTCA     8303
Pachon3      TGGTTCCTTACTTTCCCCTGACCCCTATGATGCCACTACAATCTATTTATTCTAATTTCA     8303
Pachon8      TGGTTCCTTACTTTCCCCTGACCCCTATGATGCCACTACAATCTATTTATTCTAATTTCA     8303
Pachon15     TGGTTCCTTACTTTCCCCTGACCCCTATGATGCCACTACAATCTATTTATTCTAATTTCA     8303
             ************************************************************

Rascon4      TCTTCCCTGTATAACAGATTTGTTTCCAGATTTTTTAAATACACTTTTATGTGTGTGTAC     8301
Surface      TCTTCCCTGTATAACAGATTTGTTTCCAGATTTTTTTAATACACTTTTATGTGTGTGTAC     8374
Rascon8      TCTTCCCTGTATAACAGATTTGTTTCCAGATTTTTTAAATACACTTTTATGTGTGTGTAC     8366
Rascon2      TCTTCCCTGTATAACAGATTTGTTTCCAGATTTTTTTAATACACTTTTATGTGTGTGTAC     8368
Rascon15     TCTTCCCTGTATAACAGATTTGTTTCCAGATTTTTTTAATACACTTTTATGTGTGTGTAC     8351
Rascon13     TCTTCCCTGTATAACAGATTTGTTTCCAGATTTTTTTAATACACTTTTATGTGTGTGTAC     8363
Rascon6      TCTTCCCTGTATAACGTATTTGTTTCCAGATTTTTTTAATACACTTTTATGTGTGTGTAC     8365
Pachon14     TCTTCCCTGTATAACAGATTTGTTTCCAGATTTTTTTAATACACTTTTATGTGTGTGTAC     8356
Pachon9      TCTTCCCTGTATAACAGATTTGTTTCCAGATTTTTTTAATACACTTTTATGTGTGTGTAC     8365
Pachon17     TCTTCCCTGTATAACAGATTTGTTTCCAGATTTTTTTAATACACTTTTATGTGTGTGTAC     8363
Pachon12     TCTTCCCTGTATAACAGATTTGTTTCCAGATTTTTTTAATACACTTTTATGTGTGTGTAC     8363
Pachon11     TCTTCCCTGTATAACAGATTTGTTTCCAGATTTTTTTAATACACTTTTATGTGTGTGTAC     8362
Pachon7      TCTTCCCTGTATAACAGATTTGTTTCCAGATTTTTTTAATACACTTTTATGTGTGTGTAC     8363
Pachon3      TCTTCCCTGTATAACAGATTTGTTTCCAGATTTTTTTAATACACTTTTATGTGTGTGTAC     8363
Pachon8      TCTTCCCTGTATAACAGATTTGTTTCCAGATTTTTTTAATACACTTTTATGTGTGTGTAC     8363
Pachon15     TCTTCCCTGTATAACAGATTTGTTTCCAGATTTTTTTAATACACTTTTATGTGTGTGTAC     8363
             ***************  ******************* ***********************

Rascon4      ACTCAGCCAGCTGAACTGTGTTATTTTGCAATAATGTGAACAATTCTGCAATAAAATGAC     8361
Surface      ACTCAGCCAGCTGAACTGTGTTATTTTGCAATAATGTGAACAATTCTGCAATAAAATTAC     8434
Rascon8      ACTCAGCCAGCTGAACTGTGTTATTTTGCAATAATGTGAACAATTCTGCAATAAAATGAC     8426
Rascon2      ACTCAGCCAGCTGAACTGTGTTATTTTGCAATAATGTGAACAATTCTGCAATAAAATGAC     8428
Rascon15     ACTCAGCCAGCTGAACTGTGTTATTTTGCAATAATGTGAACAATTCTGCAATAAAATGAC     8411
Rascon13     ACTCAGCCAGCTGAACTGTGTTATTTTGCAATAATGTGAACAATTCTGCAATAAAATGAC     8423
Rascon6      ACTCAGCCAGCTGAACTGTGTTATTTTGCAATAATGTGAACAATTCTGCAATAAAATGAC     8425
Pachon14     ACTCAGCCAGCTGAACTGTGTTATTTTGCAATAATGTGAACAATTCTGCAATAAAATGAC     8416
Pachon9      ACTCAGCCAGCTGAACTGTGTTATTTTGCAATAATGTGAACAATTCTGCAATAAAATGAC     8425
Pachon17     ACTCAGCCAGCTGAACTGTGTTATTTTGCAATAATGTGAACAATTCTGCAATAAAATGAC     8423
Pachon12     ACTCAGCCAGCTGAACTGTGTTATTTTGCAATAATGTGAACAATTCTGCAATAAAATGAC     8423
Pachon11     ACTCAGCCAGCTGAACTGTGTTATTTTGCAATAATGTGAACAATTCTGCAATAAAATGAC     8422
Pachon7      ACTCAGCCAGCTGAACTGTGTTATTTTGCAATAATGTGAACAATTCTGCAATAAAATGAC     8423
Pachon3      ACTCAGCCAGCTGAACTGTGTTATTTTGCAATAATGTGAACAATTCTGCAATAAAATGAC     8423
Pachon8      ACTCAGCCAGCTGAACTGTGTTATTTTGCAATAATGTGAACAATTCTGCAATAAAATGAC     8423
```

```
Pachon15      ACTCAGCCAGCTGAACTGTGTTATTTTGCAATAATGTGAACAATTCTGCAATAAAATGAC      8423
              ********************************************************* **

Rascon4       CATTTAAAAAATTCCCATGAACATGTGGTCATTTTTTTATGCATCACTGTGGGATGATCT      8421 Rx3
Surface       CATTTTAAAAATTCCCATGAACATGTGGTCATTTTTTTATGTATCACTGTGGGATGATCT      8494 3'-UTR
Rascon8       CATTTAAAAAATTCCCATGAACATGTGGTCATTTTTTTATGCATCACTGTGGGATGATCT      8486 end
Rascon2       CATTTAAAAAATTCCCATGAACATGTGCTCATTTTTTTATGCATCACTGTGGGACGATCT      8488
Rascon15      CATTTAAAAAATTCCCATGAACATGTGGTCATTTTTTTATGCATCACTGTGGGATGATCT      8471
Rascon13      CATTTAAAAAATTCCCAAGAACATGTGGTCATTTTTTTATGCATCACTGTGGGATGATCT      8483
Rascon6       CATTTAAAAAATTCCCATGAACATGTGGTCATTTTTTTATGCATCACTGTGGGATGATCT      8485
Pachon14      CATTTAAAAAATTCCCATGAACATGTGGTCATTTTTTTATGCATCACTGTGGGATGATCT      8476
Pachon9       CATTTAAAAAATTCCCATGAACATGTGGTCATTTTTTTATGCATCACTGTGGGATGATCT      8485
Pachon17      CATTTAAAAAATTCCCATGAACATGTGGTCATTTTTTTATGCATCACTGTGGGATGATCT      8483
Pachon12      CATTTAAAAAATTCCCATGAACATGTGGTCATTTTTTTATGCATCACTGTGGGATGATCT      8483
Pachon11      CATTTAAAAAATTCCCATGAACATGTGGTCATTTTTTTATGCATCACTGTGGGATGATCT      8482
Pachon7       CATTTAAAAAATTCCCATGAACATGTGGTCATTTTTTTATGCATCACTGTGGGATGATCT      8483
Pachon3       CATTTAAAAAATTCCCATGAACATGTGGTCATTTTTTTATGCATCACTGTGGGATGATCT      8483
Pachon8       CATTTAAAAAATTCCCATGAACATGTGGTCATTTTTTTATGCATCACTGTGGGATGATCT      8483
Pachon15      CATTTAAAAAATTCCCATGAACATGTGGTCATTTTTTTATGCATCACTGTGGGATGATCT      8483
              ***** *********** ********* ************* ************ *****

Rascon4       AATAAGCCAGTGCGTTTATTTGGCTGTTTATACCTACTTTCACACAAGAATCACACATCT      8481
Surface       AATAAGCCAGTGCGTTTATTTGGCTGTTTATACCTACTTTCACACAAGAATCACACATCT      8554
Rascon8       AATAAGCCAGTGCGTTTATTTGGCTGTTTATACCTACTTTCACACAAGAATCACACATCT      8546
Rascon2       AATAAGCCAGTGCGTTTATTTGGCTGTTTATACCTACTTTCACACAAGAATCACACATCT      8548
Rascon15      AATAAGCCAGTGCATTTATTTGGCTGTTTATACCTACTTTCACACAAGAATCACACATCT      8531
Rascon13      AATAAGCCAGTGCGTTTATTTGGCTGTTTATACCTACTTTCACACAAGAATCACACATCT      8543
Rascon6       AATAAGCCAGTGCATTTATTTGGCTGTTTATACCTACTTTCACACAAGAATCACACATCT      8545
Pachon14      AATAAGCCAGTGCGTTTATTTGGCTGTTTATACCTACTTTCACACAAGAATCACACATCT      8536
Pachon9       AATAAGCCAGTGCGTTTATTTGGCTGTTTATACCTACTTTCACACAAGAATCACACATCT      8545
Pachon17      AATAAGCCAGTGCGTTTATTTGGCTGTTTATACCTACTTTCACACAAGAATCACACATCT      8543
Pachon12      AATAAGCCAGTGCGTTTATTTGGCTGTTTATACCTACTTTCACACAAGAATCACACATCT      8543
Pachon11      AATAAGCCAGTGCGTTTATTTGGCTGTTTATACCTACTTTCACACAAGAATCACACATCT      8542
Pachon7       AATAAGCCAGTGCGTTTATTTGGCTGTTTATACCTACTTTCACACAAGAATCACACATCT      8543
Pachon3       AATAAGCCAGTGCGTTTATTTGGCTGTTTATACCTACTTTCACACAAGAATCACACATCT      8543
Pachon8       AATAAGCCAGTGCGTTTATTTGGCTGTTTATACCTACTTTCACACAAGAATCACACATCT      8543
Pachon15      AATAAGCCAGTGCGTTTATTTGGCTGTTTATACCTACTTTCACACAAGAATCACACATCT      8543
              ************* **********************************************

Rascon4       GTAATGCCACTATTAAAGTATTTAAGTATGGAGAATTAAGCATAAGAACAAGAACAATAA      8541
Surface       GTAATGCCACTATTAAAGTATTTAAGTATGGAGAATTAAGCATAAGAACAAGAACAATAA      8614
Rascon8       GTAATGCCACTATTAAAGTATTTAAGTATGGAGAATTAAGCATAAGAACAAGAACAATAA      8606
Rascon2       GTAATGCCACTATTAAAGTATTTAAGTATGGAGAATTAAGCATAAGAACAAGAACAATAA      8608
Rascon15      GTAATGCCACTATTAAAGTATTTAAGTATGGAGAATTAAGCATAAGAACCAGAACAATAA      8591
Rascon13      GTAATGCCACTATTAAAGTATTTAAGTATGGAGAATTAAGCATAAGAACCAGAACAATAA      8603
Rascon6       GTAATGCCACTATTAAAGTATTTAAGTATGGAGAATTAAGCATAAGAACCAGAACAATAA      8605
Pachon14      GTAATGCCACTATTAAAGTATTTAAGTATGGAGAATTAAGCATAAGAACCAGAACAATAA      8596
Pachon9       GTAATGCCACTATTAAAGTATTTAAGTATGGAGAATTAAGCATAAGAACCAGAACAATAA      8605
Pachon17      GTAATGCCACTATTAAAGTATTTAAGTATGGAGAATTAAGCATAAGAACCAGAACAATAA      8603
Pachon12      GTAATGCCACTATTAAAGTATTTAAGTATGGAGAATTAAGCATAAGAACCAGAACAATAA      8603
Pachon11      GTAATGCCACTATTAAAGTATTTAAGTATGGAGAATTAAGCATAAGAACCAGAACAATAA      8602
Pachon7       GTAATGCCACTATTAAAGTATTTAAGTATGGAGAATTAAGCATAAGAACCAGAACAATAA      8603
Pachon3       GTAATGCCACTATTAAAGTATTTAAGTATGGAGAATTAAGCATAAGAACCAGAACAATAA      8603
Pachon8       GTAATGCCACTATTAAAGTATTTAAGTATGGAGAATTAAGCATAAGAACCAGAACAATAA      8603
Pachon15      GTAATGCCACTATTAAAGTATTTAAGTATGGAGAATTAAGCATAAGAACCAGAACAATAA      8603
              ************************************************* **********

Rascon4       TGTACAGTATGTTGTGTGGATTTCTAATAACACTAAATTAAGAAAGCATGGATTAAATAC      8601
Surface       TGTACAGTATGTTGTGTGGATTTCTAATAACACTAAATTAAGAAAGCATGGATTAAATAC      8674
Rascon8       TGTACAGTATGTTGTGTGGATTTCTAATAACACTAAATTAAGAAAGCATGGATTAAATAC      8666
Rascon2       TGTACAGTATGTTGTGTGGATTTCTAATAACACTAAATTAAGAAAGCATGGATTAAATAC      8668
Rascon15      TGTACAGTATGTTGTGTGGATTTCTAATAACACTAAATTAAGAAAGCATGGATTAAATAC      8651
Rascon13      TGTACAGTATGTTGTGTGGATTTCTAATAACACTAAATTAAGAAAGCATGGATTAAATAC      8663
Rascon6       TGTACAGTATGTTGTGTGGATTTCTAATAACACTAAATTAAGAAAGCATGGATTAAATAC      8665
Pachon14      TGTACAGTATGTTGTGTGGATTTCTAATAACACTAAATTAAGAAAGCATGGATTAAATAC      8656
Pachon9       TGTACAGTATGTTGTGTGGATTTCTAATAACACTAAATTAAGAAAGCATGGATTAAATAC      8665
Pachon17      TGTACAGTATGTTGTGTGGATTTCTAATAACACTAAATTAAGAAAGCATGGATTAAATAC      8663
Pachon12      TGTACAGTATGTTGTGTGGATTTCTAATAACACTAAATTAAGAAAGCATGGATTAAATAC      8663
Pachon11      TGTACAGTATGTTGTGTGGATTTCTAATAACACTAAATTAAGAAAGCATGGATTAAATAC      8662
Pachon7       TGTACAGTATGTTGTGTGGATTTCTAATAACACTAAATTAAGAAAGCATGGATTAAATAC      8663
Pachon3       TGTACAGTATGTTGTGTGGATTTCTAATAACACTAAATTAAGAAAGCATGGATTAAATAC      8663
Pachon8       TGTACAGTATGTTGTGTGGATTTCTAATAACACTAAATTAAGAAAGCATGGATTAAATAC      8663
Pachon15      TGTACAGTATGTTGTGTGGATTTCTAATAACACTAAATTAAGAAAGCATGGATTAAATAC      8663
              ************************************************************

Rascon4       ATTAGTTTGGATTACTAAGATCATAGCTCATTGTGAAAATTAAGAGATGAATAATTATAA      8661 Not in
Surface       ATTAGTTTGGATTACTAAGATCATAACTCATTGTGAAAATTAAGAGATGAATAATTATAA      8734 Choy SF
```

```
Rascon8         ATTAGTTTGGATTACTAAGATCATAGCTCATTGTGAAAATTAAGAGATGAATAATTATAA        8726 Mix C/A
Rascon2         ATTAGTTTGGATTACTAAGATCATAGCTCATTGTGAAAATTAAGAGATGAATAATTATAA        8728
Rascon15        ATTAGTTTGGATTACTAAGATCATAGCTCATTGTGAAAATTAAGAGATGAATAATTATAA        8711
Rascon13        ATTAGTTTGGATTACTAAGATCATAGCTCATTGTGAAAATTAAGAGATGAATAATTATAA        8723
Rascon6         ATTAGTTTGGATTACTAAGATCATAGCTCATTGTGAAAATTAAGAGATGAATAATTATAA        8725
Pachon14        ATTAGTTTGGATTACTAAGATCATAGCTCATTGTGAAAATTAAGAGATGAATCATTATAA        8716
Pachon9         ATTAGTTTGGATTACTAAGATCATAGCTCATTGTGAAAATTAAGAGATGAATCATTATAA        8725
Pachon17        ATTAGTTTGGATTACTAAGATCATAGCTCATTGTGAAAATTAAGAGATGAATCATTATAA        8723
Pachon12        ATTAGTTTGGATTACTAAGATCATAGCTCATTGTGAAAATTAAGAGATGAATCATTATAA        8723
Pachon11        ATTAGTTTGGATTACTAAGATCATAGCTCATTGTGAAAATTAAGAGATGAATCATTATAA        8722
Pachon7         ATTAGTTTGGATTACTAAGATCATAGCTCATTGTGAAAATTAAGAGATGAATCATTATAA        8723
Pachon3         ATTAGTTTGGATTACTAAGATCATAGCTCATTGTGAAAATTAAGAGATGAATCATTATAA        8723
Pachon8         ATTAGTTTGGATTACTAAGATCATAGCTCATTGTGAAAATTAAGAGATGAATCATTATAA        8723
Pachon15        ATTAGTTTGGATTACTAAGATCATAGCTCATTGTGAAAATTAAGAGATGAATCATTATAA        8723
                ************************* ************************** *******

Rascon4         ATGTATAAATCACATGTTGGACATTTATTAATTGTACCTTTGCTTCCCTCTAGTGGTCAC        8721
Surface         ATGTATAAATCACATGTTGGACATTTATTAATTGTACCTTTGCTTCCCTCTAGTGGTCAC        8794
Rascon8         ATGTATAAATCACATGTTGGACATTTATTAATTGTACCTTTGCTTCCCTCTAGTGGTCAC        8786
Rascon2         ATGTATAAATCACATGTTGGACATTTATTAATTGTACCTTTGCTTCCCTCTAGTGGTCAC        8788
Rascon15        ATGTATAAATCACATGTTGGACATTTATTAATTGTACCTTTGCTTCCCTCTAGTGGTCAC        8771
Rascon13        ATGTATAAATCACATGTTGGACATTTATTAATTGTACCTTTGCTTCCCTCTAATGGTCAC        8783
Rascon6         ATGTATAAATCACATGTTGGACATTTATTAATTGTACCTTTGCTTCCCTCTAATGGTCAC        8785
Pachon14        ATGTATAAATCACATGTTGGACATTTATTAATTGTACCTTTGCTTCCCTCTAGTGGTCAC        8776
Pachon9         ATGTATAAATCACATGTTGGACATTTATTAATTGTACCTTTGCTTCCCTCTAGTGGTCAC        8785
Pachon17        ATGTATAAATCACATGTTGGACATTTATTAATTGTACCTTTGCTTCCCTCTAGTGGTCAC        8783
Pachon12        ATGTATAAATCACATGTTGGACATTTATTAATTGTACCTTTGCTTCCCTCTAGTGGTCAC        8783
Pachon11        ATGTATAAATCACATGTTGGACATTTATTAATTGTACCTTTGCTTCCCTCTAGTGGTCAC        8782
Pachon7         ATGTATAAATCACATGTTGGACATTTATTAATTGTACCTTTGCTTCCCTCTAGTGGTCAC        8783
Pachon3         ATGTATAAATCACATGTTGGACATTTATTAATTGTACCTTTGCTTCCCTCTAGTGGTCAC        8783
Pachon8         ATGTATAAATCACATGTTGGACATTTATTAATTGTACCTTTGCTTCCCTCTAGTGGTCAC        8783
Pachon15        ATGTATAAATCACATGTTGGACATTTATTAATTGTACCTTTGCTTCCCTCTAGTGGTCAC        8783
                **************************************************** *******

Rascon4         TACAACAAAATGCACTATTGGCAAAATATTTACTTACAAAAAGCAGCACAATCACCTTCT        8781
Surface         TACAACAAAATGCACTATTGGCAAAATATTTACTTACAAAAAGCAGCACAATCACCTTCT        8854
Rascon8         TACAACAAAATGCACTATTGGCAAAATATTTACTTACAAAAAGCAGCACAATCACCTTCT        8846
Rascon2         TACAACAAAATGCACTATTGGCAAAATATTTACTTACAAAAAGCAGCACAATCACCTTCT        8848
Rascon15        TACAACAAAATGCACTATTGGCAAAATATTTACTTACAAAAAGCAGCACAATCACCTTCT        8831
Rascon13        TACAACAAAATGCACTATTGGCAAAATATTTACTTACAAAAAGCAGCACAATCACCTTCT        8843
Rascon6         TACAACAAAATGCACTATTGGCAAAATATTTACTTACAAAAAGCAGCACAATCACCTTCT        8845
Pachon14        TACAACAAAATGCACTATTGGCAAAATATTTACTTACAAAAAGCAGCACAATCACCTTCT        8836
Pachon9         TACAACAAAATGCACTATTGGCAAAATATTTACTTACAAAAAGCAGCACAATCACCTTCT        8845
Pachon17        TACAACAAAATGCACTATTGGCAAAATATTTACTTACAAAAAGCAGCACAATCACCTTCT        8843
Pachon12        TACAACAAAATGCACTATTGGCAAAATATTTACTTACAAAAAGCAGCACAATCACCTTCT        8843
Pachon11        TACAACAAAATGCACTATTGGCAAAATATTTACTTACAAAAAGCAGCACAATCACCTTCT        8842
Pachon7         TACAACAAAATGCACTATTGGCAAAATATTTACTTACAAAAAGCAGCACAATCACCTTCT        8843
Pachon3         TACAACAAAATGCACTATTGGCAAAATATTTACTTACAAAAAGCAGCACAATCACCTTCT        8843
Pachon8         TACAACAAAATGCACTATTGGCAAAATATTTACTTACAAAAAGCAGCACAATCACCTTCT        8843
Pachon15        TACAACAAAATGCACTATTGGCAAAATATTTACTTACAAAAAGCAGCACAATCACCTTCT        8843
                ************************************************************

Rascon4         ACAGTATTCAGTTTCCCTCAGTTTACAGGTATCATTTACTTATTTGTAGTAATAGTAGTA        8841
Surface         ACAGTATTCAGTTTCCCTCAGTTTACAGGTATCATTTACTTATTTGTAGTAATAGTAGTA        8914
Rascon8         ACAGTATTCAGTTTCCCTCAGTTTACAGGTATCATTTACTTATTTGTAGTAATAGTAGTA        8906
Rascon2         ACAGTATTCAGTTTCCCTCAGTTTACAGGTATCATTTACTTATTTGTAGTAATAGTAGTA        8908
Rascon15        ACAGTATTCAGTTTCCCTCAGTTTACAGGTATCATTTACTTATTTGTAGTAATAGTAGTA        8891
Rascon13        ACGGTATTCAGTTTCCCTCAGTTTACAGGTATCATTTACTTATTTGTAGTAATAGTAGTA        8903
Rascon6         ACAGTATTCAGTTTCCCTCAGTTTACAGGTATCATTTACTTATTTGTAGTAATAGTAGTA        8905
Pachon14        ACAGTATTCAGTTTCCCTCAGTTTACAGGTATCATTTACTTATTTGTAGTAATAGTAGTA        8896
Pachon9         ACAGTATTCAGTTTCCCTCAGTTTACAGGTATCATTTACTTATTTGTAGTAATAGTAGTA        8905
Pachon17        ACAGTATTCAGTTTCCCTCAGTTTACAGGTATCATTTACTTATTTGTAGTAATAGTAGTA        8903
Pachon12        ACAGTATTCAGTTTCCCTCAGTTTACAGGTATCATTTACTTATTTGTAGTAATAGTAGTA        8903
Pachon11        ACAGTATTCAGTTTCCCTCAGTTTACAGGTATCATTTACTTATTTGTAGTAATAGTAGTA        8902
Pachon7         ACAGTATTCAGTTTCCCTCAGTTTACAGGTATCATTTACTTATTTGTAGTAATAGTAGTA        8903
Pachon3         ACAGTATTCAGTTTCCCTCAGTTTACAGGTATCATTTACTTATTTGTAGTAATAGTAGTA        8903
Pachon8         ACAGTATTCAGTTTCCCTCAGTTTACAGGTATCATTTACTTATTTGTAGTAATAGTAGTA        8903
Pachon15        ACAGTATTCAGTTTCCCTCAGTTTACAGGTATCATTTACTTATTTGTAGTAATAGTAGTA        8903
                ** *********************************************************

Rascon4         ACATTAACAGTAGTAACAGTAGTAGTAGTAATAGTAGTAGTATCAATAGTAGTACACAAA        8901
Surface         ACATTAACAGTAGTAACAGTAGTAGTAGTAATAGTAGTAGTATCAATAGTAGTACACAAA        8974
Rascon8         ACATTAACAGTAGTAACAGTAGTAGTAGTAATAGTAGTAGTATCAATAGTAGTACACAAA        8966
Rascon2         ACATTAACAGTAGTAACAGTAGTAGTAGTAATAGTAGTAGTATCAATAGTAGTACACAAA        8968
Rascon15        ACATTAACAGTAGTAACAGTAGTAGTAGTAATAGTAGTAGTACCAATAGTAGTACACAAA        8951
Rascon13        ACATTAACAGTAGTAACAGTAGTAGTAGTAATAGTAGTAGTATCAATAGTAGTACACAAA        8963
Rascon6         ACATTAACAGTAGTAACAGTAGTAGTAGTAATAGTAGTAGTATCAATAGTAGTACACAAA        8965
```

```
Pachon14     ACATTAACAGTAGTAACAGTAGTAGTAGTAATAGTAGTAGTATCAATAGTAGTACACAAA     8956
Pachon9      ACATTAACAGTAGTAACAGTAGTAGTAGTAATAGTAGTAGTATCAATAGTAGTACACAAA     8965
Pachon17     ACATTAACAGTAGTAACAGTAGTAGTAGTAATAGTAGTAGTATCAATAGTAGTACACAAA     8963
Pachon12     ACATTAACAGTAGTAACAGTAGTAGTAGTAATAGTAGTAGTATCAATAGTAGTACACAAA     8963
Pachon11     ACATTAACAGTAGTAACAGTAGTAGTAGTAATAGTAGTAGTATCAATAGTAGTACACAAA     8962
Pachon7      ACATTAACAGTAGTAACAGTAGTAGTAGTAATAGTAGTAGTATCAATAGTAGTACACAAA     8963
Pachon3      ACATTAACAGTAGTAACAGTAGTAGTAGTAATAGTAGTAGTATCAATAGTAGTACACAAA     8963
Pachon8      ACATTAACAGTAGTAACAGTAGTAGTAGTAATAGTAGTAGTATCAATAGTAGTACACAAA     8963
Pachon15     ACATTAACAGTAGTAACAGTAGTAGTAGTAATAGTAGTAGTATCAATAGTAGTACACAAA     8963
             ****************************************** *****************

Rascon4      GCATCAGATGTACTATCAATAATAGTAGAAGTATCAATAGTAGTAGTAGCATCAGTAACA     8961
Surface      GCATCAGATGTACTATCAATAATAGTAGAAGTATCAATAGTAGTAGTAGCATCAGTAACA     9034
Rascon8      GCATCAGATGTACTATCAATAATAGTAGAAGTATCAATAGTAGTAGTAGCATCAGTAACA     9026
Rascon2      GCATCAGATGTACTATCAATAATAGTAGAAGTATCAATAGTAGTAGTAGCATCAGTAACA     9028
Rascon15     GCATCAGATGTACTATCAATAATAGTAGAAGTATCAATAGTAGTAGTAGCATCAGTAACA     9011
Rascon13     GCATCAGATGTACTATCAATAATAGTAGAAGTATCAATAGTAGTAGTAGCATCAGTAACA     9023
Rascon6      GCATCAGATGTACTATCAATAATAGTAGAAGTATCAATAGTAGTAGTAGCATCAGTAACA     9025
Pachon14     GCATCAGATGTACTATCAATAATAGTAGAAGTATCAATAGTAGTAGTAGCATCAGTAACA     9016
Pachon9      GCATCAGATGTACTATCAATAATAGTAGAAGTATCAATAGTAGTAGTAGCATCAGTAACA     9025
Pachon17     GCATCAGATGTACTATCAATAATAGTAGAAGTATCAATAGTAGTAGTAGCATCAGTAACA     9023
Pachon12     GCATCAGATGTACTATCAATAATAGTAGAAGTATCAATAGTAGTAGTAGCATCAGTAACA     9023
Pachon11     GCATCAGATGTACTATCAATAATAGTAGAAGTATCAATAGTAGTAGTAGCATCAGTAACA     9022
Pachon7      GCATCAGATGTACTATCAATAATAGTAGAAGTATCAATAGTAGTAGTAGCATCAGTAACA     9023
Pachon3      GCATCAGATGTACTATCAATAATAGTAGAAGTATCAATAGTAGTAGTAGCATCAGTAACA     9023
Pachon8      GCATCAGATGTACTATCAATAATAGTAGAAGTATCAATAGTAGTAGTAGCATCAGTAACA     9023
Pachon15     GCATCAGATGTACTATCAATAATAGTAGAAGTATCAATAGTAGTAGTAGCATCAGTAACA     9023
             ************************************************************

Rascon4      GTAATAGCAGTCTCATTAATAGAATAAGTAGAAGTATTAATAGTACTATCAGTAGTAGTA     9021
Surface      GTAGTAGCAGTCTCATTAATAGAATAAGTAGAAGTATTAATAGTACTATCAGTAGTAGTA     9094
Rascon8      GTAATAGCAGTCTCATTAATAGAATAAGTAGAAGTATTAATAGTACTATCAGTAGTAGTA     9086
Rascon2      GTAATAGCAGTCTCATTAATAGAATAAGTAGAAGTATTAATAGTACTATCAGTAGTAGTA     9088
Rascon15     GTAATAGCAGTCTCATTAATAGAATAAGTAGAAGTATTAATAGTACTATCAGTAGTAGTA     9071
Rascon13     GTAATAGCAGTCTCATTAATAGAATAAGTAGAAGTATTAATAGTACTATCAGTAGTAGTA     9083
Rascon6      GTAATAGCAGTCTCATTAATAGAATAAGTAGAAGTATTAATAGTACTATCAGTAGTAGTA     9085
Pachon14     GTAATAGCAGTCTCATTAATAGAATAAGTAGAAGTATTAATAGTACTATCAGTAGTAGTA     9076
Pachon9      GTAATAGCAGTCTCATTAATAGAATAAGTAGAAGTATTAATAGTACTATCAGTAGTAGTA     9085
Pachon17     GTAATAGCAGTCTCATTAATAGAATAAGTAGAAGTATTAATAGTACTATCAGTAGTAGTA     9083
Pachon12     GTAATAGCAGTCTCATTAATAGAATAAGTAGAAGTATTAATAGTACTATCAGTAGTAGTA     9083
Pachon11     GTAATAGCAGTCTCATTAATAGAATAAGTAGAAGTATTAATAGTACTATCAGTAGTAGTA     9082
Pachon7      GTAATAGCAGTCTCATTAATAGAATAAGTAGAAGTATTAATAGTACTATCAGTAGTAGTA     9083
Pachon3      GTAATAGCAGTCTCATTAATAGAATAAGTAGAAGTATTAATAGTACTATCAGTAGTAGTA     9083
Pachon8      GTAATAGCAGTCTCATTAATAGAATAAGTAGAAGTATTAATAGTACTATCAGTAGTAGTA     9083
Pachon15     GTAATAGCAGTCTCATTAATAGAATAAGTAGAAGTATTAATAGTACTATCAGTAGTAGTA     9083
             *** ********************************************************

Rascon4      GTTGTATGAGTAGAAGCATCACTAGTAGTAGTAGTAGTATCAGTGGTAGTAGTAGTAGTA     9081
Surface      GTTGTATGAGTAGAAGCATCACTAGTAGTAGTAGTAGT---A---GTAGTAGTAGTAGTA     9148
Rascon8      GTTGTATGAGTAGAAGCATCACTAGTAGTAGTAGTAGTATCA---GTAGTAGTAGTAGTA     9143
Rascon2      GTTGTATGAGTAGAAGCATCACTAGTAGTAGTAGTAGT---A---GTAGTAGTAGTAGTA     9142
Rascon15     GTTGTATGAGTAGAAGCATCACTAGTAGTAGTAGTAGT---A---GTAGTAGTAGTAGTA     9125
Rascon13     GTTGTATGAGTAGAAGCATCACTAGTAGTAGTAGTAGT---A---GTAGTAGTAGTAGTA     9137
Rascon6      GTTGTATGAGTAGAAGCATCACTAGTAGTAGTAGTAGT---A---GTAGTAGTAGTAGTA     9139
Pachon14     GTTGTATGAGTAGAAGCATCACTAGTAGTAGTAGTAGT---A---GTAGTAGTAGTAGTA     9130
Pachon9      GTTGTATGAGTAGAAGCATCACTAGTAGTAGTAGTAGT---A---GTAGTAGTAGTAGTA     9139
Pachon17     GTTGTATGAGTAGAAGCATCACTAGTAGTAGTAGTAGT---A---GTAGTAGTAGTAGTA     9137
Pachon12     GTTGTATGAGTAGAAGCATCAC------------------------TAGTAGTAGTAGTA     9119
Pachon11     GTTGTATGAGTAGAAGCATCACTAGTAGTAGTAGTAGT---A---GTAGTAGTAGTAGTA     9136
Pachon7      GTTGTATGAGTAGAAGCATCACTAGTAGTAGTAGTAGT---A---GTAGTAGTAGTAGTA     9137
Pachon3      GTTGTATGAGTAGAAGCATCACTAGTAGTAGTAGTAGT---A---GTAGTAGTAGTAGTA     9137
Pachon8      GTTGTATGAGTAGAAGCATCACTAGTAGTAGTAGTAGT---A---GTAGTAGTAGTAGTA     9137
Pachon15     GTTGTATGAGTAGAAGCATCACTAGTAGTAGTAGTAGT---A---GTAGTAGTAGTAGTA     9137
             **********************                        **************

Rascon4      GTATCAGTGGTAGTAGTAGTGG-----TATCATCAGTAGTAGTTTTAATAGTAGTAGTAT     9136
Surface      GTATCAGTGGTAGTAGTAGTGGTATCATATCATCAGTAGTAGTTTTAATAGTAGTAGTAT     9208
Rascon8      GTATCAGTGGTAGTAGTAGTGG-----TATCATCAGTAGTAGTTTTAATAGTAGTAGTAT     9198
Rascon2      GTATCAGTGGTAGTAGTAGTGG-----TATCATCAGTAGTAGTTTTAATAGTAGTAGTAT     9197
Rascon15     GTATCAGTGGTAGTAGTAGTGG-----TATCATCAGTAGTAGTTTTAATAGTAGTAGTAT     9180
Rascon13     GTATCAGTGGTAGTAGTAGTGG-----TATCATCAGTAGTAGTTTTAATAGTAGTAGTAT     9192
Rascon6      GTATCAGTGGTAGTAGTAGTGG-----TATCATCAGTAGTAGTTTTAATAGTAGTAGTAT     9194
Pachon14     GTATCAGTGGTAGTAGTAGTGG-----TATCATCAGTAGTAGTTTTAATAGTAGTAGTAT     9185
Pachon9      GTATCAGTGGTAGTAGTAGTGG-----TATCATCAGTAGTAGTTTTAATAGTAGTAGTAT     9194
Pachon17     GTATCAGTGGTAGTAGTAGTGG-----TATCATCAGTAGTAGTTTTAATAGTAGTAGTAT     9192
Pachon12     GTATCAGTGGTAGTAGTAGTGG-----TATCATCAGTAGTAGTTTTAATAGTAGTAGTAT     9174
Pachon11     GTATCAGTGGTAGTAGTAGTGG-----TATCATCAGTAGTAGTTTTAATAGTAGTAGTAT     9191
```

```
Pachon7     GTATCAGTGGTAGTAGTAGTGG-----TATCATCAGTAGTAGTTTTAATAGTAGTAGTAT    9192
Pachon3     GTATCAGTGGTAGTAGTAGTGG-----TATCATCAGTAGTAGTTTTAATAGTAGTAGTAT    9192
Pachon8     GTATCAGTGGTAGTAGTAGTGG-----TATCATCAGTAGTAGTTTTAATAGTAGTAGTAT    9192
Pachon15    GTATCAGTGGTAGTAGTAGTGG-----TATCATCAGTAGTAGTTTTAATAGTAGTAGTAT    9192
            **********************     *********************************

Rascon4     TTTTATCAGTAGTAGCATTAGTAGTAGTCGTATCAGTGGCAGTATTATCAGCAGTAGC--    9194
Surface     TTTTATCAGTAGTAGCATTAGTAGTAGTCGTATCAGTGGCAGTATTATCAGCAGTAGCAT    9268
Rascon8     TTTTATCAGTAGTAGCATTAGTAGTAGTCGTATCAGTGGCAGTATTATCAGCAGTAGC--    9256
Rascon2     TTTTATCAGTAGTAGCATTAGTAGTAGTCGTATCAGTGGCAGTATTATCAGCAGTAGCAT    9257
Rascon15    TTTTATCAGTAGTAGCATTAGTAGTAGTCGTATCAGTGGCAGTATTATCAGCAGTAGCAT    9240
Rascon13    TTTTATCAGTAGTAGCATTAGTAGTAGTCGTATCAGTGGCAGTATTATCAGCAGTAGCAT    9252
Rascon6     TTTTATCAGTAGTAGCATTAGTAGTAGTCGTATCAGTGGCAGTATTATCAGCAGTAGC--    9252
Pachon14    TTTTATCAGTAGTAGCATTAGTAGTAGTCGTATCAGTGGCAGTATTATCAGCAGTAGCAT    9245
Pachon9     TTTTATCAGTAGTAGCATTAGTAGTAGTCGTATCAGTGGCAGTATTATCAGCAGTAGCAT    9254
Pachon17    TTTTATCAGTAGTAGCATTAGTAGTAGTCGTATCAGTGGCAGTATTATCAGCAGTAGCAT    9252
Pachon12    TTTTATCAGTAGTAGCATTAGTAGTAGTCGTATCAGTGGCAGTATTATCAGCAGTAGCAT    9234
Pachon11    TTTTATCAGTAGTAGCATTAGTAGTAGTCGTATCAGTGGCAGTATTATCAGCAGTAGCAT    9251
Pachon7     TTTTATCAGTAGTAGCATTAGTAGTAGTCGTATCAGTGGCAGTATTATCAGCAGTAGCAT    9252
Pachon3     TTTTATCAGTAGTAGCATTAGTAGTAGTCGTATCAGTGGCAGTATTATCAGCAGTAGCAT    9252
Pachon8     TTTTATCAGTAGTAGCATTAGTAGTAGTCGTATCAGTGGCAGTATTATCAGCAGTAGCAT    9252
Pachon15    TTTTATCAGTAGTAGCATTAGTAGTAGTCGTATCAGTGGCAGTATTATCAGCAGTAGCAT    9252
            **********************************************************

Rascon4     -AGTAGAAGTAGCATCAGTAGAAGTAGTATCAGTAGTAGTAGTATTAATAGA---AGTAG    9250 Not in
Surface     AAGTAGAAGTAGCATCAGTAGAAGTAGTATCAGTAGTAGTAGTATTAA---------TAG    9319 Choy SF
Rascon8     -AGTAGAAGTAGCATCAGTAGAAGTAGTATCAGTATTAGTAGTATTAATAG------TAG    9309
Rascon2     AAGTAGAAGTAGCATCAGTAGAAGTAGTATCAGTAGTAGTAGTATTAATAG------TAG    9311
Rascon15    AAGTAGAAGTAGCATCAGTAGAAGTAGTATCAGTAGTAGTAGTATTAATAGTAGTAGTAG    9300
Rascon13    AAGTAGAAGTAGCATCAGTAGAAGTAGTATCAGTAGTAGTAGTATTAATAG------TAG    9306
Rascon6     -AGTAGAAGTAGCATCAGTAGAAGTAGTATCAGTAGTAGTAGTATTAATAG---TATTAG    9308
Pachon14    AAGTAGAAGTAGCATCAGTAGAAGTAGTATCAGTAGTAGTAGTATTAA------------    9293
Pachon9     AAGTAGAAGTAGCATCAGTAGAAGTAGTATCAGTAGTAGTAGTATTAA------------    9302
Pachon17    AAGTAGAAGTAGCATCAGTAGAAGTAGTATCAGTAGTAGTAGTATTAA------------    9300
Pachon12    AAGTAGAAGTAGCATCAGTAGAAGTAGTATCAGTAGTAGTAGTATTAA------------    9282
Pachon11    AAGTAGAAGTAGCATCAGTAGAAGTAGTATCAGTAGTAGTAGTATTAA------------    9299
Pachon7     AAGTAGAAGTAGCATCAGTAGAAGTAGTATCAGTAGTAGTAGTATTAA------------    9300
Pachon3     AAGTAGAAGTAGCATCAGTAGAAGTAGTATCAGTAGTAGTAGTATTAA------------    9300
Pachon8     AAGTAGAAGTAGCATCAGTAGAAGTAGTATCAGTAGTAGTAGTATTAA------------    9300
Pachon15    AAGTAGAAGTAGCATCAGTAGAAGTAGTATCAGTAGTAGTAGTATTAA------------    9300
             ********************************** ************

Rascon4     TAGTAGTAGTAGTAGTATCAGTAGTAGTATCAGTGGTAAGTGTTTTAATAGTAGTAGTAT    9310
Surface     TAGTAGTAGTAGTAGTATCAGTAGTAGTATCAGTGGTAAGTGTTTTAATAGTAGTAGTAT    9379
Rascon8     TAGTAGTAGTAGTAGTATCAGTAGTAGTATCAGTGGTAAGTGTTTTAATAGTAGTAGTAT    9369
Rascon2     TAGTAGTAGTAGTAGTATCAGTAGTAGTATCAGTGGTAAGTGTTTTAATAGTAGTAGTAT    9371
Rascon15    TAGTAGTAGTAGTAGTATCAGTAGTAGTATCAGTGGTAAGTGTTTTAATAGTAGTAGTAT    9360
Rascon13    TAGTAGTAGTAGTAGTATCAGTAGTAGTATCAGTGGTAAGTGTTTTAATAGTAGTAGTAT    9366
Rascon6     TAGTAGTAGTAGTAGTATCAGTAGTAGTATCAGTGGTAAGTGTTTTAATAGTAGTAGTAT    9368
Pachon14    ---TAGTAGTAGTAGTATCAGTAGTAGTATCAGTGGTAAGTGTTTTAATAGTAGTAGTAT    9350
Pachon9     ---TAGTAGTAGTAGTATCAGTAGTAGTATCAGTGGTAAGTGTTTTAATAGTAGTAGTAT    9359
Pachon17    ---TAGTAGTAGTAGTATCAGTAGTAGTATCAGTGGTAAGTGTTTTAATAGTAGTAGTAT    9357
Pachon12    ---TAGTAGTAGTAGTATCAGTAGTAGTATCAGTGGTAAGTGTTTTAATAGTAGTAGTAT    9339
Pachon11    ---TAGTAGTAGTAGTATCAGTAGTAGTATCAGTGGTAAGTGTTTTAATAGTAGTAGTAT    9356
Pachon7     ---TAGTAATAGTAGTATCAGTAGTAGTATCAGTGGTAAGTGTTTTAATAGTAGTAGTAT    9357
Pachon3     ---TAGTAGTAGTAGTATCAGTAGTAGTATCAGTGGTAAGTGTTTTAATAGTAGTAGTAT    9357
Pachon8     ---TAGTAGTAGTAGTATCAGTAGTAGTATCAGTGGTAAGTGTTTTAATAGTAGTAGTAT    9357
Pachon15    ---TAGTAGTAGTAGTATCAGTAGTAGTATCAGTGGTAAGTGTTTTAATAGTAGTAGTAT    9357
               ***** ***************************************************

Rascon4     TTGTATTAGTAGTATC---AGTAGTAGTAGTAATATCAGTGTTAGTAGTAGTGGTAGCAT    9367
Surface     TTGTATTAGTAGTATC---AGTAGTAGTAGTAATATCAGTGTTAGTAGTAGTGGTAGCAT    9436
Rascon8     TTGTATTAGTAGTATCAGTAGTAGTAGTAGTAATATCAGTGTTAGTAGTAGTGGTAGCAT    9429
Rascon2     TTGTATTAGTAGTATC---AGTAGTAGTAGTAATATCAGTGTTAGTAGTAGTGGTAGCAT    9428
Rascon15    TTGTATTAGTAGTATC---AGTAGTAGTAGTAATATCAGTGTTAGTAGTAGTGGTAGCAT    9417
Rascon13    TTGTATTAGTAGTATC---AGTAGTAGTAGTAATATCAGTGTTAGTAGTAGTGGTAGCAT    9423
Rascon6     TTGTATTAGTAGTATCAGTAGTAGTAGTAGTAATATCAGTGTTAGTAGTAGTGGTAGCAT    9428
Pachon14    TTGTATTAGTAGTATC---AGTAGTAGTAGTAATATCAGTGTTAGTAGTAGTGGTAGCAT    9407
Pachon9     TTGTATTAGTAGTATC---AGTAGTAGTAGTAATATCAGTGTTAGTAGTAGTGGTAGCAT    9416
Pachon17    TTGTATTAGTAGTATC---AGTAGTAGTAGTAATATCAGTGTTAGTAGTAGTGGTAGCAT    9414
Pachon12    TTGTATTAGTAGTATC---AGTAGTAGTAGTAATATCAGTGTTAGTAGTAGTGGTAGCAT    9396
Pachon11    TTGTATTAGTAGTATC---AGTAGTAGTAGTAATATCAGTGTTAGTAGTAGTGGTAGCAT    9413
Pachon7     TTGTATTAGTAGTATC---AGTAGTAGTAGTAATATCAGTGTTAGTAGTAGTGGTAGCAT    9414
Pachon3     TTGTATTAGTAGTATC---AGTAGTAGTAGTAATATCAGTGTTAGTAGTAGTGGTAGCAT    9414
Pachon8     TTGTATTAGTAGTATC---AGTAGTAGTAGTAATATCAGTGTTAGTAGTAGTGGTAGCAT    9414
Pachon15    TTGTATTAGTAGTATC---AGTAGTAGTAGTAATATCAGTGTTAGTAGTAGTGGTAGCAT    9414
            ****************   *****************************************
```

```
Rascon4         TAGTAGTAATAGTAGTAACTGTTAGTAGTATTTGTATCTGTTTAGTAGTATCAGTGCTAG       9427 Not in
Surface         TAGTAGTAATAGTAGTAACTGTTAGTAGTATTTGTATCTGTTTAGTAGTATCAGTGCTAG       9496 Choy SF
Rascon8         TAGTAGTAATAGTAGTAACTGTTAGTAGTATTTGTATCTGTTTAGTAGTATCAGTGCTAG       9489 Mix C/A
Rascon2         TAGTAGTAATAGTAGTAACTGTTAGTAGTATTTGTATCTGTTTAGTAGTATCAGTGCTAG       9488
Rascon15        TAGTAGTAATAGTAGTAACTGTTAGTAGTATTTGTATCTGTTTAGTAGTATCAGTGCTAG       9477
Rascon13        TAGTAGTAATAGTAGTAACTGTTAGTAGTATTTGTATCTGTTTAGTAGTATCAGTGCTAG       9483
Rascon6         TAGTAGTAATAGTAGTAACTGTTAGTAGTATTTGTATCTGTTTAGTAGTATCAGTGCTAG       9488
Pachon14        TAGTAGTAATAGTAGTAACTGTTAGTAGTATTTGTATCTGTTTAGTAGTATGAGTGCTAG       9467
Pachon9         TAGTAGTAATAGTAGTAACTGTTAGTAGTATTTGTATCTGTTTAGTAGTATGAGTGCTAG       9476
Pachon17        TAGTAGTAATAGTAGTAACTGTTAGTAGTATTTGTATCTGTTTAGTAGTATGAGTGCTAG       9474
Pachon12        TAGTAGTAATAGTAGTAACTGTTAGTAGTATTTGTATCTGTTTAGTAGTATGAGTGCTAG       9456
Pachon11        TAGTAGTAATAGTAGTAACTGTTAGTAGTATTTGTATCTGTTTAGTAGTATGAGTGCTAG       9473
Pachon7         TAGTAGTAATAGTAGTAACTGTTAGTAGTATTTGTATCTGTTTAGTAGTATGAGTGCTAG       9474
Pachon3         TAGTAGTAATAGTAGTAACTGTTAGTAGTATTTGTATCTGTTTAGTAGTATGAGTGCTAG       9474
Pachon8         TAGTAGTAATAGTAGTAACTGTTAGTAGTATTTGTATCTGTTTAGTAGTATGAGTGCTAG       9474
Pachon15        TAGTAGTAATAGTAGTAACTGTTAGTAGTATTTGTATCTGTTTAGTAGTATGAGTGCTAG       9474
                *************************************************** ********

Rascon4         TACTATTAGTTGTAGTAGTAATAGTAGCAGTATTGTTGTTCTTTGTCCACCAGATGGTGC       9487
Surface         TACTATTAGTTGTAGTAGTAATAGTAGCATTATTGTTGTTCTTTGTCCACCAGATGGTGC       9556
Rascon8         TACTATTAGTTGTAGTAGTAATAGTAGCAGTATTGTTGTTCTTTGTCCACCAGATGGTGC       9549
Rascon2         TACTATTAGTTGTAGTAGTAATAGTAGCAGTATTGTTGTTCTTTGTCCACCAGATGGTGC       9548
Rascon15        TACTATTAGTTGTAGTAGTAATAGTAGCAGTATTGTTGTTCTTTGTCCACCAGATGGTGC       9537
Rascon13        TACTATTAGTTGTAGTAGTAATAGTAGCAGTATTGTTGTTCTTTGTCCACCAGATGGTGC       9543
Rascon6         TACTATTAGTTGTAGTAGTAATAGTAGCAGTATTGTTGTTCTTTGTCCACCAGATGGTGC       9548
Pachon14        TACTATTAGTTGTAGTAGTAATAGTAGCAGTATTGTTGTTCTTTGTCCACCAGATGGTGC       9527
Pachon9         TACTATTAGTTGTAGTAGTAATAGTAGCAGTATTGTTGTTCTTTGTCCACCAGATGGTGC       9536
Pachon17        TACTATTAGTTGTAGTAGTAATAGTAGCAGTATTGTTGTTCTTTGTCCACCAGATGGTGC       9534
Pachon12        TACTATTAGTTGTAGTAGTAATAGTAGCAGTATTGTTGTTCTTTGTCCACCAGATGGTGC       9516
Pachon11        TACTATTAGTTGTAGTAGTAATAGTAGCAGTATTGTTGTTCTTTGTCCACCAGATGGTGC       9533
Pachon7         TACTATTAGTTGTAGTAGTAATAGTAGCAGTATTGTTGTTCTTTGTCCACCAGATGGTGC       9534
Pachon3         TACTATTAGTTGTAGTAGTAATAGTAGCAGTATTGTTGTTCTTTGTCCACCAGATGGTGC       9534
Pachon8         TACTATTAGTTGTAGTAGTAATAGTAGCAGTATTGTTGTTCTTTGTCCACCAGATGGTGC       9534
Pachon15        TACTATTAGTTGTAGTAGTAATAGTAGCAGTATTGTTGTTCTTTGTCCACCAGATGGTGC       9534
                **************************** *******************************

Rascon4         TGATTAAAACTAAATCAGTTTAGTCCTATAAACGGATAATACATTTGAATATGAAGCCTA       9547
Surface         TGATTAAAACTAAATCAGTTTAGTCCTATAAACGGATAATACATTTGAATATGAAGCCTA       9616
Rascon8         TGATTAAAACTAAATCAGTTTAGTCCTATAAACGGATAATACATTTGAATATGAAGCCTA       9609
Rascon2         TGATTAAAACTAAATCAGTTTAGTCCTATAAACGGATAATACATTTGAATATGAAGCCTA       9608
Rascon15        TGATTAAAACTAAATCAGTTTAGTCCTATAAACGGATAATACATTTGAATATGAAGCCTA       9597
Rascon13        TGATTAAAACTAAATCAGTTTAGTCCTATAAACGGATAATACATTTGAATATGAAGCCTA       9603
Rascon6         TGATTAAAACTAAATCAGTTTAGTCCTATAAACGGATAATACATTTGAATATGAAGCCTA       9608
Pachon14        TGATTAAAACTAAATCAGTTTAGTCCTATAAACGGATAATACATTTGAATATGAAGCCTA       9587
Pachon9         TGATTAAAACTAAATCAGTTTAGTCCTATAAACGGATAATACATTTGAATATGAAGCCTA       9596
Pachon17        TGATTAAAACTAAATCAGTTTAGTCCTATAAACGGATAATACATTTGAATATGAAGCCTA       9594
Pachon12        TGATTAAAACTAAATCAGTTTAGTCCTATAAACGGATAATACATTTGAATATGAAGCCTA       9576
Pachon11        TGATTAAAACTAAATCAGTTTAGTCCTATAAACGGATAATACATTTGAATATGAAGCCTA       9593
Pachon7         TGATTAAAACTAAATCAGTTTAGTCCTATAAACGGATAATACATTTGAATATGAAGCCTA       9594
Pachon3         TGATTAAAACTAAATCAGTTTAGTCCTATAAACGGATAATACATTTGAATATGAAGCCTA       9594
Pachon8         TGATTAAAACTAAATCAGTTTAGTCCTATAAACGGATAATACATTTGAATATGAAGCCTA       9594
Pachon15        TGATTAAAACTAAATCAGTTTAGTCCTATAAACGGATAATACATTTGAATATGAAGCCTA       9594
                ************************************************************

Rascon4         TTCAAAAGGAATAGAGCTGAAAGAATTTTATACATAATTAATAACATTATTTGCTTTATT       9607
Surface         TTCAAAAGGAATAGAGCTGAAAGAATTTTATACATAATTAATAACATGATTTGCTTTATT       9676
Rascon8         TTCAAAAGGAATAGAGCTGAAAGAATTTTATACATAATTAATAACATTATTTGCTTTATT       9669
Rascon2         TTCAAAAGGAATAGAGCTGAAAGAATTTTATACATAATTAATAACATTATTTGCTTTATT       9668
Rascon15        TTCAAAAGGAATAGAGCTGAAAGAATTTTATACATAATTAATAACATTATTTGCTTTATT       9657
Rascon13        TTCAAAAGGAATAGAGCTGAAAGAATTTTATACATAATTAATAACATTATTTGCTTTATT       9663
Rascon6         TTCAAAAGGAATAGAGCTGAAAGAATGTTATACATAATTAATAACATTATTTGCTTTATT       9668
Pachon14        TTCAAAAGGAATAGAGCTGAAAGAATTTTATACATAATTAATAACATTATTTGCTTTATT       9647
Pachon9         TTCAAAAGGAATAGAGCTGAAAGAATTTTATACATAATTAATAACATTATTTGCTTTATT       9656
Pachon17        TTCAAAAGGAATAGAGCTGAAAGAATTTTATACATAATTAATAACATTATTTGCTTTATT       9654
Pachon12        TTCAAAAGGAATAGAGCTGAAAGAATTTTATACATAATTAATAACATTATTTGCTTTATT       9636
Pachon11        TTCAAAAGGAATAGAGCTGAAAGAATTTTATACATAATTAATAACATTATTTGCTTTATT       9653
Pachon7         TTCAAAAGGAATAGAGCTGAAAGAATTTTATACATAATTAATAACATTATTTGCTTTATT       9654
Pachon3         TTCAAAAGGAATAGAGCTGAAAGAATTTTATACATAATTAATAACATTATTTGCTTTATT       9654
Pachon8         TTCAAAAGGAATAGAGCTGAAAGAATTTTATACATAATTAATAACATTATTTGCTTTATT       9654
Pachon15        TTCAAAAGGAATAGAGCTGAAAGAATTTTATACATAATTAATAACATTATTTGCTTTATT       9654
                ************************** ******************** ************

Rascon4         CCAGAAATGAAATTTTATTTTTAATCTATGATACAGTAAGCAACATAAACCATACATTTT       9667
Surface         CCAGAAATGAAATTTTATTTTTAATCTATGATACAGTAAGCAACATAAACCATACATTTT       9736
Rascon8         CCAGAAATGAAATTTTATTTTTAATCTATGATACAGTAAGCAACATAAACCATACATTTT       9729
Rascon2         CCAGAAATGAAATTTTATTTTTAATCTATGATACAGTAAGCAACATAAACCATACATTTT       9728
```

```
Rascon15     CCAGAAATGAAATTTTATTTTTAATCTATGATACAGTAAGCAACATAAACCATACATTTT     9717
Rascon13     CCAGAAATGAAATTTTATTTTTAATCTATGATACAGTAAGCAACATAAACCATACATTTT     9723
Rascon6      CCAGAAATGAAAATTTATTTTTAATCTATGATACAGTAAGCAACATAAACCATACATTTT     9728
Pachon14     CCAGAAATGAAATTTTATTTTTAATCTATGATACAGTAAGCAACATAAACCATACATTTT     9707
Pachon9      CCAGAAATGAAATTTTATTTTTAATCTATGATACAGTAAGCAACATAAACCATACATTTT     9716
Pachon17     CCAGAAATGAAATTTTATTTTTAATCTATGATACAGTAAGCAACATAAACCATACATTTT     9714
Pachon12     CCAGAAATGAAATTTTATTTTTAATCTATGATACAGTAAGCAACATAAACCATACATTTT     9696
Pachon11     CCAGAAATGAAATTTTATTTTTAATCTATGATACAGTAAGCAACATAAACCATACATTTT     9713
Pachon7      CCAGAAATGAAATTTTATTTTTAATCTATGATACAGTAAGCAACATAAACCATACATTTT     9714
Pachon3      CCAGAAATGAAATTTTATTTTTAATCTATGATACAGTAAGCAACATAAACCATACATTTT     9714
Pachon8      CCAGAAATGAAATTTTATTTTTAATCTATGATACAGTAAGCAACATAAACCATACATTTT     9714
Pachon15     CCAGAAATGAAATTTTATTTTTAATCTATGATACAGTAAGCAACATAAACCATACATTTT     9714
             ************ ***********************************************

Rascon4      TATTTTACTTTCCATTCAGTCTATTTGATCCTTCTATTCACATAGTGTATATTACGTTAA     9727
Surface      TATTTTACTTTCCATTCAGTCTATTTGATCCTTCTATTCACATAGTGTATATTACGTTAA     9796
Rascon8      TATTTTACTTTCCATTCAGTCTATTTGATCCTTCTATTCACATAGTGTATATTACGTTAA     9789
Rascon2      TATTTTACTTTCCATTCAGTCTATTTGATCCTTCTATTCACATAGTGTATATTACGTTAA     9788
Rascon15     TATTTTACTTTCCATTCAGTCTATTTGATCCTTCTATTCACATAGTGTATATTACGTTAA     9777
Rascon13     TATTTTACTTTCCATTCAGTCTATTTGATCCTTCTATTCACATAGTGTATATTACGTTAA     9783
Rascon6      TATTTTACTTTCCATTCAGTCTATTTGATCCTTCTATTCACATAGTGTATATTACGTTAA     9788
Pachon14     TATTTTACTTTCCATTCAGTCTATTTGATCCTTCTATTCACATAGTGTATATTACGTTAA     9767
Pachon9      TATTTTACTTTCCATTCAGTCTATTTGATCCTTCTATTCACATAGTGTATATTACGTTAA     9776
Pachon17     TATTTTACTTTCCATTCAGTCTATTTGATCTTTCTATTCACATAGTGTATATTACGTTAA     9774
Pachon12     TATTTTACTTTCCATTCAGTCTATTTGATCCTTCTATTCACATAGTGTATATTACGTTAA     9756
Pachon11     TATTTTACTTTCCATTCAGTCTATTTGATCTTTCTATTCACATAGTGTATATTACGTTAA     9773
Pachon7      TATTTTACTTTCCATTCAGTCTATTTGATCCTTCTATTCACATAGTGTATATTACGTTAA     9774
Pachon3      TATTTTACTTTCCATTCAGTCTATTTGATCTTTCTATTCACATAGTGTATATTACGTTAA     9774
Pachon8      TATTTTACTTTCCATTCAGTCTATTTGATCCTTCTATTCACATAGTGTATATTACGTTAA     9774
Pachon15     TATTTTACTTTCCATTCAGTCTATTTGATCCTTCTATTCACATAGTGTATATTACGTTAA     9774
             ****************************** *****************************

Rascon4      TGTTAAGGGTGTAATATCCCTATCCTTTACTGTTGGTCATTCGACCAACCCTCTTGGATC     9787
Surface      TGTTACGGGTGTAATATCCCTATCCTTTACTGTTGGTCATTCGACCAACCCTCTTGGATC     9856
Rascon8      TGTTAAGGGTGTAATATCCCTATCCTTTACTGTTGGTCATTCGACCAACCCTCTTGGATC     9849
Rascon2      TGTTAAGGGTGTAATATCCCTATCCTTTACTGTTGGTCATTCGACCAACCCTCTTGGATC     9848
Rascon15     TGTTAAGGGTGTAATATCCCTATCCTTTACTGTTGGTCATTCGACCAACCCTCTTGGATC     9837
Rascon13     TGTTAAGGGTGTAATATCCCTATCCTTTACTGTTGGTCATTCGACCAACCCTCTTGGATC     9843
Rascon6      TGTTAAGGGTGTAATATCCCTATCCTTTACTGTTGGTCATTCGACCAACCCTCTTGGATC     9848
Pachon14     TGTTAAGGGTGTAATATCCCTATCCTTTACTGTTGGTCATTCGACCAACCCTCTTGGATC     9827
Pachon9      TGTTAAGGGTGTAATATCCCTATCCTTTACTGTTGGTCATTCGACCAACCCTCTTGGATC     9836
Pachon17     TGTTAAGGGTGTAATATCCCTATCCTTTACTGTTGGTCATTCGACCAACCCTCTTGGATC     9834
Pachon12     TGTTAAGGGTGTAATATCCCTATCCTTTACTGTTGGTCATTCGACCAACCCTCTTGGATC     9816
Pachon11     TGTTAAGGGTGTAATATCCCTATCCTTTACTGTTGGTCATTCGACCAACCCTCTTGGATC     9833
Pachon7      TGTTAAGGGTGTAATATCCCTATCCTTTACTGTTGGTCATTCGACCAACCCTCTTGGATC     9834
Pachon3      TGTTAAGGGTGTAATATCCCTATCCTTTACTGTTGGTCATTCGACCAACCCTCTTGGATC     9834
Pachon8      TGTTAAGGGTGTAATATCCCTATCCTTTACTGTTGGTCATTCGACCAACCCTCTTGGATC     9834
Pachon15     TGTTAAGGGTGTAATATCCCTATCCTTTACTGTTGGTCATTCGACCAACCCTCTTGGATC     9834
             ***** ******************************************************

Rascon4      CATCATTGCTGTAGCATAAGTATAAGGATTCGTGAGTCAGAAAGGGTGATATGTGCTTTG     9847
Surface      CATCATTGCTGTAGCATAAGTATAAGGATTCGTGAGTCAGAAAGGGTGATATGTGCTTTG     9916
Rascon8      CATCATTGCTGTAGCATAAGTATAAGGATTCGTGAGTCAGAAAGGGTGATATGTGCTTTG     9909
Rascon2      CATCATTGCTGTAGCATAAGTATAAGGATTCGTGAGTCAGAAAGGGTGATATGTGCTTTG     9908
Rascon15     CATCATTGCTGTAGCATAAGTATAAGGATTCGTGAGTCAGAAAGGGTGATATGTGCTTTG     9897
Rascon13     CATCATTGCTGTAGCATAAGTATAAGGATTCGTGAGTCAGAAAGGGTGATATGTGCTTTG     9903
Rascon6      CATCATTGCTGTAGCATAAGTATAAGGATTCGTGAGTCAGAAAGGGTGATATGTGCTTTG     9908
Pachon14     CATCATTGCTGTAGCATAAGTATAAGGATTCGTGAGTCAGAAAGGGTGATATGTGCTTTG     9887
Pachon9      CATCATTGCTGTAGCATAAGTATAAGGATTCGTGAGTCAGAAAGGGTGATATGTGCTTTG     9896
Pachon17     CATCATTGCTGTAGCATAAGTATAAGGATTCGTGAGTCAGAAAGGGTGATATGTGCTTTG     9894
Pachon12     CATCATTGCTGTAGCATAAGTATAAGGATTCGTGAGTCAGAAAGGGTGATATGTGCTTTG     9876
Pachon11     CATCATTGCTGTAGCATAAGTATAAGGATTCGTGAGTCAGAAAGGGTGATATGTGCTTTG     9893
Pachon7      CATCATTGCTGTAGCATAAGTATAAGGATTCGTGAGTCAGAAAGGGTGATATGTGCTTTG     9894
Pachon3      CATCATTGCTGTAGCATAAGTATAAGGATTCGTGAGTCAGAAAGGGTGATATGTGCTTTG     9894
Pachon8      CATCATTGCTGTAGCATAAGTATAAGGATTCGTGAGTCAGAAAGGGTGATATGTGCTTTG     9894
Pachon15     CATCATTGCTGTAGCATAAGTATAAGGATTCGTGAGTCAGAAAGGGTGATATGTGCTTTG     9894
             ************************************************************

Rascon4      TAAAATAAATGCATCCCTTCATAGGAGCCACTTCTGTCGGGTTAGATAAGCTCTCCTGCT     9907
Surface      TAAAATAAATGCATCCTTTCATAGGAGCCACTTCTGTCGTGTTAGATAAGCTCTCCTGCT     9976
Rascon8      TAAAATAAATGCATCCTTTCATAGGAGCCACTTCTGTCGGGTTAGATAAGCTCTCCTGCT     9969
Rascon2      TAAAATAAATGCATCCTTTCATAGGAGCCACTTCTGTCGGGTTAGATAAGCTCTCCTGCT     9968
Rascon15     TAAAATAAATGCATCCCTTCATAGGAGCCACTTCTGTCGGGTTAGATAAGCTCTCCTGCT     9957
Rascon13     TAAAATAAATGCATCCTTTCATAGGAGCCACTTCTGTCGGGTTAGATAAGCTCTCCTGCT     9963
Rascon6      TAAAATAAATGCATCCTTTCATAGGAGCCACTTCTGTCGGGTTAGATAAGCTCTCCTGCT     9968
Pachon14     TAAAATAAATGCATCCTTTCATAGGAGCCACTTCTGTCGGGTTAGATAAGCTCTCCTGCT     9947
Pachon9      TAAAATAAATGCATCCTTTCATAGGAGCCACTTCTGTCGGGTTAGATAAGCTCTCCTGCT     9956
```

```
Pachon17   TAAAATAAATGCATCCTTTCATAGGAGCCACTTCTGTCGGGTTAGATAAGCTCTCCTGCT   9954
Pachon12   TAAAATAAATGCATCCTTTCATAGGAGCCACTTCTGTCGGGTTAGATAAGCTCTCCTGCT   9936
Pachon11   TAAAATAAATGCATCCTTTCATAGGAGCCACTTCTGTCGGGTTAGATAAGCTCTCCTGCT   9953
Pachon7    TAAAATAAATGCATCCTTTCATAGGAGCCACTTCTGTCGGGTTAGATAAGCTCTCCTGCT   9954
Pachon3    TAAAATAAATGCATCCTTTCATAGGAGCCACTTCTGTCGGGTTAGATAAGCTCTCCTGCT   9954
Pachon8    TAAAATAAATGCATCCTTTCATAGGAGCCACTTCTGTCGGGTTAGATAAGCTCTCCTGCT   9954
Pachon15   TAAAATAAATGCATCCTTTCATAGGAGCCACTTCTGTCGGGTTAGATAAGCTCTCCTGCT   9954
           **************** ********************** ********************

Rascon4    TAGCCTAATAAATTTAGTTTAGATTCACTGCATAGTTTTGGGTTTAAGATTCTGCTCCAA   9967
Surface    TAGCCTAATAAATTTAGTTTAGATTCACTGCATAGTTTTGGGTTTAAGATTCTGCTCCAA   10036
Rascon8    TAGCCTCATAAATTTAGTTTAGATTCACTGCATAGTTTTGGGTTTAAGATTCTGCTCCAA   10029
Rascon2    TAGCCTAATAAATTTAGTTTAGATTCACTGCATAGTTTTGGGTTTAAGATTCTGCTCCAA   10028
Rascon15   TAGCCTAATAAATTTAGTTTAGATTCACTGCATAGTTTTGGGTTTAAGATTCTGCTCCAA   10017
Rascon13   TAGCCTAATAAATTTAGTTTAGATTCACTGCATAGTTTTGGGTTTAAGATTCTGCTCCAA   10023
Rascon6    TAGCCTAATAAATTTAGTTTAGATTCACTGCATAGTTTTGGGTTTAAGATTCTGCTCCAA   10028
Pachon14   TAGCCTAATAAATTTAGTTTAGATTCACTGCATAGTTTTGGGTTTAAGATTCTGCTCCAA   10007
Pachon9    TAGCCTAATAAATTTAGTTTAGATTCACTGCATAGTTTTGGGTTTAAGATTCTGCTCCAA   10016
Pachon17   TAGCCTAATAAATTTAGTTTAGATTCACTGCATAGTTTTGGGTTTAAGATTCTGCTCCAA   10014
Pachon12   TAGCCTAATAAATTTAGTTTAGATTCACTGCATAGTTTTGGGTTTAAGATTCTGCTCCAA   9996
Pachon11   TAGCCTAATAAATTTAGTTTAGATTCACTGCATAGTTTTGGGTTTAAGATTCTGCTCCAA   10013
Pachon7    TAGCCTAATAAATTTAGTTTAGATTCACTGCATAGTTTTGGGTTTAAGATTCTGCTCCAA   10014
Pachon3    TAGCCTAATAAATTTAGTTTAGATTCACTGCATAGTTTTGGGTTTAAGATTCTGCTCCAA   10014
Pachon8    TAGCCTAATAAATTTAGTTTAGATTCACTGCATAGTTTTGGGTTTAAGATTCTGCTCCAA   10014
Pachon15   TAGCCTAATAAATTTAGTTTAGATTCACTGCATAGTTTTGGGTTTAAGATTCTGCTCCAA   10014
           ****** *****************************************************

Rascon4    GCTGTTGTAGCTAATTAAAACAATCAGTTTGGAGTCTACAACAACACAAACACAAAAAAA   10027 Not in
Surface    GCTGTTGTAGCTAATTAAAACAATCAGTTTGGAGTCTACAACAACACAAACACAAAAATA   10096 Choy SF
Rascon8    GCTGTTGTAGCTAATTAAAACAATCAGTTTGGAGTCTACAACAACACAAACACAAAAATA   10089 Mix A/G
Rascon2    GCTGTTGTAGCTAATTAAAACAATCAGTTTGGAGTCTACAACAACACAAACACAAAAATA   10088
Rascon15   GCTGTTGTAGCTAATTAAAACAATCAGTTTGGAGTCTACAACAACACAAACACAAAAATA   10077
Rascon13   GCTGTTGTAGCTAATTAAAACAATCAGTTTGGAGTCTACAACAACACAAACACAAAAATA   10083
Rascon6    GCTGTTGTAGCTAATTAAAACAATCAGTTTGGAGTCTACAACAACACAAACACAAAAATA   10088
Pachon14   GCTGTTGTAGCTAATTAAAACAATCAGTTTGGAGTCTACAACAACACAAGCACAAAAATA   10067
Pachon9    GCTGTTGTAGCTAATTAAAACAATCAGTTTGGAGTCTACAACAACACAAGCACAAAAATA   10076
Pachon17   GCTGTTGTAGCTAATTAAAACAATCAGTTTGGAGTCTACAACAACACAAGCACAAAAATA   10074
Pachon12   GCTGTTGTAGCTAATTAAAACAATCAGTTTGGAGTCTACAACAACACAAGCACAAAAATA   10056
Pachon11   GCTGTTGTAGCTAATTAAAACAATCAGTTTGGAGTCTACAACAACACAAGCACAAAAATA   10073
Pachon7    GCTGTTGTAGCTAATTAAAACAATCAGTTTGGAGTCTACAACAACACAAGCACAAAAATA   10074
Pachon3    GCTGTTGTAGCTAATTAAAACAATCAGTTTGGAGTCTACAACAACACAAGCACAAAAATA   10074
Pachon8    GCTGTTGTAGCTAATTAAAACAATCAGTTTGGAGTCTACAACAACACAAGCACAAAAATA   10074
Pachon15   GCTGTTGTAGCTAATTAAAACAATCAGTTTGGAGTCTACAACAACACAAGCACAAAAATA   10074
           ************************************************* ******** *

Rascon4    AATATCATTTTAAATGATTAGGTGCTCCAAATTTCACTATTTTAGAGCTGCGCTCTACAT   10087
Surface    AATATCATTTTAAAGGATTAGGTGCTCCAAATTTCACTATTTTAGAGCTGCGCTCTACAC   10156
Rascon8    AATATCATTTTAAATGATTAGGTGCTCCAAATTCCACTATTTTAGAGCTGCGCTCTACAT   10149
Rascon2    AATATCATTTTAAATGATTAGGTGCTCCAAATTCCACTATTTTAGAGCTGTGCTCTACAT   10148
Rascon15   AATATCATTTTAAATGATTAGGTGCTCCAAATTTCACTATTTTAGAGCTGCGCTCTACAT   10137
Rascon13   AATATCATTTTAAATGATTAGGTGCTCCAAATTTCACTATTTTAGAGCTGCGCTCTACAT   10143
Rascon6    AATATCATTTTAAATGATTAGGTGCTCCAAATTTCACTATTTTAGAGCTGCGCTCTACAT   10148
Pachon14   AATATCATTTTAAAGGATTAGGTGCTCCAAATTTCACTATTTTAGAGCTGCGCTCTACAT   10127
Pachon9    AATATCATTTTAAAGGATTAGGTGCTCCAAATTTCACTATTTTAGAGCTGCGCTCTACAT   10136
Pachon17   AATATCATTTTAAAGGATTAGGTGCTCCAAATTTCACTATTTTAGAGCTGCGCTCTACAT   10134
Pachon12   AATATCATTTTAAAGGATTAGGTGCTCCAAATTTCACTATTTTAGAGCTGCGCTCTACAT   10116
Pachon11   AATATCATTTTAAAGGATTAGGTGCTCCAAATTTCACTATTTTAGAGCTGCGCTCTACAT   10133
Pachon7    AATATCATTTTAAAGGATTAGGTGCTCCAAATTTCACTATTTTAGAGCTGCGCTCTACAT   10134
Pachon3    AATATCATTTTAAAGGATTAGGTGCTCCAAATTTCACTATTTTAGAGCTGCGCTCTACAT   10134
Pachon8    AATATCATTTTAAAGGATTAGGTGCTCCAAATTTCACTATTTTAGAGCTGCGCTCTACAT   10134
Pachon15   AATATCATTTTAAAGGATTAGGTGCTCCAAATTTCACTATTTTAGAGCTGCGCTCTACAT   10134
           ************** ****************** **************** ******** 

Rascon4    GCTGAATGTGATTTTCCTTAGCTTAAGAAATATCACGATAATTCTAAAAAGCTCGCTTTT   10147
Surface    GCTGAATATGATTTTCCTTAGCTTAAGAAATATCACGATAATTCTAAAAAGCTCGCTTTT   10216
Rascon8    GCTGAATGTGATTTTCCTTAGCTTAAGAAATATCACGATAATTCTAAAAAGCTCGCTTTT   10209
Rascon2    GCTGAATGTGATTTTCCTTAGCTTAAGAAATATCACGATAATTCTAAAAAGCTCGCTTTT   10208
Rascon15   GCTGAATGTGATTTTCCTTAGCTTAAGAAATATCACGATAATTCTAAAAAGCTCGCTTTT   10197
Rascon13   GCTGAATGTGATTTTCCTTAGCTTAAGAAATATCACGATAATTCTAAAAAGCTCGCTTTT   10203
Rascon6    GCTGAATGTGATTTTCCTTAGCTTAAGAAATATCACGATAATTCTAAAAAGCTCGCTTTT   10208
Pachon14   GCTGAATGTGATTTTCCTTAGCTTAAGAAATATCACGATAATTCTAAAAAGCTCGCTTTT   10187
Pachon9    GCTGAATGTGATTTTCCTTAGCTTAAGAAATATCACGATAATTCTAAAAAGCTCGCTTTT   10196
Pachon17   GCTGAATGTGATTTTCCTTAGCTTAAGAAATATCACGATAATTCTAAAAAGCTCGCTTTT   10194
Pachon12   GCTGAATGTGATTTTCCTTAGCTTAAGAAATATCACGATAATTCTAAAAAGCTCGCTTTT   10176
Pachon11   GCTGAATGTGATTTTCCTTAGCTTAAGAAATATCACGATAATTCTAAAAAGCTCGCTTTT   10193
Pachon7    GCTGAATGTGATTTTCCTTAGCTTAAGAAATATCACGATAATTCTAAAAAGCTCGCTTTT   10194
Pachon3    GCTGAATGTGATTTTCCTTAGCTTAAGAAATATCACGATAATTCTAAAAAGCTCGCTTTT   10194
```

```
Pachon8       GCTGAATGTGATTTTCCTTAGCTTAAGAAATATCACGATAATTCTAAAAAGCTCGCTTTT     10194
Pachon15      GCTGAATGTGATTTTCCTTAGCTTAAGAAATATCACGATAATTCTAAAAAGCTCGCTTTT     10194
              ******* ****************************************************

Rascon4       GAATGGAGCATCTCTCTCCTCAAAGCATTTAAAGTCATTGAGTCAATCATATGTCATCTG     10207
Surface       GAATGGAGCATCTCTCTCCTCAAAGCATTTAAAGTCATTGAGTCAATCATATGTCATCTG     10276
Rascon8       GAATGGAGCATCTCTCTCCTCAAAGCATTTAAAGTCATTGAGTCAATCATATGTCATCTG     10269
Rascon2       GAATGGAGCATCTCTCTCCTCAAAGCATTTAAAGTCATTGAGTCAATCATATGTCATCTG     10268
Rascon15      GAATGGAGCATCTCTCTCCTCAAAGCATTTAAAGTCATTGAGTCAATCATATGTCATCTG     10257
Rascon13      GAATGGAGCATCTCTCTCCTCAAAGCATTTAAAGTCATTGAGTCAATCATATGTCATATG     10263
Rascon6       GAATGGAGCATCTCTCTCCTCAAAGCATTTAAAGTCATTGAGTCAATCATATGTCATCTG     10268
Pachon14      GAATGGAGCATCTCTCTCCTCAAAGCATTTAAAGTCATTGAGTCAATCATATGTCATCTG     10247
Pachon9       GAATGGAGCATCTCTCTCCTCAAAGCATTTAAAGTCATTGAGTCAATCATATGTCATCTG     10256
Pachon17      GAATGGAGCATCTCTCTCCTCAAAGCATTTAAAGTCATTGAGTCAATCATATGTCATCTG     10254
Pachon12      GAATGGAGCATCTCTCTCCTCAAAGCATTTAAAGTCATTGAGTCAATCATATGTCATCTG     10236
Pachon11      GAATGGAGCATCTCTCTCCTCAAAGCATTTAAAGTCATTGAGTCAATCATATGTCATCTG     10253
Pachon7       GAATGGAGCATCTCTCTCCTCAAAGCATTTAAAGTCATTGAGTCAATCATATGTCATCTG     10254
Pachon3       GAATGGAGCATCTCTCTCCTCAAAGCATTTAAAGTCATTGAGTCAATCATATGTCATCTG     10254
Pachon8       GAATGGAGCATCTCTCTCCTCAAAGCATTTAAAGTCATTGAGTCAATCATATGTCATCTG     10254
Pachon15      GAATGGAGCATCTCTCTCCTCAAAGCATTTAAAGTCATTGAGTCAATCATATGTCATCTG     10254
              ********************************************************* **

Rascon4       TAAGGCCTTCAGGACAGTTTCTATCATTATAGCATTATTAGACACGGCAAAACACACAGA     10267
Surface       TAAGGCCTTCAGGACAGTTTCTATCATTATAGCATTATTAGACACGGCAAAACACACAGA     10336
Rascon8       TAAGGCCTTCAGGACAGTTTCTATCATTATAGCATTATTAGACACGGCAAAACACACAGA     10329
Rascon2       TAAGGCCTTCAGGACAGTTTCTATCATTATAGCATTATTAGACACGGCAAAACACACAGA     10328
Rascon15      TAAGGCCTTCAGGACAGTTTCTATCATTATAGCATTATTAGACACGGCAAAACACACAGA     10317
Rascon13      TAAGGCCTTCAGGACAGTTTCTATCATTATAGCATTATTAGACACGGCAAAACACACAGA     10323
Rascon6       TTAGGCCTTCAGGACAGTTTCTATCATTATAGCATTATTAGACACGGCAAAACACACAGA     10328
Pachon14      TAAGGCCTTCAGGACAGTTTCTATCATTATAGCATTATTAGACACGGCAAAACACACAGA     10307
Pachon9       TAAGGCCTTCAGGACAGTTTCTATCATTATAGCATTATTAGACACGGCAAAACACACAGA     10316
Pachon17      TAAGGCCTTCAGGACAGTTTCTATCATTATAGCATTATTAGACACGGCAAAACACACAGA     10314
Pachon12      TAAGGCCTTCAGGACAGTTTCTATCATTATAGCATTATTAGACACGGCAAAACACACAGA     10296
Pachon11      TAAGGCCTTCAGGACAGTTTCTATCATTATAGCATTATTAGACACGGCAAAACACACAGA     10313
Pachon7       TAAGGCCTTCAGGACAGTTTCTATCATTATAGCATTATTAGACACGGCAAAACACACAGA     10314
Pachon3       TAAGGCCTTCAGGACAGTTTCTATCATTATAGCATTATTAGACACGGCAAAACACACAGA     10314
Pachon8       TAAGGCCTTCAGGACAGTTTCTATCATTATAGCATTATTAGACACGGCAAAACACACAGA     10314
Pachon15      TAAGGCCTTCAGGACAGTTTCTATCATTATAGCATTATTAGACACGGCAAAACACACAGA     10314
              * **********************************************************

Rascon4       ACA---GTTTTACATCTCCATTATTAGCTATTCATCTGTAACAAATGAGCAGAGCAAACA     10324
Surface       ACA---GTTTTACATCTCCATTATTAGCTATTCATCTGTAACAAATGAGCAGAGCAAACA     10393
Rascon8       ACA---GTTTTACATCTCCATTATTAGCTATTCATCTGTAACAAATGAGCAGAGCAAACA     10386
Rascon2       ACA---GTTTTACATCTCCATTATTAGCTATTCATCTGTAACAAATGAGCAGAGCAAACA     10385
Rascon15      ACA---GTTTTACATCTACATTATTAGCTATTCATCTGTAACAAATGAGCAGAGCAAACA     10374
Rascon13      ACAAGTGTTTTACATCTCCATTATTAGCTATTCATCTGTAACAAATGAGCAGAGAAAACA     10383
Rascon6       ACAAGTGTTTTACATCTACATTATTAGCTATTCATCTGTAACAAATGAGCAGAGCAAACA     10388
Pachon14      ACAAGTGTTTTACATCTCCATTATTAGCTATTCATCTGTAACAAATGAGCAGAGCAAACA     10367
Pachon9       ACAAGTGTTTTACATCTCCATTATTAGCTATTCATCTGTAACAAATGAGCAGAGCAAACA     10376
Pachon17      ACAAGTGTTTTACATCTCCATTATTAGCTATTCATCTGTAACAAATGAGCAGAGCAAACA     10374
Pachon12      ACAAGTGTTTTACATCTCCATTATTAGCTATTCATCTGTAACAAATGAGCAGAGCAAACA     10356
Pachon11      ACAAGTGTTTTACATCTCCATTATTAGCTATTCATCTGTAACAAATGAGCAGAGCAAACA     10373
Pachon7       ACAAGTGTTTTACATCTCCATTATTAGCTATTCATCTGTAACAAATGAGCAGAGCAAACA     10374
Pachon3       ACAAGTGTTTTACATCTCCATTATTAGCTATTCATCTGTAACAAATGAGCAGAGCAAACA     10374
Pachon8       ACAAGTGTTTTACATCTCCATTATTAGCTATTCATCTGTAACAAATGAGCAGAGCAAACA     10374
Pachon15      ACAAGTGTTTTACATCTCCATTATTAGCTATTCATCTGTAACAAATGAGCAGAGCAAACA     10374
              ***   *********** ************************************ *****

Rascon4       GCAGAGAGCAGAGCGTCAGGATGTGAGAACAGGATGCACTTAAGCACATAAAAAAGAAAC     10384
Surface       GCAGAGAGCAGAGCGTCAGGATGTGAGAACAGGATGCACTTAAGCACATAAAAAAGAAAC     10453
Rascon8       GCAGAGAGCAGAGCGTCAGGATGTGAGAACAGGATGCACTTAAGCACATAAAAAAGAAAC     10446
Rascon2       GCAGAGAGCAGAGCGTCAGGATGTGAGAACAGGATGCACTTAAGCACATAAAAAAGAAAC     10445
Rascon15      GCAGAGAGCAGAGCGTCAGGATGTGAGAACAGGATGCACTTAAGCACATAAAAAAGAAAC     10434
Rascon13      GCAGAGAGCAGAGCGTCAGGATGTGAGAACAGGATGCACTTAAGCACATAAAAAAGAAAC     10443
Rascon6       GCAGAGAGCAGAGCGTCAGGATGTGAGAACAGGATGCACTTAAGCACATAAAAAAGAAAC     10448
Pachon14      GCAGAGAGCAGAGCGTCAGGATGTGAGAACAGGATGCACTTAAGCACATAAAAAAGAAAC     10427
Pachon9       GCAGAGAGCAGAGCGTCAGGATGTGAGAACAGGATGCACTTAAGCACATAAAAAAGAAAC     10436
Pachon17      GCAGAGAGCAGAGCGTCAGGATGTGAGAACAGGATGCACTTAAGCACATAAAAAAGAAAC     10434
Pachon12      GCAGAGAGCAGAGCGTCAGGATGTGAGAACAGGATGCACTTAAGCACATAAAAAAGAAAC     10416
Pachon11      GCAGAGAGCAGAGCGTCAGGATGTGAGAACAGGATGCACTTAAGCACATAAAAAAGAAAC     10433
Pachon7       GCAGAGAGCAGAGCGTCAGGATGTGAGAACAGGATGCACTTAAGCACATAAAAAAGAAAC     10434
Pachon3       GCAGAGAGCAGAGCGTCAGGATGTGAGAACAGGATGCACTTAAGCACATAAAAAAGAAAC     10434
Pachon8       GCAGAGAGCAGAGCGTCAGGATGTGAGAACAGGATGCACTTAAGCACATAAAAAAGAAAC     10434
Pachon15      GCAGAGAGCAGAGCGTCAGGATGTGAGAACAGGATGCACTTAAGCACATAAAAAAGAAAC     10434
              ************************************************************

Rascon4       CTCAATTAACATTTCTTAATGTTAGAAGGCCCAGTGTCTGTAATTGGGCCCTTTGTAGTG     10444
```

```
Surface      CTCAATTAACATTTCTTAATGTTAGAAGGCCCAGTGTCTGTAATTGGGCCCTTTGTAGTG   10513
Rascon8      CTCAATTAACATTTCTTAATGTTAGAAGGCCCAGTGTCTGTAATTGGGCCCTTTGTAGTG   10506
Rascon2      CTCAATTAACATTTCTTAATGTTAGAAGGCCCAGTGTCTGTAATTGGGCCCTTTGTAGTG   10505
Rascon15     CTCAATTAACATTTCTTAATGTTAGAAGGCCCAGTGTCTGTAATTGGGCCCTTTGTAGTG   10494
Rascon13     CTCAATTAACATTTCTTAATGTTAGAAGGCCCAGTGTCTGTAATTGGGCCCTTTGTAGTG   10503
Rascon6      CTCAATTAACATTTCTTAATGTTAGAAGGCCCAGTGTCTGTAATTGGGCCCTTTGTAGTG   10508
Pachon14     CTCAATTAACATTTCTTAATGTTAGAAGGCCCAGTGTCTGTAATTGGGCCCTTTGTAGTG   10487
Pachon9      CTCAATTAACATTTCTTAATGTTAGAAGGCCCAGTGTCTGTAATTGGGCCCTTTGTAGTG   10496
Pachon17     CTCAATTAACATTTCTTAATGTTAGAAGGCCCAGTGTCTGTAATTGGGCCCTTTGTAGTG   10494
Pachon12     CTCAATTAACATTTCTTAATGTTAGAAGGCCCAGTGTCTGTAATTGGGCCCTTTGTAGTG   10476
Pachon11     CTCAATTAACATTTCTTAATGTTAGAAGGCCCAGTGTCTGTAATTGGGCCCTTTGTAGTG   10493
Pachon7      CTCAATTAACATTTCTTAATGTTAGAAGGCCCAGTGTCTGTAATTGGGCCCTTTGTAGTG   10494
Pachon3      CTCAATTAACATTTCTTAATGTTAGAAGGCCCAGTGTCTGTAATTGGGCCCTTTGTAGTG   10494
Pachon8      CTCAATTAACATTTCTTAATGTTAGAAGGCCCAGTGTCTGTAATTGGGCCCTTTGTAGTG   10494
Pachon15     CTCAATTAACATTTCTTAATGTTAGAAGGCCCAGTGTCTGTAATTGGGCCCTTTGTAGTG   10494
             ************************************************************

Rascon4      ATGGACAGCTGAGAGTGGGAATATAAGGCACTTTAGTCAACAGTTTTCCGCCAGCTGACA   10504
Surface      ATGGACAGCTGAGAGTGGGAATATAAGGCACTTTAGTCAACAGTTTTCCGCCAGCTGACA   10573
Rascon8      ATGGACAGCTGAGAGTGGGAATATAAGGCACTTTAGTCAACAGTTTTCCGCCAGCTGACA   10566
Rascon2      ATGGACAGCTGAGAGTGGGAATATAAGGCACTTTAGTCAACAGTTTTCCACCAGCTGACA   10565
Rascon15     ATGGACAGCTGAGAGTGGGAATATAAGGCACTTTAGTCAACAGTTTTCCGCCAGCTGACA   10554
Rascon13     ATGGACAGCTGAGAGTGGGAATATAAGGCACTTTAGTCAACAGTTTTCCGCCAGCTGACA   10563
Rascon6      ATGGACAGCTGAGAGTGGGAATATAAGGCACTTTAGTCAACAGTTTTCCGCCAGCTGACA   10568
Pachon14     ATGGACAGCTGAGAGTGGGAATATAAGGCACTTTAGTCAACAGTTTTCCGCCAGCTGACA   10547
Pachon9      ATGGACAGCTGAGAGTGGGAATATAAGGCACTTTAGTCAACAGTTTTCCGCCAGCTGACA   10556
Pachon17     ATGGACAGCTGAGAGTGGGAATATAAGGCACTTTAGTCAACAGTTTTCCGCCAGCTGACA   10554
Pachon12     ATGGACAGCTGAGAGTGGGAATATAAGGCACTTTAGTCAACAGTTTTCCGCCAGCTGACA   10536
Pachon11     ATGGACAGCTGAGAGTGGGAATATAAGGCACTTTAGTCAACAGTTTTCCGCCAGCTGACA   10553
Pachon7      ATGGACAGCTGAGAGTGGGAATATAAGGCACTTTAGTCAACAGTTTTCCGCCAGCTGACA   10554
Pachon3      ATGGACAGCTGAGAGTGGGAATATAAGGCACTTTAGTCAACAGTTTTCCGCCAGCTGACA   10554
Pachon8      ATGGACAGCTGAGAGTGGGAATATAAGGCACTTTAGTCAACAGTTTTCCGCCAGCTGACA   10554
Pachon15     ATGGACAGCTGAGAGTGGGAATATAAGGCACTTTAGTCAACAGTTTTCCGCCAGCTGACA   10554
             ************************************************* **********

Rascon4      CAGACTGAGGTTATAATTACGTCCATTAGAGCTGAAATGAGCCGAACTGTCCCTGAACAC   10564 SNP5
Surface      CAGACTGAGGTTATAATTACGTCCATTAGAGCTGAAATGAGCCGAACTGTCCCTGAACAC   10633 Also
Rascon8      CAGACTGAGGTTATAATTACGTCCATTAGAGCTGAAATGAGCCGAACTGTCCCTGAACAC   10626 fixed
Rascon2      CAGATTGAGGTTATAATTACGTCCATTAGAGCTGAAATGAGCCGAACTGTCCCTGAACAC   10625 in
Rascon15     CAGACTGAGGTTATAATTACGTCCATTAGAGCTGAAATGAGCCGAACTGTCCCTGAACAC   10614 Choy SF
Rascon13     CAGACTGAGGTTATAATTACGTCCATTAGAGCTGAAATGAGCCGAACTGTCCCTGAACAC   10623
Rascon6      CAGACTGAGGTTATAATTACGTCCATTAGAGCTGAAATGAGCCGAACTGTCCCTGAACAC   10628
Pachon14     CAGACTGAGGTTATAATTACGTCCATTAGAGCTGAAATGAGCCGAACTGTCACTGAACAC   10607
Pachon9      CAGACTGAGGTTATAATTACGTCCATTAGAGCTGAAATGAGCCGAACTGTCACTGAACAC   10616
Pachon17     CAGACTGAGGTTATAATTACGTCCATTAGAGCTGAAATGAGCCGAACTGTCACTGAACAC   10614
Pachon12     CAGACTGAGGTTATAATTACGTCCATTAGAGCTGAAATGAGCCGAACTGTCACTGAACAC   10596
Pachon11     CAGACTGAGGTTATAATTACGTCCATTAGAGCTGAAATGAGCCGAACTGTCACTGAACAC   10613
Pachon7      CAGACTGAGGTTATAATTACGTCCATTAGAGCTGAAATGAGCCGAACTGTCACTGAACAC   10614
Pachon3      CAGACTGAGGTTATAATTACGTCCATTAGAGCTGAAATGAGCCGAACTGTCACTGAACAC   10614
Pachon8      CAGACTGAGGTTATAATTACGTCCATTAGAGCTGAAATGAGCCGAACTGTCACTGAACAC   10614
Pachon15     CAGACTGAGGTTATAATTACGTCCATTAGAGCTGAAATGAGCCGAACTGTCACTGAACAC   10614
             **** ********************************************** ********

Rascon4      AGAAACTGAAGGTTTGATCCATTTACAAACTAGGATTGGATTGCGTGATCTGATCCGATT   10624
Surface      AGAAACTGAAGGTTTGATCCATTTACAAACTAGGATTGGATTGCGTGATCTGATCCGATT   10693
Rascon8      AGAAACTGAAGGTTTGATCCATTTACAAACTAGGATTGGATTGCGTGATCTGATCCGATT   10686
Rascon2      AGAAACTGAAGGTTTGATCCATTTACAAACTAGGATTGGATTGCGTGATCTGATCCGATT   10685
Rascon15     AGAAACTGAAGGTTTGATCCATTTACAAACTAGGATTGGATTGCGTGATCTGATCCGATT   10674
Rascon13     AGAAACTGAAGGTTTGATCCATTTACAAACTAGGATTGGATTGCGTGATCTGATCCGATT   10683
Rascon6      AGAAACTGAAGGTTTGATCCATTTACAAACTAGGATTGGATTGCGTGATCTGATCCGATT   10688
Pachon14     AGAAACTGAAGGTTTGATCCATTTACAAACTAGGATTGGATTGCGTGATCTGATCCGATT   10667
Pachon9      AGAAACTGAAGGTTTGATCCATTTACAAACTAGGATTGGATTGCGTGATCTGATCCGATT   10676
Pachon17     AGAAACTGAAGGTTTGATCCATTTACAAACTAGGATTGGATTGCGTGATCTGATCCGATT   10674
Pachon12     AGAAACTGAAGGTTTGATCCATTTACAAACTAGGATTGGATTGCGTGATCTGATCCGATT   10656
Pachon11     AGAAACTGAAGGTTTGATCCATTTACAAACTAGGATTGGATTGCGTGATCTGATCCGATT   10673
Pachon7      AGAAACTGAAGGTTTGATCCATTTACAAACTAGGATTGGATTGCGTGATCTGATCCGATT   10674
Pachon3      AGAAACTGAAGGTTTGATCCATTTACAAACTAGGATTGGATTGCGTGATCTGATCCGATT   10674
Pachon8      AGAAACTGAAGGTTTGATCCATTTACAAACTAGGATTGGATTGCGTGATCTGATCCGATT   10674
Pachon15     AGAAACTGAAGGTTTGATCCATTTACAAACTAGGATTGGATTGCGTGATCTGATCCGATT   10674
             ************************************************************

Rascon4      TCAGAATGCTTCTTTTCCTTTTGAACAACCCGTTTTTCAGATTTGATCCAATCCGATAAC   10684
Surface      TCAGAATGCTTCTTTTCCTTTTGAACAACCCGTTTTTCAGATTTGATCCAATCCGATAAC   10753
Rascon8      TCAGAATGCTTCTTTTCCTTTTGAACAACCCGTTTTTCAGATTTGATCCAATCCGATAAC   10746
Rascon2      TCAGAATGCTTCTTTTCCTTTTGAACAACCCGTTTTTCAGATTTGATCCAATCCGATAAC   10745
Rascon15     TCAGAATGCTTCTTTTCCTTTTGAACAACCCGTTTTTCAGATTTGATCCAATCCGATAAC   10734
Rascon13     TCAGAATGCTTCTTTTCCTTTTGAACAACCCGTTTTTCAGATTTGATCCAATCCGATAAC   10743
```

```
Rascon6      TCAGAATGCTTCTTTTCCTTTTGAACAACCCGTTTTTCAGATTTGATCCAATCCGATAAC    10748
Pachon14     TCAGAATGCTTCTTTTCCTTTTGAACAACCCGTTTTTCAGATTTGATCCAATCCGATAAC    10727
Pachon9      TCAGAATGCTTCTTTTCCTTTTGAACAACCCGTTTTTCAGATTTGATCCAATCCGATAAC    10736
Pachon17     TCAGAATGCTTCTTTTCCTTTTGAACAACCCGTTTTTCAGATTTGATCCAATCCGATAAC    10734
Pachon12     TCAGAATGCTTCTTTTCCTTTTGAACAACCCGTTTTTCAGATTTGATCCAATCCGATAAC    10716
Pachon11     TCAGAATGCTTCTTTTCCTTTTGAACAACCCGTTTTTCAGATTTGATCCAATCCGATAAC    10733
Pachon7      TCAGAATGCTTCTTTTCCTTTTGAACAACCCGTTTTTCAGATTTGATCCAATCCGATAAC    10734
Pachon3      TCAGAATGCTTCTTTTCCTTTTGAACAACCCGTTTTTCAGATTTGATCCAATCCGATAAC    10734
Pachon8      TCAGAATGCTTCTTTTCCTTTTGAACAACCCGTTTTTCAGATTTGATCCAATCCGATAAC    10734
Pachon15     TCAGAATGCTTCTTTTCCTTTTGAACAACCCGTTTTTCAGATTTGATCCAATCCGATAAC    10734
             ************************************************************

Rascon4      CAAAATCCGATCAGATTTTCTTTGAACAACTGGCCCCAGGAGACAGCAGCACCTTCGGCA    10744
Surface      CAAAATCCGATCAGATTTCCTTTGAACAACTGGCCCCAGGAGACAGCAGCACCTTCGGCA    10813
Rascon8      CAAAATCCGATCAGATTTCCTTTGAACAACTGGCCCCAGGAGACAGCAGCACCTTCGGCA    10806
Rascon2      CAAAATCCGATCAGATTTTCTTTGAACAACTGGCCCCAGGAGACAGCAGCACCTTCGGCA    10805
Rascon15     CAAAATCCGATCAGATTTCCTTTGAACAACTGGCCCCAGGAGACAGCAGCACCTTCGGCA    10794
Rascon13     CAAAATCCGATCAGATTTCCTTTGAACAACTGGCCCCAGGAGACAGCAGCACCTTCGGCA    10803
Rascon6      CAAAATCCGATCAGATTTCCTTTGAACAACTGGCCCCAGGAGACAGCAGCACCTTCGGCA    10808
Pachon14     CAAAATCCGATCAGATTTCCTTTGAACAACTGGCCCCAGGAGACAGCAGCACCTTCGGCA    10787
Pachon9      CAAAATCCGATCAGATTTCCTTTGAACAACTGGCCCCAGGAGACAGCAGCACCTTCGGCA    10796
Pachon17     CAAAATCCGATCAGATTTCCTTTGAACAACTGGCCCCAGGAGACAGCAGCACCTTCGGCA    10794
Pachon12     CAAAATCCGATCAGATTTCCTTTGAACAACTGGCCCCAGGAGACAGCAGCACCTTCGGCA    10776
Pachon11     CAAAATCCGATCAGATTTCCTTTGAACAACTGGCCCCAGGAGACAGCAGCACCTTCGGCA    10793
Pachon7      CAAAATCCGATCAGATTTCCTTTGAACAACTGGCCCCAGGAGACAGCAGCACCTTCGGCA    10794
Pachon3      CAAAATCCGATCAGATTTCCTTTGAACAACTGGCCCCAGGAGACAGCAGCACCTTCGGCA    10794
Pachon8      CAAAATCCGATCAGATTTCCTTTGAACAACTGGCCCCAGGAGACAGCAGCACCTTCGGCA    10794
Pachon15     CAAAATCCGATCAGATTTCCTTTGAACAACTGGCCCCAGGAGACAGCAGCACCTTCGGCA    10794
             ****************** *****************************************

Rascon4      CCAGGCGAGCATTAAGACCACTTCTGAACCCAATAGACGCGTAGAAAGAATTTATACAGC    10804
Surface      CCAGGCGAGCATTAAGACCACTTCTGAACCCAATAGACACGTAGAAAGAATTTATACAGC    10873
Rascon8      CCAGGCGAGCATTAAGACCACTTCTGAACCCAATAGACGCGTAGAAAGAATTTATACAGC    10866
Rascon2      CCAGACGAGCATTAAGACCACTTCTGAACCCAATAGACGCGTAGAAAGAATTTATACAGC    10865
Rascon15     CCAGGCGAGCATTAAGACCACTTCTGAACCCAATAGACGCGTAGAAAGAATTTATACAGC    10854
Rascon13     CCAGGCGAGCATTAAGACCACTTCTGAACCCAATAGACGCGTAGAAAGAATTTATACAGC    10863
Rascon6      CCAGGCGAGCATTAAGACCACTTCTGAACCCAATAGACACGTAGAAAGAATTTATACAGC    10868
Pachon14     CCAGGCGAGCATTAAGACCACTTCTGAACCCAATAGACACGTAGAAAGAATTTATACAGC    10847
Pachon9      CCAGGCGAGCATTAAGACCACTTCTGAACCCAATAGACACGTAGAAAGAATTTATACAGC    10856
Pachon17     CCAGGCGAGCATTAAGACCACTTCTGAACCCAATAGACACGTAGAAAGAATTTATACAGC    10854
Pachon12     CCAGGCGAGCATTAAGACCACTTCTGAACCCAATAGACACGTAGAAAGAATTTATACAGC    10836
Pachon11     CCAGGCGAGCATTAAGACCACTTCTGAACCCAATAGACACGTAGAAAGAATTTATACAGC    10853
Pachon7      CCAGGCGAGCATTAAGACCACTTCTGAACCCAATAGACACGTAGAAAGAATTTATACAGC    10854
Pachon3      CCAGGCGAGCATTAAGACCACTTCTGAACCCAATAGACACGTAGAAAGAATTTATACAGC    10854
Pachon8      CCAGGCGAGCATTAAGACCACTTCTGAACCCAATAGACACGTAGAAAGAATTTATACAGC    10854
Pachon15     CCAGGCGAGCATTAAGACCACTTCTGAACCCAATAGACACGTAGAAAGAATTTATACAGC    10854
             **** ********************************* *********************

Rascon4      TACAGCTATTATAACACAATCCTGTATTGATTCCAGGCAACAAC-AAAAAAATCGATAAA    10863
Surface      TACAGCTATTATAACACAATCCTGTATTGATTCCAGGGAACAAC-AAAAAAATCGATAAA    10932
Rascon8      TACAGCCATTATAACACAATCCTGTATTGATTCCAGGCAACAA-AAAAAAAATCGATAAA    10925
Rascon2      TACAGCTATTATAACACAATCCTGTATTGATTCCAGGCAACAACAAAAAAAATCGATAAA    10925
Rascon15     TACAGCTATTATAACACAATCCTGTATTGATTCCAGGCAACAAA-AAAAAAATCGATAAA    10913
Rascon13     TACAGCTATTATAACACAATCCTGTATTGATTCCAGGCAACAAA-AAAAAAATCGATAAA    10922
Rascon6      TACAGCTATTATAACACAATCCTGTATTGATTCCAGGCAACAAC-AAAAAAATCGATAAA    10927
Pachon14     TACAGCTATTATAACACAATCCTGTATTGATTCCAGGGAACAAC-AAAAAAATCGATAAA    10906
Pachon9      TACAGCTATTATAACACAATCCTGTATTGATTCCAGGGAACAAC-AAAAAAATCGATAAA    10915
Pachon17     TACAGCTATTATAACACAATCCTGTATTGATTCCAGGGAACAAC-AAAAAAATCGATAAA    10913
Pachon12     TACAGCTATTATAACACAATCCTGTATTGATTCCAGGGAACAAC-AAAAAAATCGATAAA    10895
Pachon11     TACAGCTATTATAACACAATCCTGTATTGATTCCAGGGAACAAC-AAAAAAATCGATAAA    10912
Pachon7      TACAGCTATTATAACACAATCCTGTATTGATTCCAGGGAACAAC-AAAAAAATCGATAAA    10913
Pachon3      TACAGCTATTATAACACAATCCTGTATTGATTCCAGGGAACAAC-AAAAAAATCGATAAA    10913
Pachon8      TACAGCTATTATAACACAATCCTGTATTGATTCCAGGGAACAAC-AAAAAAATCGATAAA    10913
Pachon15     TACAGCTATTATAACACAATCCTGTATTGATTCCAGGGAACAAC-AAAAAAATCGATAAA    10913
             ****** ****************************** *****  ***************

Rascon4      CGGAGGTGACTGTAATACAGGTTTGCAGAAGAAAAAACAGATCTGCTAGTGTTGTAGGAG    10923
Surface      CCGAGGTGACTGTAATACAGGTTTGCAGAAGAAAAAAACGATCTGCTAGTGTTGTAGGAG    10992
Rascon8      CGGAGGTGACTGTAATACAGGTTTGCAGAAGAAAAAACAGATCTGCTAGTGTTGTAGGAG    10985
Rascon2      CGGAGGTGACTGTAATACAGGTTTGCAGAAGAAAAAACAGATCTGCTAGTGTTGTAGGAG    10985
Rascon15     CGGAGGTGACTGTAATACAGGTTTGCAGAAGAAAAAACAGATCTGCTAGTGTTGTAGGAG    10973
Rascon13     CGGAGGTGACTGTAATACAGGTTTGCAGAAGAAAAAACAGATCTGCTAGTGTTGTAGGAG    10982
Rascon6      CGGAGGTGACTGTAATACAGGTTTGCAGAAGAAAAAACAGATCTGCTAGTGTTGTAGGAG    10987
Pachon14     CGGAGGTGACTGTAATACAGGTTTGCAGAAGAAAAAAACGATCTGCTAGTGTTGTAGGAG    10966
Pachon9      CGGAGGTGACTGTAATACAGGTTTGCAGAAGAAAAAAACGATCTGCTAGTGTTGTAGGAG    10975
Pachon17     CGGAGGTGACTGTAATACAGGTTTGCAGAAGAAAAAAACGATCTGCTAGTGTTGTAGGAG    10973
Pachon12     CGGAGGTGACTGTAATACAGGTTTGCAGAAGAAAAAAACGATCTGCTAGTGTTGTAGGAG    10955
```

```
Pachon11      CGGAGGTGACTGTAATACAGGTTTGCAGAAGAAAAAAACGATCTGCTAGTGTTGTAGGAG      10972
Pachon7       CGGAGGTGACTGTAATACAGGTTTGCAGAAGAAAAAAACGATCTGCTAGTGTTGTAGGAG      10973
Pachon3       CGGAGGTGACTGTAATACAGGTTTGCAGAAGAAAAAAACGATCTGCTAGTGTTGTAGGAG      10973
Pachon8       CGGAGGTGACTGTAATACAGGTTTGCAGAAGAAAAAAACGATCTGCTAGTGTTGTAGGAG      10973
Pachon15      CGGAGGTGACTGTAATACAGGTTTGCAGAAGAAAAAAACGATCTGCTAGTGTTGTAGGAG      10973
              * ***********************************  *********************

Rascon4       ATATCTGCGGTCATCTAGTGCTCATAATGTGACATTGCATCAACAATTCCAATGAAGTTC      10983
Surface       ATATCTGCTGTCATCTAGTGCTCATAATGTGACATTGCATCAACAATTCCAATGAAGTTC      11052
Rascon8       ATATCTGCTGTCATCTAGTGCTCATAATGTGACATTGCATCAACAATTCCAATGAAGTTC      11045
Rascon2       ATATCTGCTGTCATCTAGTGCTCATAATGTGACATTGCATCAACAATTCCAATGAAGTTC      11045
Rascon15      ATATCTGCTGTCATCTAGTGCTCATAATGTGACATTGCATCAACAATTCCAATGAAGTTC      11033
Rascon13      ATATCTGCTGTCATCTAGTGCTCATAATGTGACATTGCATCAACAATTCCAATGAAGTTC      11042
Rascon6       ATATCTGCTGTCATCTAGTGCTCATAATGTGACATTGCATCAACAATTCCAATGAAGTTC      11047
Pachon14      ATATCTGCTGTCATCTAGTGCTCATAATGTGACATTGCATCAACAATTCCAATGAAGTTC      11026
Pachon9       ATATCTGCTGTCATCTAGTGCTCATAATGTGACATTGCATCAACAATTCCAATGAAGTTC      11035
Pachon17      ATATCTGCTGTCATCTAGTGCTCATAATGTGACATTGCATCAACAATTCCAATGAAGTTC      11033
Pachon12      ATATCTGCTGTCATCTAGTGCTCATAATGTGACATTGCATCAACAATTCCAATGAAGTTC      11015
Pachon11      ATATCTGCTGTCATCTAGTGCTCATAATGTGACATTGCATCAACAATTCCAATGAAGTTC      11032
Pachon7       ATATCTGCTGTCATCTAGTGCTCATAATGTGACATTGCATCAACAATTCCAATGAAGTTC      11033
Pachon3       ATATCTGCTGTCATCTAGTGCTCATAATGTGACATTGCATCAACAATTCCAATGAAGTTC      11033
Pachon8       ATATCTGCTGTCATCTAGTGCTCATAATGTGACATTGCATCAACAATTCCAATGAAGTTC      11033
Pachon15      ATATCTGCTGTCATCTAGTGCTCATAATGTGACATTGCATCAACAATTCCAATGAAGTTC      11033
              ******** ***************************************************

Rascon4       AACAATTAAAAAAACCTGGAGAATCTACTTCAATAGATAGAATAGCAAAACAAAAAATAA      11043 Not in
Surface       AACAATTAAAAAAAACTGGAGAATCTACTTCAATAGATAGAATAGCAAAACACAAAATAA      11112 Choy SF
Rascon8       AACAATTAAAAAAAACTGGAGAATCTACTTCAATAGATAGAATAGCAAAACAAAAAATAA      11105
Rascon2       AACAATTAAAAAAAACTGGAGAATCTACTTCAATAGATAGAATAGCAAAACAAAAAATAA      11105
Rascon15      AACAATTAAAAAAAACTGGAGAATCTACTTCAATAGATAGAATAGCAAAACAAAAAATAA      11093
Rascon13      AACAATTAAAAAAAACTGGAGAATCTACTTCAATAGATAGAATAGCAAAACAAAAAATAA      11102
Rascon6       AACAATTAAAAAAAACTGGAGAATCTAATTCAATAGATAGAATAGCAAAACAAAAAATAA      11107
Pachon14      AACAATTAAAAAAAA-TGGAGAATCTACTTCAATAGATAGAATAGCAAAACACAAAATAA      11085
Pachon9       AACAATTAAAAAAAA-TGGAGAATCTACTTCAATAGATAGAATAGCAAAACACAAAATAA      11094
Pachon17      AACAATTAAAAAAAA-TGGAGAATCTACTTCAATAGATAGAATAGCAAAACACAAAATAA      11092
Pachon12      AACAATTAAAAAAAA-TGGAGAATCTACTTCAATAGATAGAATAGCAAAACACAAAATAA      11074
Pachon11      AACAATTAAAAAAAA-TGGAGAATCTACTTCAATAGATAGAATAGCAAAACACAAAATAA      11091
Pachon7       AACAATTAAAAAAAA-TGGAGAATCTACTTCAATAGATAGAATAGCAAAACACAAAATAA      11092
Pachon3       AACAATTAAAAAAAA-TGGAGAATCTACTTCAATAGATAGAATAGCAAAACACAAAATAA      11092
Pachon8       AACAATTAAAAAAAA-TGGAGAATCTACTTCAATAGATAGAATAGCAAAACACAAAATAA      11092
Pachon15      AACAATTAAAAAAAA-TGGAGAATCTACTTCAATAGATAGAATAGCAAAACACAAAATAA      11092
              **************  *********** ************************ *******

Rascon4       ACGCATGAACACAATTTTGTTCGTCAGTTTACACAATTTTGGCAGGTGATATATAACATT      11103 Not in
Surface       ACGCATGAACACAATTTTGTTCGTCAGTTTACACAATTTTGGCAGGTGATATATAACATT      11172 Choy SF
Rascon8       ACGCATGAACACAATTTTGTTCGTCAGTTTACACAATTTTGGCAGGTGATATATAACATT      11165 Mix A/G
Rascon2       ACGCAGGAACACAATTTTGTTCGTCAGTTTACACAATTTTGGCAGGTGATATATAACATT      11165
Rascon15      ACGCATGAACACAATTTTGTTCGTCAGTTTACACAATTTTGGCAGGTGATATATAACATT      11153
Rascon13      ACGCATGAACACAATTTTGTTCGTCAGTTTACACAATTTTGGCAGGTGATATATAACATT      11162
Rascon6       ACGCATGAACACAATTTTGTTCGTCAGTTTACACAATTTTGGCAGGTGATATATAACATT      11167
Pachon14      ACACATGAACACAATTTTGTTCGTCAGTTTACACAATTTTGGCAGGTGATATATAACATT      11145
Pachon9       ACACATGAACACAATTTTGTTCGTCAGTTTACACAATTTTGGCAGGTGATATATAACATT      11154
Pachon17      ACACATGAACACAATTTTGTTCGTCAGTTTACACAATTTTGGCAGGTGATATATAACATT      11152
Pachon12      ACACATGAACACAATTTTGTTCGTCAGTTTACACAATTTTGGCAGGTGATATATAACATT      11134
Pachon11      ACACATGAACACAATTTTGTTCGTCAGTTTACACAATTTTGGCAGGTGATATATAACATT      11151
Pachon7       ACACATGAACACAATTTTGTTCGTCAGTTTACACAATTTTGGCAGGTGATATATAACATT      11152
Pachon3       ACACATGAACACAATTTTGTTCGTCAGTTTACACAATTTTGGCAGGTGATATATAACATT      11152
Pachon8       ACACATGAACACAATTTTGTTCGTCAGTTTACACAATTTTGGCAGGTGATATATAACATT      11152
Pachon15      ACACATGAACACAATTTTGTTCGTCAGTTTACACAATTTTGGCAGGTGATATATAACATT      11152
              ** ** ******************************************************

Rascon4       ATAAAAGTCTATTATATCAAAAAATCACTTTTTTTCTTTTTTATTTTATTTCCTTTACTA      11163
Surface       ATAAAAGTCTATTATATCAAAAAAACACTTTTTTTCTTTTTTCTTTTATTTCCTTTACTA      11232
Rascon8       ATAAAAGTCTATTATATCAAAAAATCACTTTTTTTCTTTTTTATTTTATTTCCTTTACTA      11225
Rascon2       ATAAAAGTCTATTATATCAAAAAATCACTTTTTTTCTTTTTTATTTTATTTCTTTTACTA      11225
Rascon15      ATAAAAGTCTATTATATCAAAAAATCACTTTTTT--TTTTTTATTTTATTTCCTTTACTA      11211
Rascon13      ATAAAAGTCTATTATATCAAAAAATCACTTTTTTTCTTTTTTATTTTATTTCCTTTACTA      11222
Rascon6       ATAAAAGTCTATTATATCAAAAAATCACTTTTTTTCTTTTTTATTTTATTTCTTTTACTA      11227
Pachon14      ATAAAAGTCTATTATATCAAAAAAACACTTTTTTTCTTTTTTATTTTATTTCCTTTACTA      11205
Pachon9       ATAAAAGTCTATTATATCAAAAAAACACTTTTTTTCTTTTTTATTTTATTTCCTTTACTA      11214
Pachon17      ATAAAAGTCTATTATATCAAAAAAACACTTTTTTTCTTTTTTATTTTATTTCCTTTACTA      11212
Pachon12      ATAAAAGTCTATTATATCAAAAAAACACTTTTTTTCTTTTTTATTTTATTTCCTTTACTA      11194
Pachon11      ATAAAAGTCTATTATATCAAAAAAACACTTTTTTTCTTTTTTATTTTATTTCCTTTACTA      11211
Pachon7       ATAAAAGTCTATTATATCAAAAAAACACTTTTTTTCTTTTTTATTTTATTTCCTTTACTA      11212
Pachon3       ATAAAAGTCTATTATATCAAAAAAACACTTTTTTTCTTTTTTATTTTATTTCCTTTACTA      11212
Pachon8       ATAAAAGTCTATTATATCAAAAAAACACTTTTTTTCTTTTTTATTTTATTTCCTTTACTA      11212
Pachon15      ATAAAAGTCTATTATATCAAAAAAACACTTTTTTTCTTTTTTATTTTATTTCCTTTACTA      11212
```

```
                    ************************ *********  ****** ********* *******

Rascon4             TCATTCTACTTGTACAATGCAAT-AAAAAATGCATATCCTATAGAGCCAGGGTGAGCCAT     11222 SNP6
Surface             TCATTCTACTTGTACAATGCAAT-AAAAAATGCATATCCTATAGAGCCAGGGTGAGCCAT     11291 Also
Rascon8             TCATTCTACTTGTACAATGCAAT-AAAAAATGCATATCCTATAGAGCCAGGGTGAGCCAT     11284 fixed
Rascon2             TCATTCTACTTGTACAATGCAAT-AAAAAATGCATATCCTATAGAGCCAGGGTGAGCCAT     11284 in
Rascon15            TCATTCTACTTGTACAATGCAAT-AAAAAATGCATATCCTATAGAGCCAGGGTGAGCCAT     11270 Choy SF
Rascon13            TCATTCTACTTGTACAATGCAAT-AAAAAATGCATATCCTATAGAGCCAGGGTGAGCCAT     11281
Rascon6             TCATTCTACTTGTACAATGCAAT-AAAAAATGCATATCCTATAGAGCCAGGGTGAGCCAT     11286
Pachon14            TCATTCTACTTGTACAATGCAATAAAAAAATGCATATCCTATAGAGCCAGGGTGAGCCAT     11265
Pachon9             TCATTCTACTTGTACAATGCAATAAAAAAATGCATATCCTATAGAGCCAGGGTGAGCCAT     11274
Pachon17            TCATTCTACTTGTACAATGCAATAAAAAAATGCATATCCTATAGAGCCAGGGTGAGCCAT     11272
Pachon12            TCATTCTACTTGTACAATGCAATAAAAAAATGCATATCCTATAGAGCCAGGGTGAGCCAT     11254
Pachon11            TCATTCTACTTGTACAATGCAATAAAAAAATGCATATCCTATAGAGCCAGGGTGAGCCAT     11271
Pachon7             TCATTCTACTTGTACAATGCAATAAAAAAATGCATATCCTATAGAGCCAGGGTGAGCCAT     11272
Pachon3             TCATTCTACTTGTACAATGCAATAAAAAAATGCATATCCTATAGAGCCAGGGTGAGCCAT     11272
Pachon8             TCATTCTACTTGTACAATGCAATAAAAAAATGCATATCCTATAGAGCCAGGGTGAGCCAT     11272
Pachon15            TCATTCTACTTGTACAATGCAATAAAAAAATGCATATCCTATAGAGCCAGGGTGAGCCAT     11272
                    *********************** ************************************

Rascon4             TTTCCCTTGCTTTTATGACTATGAAAGTAGTGGTTTTAAAGTACTTGATTTAGCAAGGTT     11282
Surface             TTTCCCTTGCTTTTATGACTATGAAAGTAGTGGTTTTAAAGTACTTGATTTAGCAAGGTT     11351
Rascon8             TTTCCCTTGCTTTTATGACTATGAAAGTAGTGGTTTTAAAGTACTTGATTTAGCAAGGTT     11344
Rascon2             TTTCCCTTGCTTTTATGACTATGAAAGTAGTGGTTTTAAAGTACTTGATTTAGCAAGGTT     11344
Rascon15            TTTCCCTTGCTTTTATGACTATGAAAGTAGTGGTTTTAAAGTACTTGATTTAGCAAGGTT     11330
Rascon13            TTTCCCTTGCTTTTATGACTATGAAAGTAGTGGTTTTAAAGTACTTGATTTAGCAAGGTT     11341
Rascon6             TTTCCCTTGCTTTTATGACTATGAAAGTAGTGGTTTTAAAGTACTTGATTTAGCAAGGTT     11346
Pachon14            TTTCCCTTGCTTTTATGACTATGAAAGTAGTGGTTTTAAAGTACTTGATTTAGCAAGGTT     11325
Pachon9             TTTCCCTTGCTTTTATGACTATGAAAGTAGTGGTTTTAAAGTACTTGATTTAGCAAGGTT     11334
Pachon17            TTTCCCTTGCTTTTATGACTATGAAAGTAGTGGTTTTAAAGTACTTGATTTAGCAAGGTT     11332
Pachon12            TTTCCCTTGCTTTTATGACTATGAAAGTAGTGGTTTTAAAGTACTTGATTTAGCAAGGTT     11314
Pachon11            TTTCCCTTGCTTTTATGACTATGAAAGTAGTGGTTTTAAAGTACTTGATTTAGCAAGGTT     11331
Pachon7             TTTCCCTTGCTTTTATGACTATGAAAGTAGTGGTTTTAAAGTACTTGATTTAGCAAGGTT     11332
Pachon3             TTTCCCTTGCTTTTATGACTATGAAAGTAGTGGTTTTAAAGTACTTGATTTAGCAAGGTT     11332
Pachon8             TTTCCCTTGCTTTTATGACTATGAAAGTAGTGGTTTTAAAGTACTTGATTTAGCAAGGTT     11332
Pachon15            TTTCCCTTGCTTTTATGACTATGAAAGTAGTGGTTTTAAAGTACTTGATTTAGCAAGGTT     11332
                    ************************************************************

Rascon4             TGTGCAGATAAGTTTCAAAACAGGACTGCCTACTA-TTTTTTTTTCCTTTTTACAAACAA     11341
Surface             TGTGCAGATAAGTTTCAAAACAGGACTGCCTACTATTTTTTTTTTCCTTTTTACAAACAA     11411
Rascon8             TGTGCAGATAAGTTTCAAAACAGGACTGCCTACTA-TTTTTTTTTCCTTTTTACAAACAA     11403
Rascon2             TGTGCAGATAAGTTTCAAAACAGGACTGCCTACTA-TTTTTTTTTCCTTTTTACAAACAA     11403
Rascon15            TGTGCAGATAAGTTTCAAAACAGGACTGCCTACTATTTTTTTTTTCCTTTTTACAAACAA     11390
Rascon13            TGTGCAGATAAGTTTCAAAACAGGACTGCCTACTA-TTTTTTTTTCCTTTTTACAAACAA     11400
Rascon6             TGTGCAGATAAGTTTCAAAACAGGACTGCCTACTATTTTTTTTTTCCTTTTTACAAACAA     11406
Pachon14            TGTGCAGATAAGTTTCAAAACAGGACTGCCTACTA--TTTTTTTTCCTTTTTACAAACAA     11383
Pachon9             TGTGCAGATAAGTTTCAAAACAGGACTGCCTACTA--TTTTTTTTCCTTTTTACAAACAA     11392
Pachon17            TGTGCAGATAAGTTTCAAAACAGGACTGCCTACTA--TTTTTTTTCCTTTTTACAAACAA     11390
Pachon12            TGTGCAGATAAGTTTCAAAACAGGACTGCCTACTA--TTTTTTTTCCTTTTTACAAACAA     11372
Pachon11            TGTGCAGATAAGTTTCAAAACAGGACTGCCTACTA--TTTTTTTTCCTTTTTACAAACAA     11389
Pachon7             TGTGCAGATAAGTTTCAAAACAGGACTGCCTACTA--TTTTTTTTCCTTTTTACAAACAA     11390
Pachon3             TGTGCAGATAAGTTTCAAAACAGGACTGCCTACTA--TTTTTTTTCCTTTTTACAAACAA     11390
Pachon8             TGTGCAGATAAGTTTCAAAACAGGACTGCCTACTA--TTTTTTTTCCTTTTTACAAACAA     11390
Pachon15            TGTGCAGATAAGTTTCAAAACAGGACTGCCTACTA--TTTTTTTTCCTTTTTACAAACAA     11390
                    ***********************************  ***********************

Rascon4             ACAGGCGGATAACCTTGTTTACCATCTTTAATTATTCATAAAGCTGATGTTTCACTTTCA     11401
Surface             ACAGGCGGATAACCTTGTTTACCATCTTTAATTATTCATAAAGCTGATGTTTCACTTTCA     11471
Rascon8             ACAGGCGGATAACCTTGTTTACCATCTTTAATTATTCATAAAGCTGATGTTTCACTTTCA     11463
Rascon2             ACAGGCGGATAACCTTGTTTACCATCTTTAATTATTCATAAAGCTGATGTTTCACTTTCA     11463
Rascon15            ACAGGCGGATAACCTTGTTTACCATCTTTAATTATTCATAAAGCTGATGTTTCACTTTCA     11450
Rascon13            ACAGGCGGATAACCTTGTTTACCATCTTTAATTATTCATAAAGCTGATGTTTCACTTTCA     11460
Rascon6             ACAGGCGGATAACCTTGTTTACCATCTTTAATTATTCATAAAGCTGATGTTTCACTTTCA     11466
Pachon14            ACAGGCGGATAACCTTGTTTACCATCTTTAATTATTCATAAAGCTGATGTTTCACTTTCA     11443
Pachon9             ACAGGCGGATAACCTTGTTTACCATCTTTAATTATTCATAAAGCTGATGTTTCACTTTCA     11452
Pachon17            ACAGGCGGATAACCTTGTTTACCATCTTTAATTATTCATAAAGCTGATGTTTCACTTTCA     11450
Pachon12            ACAGGCGGATAACCTTGTTTACCATCTTTAATTATTCATAAAGCTGATGTTTCACTTTCA     11432
Pachon11            ACAGGCGGATAACCTTGTTTACCATCTTTAATTATTCATAAAGCTGATGTTTCACTTTCA     11449
Pachon7             ACAGGCGGATAACCTTGTTTACCATCTTTAATTATTCATAAAGCTGATGTTTCACTTTCA     11450
Pachon3             ACAGGCGGATAACCTTGTTTACCATCTTTAATTATTCATAAAGCTGATGTTTCACTTTCA     11450
Pachon8             ACAGGCGGATAACCTTGTTTACCATCTTTAATTATTCATAAAGCTGATGTTTCACTTTCA     11450
Pachon15            ACAGGCGGATAACCTTGTTTACCATCTTTAATTATTCATAAAGCTGATGTTTCACTTTCA     11450
                    ************************************************************

Rascon4             CTAAATCCAATTAATAAAAAACTGCTTTCTTAATGTTAATTACATTGGAAGCCTGTGTTT     11461
Surface             CTAAATCCAATTAAT-AAAAACTGCTTTCTTAATGTTAATTACATTGGAAGCCTGTGTTT     11530
Rascon8             CTAAATCCAATTAATAAAAAACTGCTTTCTTAATGTTAATTACATTGGAAGCCTGTGTTT     11523
```

```
Rascon2     CTAAATCCAATTAATAAAAAACTGCTTTCTTAATGTTAATTACATTGGAAGCCTGTGTTT     11523
Rascon15    CTAAATCCAATTAATAAAAAACGGCTTTCTTAATGTTAATTACATTGGAAGCCTGTGTTT     11510
Rascon13    CTAAATCCAATTAATAAAAAACTGCTTTCTTAATGTTAATTACATTGGAAGCCTGTGTTT     11520
Rascon6     CTAAATCCAATTAATAAAAAACTGCTTTCTTAATGTTAATTACATTGGAAGCCTGTGTTT     11526
Pachon14    CTAAATCCAATTAATAAAAAACTGCTTTCTTAATGTTAATTACATTGGAAGCCTGTGTTT     11503
Pachon9     CTAAATCCAATTAATAAAAAACTGCTTTCTTAATGTTAATTACATTGGAAGCCTGTGTTT     11512
Pachon17    CTAAATCCAATTAATAAAAAACTGCTTTCTTAATGTTAATTACATTGGAAGCCTGTGTTT     11510
Pachon12    CTAAATCCAATTAATAAAAAACTGCTTTCTTAATGTTAATTACATTGGAAGCCTGTGTTT     11492
Pachon11    CTAAATCCAATTAATAAAAAACTGCTTTCTTAATGTTAATTACATTGGAAGCCTGTGTTT     11509
Pachon7     CTAAATCCAATTAATAAAAAACTGCTTTCTTAATGTTAATTACATTGGAAGCCTGTGTTT     11510
Pachon3     CTAAATCCAATTAATAAAAAACTGCTTTCTTAATGTTAATTACATTGGAAGCCTGTGTTT     11510
Pachon8     CTAAATCCAATTAATAAAAAACTGCTTTCTTAATGTTAATTACATTGGAAGCCTGTGTTT     11510
Pachon15    CTAAATCCAATTAATAAAAAACTGCTTTCTTAATGTTAATTACATTGGAAGCCTGTGTTT     11510
            *************** ****** *************************************

Rascon4     AACATTTTTATCTTATTGGATATTTAAAACTGACCTTTTTATATCATAATCCGTACACTG     11521
Surface     AACATTTTTATCTTATTGGATATTTAAAACTGACCTTTTTATATCATAATCCGTACACTG     11590
Rascon8     AACATTTTTATCTTATTGGATATTTAAAACTGACCTTTTTATATCATAATCCGTACACTG     11583
Rascon2     AACATTTTTATCTTATTGGATATTTAAAACTGACCTTTTTATATCATAATCCGTACACTG     11583
Rascon15    AACATTTTTATCTTATTGGATATTTAAAACTGACCTTTTTATATCATAATCCGTACACTG     11570
Rascon13    AACATTTTTATCTTATTGGATATTTAAAACTGACCTTTTTATATCATAATCCGTACACTG     11580
Rascon6     AACATTTTTATCTTATTGGATATTTAAAACTGACCTTTTTATATCATAATCCGTACACTG     11586
Pachon14    AACATTTTTATCTTATTGGATATTTAAAACTGACCTTTTTATATCATAATCCGTACACTG     11563
Pachon9     AACATTTTTATCTTATTGGATATTTAAAACTGACCTTTTTATATCATAATCCGTACACTG     11572
Pachon17    AACATTTTTATCTTATTGGATATTTAAAACTGACCTTTTTATATCATAATCCGTACACTG     11570
Pachon12    AACATTTTTATCTTATTGGATATTTAAAACTGACCTTTTTATATCATAATCCGTACACTG     11552
Pachon11    AACATTTTTATCTTATTGGATATTTAAAACTGACCTTTTTATATCATAATCCGTACACTG     11569
Pachon7     AACATTTTTATCTTATTGGATATTTAAAACTGACCTTTTTATATCATAATCCGTACACTG     11570
Pachon3     AACATTTTTATCTTATTGGATATTTAAAACTGACCTTTTTATATCATAATCCGTACACTG     11570
Pachon8     AACATTTTTATCTTATTGGATATTTAAAACTGACCTTTTTATATCATAATCCGTACACTG     11570
Pachon15    AACATTTTTATCTTATTGGATATTTAAAACTGACCTTTTTATATCATAATCCGTACACTG     11570
            ************************************************************

Rascon4     TAATGAATGCAGGGTCTGAAAGGGTTACAGCAACTCATGGGTTAAATGTAATCAGTTAAT     11581
Surface     TAATGAAGGCAGGGTCTGAAAGGGTTACAGCAACTCATGGGTTAAATGTAATCAGTTAAT     11650
Rascon8     TAATGAATGCAGGGTCTGAAAGGGTTACAGCAACTCATGGGTTAAATGTAATCAGTTAAT     11643
Rascon2     TAATGAATGCAGGGTCTGAAAGGGTTACAGCAACTCAGGGGTTAAATGTAATCAGTTAAT     11643
Rascon15    TAATGAATGCAGGGTCTGAAAGGGTTACAGCAACTCATGGGTTAAATGTAATCAGTTAAT     11630
Rascon13    TAATGAATGCAGGGTCTGAAAGGGTTACAGCAACTCATGGGTTAAATGTAATCAGTTAAT     11640
Rascon6     TAATGAATGCAGGGTCTGAAAGGGTTACAGCAACTCATGGGTTAAATGTAATCAGTTAAT     11646
Pachon14    TAATGAATGCAGGGTCTGAAAGGGTTACAGCAACTCATGGGTTAAATGTAATCAGTTAAT     11623
Pachon9     TAATGAATGCAGGGTCTGAAAGGGTTACAGCAACTCATGGGTTAAATGTAATCAGTTAAT     11632
Pachon17    TAATGAATGCAGGGTCTGAAAGGGTTACAGCAACTCATGGGTTAAATGTAATCAGTTAAT     11630
Pachon12    TAATGAATGCAGGGTCTGAAAGGGTTACAGCAACTCATGGGTTAAATGTAATCAGTTAAT     11612
Pachon11    TAATGAATGCAGGGTCTGAAAGGGTTACAGCAACTCATGGGTTAAATGTAATCAGTTAAT     11629
Pachon7     TAATGAATGCAGGGTCTGAAAGGGTTACAGCAACTCATGGGTTAAATGTAATCAGTTAAT     11630
Pachon3     TAATGAATGCAGGGTCTGAAAGGGTTACAGCAACTCATGGGTTAAATGTAATCAGTTAAT     11630
Pachon8     TAATGAATGCAGGGTCTGAAAGGGTTACAGCAACTCATGGGTTAAATGTAATCAGTTAAT     11630
Pachon15    TAATGAATGCAGGGTCTGAAAGGGTTACAGCAACTCATGGGTTAAATGTAATCAGTTAAT     11630
            ******* **************************** ***********************

Rascon4     ACCACTCCTCTTGTGACTTGTGTGCTCAGTCGCTGTGTTGTGTGTACAGACAGCACCTCT     11641
Surface     ACCACTCCTCTTGTGACTTGTGTGCTCAGACGCTGTGTTGTGTGTACAGACAGCACCTCT     11710
Rascon8     ACCACTCCTCTTGTGACTTGTGTGCTCAGTCGCTGTGTTGTGTGTACAGACAGCACCTCT     11703
Rascon2     ACCA-----CTTGTGACTTGTGTGCTCAGTCGCTGTGTTGTGTGTACAGACAGCACCTCT     11698
Rascon15    ACCACTCCTCTTGTGACTTGTGTGCTCAGTCGCTGTGTTGTGTGTACAGACAGCACCTCT     11690
Rascon13    ACCACTCCTCTTGTGACTTGTGTGCTCAGTCGCTGTGTTGTGTGTACAGACAGCACCTCT     11700
Rascon6     ACCA-----CTTGTGACTTGTGTGCTCAGTCGCTGTGTTGTGTGTACAGACAGCACCTCT     11701
Pachon14    ACCACTCCTCTTGTGACTTGTGTGCTCAGTCGCTGTGTTGTGTGTACAGACAGCACCTCT     11683
Pachon9     ACCACTCCTCTTGTGACTTGTGTGCTCAGTCGCTGTGTTGTGTGTACAGACAGCACCTCT     11692
Pachon17    ACCACTCCTCTTGTGACTTGTGTGCTCAGTCGCTGTGTTGTGTGTACAGACAGCACCTCT     11690
Pachon12    ACCACTCCTCTTGTGACTTGTGTGCTCAGTCGCTGTGTTGTGTGTACAGACAGCACCTCT     11672
Pachon11    ACCACTCCTCTTGTGACTTGTGTGCTCAGTCGCTGTGTTGTGTGTACAGACAGCACCTCT     11689
Pachon7     ACCACTCCTCTTGTGACTTGTGTGCTCAGTCGCTGTGTTGTGTGTACAGACAGCACCTCT     11690
Pachon3     ACCACTCCTCTTGTGACTTGTGTGCTCAGTCGCTGTGTTGTGTGTACAGACAGCACCTCT     11690
Pachon8     ACCACTCCTCTTGTGACTTGTGTGCTCAGTCGCTGTGTTGTGTGTACAGACAGCACCTCT     11690
Pachon15    ACCACTCCTCTTGTGACTTGTGTGCTCAGTCGCTGTGTTGTGTGTACAGACAGCACCTCT     11690
            ****     ******************** ******************************

Rascon4     TACTCACACTGCAGTTTATTGAAATAGTTATAGCTTTATTGAATATGTTGGATGAAACAT     11701 Grp
Surface     TACTCACACTGCAGTTTATTGAAATAGTTATAGCTTTATTGAATATGTTGGATGAAACAT     11770 3'-UTR
Rascon8     TACTCACACTGCAGTTTATTGAAATAGTTATAGCTTTATTGAATATGTTGGATGAAACAT     11763 end
Rascon2     TACTCACACTGCAGTTTATTGAAATAGTTATAGCTTTATTGAATATGTTGGATGAAACAT     11758
Rascon15    TACTCACACTGCAGTTTATTGAAATAGTTATAGCTTTATTGAATATGTTGGATGAAACAT     11750
Rascon13    TACTCACACTGCAGTTTATTGAAATAGTTATAGCTTTATTGAATATGTTGGATGAAACAT     11760
Rascon6     TACTCACACTGCAGTTTATTGAAATAGTTATAGCTTTATTGAATATGTTGGATGAAACAT     11761
Pachon14    TACTCACACTGCAGTTTATTGAAATAGTTATAGCTTTATTGAATATGTTGGATGAAACAT     11743
```

```
Pachon9          TACTCACACTGCAGTTTATTGAAATAGTTATAGCTTTATTGAATATGTTGGATGAAACAT     11752
Pachon17         TACTCACACTGCAGTTTATTGAAATAGTTATAGCTTTATTGAATATGTTGGATGAAACAT     11750
Pachon12         TACTCACACTGCAGTTTATTGAAATAGTTATAGCTTTATTGAATATGTTGGATGAAACAT     11732
Pachon11         TACTCACACTGCAGTTTATTGAAATAGTTATAGCTTTATTGAATATGTTGGATGAAACAT     11749
Pachon7          TACTCACACTGCAGTTTATTGAAATAGTTATAGCTTTATTGAATATGTTGGATGAAACAT     11750
Pachon3          TACTCACACTGCAGTTTATTGAAATAGTTATAGCTTTATTGAATATGTTGGATGAAACAT     11750
Pachon8          TACTCACACTGCAGTTTATTGAAATAGTTATAGCTTTATTGAATATGTTGGATGAAACAT     11750
Pachon15         TACTCACACTGCAGTTTATTGAAATAGTTATAGCTTTATTGAATATGTTGGATGAAACAT     11750
                 ************************************************************

Rascon4          CACCACATTAATAATTTAAAGGTAGATAAATTAATGTGTAGGCCTATAAAACAAAAGTAA     11761
Surface          CACCACATTAATAATTTAAAAGTAGATAAATTAATGTGTAGGCCTATAAAACAAAAGTAA     11830
Rascon8          CACCACATTAATAATTTAAAGGTAGATAAATTAATGTGTAGGCCTATAAAACAAAAGTAA     11823
Rascon2          CACCACATTAATAATTTAAAGGTAGATAAATTAATGTGTAGGCCTATAAAACAAAAGTAA     11818
Rascon15         CACCACATTAATAATTTAAAGGTAGATAAATTAATGTGTAGGCCTATAAAACAAAAGTAA     11810
Rascon13         CACCACATTAATAATTTAAAGGTAGATAAATTAATGTGTAGGCCTATAAAACAAAAGTAA     11820
Rascon6          CACCACATTAATAATTTAAAGGTAGATAAATTAATGTGTAGGCCTATAAAACAAAAGTAA     11821
Pachon14         CACCACATTAATAATTTAAAGGTAGATAAATTAATGTGTAGGCCTATAAAACAAAAGTAA     11803
Pachon9          CACCACATTAATAATTTAAAGGTAGATAAATTAATGTGTAGGCCTATAAAACAAAAGTAA     11812
Pachon17         CACCACATTAATAATTTAAAGGTAGATAAATTAATGTGTAGGCCTATAAAACAAAAGTAA     11810
Pachon12         CACCACATTAATAATTTAAAGGTAGATAAATTAATGTGTAGGCCTATAAAACAAAAGTAA     11792
Pachon11         CACCACATTAATAATTTAAAGGTAGATAAATTAATGTGTAGGCCTATAAAACAAAAGTAA     11809
Pachon7          CACCACATTAATAATTTAAAGGTAGATAAATTAATGTGTAGGCCTATAAAACAAAAGTAA     11810
Pachon3          CACCACATTAATAATTTAAAGGTAGATAAATTAATGTGTAGGCCTATAAAACAAAAGTAA     11810
Pachon8          CACCACATTAATAATTTAAAGGTAGATAAATTAATGTGTAGGCCTATAAAACAAAAGTAA     11810
Pachon15         CACCACATTAATAATTTAAAGGTAGATAAATTAATGTGTAGGCCTATAAAACAAAAGTAA     11810
                 ******************** ***************************************

Rascon4          ATGTGAATTCATGTCCTGGGTGTTTTAGATTTTACATACAAAAATATAACTTACATTTA-     11820
Surface          ATGTGAATTCATGTCCTGGGTGTTTTAGATTTTACATACAAAAATATAACTTACATTTAT     11890
Rascon8          ATGTGAATTCATGTCCTGGGTGTTTTAGATTTTACATACAAAAATATAACTTACATTTA-     11882
Rascon2          ATGTGAATTCATGTCCTGGGTGTTTTAGATTTTACATACAAAAATATAACTTACATTTA-     11877
Rascon15         ATGTGAATTCATGTCCTGGGTGTTTTAGATTTTACATACAAAAATATAACTTACATTTAT     11870
Rascon13         ATGTGAATTCATGTCCTGGGTGTTTTAGATTTTACATACAAAAATATAACTTACATTTAT     11880
Rascon6          ATGTGAATTCATGTCCTGGGTGTTTTAGATTTTACATACAAAAATATAACTTACATTTAT     11881
Pachon14         ATGTGAATTCATGTCCTGGGTGTTTTAGATTTTACATACAAAAATATAACTTACATTTAT     11863
Pachon9          ATGTGAATTCATGTCCTGGGTGTTTTAGATTTTACATACAAAAATATAACTTACATTTAT     11872
Pachon17         ATGTGAATTCATGTCCTGGGTGTTTTAGATTTTACATACAAAAATATAACTTACATTTAT     11870
Pachon12         ATGTGAATTCATGTCCTGGGTGTTTTAGATTTTACATACAAAAATATAACTTACATTTAT     11852
Pachon11         ATGTGAATTCATGTCCTGGGTGTTTTAGATTTTACATACAAAAATATAACTTACATTTAT     11869
Pachon7          ATGTGAATTCATGTCCTGGGTGTTTTAGATTTTACATACAAAAATATAACTTACATTTAT     11870
Pachon3          ATGTGAATTCATGTCCTGGGTGTTTTAGATTTTACATACAAAAATATAACTTACATTTAT     11870
Pachon8          ATGTGAATTCATGTCCTGGGTGTTTTAGATTTTACATACAAAAATATAACTTACATTTAT     11870
Pachon15         ATGTGAATTCATGTCCTGGGTGTTTTAGATTTTACATACAAAAATATAACTTACATTTAT     11870
                 ***********************************************************

Rascon4          TCGTCAGTAGATGCAGTGCAGAATGAAAAGACTATAATGCA--------TATTTTTCGAA     11872
Surface          TCGTCAGTAGATGCAGTGCAGAATGAAAAGACTATAATGCATATTACTTTATTTTTCGAA     11950
Rascon8          TCGTCAGTAGATGCAGTGCAGAATGAAAAGACTATAATGCA--------TATTTTTCGAA     11934
Rascon2          TCGTCAGTAGATGCAGTGCAGAATGAAAAGACTATAATGCA--------TATTTTTCGAA     11929
Rascon15         TCGTCAGTAGATGCAGTGCAGAATGAAAAGACTATAATGCATATTACTTTATTTTTCGAA     11930
Rascon13         TCGTCAGTAGATGCAGTGCAGAATGAAAAGACTATAATGCATATTACTTTATTTTTCGAA     11940
Rascon6          TCGTCAGTAGATGCAGTGCAGAATGAAAAGACTATAATGCATATTACTTTATTTTTCGAA     11941
Pachon14         TCGTCAGTAGATGCAGTGCAGAATGAAAAGACTATAATGCATATTACTTTATTTTTCGAA     11923
Pachon9          TCGTCAGTAGATGCAGTGCAGAATGAAAAGACTATAATGCATATTACTTTATTTTTCGAA     11932
Pachon17         TCGTCAGTAGATGCAGTGCAGAATGAAAAGACTATAATGCATATTACTTTATTTTTCGAA     11930
Pachon12         TCGTCAGTAGATGCAGTGCAGAATGAAAAGACTATAATGCATATTACTTTATTTTTCGAA     11912
Pachon11         TCGTCAGTAGATGCAGTGCAGAATGAAAAGACTATAATGCATATTACTTTATTTTTCGAA     11929
Pachon7          TCGTCAGTAGATGCAGTGCAGAATGAAAAGACTATAATGCATATTACTTTATTTTTCGAA     11930
Pachon3          TCGTCAGTAGATGCAGTGCAGAATGAAAAGACTATAATGCATATTACTTTATTTTTCGAA     11930
Pachon8          TCGTCAGTAGATGCAGTGCAGAATGAAAAGACTATAATGCATATTACTTTATTTTTCGAA     11930
Pachon15         TCGTCAGTAGATGCAGTGCAGAATGAAAAGACTATAATGCATATTACTTTATTTTTCGAA     11930
                 *****************************************        ***********

Rascon4          CAGAGACAGCTGTGAGGCAGTGTTATCACGTTTCTGCTGCTTCCAACCACAAAGTCTTTG     11932
Surface          CAGAGACAGCTGTGAGGCAGTGTTATCACGTTTCTGCTGCTTCCAACCACAAAGTCTTTG     12010
Rascon8          CAGAGACAGCTGTGAGGCAGTGTTATCACGTTTCTGCTGCTTCCAACCACAAAGTCTTTG     11994
Rascon2          CAGAGACAGCTGTGAGGCAGTGTTATCACATTTCTGCTGCTTCCAACCACAAAGTCTTTG     11989
Rascon15         CAGAGACAGCTGTGAGGCAGTGTTATCACGTTTCTGCTGCTTCCAACCACAAAGTCTTTG     11990
Rascon13         CAGAGACAGCTGTGAGGCAGTGTTATCACGTTTCTGCTGCTTCCAACCACAAAGTCTTTG     12000
Rascon6          CAGAGACAGCTGTGAGGCAGTGTTATCACGTTTCTGCTGCTTCCAACCACAAAGTCTTTG     12001
Pachon14         CAGAGACAGCTGTGAGGCAGTGTTATCACGTTTCTGCTGCTTCCAACCACAAAGTCTTTG     11983
Pachon9          CAGAGACAGCTGTGAGGCAGTGTTATCACGTTTCTGCTGCTTCCAACCACAAAGTCTTTG     11992
Pachon17         CAGAGACAGCTGTGAGGCAGTGTTATCACGTTTCTGCTGCTTCCAACCACAAAGTCTTTG     11990
Pachon12         CAGAGACAGCTGTGAGGCAGTGTTATCACGTTTCTGCTGCTTCCAACCACAAAGTCTTTG     11972
Pachon11         CAGAGACAGCTGTGAGGCAGTGTTATCACGTTTCTGCTGCTTCCAACCACAAAGTCTTTG     11989
Pachon7          CAGAGACAGCTGTGAGGCAGTGTTATCACGTTTCTGCTGCTTCCAACCACAAAGTCTTTG     11990
```

```
Pachon3      CAGAGACAGCTGTGAGGCAGTGTTATCACGTTTCTGCTGCTTCCAACCACAAAGTCTTTG      11990
Pachon8      CAGAGACAGCTGTGAGGCAGTGTTATCACGTTTCTGCTGCTTCCAACCACAAAGTCTTTG      11990
Pachon15     CAGAGACAGCTGTGAGGCAGTGTTATCACGTTTCTGCTGCTTCCAACCACAAAGTCTTTG      11990
             ***************************** ******************************

Rascon4      AAAAATCAGGCCACAGTGCTCATCTTCAGCTTGTTTTGGAGTCCCTCATGTTTAAAGCCA      11992 Grp
Surface      ATAAATCAGGCCACAGTGCTCATCTTCAGCTTGTTTTGGAGTCCCTCATGTTTAAAGCCA      12070 3'-UTR
Rascon8      AAAAATCAGGCCACAGTGCTCATCTTCAGCTTGTTTTGGAGTCCCTCATGTTTAAAGCCA      12054 start
Rascon2      AAAAATCAGGCCACAGTGCTCATCTTCAGCTTGTTTTGGAGTCCCTCATGTTTAAAGCCA      12049 exon 3
Rascon15     ATAAATCAGGCCACAGTGCTCATCTTCAGCTTGTTTTGGAGTCCCTCATGTTTAAAGCCA      12050 end
Rascon13     ATAAATCAGGCCACAGTGCTCATCTTCAGCTTGTTTTGGAGTCCCTCATGTTTAAAGCCA      12060
Rascon6      ATAAATCAGGCCACAGTGCTCATCTTCAGCTTGTTTTGGAGTCCCTCATGTTTAAAGCCA      12061
Pachon14     ATAAATCAGGCCACAGTGCTCATCTTCAGCTTGTTTTGGAGTCCCTCATGTTTAAAGCCA      12043
Pachon9      ATAAATCAGGCCACAGTGCTCATCTTCAGCTTGTTTTGGAGTCCCTCATGTTTAAAGCCA      12052
Pachon17     ATAAATCAGGCCACAGTGCTCATCTTCAGCTTGTTTTGGAGTCCCTCATGTTTAAAGCCA      12050
Pachon12     ATAAATCAGGCCACAGTGCTCATCTTCAGCTTGTTTTGGAGTCCCTCATGTTTAAAGCCA      12032
Pachon11     ATAAATCAGGCCACAGTGCTCATCTTCAGCTTGTTTTGGAGTCCCTCATGTTTAAAGCCA      12049
Pachon7      ATAAATCAGGCCACAGTGCTCATCTTCAGCTTGTTTTGGAGTCCCTCATGTTTAAAGCCA      12050
Pachon3      ATAAATCAGGCCACAGTGCTCATCTTCAGCTTGTTTTGGAGTCCCTCATGTTTAAAGCCA      12050
Pachon8      ATAAATCAGGCCACAGTGCTCATCTTCAGCTTGTTTTGGAGTCCCTCATGTTTAAAGCCA      12050
Pachon15     ATAAATCAGGCCACAGTGCTCATCTTCAGCTTGTTTTGGAGTCCCTCATGTTTAAAGCCA      12050
             * **********************************************************

Rascon4      GTAGCAAAATCTTTGCTGCCTGTATGAAAGAATAGTGACAGAGAGAGCATTAGTCATTAT      12052 Grp
Surface      GTAGCAAAATCTTTGCTGCCTGTATGAAAGAATAGTGACAGAGAGAGCATTAGTCATTAT      12130 exon 3
Rascon8      GTAGCAAAATCTTTGCTGCCTGTATGAAAGAATAGTGACAGAGAGAGCATTAGTCATTAT      12114 start
Rascon2      GTAGCAAAATCTTTGCTGCCTGTATGAAAGAATAGTGACAGAGAGAGCATTAGTCATTAT      12109 Intron2
Rascon15     GTAGCAAAATCTTTGCTGCCTGTATGAAAGAATAGTGACAGAGAGAGCATTAGTCATTAT      12110 end
Rascon13     GTAGCAAAATCTTTGCTGCCTGTATGAAAGAATAGTGACAGAGAGAGCATTAGTCATTAT      12120
Rascon6      GTAGCAAAATCTTTGCTGCCTGTATGAAAGAATAGTGACAGAGAGAGCATTAGTCATTAT      12121
Pachon14     GTAGCAAAATCTTTGCTGCCTGTATGAAAGAATAGTGACAGAGAGAGCATTAGTCATTAT      12103
Pachon9      GTAGCAAAATCTTTGCTGCCTGTATGAAAGAATAGTGACAGAGAGAGCATTAGTCATTAT      12112
Pachon17     GTAGCAAAATCTTTGCTGCCTGTATGAAAGAATAGTGACAGAGAGAGCATTAGTCATTAT      12110
Pachon12     GTAGCAAAATCTTTGCTGCCTGTATGAAAGAATAGTGACAGAGAGAGCATTAGTCATTAT      12092
Pachon11     GTAGCAAAATCTTTGCTGCCTGTATGAAAGAATAGTGACAGAGAGAGCATTAGTCATTAT      12109
Pachon7      GTAGCAAAATCTTTGCTGCCTGTATGAAAGAATAGTGACAGAGAGAGCATTAGTCATTAT      12110
Pachon3      GTAGCAAAATCTTTGCTGCCTGTATGAAAGAATAGTGACAGAGAGAGCATTAGTCATTAT      12110
Pachon8      GTAGCAAAATCTTTGCTGCCTGTATGAAAGAATAGTGACAGAGAGAGCATTAGTCATTAT      12110
Pachon15     GTAGCAAAATCTTTGCTGCCTGTATGAAAGAATAGTGACAGAGAGAGCATTAGTCATTAT      12110
             ************************************************************

Rascon4      TCAATATTTTACCCAGTAATTAAGCAGATCTGCTTAATTAACAAGGTGCAATCCCATTTC      12112
Surface      TCAATATTTTACCCAGTAATTAAGCAGATCTGCTTAATTAACAAGGTGCAATCCCATTTC      12190
Rascon8      TCAATATTTTACCCAGTAATTAAGCAGATCTGCTTAATTAACAAGGTGCAATCCCATTTC      12174
Rascon2      TCAATATTTTACCCAGTAATTAAGCAGATCTGCTTAATTAACAAGGTGCAATCCCATTTC      12169
Rascon15     TCAATATTTTACCCAGTAATTAAGCAGATCTGCTTAATTAACAAGGTGCAATCCCATTTC      12170
Rascon13     TCAATATTTTACCCAGTAATTAAGCAGATCTGCTTAATTAACAAGGTGCAATCCCATTTC      12180
Rascon6      TCAATATTTTACCCAGTAATTAAGCAGATCTGCTTAATTAACAAGGTGCAATCCCATTTC      12181
Pachon14     TCAATATTTTACCCAGTAATTAAGCAGATCTGCTTAATTAACAAGGTGCAATCCCATTTC      12163
Pachon9      TCAATATTTTACCCAGTAATTAAGCAGATCTGCTTAATTAACAAGGTGCAATCCCATTTC      12172
Pachon17     TCAATATTTTACCCAGTAATTAAGCAGATCTGCTTAATTAACAAGGTGCAATCCCATTTC      12170
Pachon12     TCAATATTTTACCCAGTAATTAAGCAGATCTGCTTAATTAACAAGGTGCAATCCCATTTC      12152
Pachon11     TCAATATTTTACCCAGTAATTAAGCAGATCTGCTTAATTAACAAGGTGCAATCCCATTTC      12169
Pachon7      TCAATATTTTACCCAGTAATTAAGCAGATCTGCTTAATTAACAAGGTGCAATCCCATTTC      12170
Pachon3      TCAATATTTTACCCAGTAATTAAGCAGATCTGCTTAATTAACAAGGTGCAATCCCATTTC      12170
Pachon8      TCAATATTTTACCCAGTAATTAAGCAGATCTGCTTAATTAACAAGGTGCAATCCCATTTC      12170
Pachon15     TCAATATTTTACCCAGTAATTAAGCAGATCTGCTTAATTAACAAGGTGCAATCCCATTTC      12170
             ************************************************************

Rascon4      ATGTGGCATATCAGTGTAAAACTGGTAATGCATCATAATGCAAGGCCTTTGTAATGTTTC      12172
Surface      ATGTGGCATATCAGTGTAAAACTGGTAATGCATCATAATGCAAGGCCTTTGTAATGTTTC      12250
Rascon8      ATGTGGCATATCAGTGTAAAACTGGTAATGCATCATAATGCAAGGCCTTTGTAATGTTTC      12234
Rascon2      ATGTGGCATATCAGTGTAAAACTGGTAATGCATCATAATGCAAGGCCTTTGTAATGTTTC      12229
Rascon15     ATGTGGCATATCAGTGTAAAACTGGTAATGCATCATAATGCAAGGCCTTTGTAATGTTTC      12230
Rascon13     ATGTGGCATATCAGTGTAAAACTGGTAATGCATCATAATGCAAGGCCTTTGTAATGTTTC      12240
Rascon6      ATGTGGCATATCAGTGTAAAACTGGTAATGCATCATAATGCAAGGCCTTTGTAATGTTTC      12241
Pachon14     ATGTGGCATATCAGTGTAAAACTGGTAATGCATCATAATGCAAGGCCTTTGTAATGTTTC      12223
Pachon9      ATGTGGCATATCAGTGTAAAACTGGTAATGCATCATAATGCAAGGCCTTTGTAATGTTTC      12232
Pachon17     ATGTGGCATATCAGTGTAAAACTGGTAATGCATCATAATGCAAGGCCTTTGTAATGTTTC      12230
Pachon12     ATGTGGCATATCAGTGTAAAACTGGTAATGCATCATAATGCAAGGCCTTTGTAATGTTTC      12212
Pachon11     ATGTGGCATATCAGTGTAAAACTGGTAATGCATCATAATGCAAGGCCTTTGTAATGTTTC      12229
Pachon7      ATGTGGCATATCAGTGTAAAACTGGTAATGCATCATAATGCAAGGCCTTTGTAATGTTTC      12230
Pachon3      ATGTGGCATATCAGTGTAAAACTGGTAATGCATCATAATGCAAGGCCTTTGTAATGTTTC      12230
Pachon8      ATGTGGCATATCAGTGTAAAACTGGTAATGCATCATAATGCAAGGCCTTTGTAATGTTTC      12230
Pachon15     ATGTGGCATATCAGTGTAAAACTGGTAATGCATCATAATGCAAGGCCTTTGTAATGTTTC      12230
             ************************************************************
```

```
Rascon4      TGCCTGGTCTGCCAAGACAAAATGTACTTTGCTTTTTGAATTAAAACATTTAAACCACTG    12232 Grp
Surface      TGCCTGGTCTGCCAAGATAAAATGTACTTTGCTTTTTGAATTAAAACATTTAAACCACTG    12310 Intron2
Rascon8      TGCCTGGTCTGCCAAGATAAAATGTACTTTGCTTTTTGAATTAAAACATTTAAACCACTG    12294
Rascon2      TGCCTGGTCTGCCAAGACAAAATGTACTTTGCTTTTTGAATTAAAACATTTAAACCACTG    12289
Rascon15     TGCCTGGTCTGCCAAGATAAAATGTACTTTGCTTTTTGAATTAAAACATTTAAACCACTG    12290
Rascon13     TGCCTGGTCTGCCAAGACAAAATGTACTTTGCTTTTTGAATTAAAACATTTAAACCACTG    12300
Rascon6      TGCCTGGTCTGCCAAGATAAAATGTACTTTGCTTTTTGAATTAAAACATTTAAACCACTG    12301
Pachon14     TGCCTGGTCTGCCAAGATAAAATGTACTTTGCTTTTTGAATTAAAACATTTAAACCACTG    12283
Pachon9      TGCCTGGTCTGCCAAGATAAAATGTACTTTGCTTTTTGAATTAAAACATTTAAACCACTG    12292
Pachon17     TGCCTGGTCTGCCAAGATAAAATGTACTTTGCTTTTTGAATTAAAACATTTAAACCACTG    12290
Pachon12     TGCCTGGTCTGCCAAGATAAAATGTACTTTGCTTTTTGAATTAAAACATTTAAACCACTG    12272
Pachon11     TGCCTGGTCTGCCAAGATAAAATGTACTTTGCTTTTTGAATTAAAACATTTAAACCACTG    12289
Pachon7      TGCCTGGTCTGCCAAGATAAAATGTACTTTGCTTTTTGAATTAAAACATTTAAACCACTG    12290
Pachon3      TGCCTGGTCTGCCAAGATAAAATGTACTTTGCTTTTTGAATTAAAACATTTAAACCACTG    12290
Pachon8      TGCCTGGTCTGCCAAGATAAAATGTACTTTGCTTTTTGAATTAAAACATTTAAACCACTG    12290
Pachon15     TGCCTGGTCTGCCAAGATAAAATGTACTTTGCTTTTTGAATTAAAACATTTAAACCACTG    12290
             ***************** ******************************************

Rascon4      TGGAAATTCAAGCACAAGTTGTGAGAGTCTGAATTAATGTGCATTAGCTCATAGTGTGGT    12292
Surface      TGGAAATTCAAGCACAAGTTGTGAGAGTCTGAATTAATGTGCATTAGCTCATAGTGTGGT    12370
Rascon8      TGGAAATTCAAGCACAAGTTGTGAGAGTCTGAATTAATGTGCATTAGCTCATAGTGTGGT    12354
Rascon2      TGGAAATTCAAGCACAAGTTGTGAGAGTCTGAATTAATGTGCATTAGCTCATAGTGTGGT    12349
Rascon15     TGGAAATTCAAGCACAAGTTGTGAGAGTCTGAATTAATGTGCATTAGCTCATAGTGTGGT    12350
Rascon13     TGGAAATTCAAGCACAAGTTGTGAGAGTCTGAATTAATGTGCATTAGCTCATAGTGTGGT    12360
Rascon6      TGGAAATTCAAGCACAAGTTGTGAGAGTCTGAATTAATGTGCATTAGCTCATAGTGTGGT    12361
Pachon14     TGGAAATTCAAGCACAAGTTGTGAGAGTCTGAATTAATGTGCATTAGCTCATAGTGTGGT    12343
Pachon9      TGGAAATTCAAGCACAAGTTGTGAGAGTCTGAATTAATGTGCATTAGCTCATAGTGTGGT    12352
Pachon17     TGGAAATTCAAGCACAAGTTGTGAGAGTCTGAATTAATGTGCATTAGCTCATAGTGTGGT    12350
Pachon12     TGGAAATTCAAGCACAAGTTGTGAGAGTCTGAATTAATGTGCATTAGCTCATAGTGTGGT    12332
Pachon11     TGGAAATTCAAGCACAAGTTGTGAGAGTCTGAATTAATGTGCATTAGCTCATAGTGTGGT    12349
Pachon7      TGGAAATTCAAGCACAAGTTGTGAGAGTCTGAATTAATGTGCATTAGCTCATAGTGTGGT    12350
Pachon3      TGGAAATTCAAGCACAAGTTGTGAGAGTCTGAATTAATGTGCATTAGCTCATAGTGTGGT    12350
Pachon8      TGGAAATTCAAGCACAAGTTGTGAGAGTCTGAATTAATGTGCATTAGCTCATAGTGTGGT    12350
Pachon15     TGGAAATTCAAGCACAAGTTGTGAGAGTCTGAATTAATGTGCATTAGCTCATAGTGTGGT    12350
             ************************************************************

Rascon4      TAGCTGTATTTGTTAGCTCTACTGTGTTTTGATATTCATTGCACTGGGCTAAGTGATGGA    12352
Surface      TAGCTGTATTTGTTAGCTCTACTGTGTTTTGATATTCATTGCACTGGGCTAAGTGATGGA    12430
Rascon8      TAGCTGTATTTGTTAGCTCTACTGTGTTTTGATATTCATTGCACTGGGCTAAGTGATGGA    12414
Rascon2      TAGCTGTATTTGTTAGCTCTACTGTGTTTTGATATTCATTGCACTGGGCTAAGTGATGGA    12409
Rascon15     TAGCTGTATTTGTTAGCTCTACTGTGTTTTGATATTCATTGCACTGGGCTAAGTGATGGA    12410
Rascon13     TAGCTGTATTTGTTAGCTCTACTGTGTTTTGATATTCATTGCACTGGGCTAAGTGATGGA    12420
Rascon6      TAGCTGTATTTGTTAGCTCTACTGTGTTTTGATATTCATTGCACTGGGCTAAGTGATGGA    12421
Pachon14     TAGCTGTATTTGTTAGCTCTACTGTGTTTTGATATTCATTGCACTGGGCTAAGTGATGGA    12403
Pachon9      TAGCTGTATTTGTTAGCTCTACTGTGTTTTGATATTCATTGCACTGGGCTAAGTGATGGA    12412
Pachon17     TAGCTGTATTTGTTAGCTCTACTGTGTTTTGATATTCATTGCACTGGGCTAAGTGATGGA    12410
Pachon12     TAGCTGTATTTGTTAGCTCTACTGTGTTTTGATATTCATTGCACTGGGCTAAGTGATGGA    12392
Pachon11     TAGCTGTATTTGTTAGCTCTACTGTGTTTTGATATTCATTGCACTGGGCTAAGTGATGGA    12409
Pachon7      TAGCTGTATTTGTTAGCTCTACTGTGTTTTGATATTCATTGCACTGGGCTAAGTGATGGA    12410
Pachon3      TAGCTGTATTTGTTAGCTCTACTGTGTTTTGATATTCATTGCACTGGGCTAAGTGATGGA    12410
Pachon8      TAGCTGTATTTGTTAGCTCTACTGTGTTTTGATATTCATTGCACTGGGCTAAGTGATGGA    12410
Pachon15     TAGCTGTATTTGTTAGCTCTACTGTGTTTTGATATTCATTGCACTGGGCTAAGTGATGGA    12410
             ************************************************************

Rascon4      CTTTCTTGAGAGGGG-TATCCAGTTAGTGTTCTGACGCTCAATGTTATGGATGGACCGAA    12411
Surface      CTTTCTTGAGAGGGG-TATCCAGTTAGTGTTCTGACGCTCAATGTTATGGATGGACCGAA    12489
Rascon8      CTTTCTTGAGAGGGG-TATCCAGTTAGTGTTCTGACGCTCAATGTTATGGATGGACCGAA    12473
Rascon2      CTTTCTTGAGAGGGG-TATCCAGTTAGTGTTCTGACGCTCAATGTTATGGATGGACCGAA    12468
Rascon15     CTTTCTTGAGAGGGG-TATCCAGTTAGTGTTCTGACGCTCAATGTTATGGATGGACCGAA    12469
Rascon13     CTTTCTTGAGAGGGG-TATCCAGTTAGTGTTCTGACGCTCAATGTTATGGATGGACCGAA    12479
Rascon6      CTTTCTTGAGAGGGGTTATCCAGTTAGTGTTCTGACGCTCAATGTTATGGATGGACCGAA    12481
Pachon14     CTTTCTTGAGAGGGG-TATCCAGTTAGTGTTCTGACGCTCAATGTTATGGATGGACCGAA    12462
Pachon9      CTTTCTTGAGAGGGG-TATCCAGTTAGTGTTCTGACGCTCAATGTTATGGATGGACCGAA    12471
Pachon17     CTTTCTTGAGAGGGG-TATCCAGTTAGTGTTCTGACGCTCAATGTTATGGATGGACCGAA    12469
Pachon12     CTTTCTTGAGAGGGG-TATCCAGTTAGTGTTCTGACGCTCAATGTTATGGATGGACCGAA    12451
Pachon11     CTTTCTTGAGAGGGG-TATCCAGTTAGTGTTCTGACGCTCAATGTTATGGATGGACCGAA    12468
Pachon7      CTTTCTTGAGAGGGG-TATCCAGTTAGTGTTCTGACGCTCAATGTTATGGATGGACCGAA    12469
Pachon3      CTTTCTTGAGAGGGG-TATCCAGTTAGTGTTCTGACGCTCAATGTTATGGATGGACCGAA    12469
Pachon8      CTTTCTTGAGAGGGG-TATCCAGTTAGTGTTCTGACGCTCAATGTTATGGATGGACCGAA    12469
Pachon15     CTTTCTTGAGAGGGG-TATCCAGTTAGTGTTCTGACGCTCAATGTTATGGATGGACCGAA    12469
             *************** ********************************************

Rascon4      CAGTGTTTTTGTCTTTCAGCTCGACTGGCCTATCAAGGGCAGGATAAGATGCCACATTAA    12471
Surface      CAGTGTTTTTGTCTTTCAGCTCGACTGGCCTATCAAGGGCAGGATAAGATGCCACATTAA    12549
Rascon8      CAGTGTTTTTGTCTTTCAGCTCGACTGGCCTATCAAGGGCAGGATAAGATGCCACATTAA    12533
Rascon2      CAGTGTTTTTGTCTTTCAGCTCGACTGGCCTATCAAGGGCAGGATAAGATGCCACATTAA    12528
Rascon15     CAGTGTTTTTGTCTTTCAGCTCGACTGGCCTATCAAGGGCAGGATAAGATGCCACATTAA    12529
```

```
Rascon13     CAGTGTTTTTGTCTTTCAGCTCGACTGGCCTATCAAGGGCAGGATAAGATGCCACATTAA     12539
Rascon6      CAGTGTTTTTGTCTTTCAGCTCGACTGGCCTATCAAGGGCAGGATAAGATGCCACATTAA     12541
Pachon14     CAGTGTTTTTGTCTTTCAGCTCGACTGGCCTATCAAGGGCAGGATAAGATGCCACATTAA     12522
Pachon9      CAGTGTTTTTGTCTTTCAGCTCGACTGGCCTATCAAGGGCAGGATAAGATGCCACATTAA     12531
Pachon17     CAGTGTTTTTGTCTTTCAGCTCGACTGGCCTATCAAGGGCAGGATAAGATGCCACATTAA     12529
Pachon12     CAGTGTTTTTGTCTTTCAGCTCGACTGGCCTATCAAGGGCAGGATAAGATGCCACATTAA     12511
Pachon11     CAGTGTTTTTGTCTTTCAGCTCGACTGGCCTATCAAGGGCAGGATAAGATGCCACATTAA     12528
Pachon7      CAGTGTTTTTGTCTTTCAGCTCGACTGGCCTATCAAGGGCAGGATAAGATGCCACATTAA     12529
Pachon3      CAGTGTTTTTGTCTTTCAGCTCGACTGGCCTATCAAGGGCAGGATAAGATGCCACATTAA     12529
Pachon8      CAGTGTTTTTGTCTTTCAGCTCGACTGGCCTATCAAGGGCAGGATAAGATGCCACATTAA     12529
Pachon15     CAGTGTTTTTGTCTTTCAGCTCGACTGGCCTATCAAGGGCAGGATAAGATGCCACATTAA     12529
             ************************************************************

Rascon4      TGCCACATAGATGCCAAACTTACCAGA-TGGCCACCTTTAAGCTGTAGATTAATAGGCTG     12530 Grp
Surface      TGCCACATAGATGCCAAACTTACCAGA-TGGCCACCTTTAAGCTGTAGATTAATAGGCTG     12608 Intron2
Rascon8      TGCCACATAGATGCCAAACTTACCAGATTGGCCACCTTTAAGCTGTAGATTAATAGGCTG     12593
Rascon2      TGCCACATAGATGCCAAACTTACCAGA-TGGCCACCTTTAAGCTGTAGATTAATAGGCTG     12587
Rascon15     TGCCACATAGATGCCAAACTTACCAGA-TGGCCACCTTTAAGCTGTAGATTAATAGGCTG     12588
Rascon13     TGCCACATAGATGCCAAACTTACCAGATTGGCCACCTTTAAGCTGTAGATTAATAGGCTG     12599
Rascon6      TGCCACATAGATGCCAAACTTACCAGA-TGGCCACCTTTAAGCTGTAGATTAATAGGCTG     12600
Pachon14     TGCCACATAGATGCCAAACTTACCAGA-TGGCCACCTTTAAGCTGTAGATTAATAGGCTG     12581
Pachon9      TGCCACATAGATGCCAAACTTACCAGA-TGGCCACCTTTAAGCTGTAGATTAATAGGCTG     12590
Pachon17     TGCCACATAGATGCCAAACTTACCAGA-TGGCCACCTTTAAGCTGTAGATTAATAGGCTG     12588
Pachon12     TGCCACATAGATGCCAAACTTACCAGA-TGGCCACCTTTAAGCTGTAGATTAATAGGCTG     12570
Pachon11     TGCCACATAGATGCCAAACTTACCAGA-TGGCCACCTTTAAGCTGTAGATTAATAGGCTG     12587
Pachon7      TGCCACATAGATGCCAAACTTACCAGA-TGGCCACCTTTAAGCTGTAGATTAATAGGCTG     12588
Pachon3      TGCCACATAGATGCCAAACTTACCAGA-TGGCCACCTTTAAGCTGTAGATTAATAGGCTG     12588
Pachon8      TGCCACATAGATGCCAAACTTACCAGA-TGGCCACCTTTAAGCTGTAGATTAATAGGCTG     12588
Pachon15     TGCCACATAGATGCCAAACTTACCAGA-TGGCCACCTTTAAGCTGTAGATTAATAGGCTG     12588
             *************************** ********************************

Rascon4      TTAAACAAACATAACAAAGCTGAGACAGTGAAAATTGCTGCTTATCTTAAGAATATAAAT     12590
Surface      TTAAACAAACATAACAAAGCTGAGACAGTGAAAATTGCTGCTTATCTTAAGAATATAAAT     12668
Rascon8      TTAAACAAACATAACAAAGCTGAGACAGTGAAAATTGCTGCTTATCTTAAGAATATAAAT     12653
Rascon2      TTAAACAAACATAACAAAGCTGAGACAGTGAAAATTGCTGCTTATCTTAAGAATATAAAT     12647
Rascon15     TTAAACAAACATAACAAAGCTGAGACAGTGAAAATTGCTGCTTATCTTAAGAATATAAAT     12648
Rascon13     TTAAACAAACATAACAAAGCTGAGACAGTGAAAATTGCTGCTTATCTTAAGAATATAAAT     12659
Rascon6      TTAAACAAACATAACAAAGCTGAGACAGTGAAAATTGCTGCTTATCTTAAGAATATAAAT     12660
Pachon14     TTAAACAAACATAACAAAGCTGAGACAGTGAAAATTGCTGCTTATCTTAAGAATATAAAT     12641
Pachon9      TTAAACAAACATAACAAAGCTGAGACAGTGAAAATTGCTGCTTATCTTAAGAATATAAAT     12650
Pachon17     TTAAACAAACATAACAAAGCTGAGACAGTGAAAATTGCTGCTTATCTTAAGAATATAAAT     12648
Pachon12     TTAAACAAACATAACAAAGCTGAGACAGTGAAAATTGCTGCTTATCTTAAGAATATAAAT     12630
Pachon11     TTAAACAAACATAACAAAGCTGAGACAGTGAAAATTGCTGCTTATCTTAAGAATATAAAT     12647
Pachon7      TTAAACAAACATAACAAAGCTGAGACAGTGAAAATTGCTGCTTATCTTAAGAATATAAAT     12648
Pachon3      TTAAACAAACATAACAAAGCTGAGACAGTGAAAATTGCTGCTTATCTTAAGAATATAAAT     12648
Pachon8      TTAAACAAACATAACAAAGCTGAGACAGTGAAAATTGCTGCTTATCTTAAGAATATAAAT     12648
Pachon15     TTAAACAAACATAACAAAGCTGAGACAGTGAAAATTGCTGCTTATCTTAAGAATATAAAT     12648
             ************************************************************

Rascon4      ATTTTGAATATGATTTCTTTAGCTTTCTGACTGGTGCTGCTGCTGTACTTTTGCTAGAGG     12650 SNP7
Surface      ATTTTGAATATGATTTCTTTAGCTTTCTGACTGGTGCTGCTGCTGTACTTTTGCTAGAGG     12728 Also
Rascon8      ATTTTGAATATGGTTTCTTTAGCTTTCTGACTGGTGCTGCTGCTGTACTTTTGCTAGAGG     12713 fixed
Rascon2      ATTTTGAATATGATTTCTTTAGCTTTCTGACTGGTGCTGCTGCTGTACTTTTGCTAGAGG     12707 in
Rascon15     ATTTTGAATATGGTTTCTTTAGCTTTCTGACTGGTGCTGCTGCTGTACTTTTGCTAGAGG     12708 Choy SF
Rascon13     ATTTTGAATATGGTTTCTTTAGCTTTCTGACTGGTGCTGCTGCTGTACTTTTGCTAGAGG     12719
Rascon6      ATTTTGAATATGGTTTCTTTAGCTTTCTGACTGGTGCTGCTGCTGTACTTTTGCTAGAGG     12720
Pachon14     ATTTTTAATATGATTTCTTTAGCTTTCTGACTGGTGCTGCTGCTGTACTTTTGCTAGAGG     12701
Pachon9      ATTTTTAATATGATTTCTTTAGCTTTCTGACTGGTGCTGCTGCTGTACTTTTGCTAGAGG     12710
Pachon17     ATTTTTAATATGATTTCTTTAGCTTTCTGACTGGTGCTGCTGCTGTACTTTTGCTAGAGG     12708
Pachon12     ATTTTTAATATGATTTCTTTAGCTTTCTGACTGGTGCTGCTGCTGTACTTTTGCTAGAGG     12690
Pachon11     ATTTTTAATATGATTTCTTTAGCTTTCTGACTGGTGCTGCTGCTGTACTTTTGCTAGAGG     12707
Pachon7      ATTTTTAATATGATTTCTTTAGCTTTCTGACTGGTGCTGCTGCTGTACTTTTGCTAGAGG     12708
Pachon3      ATTTTTAATATGATTTCTTTAGCTTTCTGACTGGTGCTGCTGCTGTACTTTTGCTAGAGG     12708
Pachon8      ATTTTTAATATGATTTCTTTAGCTTTCTGACTGGTGCTGCTGCTGTACTTTTGCTAGAGG     12708
Pachon15     ATTTTTAATATGATTTCTTTAGCTTTCTGACTGGTGCTGCTGCTGTACTTTTGCTAGAGG     12708
             ***** ****** ***********************************************

Rascon4      TTAAATGTGCTCTACTAATCAATTTATGGGCAGAAAGATGCACGATTATAAGCAATATTA     12710
Surface      TTAAATGTGCTCTACTAATCAATTTATGGGCAGAAAGATGCACGATTATAAGCAATATTA     12788
Rascon8      TTAAATGTGCTCTACTAATCAATTTATGGGCAGAAAGATGCACGATTATAAGCAATATTA     12773
Rascon2      TTAAATGTGCTCTACTAATCAATTTATGGGCAGAAAGATGCACGATTATAAGCAATATTA     12767
Rascon15     TTAAATGTGCTCTACTAATCAATTTACGGGCAGAAAGATGCACGATTATAAGCAATATTA     12768
Rascon13     TTAAATGTGCTCTACTAATCAATTTATGGGCAGAAAGATGCACGATTATAAGCAATATTA     12779
Rascon6      TTAAATGTGCTCTACTAATCAATTTACGGGCAGAAAGATGCACGATTATAAGCAATATTA     12780
Pachon14     TTAAATGTGCTCTACTAATCAATTTATGGGCAGAAAGATGCACGATTATAAGCAATATTA     12761
Pachon9      TTAAATGTGCTCTACTAATCAATTTATGGGCAGAAAGATGCACGATTATAAGCAATATTA     12770
Pachon17     TTAAATGTGCTCTACTAATCAATTTATGGGCAGAAAGATGCACGATTATAAGCAATATTA     12768
```

```
Pachon12     TTAAATGTGCTCTACTAATCAATTTATGGGCAGAAAGATGCACGATTATAAGCAATATTA     12750
Pachon11     TTAAATGTGCTCTACTAATCAATTTATGGGCAGAAAGATGCACGATTATAAGCAATATTA     12767
Pachon7      TTAAATGTGCTCTACTAATCAATTTATGGGCAGAAAGATGCACGATTATAAGCAATATTA     12768
Pachon3      TTAAATGTGCTCTACTAATCAATTTATGGGCAGAAAGATGCACGATTATAAGCAATATTA     12768
Pachon8      TTAAATGTGCTCTACTAATCAATTTATGGGCAGAAAGATGCACGATTATAAGCAATATTA     12768
Pachon15     TTAAATGTGCTCTACTAATCAATTTATGGGCAGAAAGATGCACGATTATAAGCAATATTA     12768
             ************************* **********************************

Rascon4      TTTGGTAAGTGGGATTAAAATCTCTTTTCACCACGATGTACGTTAATCCTACACATATGA     12770 Grp
Surface      TTTGGTAAGTGGGATTAAAATCTCTTTTCACCACGATGTATGTTAATCCTACACATATGA     12848 Intron2
Rascon8      TTTGGTAAGTGGGATTAAAATCTCTTTTCACCACGATGTACGTTAATCCTACACATATGA     12833
Rascon2      TTTGGTAAGTGGGATTAAAATCTCTTTTCACCACGATGTACGTTAATCCTACACATATGA     12827
Rascon15     TTTGGTAAGTGGGATTAAAATCTCTTTTCACCACGATGTACGTTAATCCTACACATATGA     12828
Rascon13     TTTGGTAAGTGGGATTAAAATCTCTTTTCACCACGATGTACGTTAATCCTACACATATGA     12839
Rascon6      TTTGGTAAGTGGGATTAAAATCTCTTTTCACCACGATGTACGTTAATCCTACACATATGA     12840
Pachon14     TTTGGTAAGTGGGATTAAAATCTCTTTTCACCACGATGTACGTTAATCCTACACATATGA     12821
Pachon9      TTTGGTAAGTGGGATTAAAATCTCTTTTCACCACGATGTACGTTAATCCTACACATATGA     12830
Pachon17     TTTGGTAAGTGGGATTAAAATCTCTTTTCACCACGATGTACGTTAATCCTACACATATGA     12828
Pachon12     TTTGGTAAGTGGGATTAAAATCTCTTTTCACCACGATGTACGTTAATCCTACACATATGA     12810
Pachon11     TTTGGTAAGTGGGATTAAAATCTCTTTTCACCACGATGTACGTTAATCCTACACATATGA     12827
Pachon7      TTTGGTAAGTGGGATTAAAATCTCTTTTCACCACGATGTACGTTAATCCTACACATATGA     12828
Pachon3      TTTGGTAAGTGGGATTAAAATCTCTTTTCACCACGATGTACGTTAATCCTACACATATGA     12828
Pachon8      TTTGGTAAGTGGGATTAAAATCTCTTTTCACCACGATGTACGTTAATCCTACACATATGA     12828
Pachon15     TTTGGTAAGTGGGATTAAAATCTCTTTTCACCACGATGTACGTTAATCCTACACATATGA     12828
             **************************************** *******************

Rascon4      ATGGCAAAAACAGTAAAACTAAACAAAACTCCAGACGGAGTGAGCAGAGCTGAGCCTGTA     12830
Surface      ATGGCAAAAACAGTAAAACTAAACAAAACTCCAGACGGAGTGAGCAGAGCTGAGCCTGTA     12908
Rascon8      ATGGCAAAAACAGTAAAACTAAACAAAACTCCAGACGGAGTGAGCAGAGCTGAGCCTGTA     12893
Rascon2      ATGGCAAAAACAGTAAAACTAAACAAAACTCCAGACGGAGTGAGCAGAGCTGAGCCTGTA     12887
Rascon15     ATGGCAAAAACAGTAAAACTAAACAAAACTCCAGACGGAGTGAGCAGAGCTGAGCCTGTA     12888
Rascon13     ATGGCAAAAACAGTAAAACTAAACAAAACTCCAGACGGAGTGAGCAGAGCTGAGCCTGTA     12899
Rascon6      ATGGCAAAAACAGTAAAACTAAACAAAACTCCAGACGGAGTGAGCAGAGCTGAGCCTGTA     12900
Pachon14     ATGGCAAAAACAGTAAAACTAAACAAAACTCCAGACGGAGTGAGCAGAGCTGAGCCTGTA     12881
Pachon9      ATGGCAAAAACAGTAAAACTAAACAAAACTCCAGACGGAGTGAGCAGAGCTGAGCCTGTA     12890
Pachon17     ATGGCAAAAACAGTAAAACTAAACAAAACTCCAGACGGAGTGAGCAGAGCTGAGCCTGTA     12888
Pachon12     ATGGCAAAAACAGTAAAACTAAACAAAACTCCAGACGGAGTGAGCAGAGCTGAGCCTGTA     12870
Pachon11     ATGGCAAAAACAGTAAAACTAAACAAAACTCCAGACGGAGTGAGCAGAGCTGAGCCTGTA     12887
Pachon7      ATGGCAAAAACAGTAAAACTAAACAAAACTCCAGACGGAGTGAGCAGAGCTGAGCCTGTA     12888
Pachon3      ATGGCAAAAACAGTAAAACTAAACAAAACTCCAGACGGAGTGAGCAGAGCTGAGCCTGTA     12888
Pachon8      ATGGCAAAAACAGTAAAACTAAACAAAACTCCAGACGGAGTGAGCAGAGCTGAGCCTGTA     12888
Pachon15     ATGGCAAAAACAGTAAAACTAAACAAAACTCCAGACGGAGTGAGCAGAGCTGAGCCTGTA     12888
             ************************************************************

Rascon4      GCCCTGGCTGACTCAGCTGAGAGTGGTTATTATTATTAAAAATTGAAGCATTATTGAACC     12890
Surface      GCCCTGGCTGACTCAGCTGAGAGTGGTTATTATTATTAAAAATTGAAGCATTATTGAACC     12968
Rascon8      GCCCTGGCTGACTCAGCTGAGAGTGGTTATTATTATTAAAAATTGAAGCATTATTGAACC     12953
Rascon2      GCCCTGGCTGACTCAGCTGAGAGTGGTTATTATTATTAAAAATTGAAGCATTATTGAACC     12947
Rascon15     GCCCTGGCTGACTCAGCTGAGAGTGGTTATTATTATTAAAAATTGAAGCATTATTGAACC     12948
Rascon13     GCCCTGGCTGACTCAGCTGAGAGTGGTTATTATTATTAAAAATTGAAGCATTATTGAACC     12959
Rascon6      GCCCTGGCTGACTCAGCTGAGAGTGGTTATTATTATTAAAAATTGAAGCATTATTGAACC     12960
Pachon14     GCCCTGGCTGACTCAGCTGAGAGTGGTTATTATTATTAAAAATTGAAGCATTATTGAACC     12941
Pachon9      GCCCTGGCTGACTCAGCTGAGAGTGGTTATTATTATTAAAAATTGAAGCATTATTGAACC     12950
Pachon17     GCCCTGGCTGACTCAGCTGAGAGTGGTTATTATTATTAAAAATTGAAGCATTATTGAACC     12948
Pachon12     GCCCTGGCTGACTCAGCTGAGAGTGGTTATTATTATTAAAAATTGAAGCATTATTGAACC     12930
Pachon11     GCCCTGGCTGACTCAGCTGAGAGTGGTTATTATTATTAAAAATTGAAGCATTATTGAACC     12947
Pachon7      GCCCTGGCTGACTCAGCTGAGAGTGGTTATTATTATTAAAAATTGAAGCATTATTGAACC     12948
Pachon3      GCCCTGGCTGACTCAGCTGAGAGTGGTTATTATTATTAAAAATTGAAGCATTATTGAACC     12948
Pachon8      GCCCTGGCTGACTCAGCTGAGAGTGGTTATTATTATTAAAAATTGAAGCATTATTGAACC     12948
Pachon15     GCCCTGGCTGACTCAGCTGAGAGTGGTTATTATTATTAAAAATTGAAGCATTATTGAACC     12948
             ************************************************************

Rascon4      ACAGCAACTCAGCCACATGTCAGATCAC--AAAGTTACAGAGTGGTGTCTCTAAGTGCTA     12948
Surface      ACAGCAACTCAGCCACATGTCAGATCACATAAAGTTACAGAGTGGTGTCTCTAAGTGCTA     13028
Rascon8      A--------CAGCCACATGTCAGATCACATAAAGTTACAGAGTGGTGTCTCTAAGTGCTA     13005
Rascon2      ACAGCAACTCAGCCACATGTCAGATCAC--AAAGTTACAGAGTGGTGTCTCTAAGTGCTA     13005
Rascon15     ACAGCAACTCAGCCACATGTCAGATCAC--AAAGTTACAGAGTGGTGTCTCTAAGTGCTA     13006
Rascon13     ACAGCAACTCAGCCACATGTCAGATCACATAAAGTTACAGAGTGGTGTCTCTAAGTGCTA     13019
Rascon6      ACAGCAACTCAGCCACATGTCAGATCACATAAAGTTACAGAGTGGTGTCTCTAAGTGCTA     13020
Pachon14     ACAGCAACTCAGCCACATGTCAGATCACATAAAGTTACAGAGTGGTGTCTCTAAGTGCTA     13001
Pachon9      ACAGCAACTCAGCCACATGTCAGATCACATAAAGTTACAGAGTGGTGTCTCTAAGTGCTA     13010
Pachon17     ACAGCAACTCAGCCACATGTCAGATCACATAAAGTTACAGAGTGGTGTCTCTAAGTGCTA     13008
Pachon12     ACAGCAACTCAGCCACATGTCAGATCACATAAAGTTACAGAGTGGTGTCTCTAAGTGCTA     12990
Pachon11     ACAGCAACTCAGCCACATGTCAGATCACATAAAGTTACAGAGTGGTGTCTCTAAGTGCTA     13007
Pachon7      ACAGCAACTCAGCCACATGTCAGATCACATAAAGTTACAGAGTGGTGTCTCTAAGTGCTA     13008
Pachon3      ACAGCAACTCAGCCACATGTCAGATCACATAAAGTTACAGAGTGGTGTCTCTAAGTGCTA     13008
Pachon8      ACAGCAACTCAGCCACATGTCAGATCACATAAAGTTACAGAGTGGTGTCTCTAAGTGCTA     13008
```

```
Pachon15    ACAGCAACTCAGCCACATGTCAGATCACATAAAGTTACAGAGTGGTGTCTCTAAGTGCTA    13008
            *        *******************  ******************************

Rascon4     AGATTTTACAATGTTTTCTTATTGTAACAGTGGTAAACTAGCAGCTAATTTTGTAGTAGC    13008 Grp
Surface     AGATTTTACAATGTTTTCTTATTGTAACAGTGGAAAACTAGCAGCTAATTTTGTAGTAGC    13088 Intron2
Rascon8     AGATTTTACAATGTTTTCTTATTGTAACAGTGGTAAACTAGCAGCTAATTTTGTAGTAGC    13065
Rascon2     AGATTTTACAATGTTTTCTTATTGTAACAGTGGTAAACTAGCAGCTAATTTTGTAGTAGC    13065
Rascon15    AGATTTTACAATGTTTTCTTATTGTAACAGTGGTAAACTAGCAGCTAATTTTGTAGTAGC    13066
Rascon13    AGATTTTACAATGTTTTCTTATTGTAACAGTGGTAAACTAGCAGCTAATTTTGTAGTAGC    13079
Rascon6     AGATTTTACAATGTTTTCTTATTGTAACAGTGGTAAACTAGCAGCTAATTTTGTAGTAGC    13080
Pachon14    AGATTTTACAATGTTTTCTTATTGTAACAGTGGTAAACTAGCAGCTAATTTTGTAGTAGC    13061
Pachon9     AGATTTTACAATGTTTTCTTATTGTAACAGTGGTAAACTAGCAGCTAATTTTGTAGTAGC    13070
Pachon17    AGATTTTACAATGTTTTCTTATTGTAACAGTGGTAAACTAGCAGCTAATTTTGTAGTAGC    13068
Pachon12    AGATTTTACAATGTTTTCTTATTGTAACAGTGGTAAACTAGCAGCTAATTTTGTAGTAGC    13050
Pachon11    AGATTTTACAATGTTTTCTTATTGTAACAGTGGTAAACTAGCAGCTAATTTTGTAGTAGC    13067
Pachon7     AGATTTTACAATGTTTTCTTATTGTAACAGTGGTAAACTAGCAGCTAATTTTGTAGTAGC    13068
Pachon3     AGATTTTACAATGTTTTCTTATTGTAACAGTGGTAAACTAGCAGCTAATTTTGTAGTAGC    13068
Pachon8     AGATTTTACAATGTTTTCTTATTGTAACAGTGGTAAACTAGCAGCTAATTTTGTAGTAGC    13068
Pachon15    AGATTTTACAATGTTTTCTTATTGTAACAGTGGTAAACTAGCAGCTAATTTTGTAGTAGC    13068
            ********************************* **************************

Rascon4     AGTATATCATGAAGTTTCTAATCTGTGATTTTTAGATGCTAAGCACTTCCGAGATCTAGC    13068
Surface     AGTATATCATGAAGTTTCTAATCTGTGATTTTTAGATGCTAAGCACTTCCGAGATCTAGC    13148
Rascon8     AGTATATCATGAAGTTTCTAATCTGTGATTTTTAGATGCTAAGCACTTCCGAGATCTAGC    13125
Rascon2     AGTATATCATGAAGTTTCTAATCTGTGATTTTTAGATGCTAAGCACTTCCGAGATCTAGC    13125
Rascon15    AGTATATCATGAAGTTTCTAATCTGTGATTTTTAGATGCTAAGCACTTCCGAGATCTAGC    13126
Rascon13    AGTATATCATGAAGTTTCTAATCTGTGATTTTTAGATGCTGAGCACTTCCGAGATCTAGC    13139
Rascon6     AGTATATCATGAAGTTTCTAATCTGTGATTTTTAGATGCTAAGCACTTCCGAGATCTAGC    13140
Pachon14    AGTATATCATGAAGTTTCTAATCTGTGATTTTTAGATGCTAAGCACTTCCGAGATCTAGC    13121
Pachon9     AGTATATCATGAAGTTTCTAATCTGTGATTTTTAGATGCTAAGCACTTCCGAGATCTAGC    13130
Pachon17    AGTATATCATGAAGTTTCTAATCTGTGATTTTTAGATGCTAAGCACTTCCGAGATCTAGC    13128
Pachon12    AGTATATCATGAAGTTTCTAATCTGTGATTTTTAGATGCTAAGCACTTCCGAGATCTAGC    13110
Pachon11    AGTATATCATGAAGTTTCTAATCTGTGATTTTTAGATGCTAAGCACTTCCGAGATCTAGC    13127
Pachon7     AGTATATCATGAAGTTTCTAATCTGTGATTTTTAGATGCTAAGCACTTCCGAGATCTAGC    13128
Pachon3     AGTATATCATGAAGTTTCTAATCTGTGATTTTTAGATGCTAAGCACTTCCGAGATCTAGC    13128
Pachon8     AGTATATCATGAAGTTTCTAATCTGTGATTTTTAGATGCTAAGCACTTCCGAGATCTAGC    13128
Pachon15    AGTATATCATGAAGTTTCTAATCTGTGATTTTTAGATGCTAAGCACTTCCGAGATCTAGC    13128
            **************************************** *******************

Rascon4     TCTTTCACTCATTCACTTACACACTTATTTAGTCACATTCAACCAGGTGGAGGAATCTCC    13128
Surface     TCTTTCACTCATTCACTTACACACTTATTTAGTCACATTCAACCAGGTGGAGGAATCTCC    13208
Rascon8     TCTTTCCCTCATTCACTTACACACTTATTTAGTCACATTCAACCAGGTGGAGGAATCTCC    13185
Rascon2     TCTTTCCCTCATTCACTTACACACTTATTTAGTCACATTCAACCAGGTGGAGGAATCTCC    13185
Rascon15    TCTTTCCCTCATTCACTTACACACTTATTTAGTCACATTCAACCAGGTGGAGGAATCTCC    13186
Rascon13    TCTTTCCCTCATTCACTTACACACTTATTTAGTCACATTCAACCAGGTGGAGGAATCTCC    13199
Rascon6     TCTTTCCCTCATTCACTTACACACTTATTTAGTCACATTCAACCAGGTGGAGGAATCTCC    13200
Pachon14    TCTTTCCCTCATTCACTTACACACTTATTTAGTCACATTCAACCAGGTGGAGGAATCTCC    13181
Pachon9     TCTTTCCCTCATTCACTTACACACTTATTTAGTCACATTCAACCAGGTGGAGGAATCTCC    13190
Pachon17    TCTTTCCCTCATTCACTTACACACTTATTTAGTCACATTCAACCAGGTGGAGGAATCTCC    13188
Pachon12    TCTTTCCCTCATTCACTTACACACTTATTTAGTCACATTCAACCAGGTGGAGGAATCTCC    13170
Pachon11    TCTTTCCCTCATTCACTTACACACTTATTTAGTCACATTCAACCAGGTGGAGGAATCTCC    13187
Pachon7     TCTTTCCCTCATTCACTTACACACTTATTTAGTCACATTCAACCAGGTGGAGGAATCTCC    13188
Pachon3     TCTTTCCCTCATTCACTTACACACTTATTTAGTCACATTCAACCAGGTGGAGGAATCTCC    13188
Pachon8     TCTTTCCCTCATTCACTTACACACTTATTTAGTCACATTCAACCAGGTGGAGGAATCTCC    13188
Pachon15    TCTTTCCCTCATTCACTTACACACTTATTTAGTCACATTCAACCAGGTGGAGGAATCTCC    13188
            ****** *****************************************************

Rascon4     CCAAGGAAGTGCTCTTAGATTTCAAATAGCTTTTGGAGCTGTACATTAATGGCACTGTGT    13188
Surface     CCAAGGAAGTGCTCTTAGATTTCAAATAGCTTTTGGAGCTGTACATTAATGGCACTGTGT    13268
Rascon8     CCAAGGAAGTGCTCTTAGATTTCAAATAGCTTTTGGAGCTGTACATTAATGGCACTGTGT    13245
Rascon2     CCAAGGAAGTGCTCTTAGATTTCAAATAGCTTTTGGAGCTGTACATTAATGGCACTGTGT    13245
Rascon15    CCAAGGAAGTGCTCTTAGATTTCAAATAGCTTTTGGAGCTGTACATTAATGGCACTGTGT    13246
Rascon13    CCAAGGAAGTGCTCTTAGATTTCAAATAGCTTTTGGAGCTGTACATTAATGGCACTGTGT    13259
Rascon6     CCAAGGAAGTGCTCTTAGATTTCAAATAGCTTTTGGAGCTGTACATTAATGGCACTGTGT    13260
Pachon14    CCAAGGAAGTGCTCTTAGATTTCAAATAGCTTTTGGAGCTGTACATTAATGGCACTGTGT    13241
Pachon9     CCAAGGAAGTGCTCTTAGATTTCAAATAGCTTTTGGAGCTGTACATTAATGGCACTGTGT    13250
Pachon17    CCAAGGAAGTGCTCTTAGATTTCAAATAGCTTTTGGAGCTGTACATTAATGGCACTGTGT    13248
Pachon12    CCAAGGAAGTGCTCTTAGATTTCAAATAGCTTTTGGAGCTGTACATTAATGGCACTGTGT    13230
Pachon11    CCAAGGAAGTGCTCTTAGATTTCAAATAGCTTTTGGAGCTGTACATTAATGGCACTGTGT    13247
Pachon7     CCAAGGAAGTGCTCTTAGATTTCAAATAGCTTTTGGAGCTGTACATTAATGGCACTGTGT    13248
Pachon3     CCAAGGAAGTGCTCTTAGATTTCAAATAGCTTTTGGAGCTGTACATTAATGGCACTGTGT    13248
Pachon8     CCAAGGAAGTGCTCTTAGATTTCAAATAGCTTTTGGAGCTGTACATTAATGGCACTGTGT    13248
Pachon15    CCAAGGAAGTGCTCTTAGATTTCAAATAGCTTTTGGAGCTGTACATTAATGGCACTGTGT    13248
            ************************************************************

Rascon4     TCTTATTTAAACTAAATAAAGAAGTTGAACTGATTCATTATTACAGTAGTAATGATACCT    13248
Surface     TCTTATTTAAACTAAATAAAGAAGTTGAACTGATTCATTATTACAGTAGTAATGATACCT    13328
```

```
Rascon8      TCTTATTTAAACTAAATAAAGAAGTTGAACTGATTCATTATTACAGTAGTAATGATACCT    13305
Rascon2      TCTTATTTAAACTAAATAAAGAAGTTGAACTGATTCATTATTACAGTAGTAATGATACCT    13305
Rascon15     TCTTATTTAAACTAAATAAAGAAGTTGAACTGATTCATTATTACAGTAGTAATGATACCT    13306
Rascon13     TCTTATTTAAACTAAATAAAGAAGTTGAACTGATTCATTATTACAGTAGTAATGATACCT    13319
Rascon6      TCTTATTTAAACTAAATAAAGAAGTTGAACTGATTCATTATTACAGTAGTAATGATACCT    13320
Pachon14     TCTTATTTAAACTAAATAAAGAAGTTGAACTGATTCATTATTACAGTAGTAATGATACCT    13301
Pachon9      TCTTATTTAAACTAAATAAAGAAGTTGAACTGATTCATTATTACAGTAGTAATGATACCT    13310
Pachon17     TCTTATTTAAACTAAATAAAGAAGTTGAACTGATTCATTATTACAGTAGTAATGATACCT    13308
Pachon12     TCTTATTTAAACTAAATAAAGAAGTTGAACTGATTCATTATTACAGTAGTAATGATACCT    13290
Pachon11     TCTTATTTAAACTAAATAAAGAAGTTGAACTGATTCATTATTACAGTAGTAATGATACCT    13307
Pachon7      TCTTATTTAAACTAAATAAAGAAGTTGAACTGATTCATTATTACAGTAGTAATGATACCT    13308
Pachon3      TCTTATTTAAACTAAATAAAGAAGTTGAACTGATTCATTATTACAGTAGTAATGATACCT    13308
Pachon8      TCTTATTTAAACTAAATAAAGAAGTTGAACTGATTCATTATTACAGTAGTAATGATACCT    13308
Pachon15     TCTTATTTAAACTAAATAAAGAAGTTGAACTGATTCATTATTACAGTAGTAATGATACCT    13308
             ************************************************************

Rascon4      ATTGCCTCATCATCCATTTCTACTAAAAGTTAATAATACACCACATGATTGACATTTTAG    13308 Grp
Surface      ATTGCCTCATCATCCATTTCTACTAAAAGTTAATAATACACCACATGATTGACATTTTAG    13388 Intron2
Rascon8      ATTGCCTCATCATCCATTTCCACTAAAAGTTAATAATACACCACATGATTGACATTTTAG    13365 start
Rascon2      ATTGCCTCATCATCCATTTCTACTAAAAGTTAATAATACACCACATGATTGACATTTTAG    13365
Rascon15     ATTGCCTCATCATCCATTTCTACTAAAAGTTAATAATACACCACATGATTGACATTTTAG    13366
Rascon13     ATTGCCTCATCATCCATTTCTACTAAAAGTTAATAATACACCACATGATTGACATTTTAG    13379
Rascon6      ATTGCCTCATCATCCATTTCTACTAAACGTTAATAATACACCACATGATTGACATTTTAG    13380
Pachon14     ATTGCCTCATCATCCATTTCTACTAAAAGTTAATAATACACCACATGATTGACATTTTAG    13361
Pachon9      ATTGCCTCATCATCCATTTCTACTAAAAGTTAATAATACACCACATGATTGACATTTTAG    13370
Pachon17     ATTGCCTCATCATCCATTTCTACTAAAAGTTAATAATACACCACATGATTGACATTTTAG    13368
Pachon12     ATTGCCTCATCATCCATTTCTACTAAAAGTTAATAATACACCACATGATTGACATTTTAG    13350
Pachon11     ATTGCCTCATCATCCATTTCTACTAAAAGTTAATAATACACCACATGATTGACATTTTAG    13367
Pachon7      ATTGCCTCATCATCCATTTCTACTAAAAGTTAATAATACACCACATGATTGACATTTTAG    13368
Pachon3      ATTGCCTCATCATCCATTTCTACTAAAAGTTAATAATACACCACATGATTGACATTTTAG    13368
Pachon8      ATTGCCTCATCATCCATTTCTACTAAAAGTTAATAATACACCACATGATTGACATTTTAG    13368
Pachon15     ATTGCCTCATCATCCATTTCTACTAAAAGTTAATAATACACCACATGATTGACATTTTAG    13368
             ******************** ****** ********************************

Rascon4      GATGGAAGAAAGACTCGCCTGTTCTCTCTCTCGCTCAAGCTGCTCTTCTAAGAGGTTCCT    13368 Grp
Surface      G-----AGAAAGACTCGCCTGTTCTCTCTCTCGCTCAAGCTGCTCTTCTAAGAGGTTCCT    13443 Exon2
Rascon8      G-----AGAAAGACTCGCCTGTTCTCTCTCTCGCTCAAGCTGCTCTTCTAAGAGGTTCCT    13420 End
Rascon2      GATGGAAGAAAGACTCGCCTGTTCTCTCTCTCGCTCAAGCTGCTCTTCTAAGAGGTTCCT    13425
Rascon15     GATGGAAGAAAGACTCGCCTGTTCTCTCTCTCGCTCAAGCTGCTCTTCTAAGAGGTTCCT    13426
Rascon13     G-----AGAAAGACTCGCCTGTTCTCTCTCTCGCTCAAGCTGCTCTTCTAAGAGGTTCCT    13434
Rascon6      GATGGAAGAAAGACTCGCCTGTTCTCTCTCTCGCTCAAGCTGCTCTTCTAAGAGGTTCCT    13440
Pachon14     GATGGAAGAAAGACTCGCCTGTTCTCTCTCTCGCTCAAGCTGCTCTTCTAAGAGGTTCCT    13421
Pachon9      GATGGAAGAAAGACTCGCCTGTTCTCTCTCTCGCTCAAGCTGCTCTTCTAAGAGGTTCCT    13430
Pachon17     GATGGAAGAAAGACTCGCCTGTTCTCTCTCTCGCTCAAGCTGCTCTTCTAAGAGGTTCCT    13428
Pachon12     GATGGAAGAAAGACTCGCCTGTTCTCTCTCTCGCTCAAGCTGCTCTTCTAAGAGGTTCCT    13410
Pachon11     GATGGAAGAAAGACTCGCCTGTTCTCTCTCTCGCTCAAGCTGCTCTTCTAAGAGGTTCCT    13427
Pachon7      GATGGAAGAAAGACTCGCCTGTTCTCTCTCTCGCTCAAGCTGCTCTTCTAAGAGGTTCCT    13428
Pachon3      GATGGAAGAAAGACTCGCCTGTTCTCTCTCTCGCTCAAGCTGCTCTTCTAAGAGGTTCCT    13428
Pachon8      GATGGAAGAAAGACTCGCCTGTTCTCTCTCTCGCTCAAGCTGCTCTTCTAAGAGGTTCCT    13428
Pachon15     GATGGAAGAAAGACTCGCCTGTTCTCTCTCTCGCTCAAGCTGCTCTTCTAAGAGGTTCCT    13428
             *     ******************************************************

Rascon4      CTGCTCCAGAGCACGCTGCTGTCTCAGGACGGTGTCTCTCTCTCTTCCTCTTCCTCTCGC    13428 SNP8
Surface      CTGCTCCAGAGCACGCTGCTGTCTCAGGACGGTGTCTCTCTCTCTTCCTCTTCCTCTCGC    13503 Also
Rascon8      CTGCTCCAGAGCACGCTGCTGTCTCAGGACGGTGTCTCTCTCTCTTCCTCTTCCTCTCGC    13480 fixed
Rascon2      CTGCTCCAGAGCACGCTGCTGTCTCAGGACGGTGTCTCTCTCTCTTCCTCTTCCTCTCGC    13485 in
Rascon15     CTGCTCCAGAGCACGCTGCTGTCTCAGGACGGTGTCTCTCTCTCTTCCTCTTCCTCTCGC    13486 Choy SF
Rascon13     CTGCTCCAGAGCACGCTGCTGTCTCAGGACGGTGTCTCTCTCTCTTCCTCTTCCTCTCGC    13494
Rascon6      CTGCTCCAGAGCACGCTGCTGTCTCAGGACGGTGTCTCTCTCTCTTCCTCTTCCTCTCGC    13500
Pachon14     CTGCTCCAGAGCACGCTGCTGTCTCAGGACGGTGTCTCTCT------CTCTTCCTCTCGC    13475
Pachon9      CTGCTCCAGAGCACGCTGCTGTCTCAGGACGGTGTCTCTCT------CTCTTCCTCTCGC    13484
Pachon17     CTGCTCCAGAGCACGCTGCTGTCTCAGGACGGTGTCTCTCT------CTCTTCCTCTCGC    13482
Pachon12     CTGCTCCAGAGCACGCTGCTGTCTCAGGACGGTGTCTCTCT------CTCTTCCTCTCGC    13464
Pachon11     CTGCTCCAGAGCACGCTGCTGTCTCAGGACGGTGTCTCTCT------CTCTTCCTCTCGC    13481
Pachon7      CTGCTCCAGAGCACGCTGCTGTCTCAGGACGGTGTCTCTCT------CTCTTCCTCTCGC    13482
Pachon3      CTGCTCCAGAGCACGCTGCTGTCTCAGGACGGTGTCTCTCT------CTCTTCCTCTCGC    13482
Pachon8      CTGCTCCAGAGCACGCTGCTGTCTCAGGACGGTGTCTCTCT------CTCTTCCTCTCGC    13482
Pachon15     CTGCTCCAGAGCACGCTGCTGTCTCAGGACGGTGTCTCTCT------CTCTTCCTCTCGC    13482
             *****************************************      *************

Rascon4      TCTTGCTCTCTCTGGTCCAGCCAGAGCAGTGATGAGCGCCTGCAGAAATCCGGACGGCTG    13488
Surface      TCTTGCTCTCTCTGGTCCAGCCAGAGCAGTGATGAGCGCCTGCAGAAATCCGGACGGCTG    13563
Rascon8      TCTTGCTCTCTCTGGTCCAGCCAGAGCAGTGATGAGCGCCTGCAGAAATCCGGACGGCTG    13540
Rascon2      TCTTGCTCTCTCTGGTCCAGCCAGAGCAGTGATGAGCGCCTGCAGAAATCCGGACGGCTG    13545
Rascon15     TCTTGCTCTCTCTGGTCCAGCCAGAGCAGTGATGAGCGCCTGCAGAAATCCGGACGGCTG    13546
Rascon13     TCTTGCTCTCTCTGGTCCAGCCAGAGCAGTGATGAGCGCCTGCAGAAATCCGGACGGCTG    13554
Rascon6      TCTTGCTCTCTCTGGTCCAGCCAGAGCAGTGATGAGCGCCTGCAGAAATCCGGACGGCTG    13560
```

```
Pachon14    TCTTGCTCTCTCTGGTCCAGCCAGAGCAGTGATGAGCGCCTGCAGAAATCCGGACGGCTG    13535
Pachon9     TCTTGCTCTCTCTGGTCCAGCCAGAGCAGTGATGAGCGCCTGCAGAAATCCGGACGGCTG    13544
Pachon17    TCTTGCTCTCTCTGGTCCAGCCAGAGCAGTGATGAGCGCCTGCAGAAATCCGGACGGCTG    13542
Pachon12    TCTTGCTCTCTCTGGTCCAGCCAGAGCAGTGATGAGCGCCTGCAGAAATCCGGACGGCTG    13524
Pachon11    TCTTGCTCTCTCTGGTCCAGCCAGAGCAGTGATGAGCGCCTGCAGAAATCCGGACGGCTG    13541
Pachon7     TCTTGCTCTCTCTGGTCCAGCCAGAGCAGTGATGAGCGCCTGCAGAAATCCGGACGGCTG    13542
Pachon3     TCTTGCTCTCTCTGGTCCAGCCAGAGCAGTGATGAGCGCCTGCAGAAATCCGGACGGCTG    13542
Pachon8     TCTTGCTCTCTCTGGTCCAGCCAGAGCAGTGATGAGCGCCTGCAGAAATCCGGACGGCTG    13542
Pachon15    TCTTGCTCTCTCTGGTCCAGCCAGAGCAGTGATGAGCGCCTGCAGAAATCCGGACGGCTG    13542
            ************************************************************

Rascon4     CGTGTATTCCTCAACCTCTGCTGTGGTTAAATATCTCTGGCCTCCACTGTCTTGTTCGAC    13548
Surface     CGTGTATTCCTCAACCTCTGCTGTGGTTAAATATCTCTGGCCTCCACTGTCTTGTTCGAC    13623
Rascon8     CATGTATTCCTCAACCTCTGCTGTGGTTAAATATCTCTGGCCTCCACTGTCTTGTTCGAC    13600
Rascon2     CGTGTATTCCTCAACCTCTGCTGTGGTTAAATATCTCTGGCCTCCACTGTCTTGTTCGAC    13605
Rascon15    CGTGTATTCCTCAACCTCTGCTGTGGTTAAATATCTCTGGCCTCCACTGTCTTGTTCGAC    13606
Rascon13    CGTGTATTCCTCAACCTCTGCTGTGGTTAAATATCTCTGGCCTCCACTGTCTTGTTCGAC    13614
Rascon6     CGTGTATTCCTCAACCTCTGCTGTGGTTAAATATCTCTGGCCTCTACTGTCTTGTTCGAC    13620
Pachon14    CGTGTATTCCTCAACCTCTGCTGTGGTTAAATATCTCTGGCCTCCACTGTCTTGTTCGAC    13595
Pachon9     CGTGTATTCCTCAACCTCTGCTGTGGTTAAATATCTCTGGCCTCCACTGTCTTGTTCGAC    13604
Pachon17    CGTGTATTCCTCAACCTCTGCTGTGGTTAAATATCTCTGGCCTCCACTGTCTTGTTCGAC    13602
Pachon12    CGTGTATTCCTCAACCTCTGCTGTGGTTAAATATCTCTGGCCTCCACTGTCTTGTTCGAC    13584
Pachon11    CGTGTATTCCTCAACCTCTGCTGTGGTTAAATATCTCTGGCCTCCACTGTCTTGTTCGAC    13601
Pachon7     CGTGTATTCCTCAACCTCTGCTGTGGTTAAATATCTCTGGCCTCCACTGTCTTGTTCGAC    13602
Pachon3     CGTGTATTCCTCAACCTCTGCTGTGGTTAAATATCTCTGGCCTCCACTGTCTTGTTCGAC    13602
Pachon8     CGTGTATTCCTCAACCTCTGCTGTGGTTAAATATCTCTGGCCTCCACTGTCTTGTTCGAC    13602
Pachon15    CGTGTATTCCTCAACCTCTGCTGTGGTTAAATATCTCTGGCCTCCACTGTCTTGTTCGAC    13602
            * ****************************************** ***************

Rascon4     CGAGTCCAGCGACTCATCTATGCTCTTCTTCCCCATCAGGTGTCCTGTGGAAAATGAGCA    13608 Grp
Surface     CGAGTCCAGCGACTCATCTATGCTCTTCTTCCCCATCAGGTGTCCTGTGGAAAATGAGCA    13683 Exon2
Rascon8     CGAGTCCAGCGACTCATCTATGCTCTTCTTCCCCATCAGGTGTCCTGTGGAAAATGAGCA    13660 start
Rascon2     CGAGTCCAGCGACTCATCTATGCTCTTCTTCCCCATCAGGTGTCCTGTGGAAAATGAGCA    13665 Intron1
Rascon15    CGAGTCCAGCGACTCATCTATGCTCTTCTTCCCCATCAGGTGTCCTGTGGAAAATGAGCA    13666 end
Rascon13    CGAGTCCAGCGACTCATCTATGCTCTTCTTCCCCATCAGGTGTCCTGTGGAAAATGAGCA    13674
Rascon6     CGAGTCCAGCGACTCATCTATGCTCTTCTTCCCCATCAGGTGTCCTGTGGAAAATGAGCA    13680
Pachon14    CGAGTCCAGCGACTCATCTATGCTCTTCTTCCCCATCAGGTGTCCTGTGGAAAATGAGCA    13655
Pachon9     CGAGTCCAGCGACTCATCTATGCTCTTCTTCCCCATCAGGTGTCCTGTGGAAAATGAGCA    13664
Pachon17    CGAGTCCAGCGACTCATCTATGCTCTTCTTCCCCATCAGGTGTCCTGTGGAAAATGAGCA    13662
Pachon12    CGAGTCCAGCGACTCATCTATGCTCTTCTTCCCCATCAGGTGTCCTGTGGAAAATGAGCA    13644
Pachon11    CGAGTCCAGCGACTCATCTATGCTCTTCTTCCCCATCAGGTGTCCTGTGGAAAATGAGCA    13661
Pachon7     CGAGTCCAGCGACTCATCTATGCTCTTCTTCCCCATCAGGTGTCCTGTGGAAAATGAGCA    13662
Pachon3     CGAGTCCAGCGACTCATCTATGCTCTTCTTCCCCATCAGGTGTCCTGTGGAAAATGAGCA    13662
Pachon8     CGAGTCCAGCGACTCATCTATGCTCTTCTTCCCCATCAGGTGTCCTGTGGAAAATGAGCA    13662
Pachon15    CGAGTCCAGCGACTCATCTATGCTCTTCTTCCCCATCAGGTGTCCTGTGGAAAATGAGCA    13662
            ************************************************************

Rascon4     GAAATTGTTAAAGAAGGCGATCAGGTTTCTTCTTGTGAGGTAAATATGATGTACAGAACA    13668
Surface     GAAATTGTTAGAGAAGGCGATCAGGTTTCTTCTTGTGAGGTAAATATGATGTACAGAACA    13743
Rascon8     GAAATTGTTAAAGAAGGCGATCAGGTTTCTTCTTGTGAGGTAAATATGATGTACAGAACA    13720
Rascon2     GAAATTGTTAAAGAAGGCGATCAGGTTTCTTCTTGTGAGGTAAATATGATGTACAGAACA    13725
Rascon15    GAAATTGTTAAAGAAGGCGATCAGGTTTCTTCTTGTGAGGTAAATATGATGTACAGAACA    13726
Rascon13    GAAATTGTTAAAGAAGGCAATCAAGTTTCTTCTTGTGAGGTAAATATGATGTACAGAACA    13734
Rascon6     TAAACTGTTAAAGAAGGCGATCAGGTTTCTTCTTGTGAGGTAAATATGATGTACAGAACA    13740
Pachon14    GAAATTGTTAAAGAAGGCGATCAGGTTTCTTCTTGTGAGGTAAATATGATGTACAGAACA    13715
Pachon9     GAAATTGTTAAAGAAGGCGATCAGGTTTCTTCTTGTGAGGTAAATATGATGTACAGAACA    13724
Pachon17    GAAATTGTTAAAGAAGGCGATCAGGTTTCTTCTTGTGAGGTAAATATGATGTACAGAACA    13722
Pachon12    GAAATTGTTAAAGAAGGCGATCAGGTTTCTTCTTGTGAGGTAAATATGATGTACAGAACA    13704
Pachon11    GAAATTGTTAAAGAAGGCGATCAGGTTTCTTCTTGTGAGGTAAATATGATGTACAGAACA    13721
Pachon7     GAAATTGTTAAAGAAGGCGATCAGGTTTCTTCTTGTGAGGTAAATATGATGTACAGAACA    13722
Pachon3     GAAATTGTTAAAGAAGGCGATCAGGTTTCTTCTTGTGAGGTAAATATGATGTACAGAACA    13722
Pachon8     GAAATTGTTAAAGAAGGCGATCAGGTTTCTTCTTGTGAGGTAAATATGATGTACAGAACA    13722
Pachon15    GAAATTGTTAAAGAAGGCGATCAGGTTTCTTCTTGTGAGGTAAATATGATGTACAGAACA    13722
             *** ***** ******* **** ************************************

Rascon4     CAAAACATAATCTGGATGTTCATCCAAGACTACTCTTAACACAGTTTAGAGACAGTCCTG    13728
Surface     CAAAACATAATCTGGATGTTCATCCAAGACTACTCTTAACACAGTTTAGAGACAGTCCTG    13803
Rascon8     CAAAACATAATCTGGATGTTCATCCAAGACTACTCTTAACACAGTTTAGAGACAGTCCTG    13780
Rascon2     CAAAACATAATCTGGATGTTCATCCAAGACTACTCTTAACACAGTTTAGAGACAGTCCTG    13785
Rascon15    CAAAACATAATCTGGATGTTCATCCAAGACTACTCTTAACACAGTTTAGAGACAGTCCTG    13786
Rascon13    CAAAACATAATCTGGATGTTCATCCAAGACTACTCTTAACACAGTTTAGAGACAGTCCTG    13794
Rascon6     CAAAACATAATCTGGATGTTCATCCAAGACTACTCTTAACACAGTTTAGAGACAGTCCTG    13800
Pachon14    CAAAACATAATCTGGATGTTCATCCAAGACTACTCTTAACACAGTTTAGAGACAGTCCTG    13775
Pachon9     CAAAACATAATCTGGATGTTCATCCAAGACTACTCTTAACACAGTTTAGAGACAGTCCTG    13784
Pachon17    CAAAACATAATCTGGATGTTCATCCAAGACTACTCTTAACACAGTTTAGAGACAGTCCTG    13782
Pachon12    CAAAACATAATCTGGATGTTCATCCAAGACTACTCTTAACACAGTTTAGAGACAGTCCTG    13764
Pachon11    CAAAACATAATCTGGATGTTCATCCAAGACTACTCTTAACACAGTTTAGAGACAGTCCTG    13781
```

```
Pachon7      CAAAACATAATCTGGATGTTCATCCAAGACTACTCTTAACACAGTTTAGAGACAGTCCTG   13782
Pachon3      CAAAACATAATCTGGATGTTCATCCAAGACTACTCTTAACACAGTTTAGAGACAGTCCTG   13782
Pachon8      CAAAACATAATCTGGATGTTCATCCAAGACTACTCTTAACACAGTTTAGAGACAGTCCTG   13782
Pachon15     CAAAACATAATCTGGATGTTCATCCAAGACTACTCTTAACACAGTTTAGAGACAGTCCTG   13782
             ************************************************************

Rascon4      AAATGCACAGATGACACCGAAGGACTTCCTCAGCCATAGATTACCATGCAGTTAAATTCT   13788 Grp
Surface      AAATGCACAGATGACACCGAAGGACTTCCTCAGCCATAGATTACCATGCAGTTACGTTAT   13863 Intron1
Rascon8      AAATGCACAGATGACACCGAAGGACTTCCTCAGCCATAGATTACCATGCAGTTAAGTTCT   13840
Rascon2      AAATGCACAGATGACACCGAAGGACTTCCTCAGCCATAGATTACCATGCAGTTACGTTCT   13845
Rascon15     AAATGCACAGATGACACCGAAGGACTTCCTCAGCCATAGATTACCATGCAGTTAAGTTCT   13846
Rascon13     AAATGCACAGATGACACCGAAGGACTTCCTCAGCCATAGATTACCATGCAGTTAAATTCT   13854
Rascon6      AAATGCACAGATGACACCGAAGGACTTCCTCAGCCATAGATTACCATGCAGTTACATTCT   13860
Pachon14     AAATGCACAGATGACACCGAAGGACTTCCTCAGCCATAGATTACCATGCAGTTACATTCT   13835
Pachon9      AAATGCACAGATGACACCGAAGGACTTCCTCAGCCATAGATTACCATGCAGTTACATTCT   13844
Pachon17     AAATGCACAGATGACACCGAAGGACTTCCTCAGCCATAGATTACCATGCAGTTACATTCT   13842
Pachon12     AAATGCACAGATGACACCGAAGGACTTCCTCAGCCATAGATTACCATGCAGTTACATTCT   13824
Pachon11     AAATGCACAGATGACACCGAAGGACTTCCTCAGCCATAGATTACCATGCAGTTACATTCT   13841
Pachon7      AAATGCACAGATGACACCGAAGGACTTCCTCAGCCATAGATTACCATGCAGTTACATTCT   13842
Pachon3      AAATGCACAGATGACACCGAAGGACTTCCTCAGCCATAGATTACCATGCAGTTACATTCT   13842
Pachon8      AAATGCACAGATGACACCGAAGGACTTCCTCAGCCATAGATTACCATGCAGTTACATTCT   13842
Pachon15     AAATGCACAGATGACACCGAAGGACTTCCTCAGCCATAGATTACCATGCAGTTACATTCT   13842
             ******************************************************  ** *

Rascon4      TGAAAACATTAAGGACTAGCAAATTCCACATGAAGGTCTGTCACGGAAATATGATACATG   13848
Surface      TGAAAACATTAAGGACTAGCAAATTCCACATGAAGGTCTGTCACGGAAATATGATACATG   13923
Rascon8      TGAAAACATTAAGGACTAGCAAATTACACATGAAGGTCTGTCACGGAAATATGATACATG   13900
Rascon2      TGAAAACATTAAGGACTAGCAAATTCCACATGAAGGTCTGTCACGGAAACATGATACATG   13905
Rascon15     TGAAAACATTAAGGACTAGCAAATTCCACATGAAGGTCTGTCACGGAAATATGATACATG   13906
Rascon13     TGAAAACATTAAGGACTAGCAAATTCCACATGAAGGTCTGTCACGGAAATATGATACATG   13914
Rascon6      TGAAAACATTAAGGACTAGCAAATTCCACATGAAGGTCTGTCACGGAAATATGATACATG   13920
Pachon14     TGAAAACATTAAGGACTAGCAAATTCCACATGAAGGTCTGTCACGGAAATATGATACATG   13895
Pachon9      TGAAAACATTAAGGACTAGCAAATTCCACATGAAGGTCTGTCACGGAAATATGATACATG   13904
Pachon17     TGAAAACATTAAGGACTAGCAAATTCCACATGAAGGTCTGTCACGGAAATATGATACATG   13902
Pachon12     TGAAAACATTAAGGACTAGCAAATTCCACATGAAGGTCTGTCACGGAAATATGATACATG   13884
Pachon11     TGAAAACATTAAGGACTAGCAAATTCCACATGAAGGTCTGTCACGGAAATATGATACATG   13901
Pachon7      TGAAAACATTAAGGACTAGCAAATTCCACATGAAGGTCTGTCACGGAAATATGATACATG   13902
Pachon3      TGAAAACATTAAGGACTAGCAAATTCCACATGAAGGTCTGTCACGGAAATATGATACATG   13902
Pachon8      TGAAAACATTAAGGACTAGCAAATTCCACATGAAGGTCTGTCACGGAAATATGATACATG   13902
Pachon15     TGAAAACATTAAGGACTAGCAAATTCCACATGAAGGTCTGTCACGGAAATATGATACATG   13902
             ************************* *********************** **********

Rascon4      TACATGTAATAGAGCTGCAACTTTTGATAATTTTGGTAGTCTACTAATCTATTTATTTAT   13908
Surface      TACATGTAATAGAGCTGCAACTTTTGATAATTTTGGTAGTCTACTAATCTATTTATTTAT   13983
Rascon8      TACATGTAATAGAGCTGCAACTTTTGATAATTTTGGTAGTCTACTAATCTATTTATTTAT   13960
Rascon2      TACATGTAATAGAGCTGCAACTTTTGATAATTTTGGTAGTCTACTAATCTATTTATTTAT   13965
Rascon15     TACATGTAATAGAGCTGCAACTTTTGATAATTTTGGTAGTCTACTAATCTATTTATTTAT   13966
Rascon13     TACATGTAATAGAGCTGCAACTTTTGATAATTTTGGTAGTCTACTAATCTATTTATTTAT   13974
Rascon6      TACATGTAATAGAGCTGCAACTTTTGATAATTTTGGTAGTCTACTAATCTATTTATTTAT   13980
Pachon14     TACATGTAATAGAGCTGCAACTTTTGATAATTTTGGTAGTCTACTAATCTATTTATTTAT   13955
Pachon9      TACATGTAATAGAGCTGCAACTTTTGATAATTTTGGTAGTCTACTAATCTATTTATTTAT   13964
Pachon17     TACATGTAATAGAGCTGCAACTTTTGATAATTTTGGTAGTCTACTAATCTATTTATTTAT   13962
Pachon12     TACATGTAATAGAGCTGCAACTTTTGATAATTTTGGTAGTCTACTAATCTATTTATTTAT   13944
Pachon11     TACATGTAATAGAGCTGCAACTTTTGATAATTTTGGTAGTCTACTAATCTATTTATTTAT   13961
Pachon7      TACATGTAATAGAGCTGCAACTTTTGATAATTTTGGTAGTCTACTAATCTATTTATTTAT   13962
Pachon3      TACATGTAATAGAGCTGCAACTTTTGATAATTTTGGTAGTCTACTAATCTATTTATTTAT   13962
Pachon8      TACATGTAATAGAGCTGCAACTTTTGATAATTTTGGTAGTCTACTAATCTATTTATTTAT   13962
Pachon15     TACATGTAATAGAGCTGCAACTTTTGATAATTTTGGTAGTCTACTAATCTATTTATTTAT   13962
             ************************************************************

Rascon4      TCAATAAGTCAAATAATCAACAATTATTTCTAACATGCACCTCCATCTATGCTTCCAGTG   13968
Surface      TCAATAAGTCAAATAATCAACAATTATTTCTAACATGCACCTCCATCTATGCTTCCAGTG   14043
Rascon8      TCAATAAGTCAAATAATCAACAATTATTTCTAACATGCACCTCCATCTATGCTTCCAGTG   14020
Rascon2      TCAATAAGTCAAATAATCAACAATTATTTCTAACATGCACCTCCATCTATGCTTCCAGTG   14025
Rascon15     TCAATAAGTCAAATAATCAACAATTATTTCTAACATGCACCTCCATCTATGCTTCCAGTG   14026
Rascon13     TCAATAAGTCAAATAATCAACAATTATTTCTAACATGCACCTCCATCTATGCTTCCAGTG   14034
Rascon6      TCAATAAGTCAAATAATCAACAATTATTTCTAACATGCACCTCCATCTATGCTTCCAGTG   14040
Pachon14     TCAATAAGTCAAATAATCAACAATTATTTCTAACATGCACCTCCATCTATGCTTCCAGTG   14015
Pachon9      TCAATAAGTCAAATAATCAACAATTATTTCTAACATGCACCTCCATCTATGCTTCCAGTG   14024
Pachon17     TCAATAAGTCAAATAATCAACAATTATTTCTAACATGCACCTCCATCTATGCTTCCAGTG   14022
Pachon12     TCAATAAGTCAAATAATCAACAATTATTTCTAACATGCACCTCCATCTATGCTTCCAGTG   14004
Pachon11     TCAATAAGTCAAATAATCAACAATTATTTCTAACATGCACCTCCATCTATGCTTCCAGTG   14021
Pachon7      TCAATAAGTCAAATAATCAACAATTATTTCTAACATGCACCTCCATCTATGCTTCCAGTG   14022
Pachon3      TCAATAAGTCAAATAATCAACAATTATTTCTAACATGCACCTCCATCTATGCTTCCAGTG   14022
Pachon8      TCAATAAGTCAAATAATCAACAATTATTTCTAACATGCACCTCCATCTATGCTTCCAGTG   14022
Pachon15     TCAATAAGTCAAATAATCAACAATTATTTCTAACATGCACCTCCATCTATGCTTCCAGTG   14022
             ************************************************************
```

```
Rascon4      GATCATTTTCCAAGCTTTCCCTATCGCTTCAAAACAATAGAAACAACTAAAATGTGATTT    14028 Grp
Surface      GATCCTTTTCCAAGCTTTCCCTATCGCTTCAAAACAATAGAAACAACTAAAATGTGATTT    14103 Intron1
Rascon8      GATCCTTTTCCAAGCTTTCCCTATCGCTTCAAAACAATAGAAACAACTAAAATGTGATTT    14080
Rascon2      GATCCTTTTCCAAGCTTTCCCTATCGCTTCAAAACAATAGAAACAACTAAAATGTGATTT    14085
Rascon15     GATCCTTTTCCAAGCTTTCCCTATCGCTTCAAAACAATAGAAACAACCTAAATGTGATTT    14086
Rascon13     GATCCTTTTCCAAGCTTTCCCTATCGCTTCAAAACAATAGAAACAACTAAAATGTGATTT    14094
Rascon6      GATCCTTTTCCAAGCTTTCCCTATCGCTTCAAAACAATAGAAACAACTAAAATGTGATTT    14100
Pachon14     GATCCTTTTCCAAGCTTTCCCTATCGCTTCAAAACAATAGAAACAACTAAAATGTGATTT    14075
Pachon9      GATCCTTTTCCAAGCTTTCCCTATCGCTTCAAAACAATAGAAACAACTAAAATGTGATTT    14084
Pachon17     GATCCTTTTCCAAGCTTTCCCTATCGCTTCAAAACAATAGAAACAACTAAAATGTGATTT    14082
Pachon12     GATCCTTTTCCAAGCTTTCCCTATCGCTTCAAAACAATAGAAACAACTAAAATGTGATTT    14064
Pachon11     GATCCTTTTCCAAGCTTTCCCTATCGCTTCAAAACAATAGAAACAACTAAAATGTGATTT    14081
Pachon7      GATCCTTTTCCAAGCTTTCCCTATCGCTTCAAAACAATAGAAACAACTAAAATGTGATTT    14082
Pachon3      GATCCTTTTCCAAGCTTTCCCTATCGCTTCAAAACAATAGAAACAACTAAAATGTGATTT    14082
Pachon8      GATCCTTTTCCAAGCTTTCCCTATCGCTTCAAAACAATAGAAACAACTAAAATGTGATTT    14082
Pachon15     GATCCTTTTCCAAGCTTTCCCTATCGCTTCAAAACAATAGAAACAACTAAAATGTGATTT    14082
             **** ******************************************  ***********

Rascon4      AAACATCCTTTAATTTAATTCACAGATGGTATGTAAATGTGAAATGTCATTTTACTGTAC    14088
Surface      AAACATCCTTTAATTTAATTCACAGATGGTATGTAAATGTGAAATGTCATTTTACTGTAC    14163
Rascon8      AAACATCCTTTAAGTTAATTCACAGATGGTATGTAAATGTGAAATGTCATTTTACTGTAC    14140
Rascon2      AAACATCCTTTAATTTAATTCACAGATGGTATGTAAATGTGAAATGTCATTTTACTGTAC    14145
Rascon15     AAACATCCTTTAATTTAATTCACAGATGGTATGTAAATGTGAAATGTCATTTTACTGTAC    14146
Rascon13     AAACATCCTTTAATTTAATTCACAGATGGTATGTAAATGTGAAATGTCATTTTACTGTAC    14154
Rascon6      AAACATCCTTTAATTTAATTCACAGATGGTATGTAAATGTGAAATGTCATTTTACTGTAC    14160
Pachon14     AAACATCCTTTAATTTAATTCACAGATGGTATGTAAATGTGAAATGTCATTTTACTGTAC    14135
Pachon9      AAACATCCTTTAATTTAATTCACAGATGGTATGTAAATGTGAAATGTCATTTTACTGTAC    14144
Pachon17     AAACATCCTTTAATTTAATTCACAGATGGTATGTAAATGTGAAATGTCATTTTACTGTAC    14142
Pachon12     AAACATCCTTTAATTTAATTCACAGATGGTATGTAAATGTGAAATGTCATTTTACTGTAC    14124
Pachon11     AAACATCCTTTAATTTAATTCACAGATGGTATGTAAATGTGAAATGTCATTTTACTGTAC    14141
Pachon7      AAACATCCTTTAATTTAATTCACAGATGGTATGTAAATGTGAAATGTCATTTTACTGTAC    14142
Pachon3      AAACATCCTTTAATTTAATTCACAGATGGTATGTAAATGTGAAATGTCATTTTACTGTAC    14142
Pachon8      AAACATCCTTTAATTTAATTCACAGATGGTATGTAAATGTGAAATGTCATTTTACTGTAC    14142
Pachon15     AAACATCCTTTAATTTAATTCACAGATGGTATGTAAATGTGAAATGTCATTTTACTGTAC    14142
             ************* **********************************************

Rascon4      AATATATGGGGACATTACCTCAATTTTGGCTGACCTCCATATTTTCTGTTGTGTTTAGGT    14148
Surface      AATATATGGGGACATTACCTCAATTTTGGCTGACCTCCATATTTTCTGTTGTGTTTAGGT    14223
Rascon8      AATATATGGGGACATTACCTCAATTTTGGCTGACCTCCATAGTTTCTGTTGTGTTTAGGT    14200
Rascon2      AATATATGGGGACATTACCTCAATTTTGGCTGACCTCCATATTTTCTGTTGTGTTTAGGT    14205
Rascon15     AATATATGGGGACATTACCTCAATTTTGGCTGACCTCCATATTTTCTGTTGTGTTTAGGT    14206
Rascon13     AATATATGGGGACATTACCTCAATTTTGGCTGACCTCCATATTTTCTGTTGTGTTTAGGT    14214
Rascon6      AATATATGGGGACATTACCTCAATTTTGGCTGACCTCCATATTTTCTGTTGTGTTTAGGT    14220
Pachon14     AATATATGGGGACATTACCTCAATTTTGGCTGACCTCCATATTTTCTGTTGTGTTTAGGT    14195
Pachon9      AATATATGGGGACATTACCTCAATTTTGGCTGACCTCCATATTTTCTGTTGTGTTTAGGT    14204
Pachon17     AATATATGGGGACATTACCTCAATTTTGGCTGACCTCCATATTTTCTGTTGTGTTTAGGT    14202
Pachon12     AATATATGGGGACATTACCTCAATTTTGGCTGACCTCCATATTTTCTGTTGTGTTTAGGT    14184
Pachon11     AATATATGGGGACATTACCTCAATTTTGGCTGACCTCCATATTTTCTGTTGTGTTTAGGT    14201
Pachon7      AATATATGGGGACATTACCTCAATTTTGGCTGACCTCCATATTTTCTGTTGTGTTTAGGT    14202
Pachon3      AATATATGGGGACATTACCTCAATTTTGGCTGACCTCCATATTTTCTGTTGTGTTTAGGT    14202
Pachon8      AATATATGGGGACATTACCTCAATTTTGGCTGACCTCCATATTTTCTGTTGTGTTTAGGT    14202
Pachon15     AATATATGGGGACATTACCTCAATTTTGGCTGACCTCCATATTTTCTGTTGTGTTTAGGT    14202
             ***************************************** ******************

Rascon4      GAATGTTACTTTATATAAGAATGTACAGTACCAGTCAAAAGTTAAGACACACATCCTATA    14208
Surface      GAATGTTACTTTATATAAGAATGTACAGTACCAGTCAAAAGTTAAGACACACATCCTATA    14283
Rascon8      GAATGTTACTTTATAAAAGAATGTACAGTACCAGTCAAAAGTTAAGACACACATCCTATA    14260
Rascon2      GAATGTTACTTTATATAAGAATGTACAGTACCAGTCAAAAGTTAAGACACACATCCTATA    14265
Rascon15     GAATGTTACTTTATATAAGAATGTACAGTACCAGTCAAAAGTTAAGACACACATCCTATA    14266
Rascon13     GAATGTTACTTTATATAAGAATGTACAGTACCAGTCAAAAGTTAAGACACACATCCTATA    14274
Rascon6      GAATGTTACTTTATATAAGAATGTACAGTACCAGTCAAAAGTTAAGACACACATCCTATA    14280
Pachon14     GAATGTTACTTTATATAAGAATGTACAGTACCAGTCAAAAGTTAAGACACACATCCTATA    14255
Pachon9      GAATGTTACTTTATATAAGAATGTACAGTACCAGTCAAAAGTTAAGACACACATCCTATA    14264
Pachon17     GAATGTTACTTTATATAAGAATGTACAGTACCAGTCAAAAGTTAAGACACACATCCTATA    14262
Pachon12     GAATGTTACTTTATATAAGAATGTACAGTACCAGTCAAAAGTTAAGACACACATCCTATA    14244
Pachon11     GAATGTTACTTTATATAAGAATGTACAGTACCAGTCAAAAGTTAAGACACACATCCTATA    14261
Pachon7      GAATGTTACTTTATATAAGAATGTACAGTACCAGTCAAAAGTTAAGACACACATCCTATA    14262
Pachon3      GAATGTTACTTTATATAAGAATGTACAGTACCAGTCAAAAGTTAAGACACACATCCTATA    14262
Pachon8      GAATGTTACTTTATATAAGAATGTACAGTACCAGTCAAAAGTTAAGACACACATCCTATA    14262
Pachon15     GAATGTTACTTTATATAAGAATGTACAGTACCAGTCAAAAGTTAAGACACACATCCTATA    14262
             *************** ********************************************

Rascon4      TAAGAACACATGCAGAAGTGTGTGGATTGTGTGCATCAAAAACTGTATTTATCACTGTCA    14268
Surface      TAAGAACACATGCAGAAGTGTGTGGATTGTGTGCATCAAAAACTGTATTTATCACTGTCA    14343
Rascon8      TAAGAACACATGCAGAAGTGTGTGGATTGTGTGCATCAAAAACTGTATTTATCACTGTCA    14320
Rascon2      TAAGAACACATGCAGAAGTGTGTGGATTGTGTGCATCAAAAACTGTATTTATCACTGTCA    14325
```

```
Rascon15      TAAGAACACATGCAGAAGTGTGTGGATTGTGTGCATCAAAAACTGTATTTATCACTGTCA      14326 Grp
Rascon13      TAAGAACACATGCAGAAGTGTGTGGATTGTGTGCATCAAAAACTGTATTTATCACTGTCA      14334 Intron1
Rascon6       TAAGAACACATGCAGAAGTGTGTGGATTGTGTGCATCAAAAACTGTATTTATCACTGTCA      14340
Pachon14      TAAGAACACATGCAGAAGTGTGTGGATTGTGTGCATCAAAAACTGTATTTATCACTGTCA      14315
Pachon9       TAAGAACACATGCAGAAGTGTGTGGATTGTGTGCATCAAAAACTGTATTTATCACTGTCA      14324
Pachon17      TAAGAACACATGCAGAAGTGTGTGGATTGTGTGCATCAAAAACTGTATTTATCACTGTCA      14322
Pachon12      TAAGAACACATGCAGAAGTGTGTGGATTGTGTGCATCAAAAACTGTATTTATCACTGTCA      14304
Pachon11      TAAGAACACATGCAGAAGTGTGTGGATTGTGTGCATCAAAAACTGTATTTATCACTGTCA      14321
Pachon7       TAAGAACACATGCAGAAGTGTGTGGATTGTGTGCATCAAAAACTGTATTTATCACTGTCA      14322
Pachon3       TAAGAACACATGCAGAAGTGTGTGGATTGTGTGCATCAAAAACTGTATTTATCACTGTCA      14322
Pachon8       TAAGAACACATGCAGAAGTGTGTGGATTGTGTGCATCAAAAACTGTATTTATCACTGTCA      14322
Pachon15      TAAGAACACATGCAGAAGTGTGTGGATTGTGTGCATCAAAAACTGTATTTATCACTGTCA      14322
              ************************************************************

Rascon4       CACAAATCCCTTATCTCCAAAAATCTCCAAAATAGCAGAAAAAGAAATCTTAATTTCAAT      14328
Surface       CACAAATCCCTTATCTCCAAAAATCTCCAAAATAGCAGAAAAAGAAATCTTAATTTCAAT      14403
Rascon8       CACAAATCCCTTATCTCCAAAAATCTCCAAAATAGCAGAAAAAGAAATCTTAATTTCAAT      14380
Rascon2       CACAAATCCCTTTTCTCCAAAAATCTCCAAAATAGCAGAAAAAGAAATCTTAATTTCAAT      14385
Rascon15      CACAAATCCCTTATCTCCAAAAATCTCCAAAATAGCAGAAAAAGAAATCTTAATTTCAAT      14386
Rascon13      CACAAATCCCTTATCTCCAAAAATCTCCAAAATAGCAGAAAAAGAAATCTTAATTTCAAT      14394
Rascon6       CACAAATCCCTTATCTCCAAAAATCTCCAAAATAGCAGAAAAAGAAATCTTAATTTCAAT      14400
Pachon14      CACAAATCCCTTATCTCCAAAAATCTCCAAAATAGCAGAAAAAGAAATCTTAATTTCAAT      14375
Pachon9       CACAAATCCCTTATCTCCAAAAATCTCCAAAATAGCAGAAAAAGAAATCTTAATTTCAAT      14384
Pachon17      CACAAATCCCTTATCTCCAAAAATCTCCAAAATAGCAGAAAAAGAAATCTTAATTTCAAT      14382
Pachon12      CACAAATCCCTTATCTCCAAAAATCTCCAAAATAGCAGAAAAAGAAATCTTAATTTCAAT      14364
Pachon11      CACAAATCCCTTATCTCCAAAAATCTCCAAAATAGCAGAAAAAGAAATCTTAATTTCAAT      14381
Pachon7       CACAAATCCCTTATCTCCAAAAATCTCCAAAATAGCAGAAAAAGAAATCTTAATTTCAAT      14382
Pachon3       CACAAATCCCTTATCTCCAAAAATCTCCAAAATAGCAGAAAAAGAAATCTTAATTTCAAT      14382
Pachon8       CACAAATCCCTTATCTCCAAAAATCTCCAAAATAGCAGAAAAAGAAATCTTAATTTCAAT      14382
Pachon15      CACAAATCCCTTATCTCCAAAAATCTCCAAAATAGCAGAAAAAGAAATCTTAATTTCAAT      14382
              ************ ***********************************************

Rascon4       AAAATTTCATGTAAAAACAATGTCATTTTCAAGCATTTCTATTTATCTACAGAATTTTGA      14388
Surface       AAAATTTCATGTAAAAACAATGTCATTTTCAAGCATTTCTATTTATCTACAGAATTGTGA      14463
Rascon8       AAAATTTCATGTAAAAACAATGTCATTTTCAAGCATTTCTATTTATCTACAGAATTTTGA      14440
Rascon2       AAAATTTCATGTAAAAACAATGTCATTTTCAAGCATTTCTATTTATCTACAGAATTTTGA      14445
Rascon15      AAAATTTCATGTAAAAACAATGTCATTTTCAAGCATTTCTATTTATCTACAGAATTTTGA      14446
Rascon13      AAAATTTCATGTAAAAACAATGTCATTTTCAAGCATTTCTATTTATCTACAGAATTTTGA      14454
Rascon6       AAAATTTCATGTAAAAACAATGTCATTTTCAAGCATTTCTATTTATCTACAGAATTTTGA      14460
Pachon14      AAAATTTCATGTAAAAACAATGTCATTTTCAAGCATTTCTATTTATCTACAGAATTTTGA      14435
Pachon9       AAAATTTCATGTAAAAACAATGTCATTTTCAAGCATTTCTATTTATCTACAGAATTTTGA      14444
Pachon17      AAAATTTCATGTAAAAACAATGTCATTTTCAAGCATTTCTATTTATCTACAGAATTTTGA      14442
Pachon12      AAAATTTCATGTAAAAACAATGTCATTTTCAAGCATTTCTATTTATCTACAGAATTTTGA      14424
Pachon11      AAAATTTCATGTAAAAACAATGTCATTTTCAAGCATTTCTATTTATCTACAGAATTTTGA      14441
Pachon7       AAAATTTCATGTAAAAACAATGTCATTTTCAAGCATTTCTATTTATCTACAGAATTTTGA      14442
Pachon3       AAAATTTCATGTAAAAACAATGTCATTTTCAAGCATTTCTATTTATCTACAGAATTTTGA      14442
Pachon8       AAAATTTCATGTAAAAACAATGTCATTTTCAAGCATTTCTATTTATCTACAGAATTTTGA      14442
Pachon15      AAAATTTCATGTAAAAACAATGTCATTTTCAAGCATTTCTATTTATCTACAGAATTTTGA      14442
              ******************************************************** ***

Rascon4       CTTAGTGTAACCAAACGTTGACTTCATAGAAATAATTCAGTAAAAATAAGGATCAAATGT      14448
Surface       CTTAGTGTAACCAAACGTTGACTTCATAGAAATAATTCAGTAAAAATAAGGATAAAATGT      14523
Rascon8       CTTAGTGTAACCAAACGTTGACTTCATAGAAATAATTCAGTAAAAATAAGGATAAAATGT      14500
Rascon2       CTTAGTGTAACCAAACGTTGACTTCATAGAAATAATTCAGTAAAAATAAGGATCAAATGT      14505
Rascon15      CTTAGTGTAACCAAACGTTGACTTCATAGAAATAATTCAGTAAAAATAAGGATTAAATGT      14506
Rascon13      CTTAGTGTAACCAAACGTTGACTTCATAGAAATAATTCAGTAAAAATAAGGATAAAATGT      14514
Rascon6       CTTAGTGTAACCAAACGTTGACTTCATAGAAATAATTCAGTAAAAATAAGGATTAAATGT      14520
Pachon14      CTTAGTGTAACCAAACGTTGACTTCATAGAAATAATTCAGTAAAAATAAGGATAAAATGT      14495
Pachon9       CTTAGTGTAACCAAACGTTGACTTCATAGAAATAATTCAGTAAAAATAAGGATAAAATGT      14504
Pachon17      CTTAGTGTAACCAAACGTTGACTTCATAGAAATAATTCAGTAAAAATAAGGATAAAATGT      14502
Pachon12      CTTAGTGTAACCAAACGTTGACTTCATAGAAATAATTCAGTAAAAATAAGGATAAAATGT      14484
Pachon11      CTTAGTGTAACCAAACGTTGACTTCATAGAAATAATTCAGTAAAAATAAGGATAAAATGT      14501
Pachon7       CTTAGTGTAACCAAACGTTGACTTCATAGAAATAATTCAGTAAAAATAAGGATAAAATGT      14502
Pachon3       CTTAGTGTAACCAAACGTTGACTTCATAGAAATAATTCAGTAAAAATAAGGATAAAATGT      14502
Pachon8       CTTAGTGTAACCAAACGTTGACTTCATAGAAATAATTCAGTAAAAATAAGGATAAAATGT      14502
Pachon15      CTTAGTGTAACCAAACGTTGACTTCATAGAAATAATTCAGTAAAAATAAGGATAAAATGT      14502
              ***************************************************** ******

Rascon4       TTTTGTGAAACAGCGTCAATATAATCACCAACCTCTGATACTTTTGATGTCCTGCCACTT      14508
Surface       TTTTGTGAAACAGCGACAATATAATCACCAACCTCTGATACTTTTGATGTCCTGCCACTT      14583
Rascon8       TTTTGTGAAACAGCGTCAATATAATCACCAACCTCTGATACTTTTGATGTCCTGCCACTT      14560
Rascon2       TTTTGTGAAACAGCGTCAATATAATCACCAACCTCTGATACTTTTGATGTCCTGCCACTT      14565
Rascon15      TTTTGTGAAACAGCGTCAATATAATCACCAACCTCTGATATTTTTGATGTCCTGCCACTT      14566
Rascon13      TTTTGTGAAACAGCGTCAATATAATCACCAACCTCTGATACTTTTGATGTCCTGCCACTT      14574
Rascon6       TTTTGTGAAACAGCGTCAATATAATCACCAACCTCTGATACTTTTGATGTCCTGCCACTT      14580
Pachon14      TTTTGTGAAACAGCGTCAATATAATCACCAACCTCTGATACTTTTGATGTCCTGCCACTT      14555
Pachon9       TTTTGTGAAACAGCGTCAATATAATCACCAACCTCTGATACTTTTGATGTCCTGCCACTT      14564
```

```
Pachon17     TTTTGTGAAACAGCGTCAATATAATCACCAACCTCTGATACTTTTGATGTCCTGCCACTT     14562
Pachon12     TTTTGTGAAACAGCGTCAATATAATCACCAACCTCTGATACTTTTGATGTCCTGCCACTT     14544
Pachon11     TTTTGTGAAACAGCGTCAATATAATCACCAACCTCTGATACTTTTGATGTCCTGCCACTT     14561
Pachon7      TTTTGTGAAACAGCGTCAATATAATCACCAACCTCTGATACTTTTGATGTCCTGCCACTT     14562
Pachon3      TTTTGTGAAACAGCGTCAATATAATCACCAACCTCTGATACTTTTGATGTCCTGCCACTT     14562
Pachon8      TTTTGTGAAACAGCGTCAATATAATCACCAACCTCTGATACTTTTGATGTCCTGCCACTT     14562
Pachon15     TTTTGTGAAACAGCGTCAATATAATCACCAACCTCTGATACTTTTGATGTCCTGCCACTT     14562
             *************** *********************** ********************

Rascon4      AATACAATTGTGATTAAGTTAATGTGTGTTCTTTTCTAAGCAGATGGAAATTACTGTCAT     14568 Grp
Surface      AATACAATTGTGATTAAGTTAATGTGTGTTCTTTTCTAAGCAGATGGAAATTACTGTCAT     14643 Intron1
Rascon8      AATACAATTGTGATTAAGTTAATGTGTGTTCTTTTCTAAGCAGATGGAAATTACTGTCAT     14620
Rascon2      AATACAATTGTGATTAAGTTAATGTGTGTTCTTTTCTAAGCAGATGGAAATTACTGTCAT     14625
Rascon15     AATACAATTGTGATTAAGTTAATGTGTGTTCTTTTCTAAGCAGATGGAAATTACTGTCAT     14626
Rascon13     AATACAATTGTGATTAAGTTAATGTGTGTTCTTTTCTAAGCAGATGGAAATTACTGTCAT     14634
Rascon6      AATACAATTGTGATTAAGTTAATGTGTGTTCTTTTCTAAGCAGATGGAAATTACTGTCAT     14640
Pachon14     AATACAATTGTGATTAAGTTAATGTGTGTTCTTTTCTAAGCAGATGGAAATTACTGTCAT     14615
Pachon9      AATACAATTGTGATTAAGTTAATGTGTGTTCTTTTCTAAGCAGATGGAAATTACTGTCAT     14624
Pachon17     AATACAATTGTGATTAAGTTAATGTGTGTTCTTTTCTAAGCAGATGGAAATTACTGTCAT     14622
Pachon12     AATACAATTGTGATTAAGTTAATGTGTGTTCTTTTCTAAGCAGATGGAAATTACTGTCAT     14604
Pachon11     AATACAATTGTGATTAAGTTAATGTGTGTTCTTTTCTAAGCAGATGGAAATTACTGTCAT     14621
Pachon7      AATACAATTGTGATTAAGTTAATGTGTGTTCTTTTCTAAGCAGATGGAAATTACTGTCAT     14622
Pachon3      AATACAATTGTGATTAAGTTAATGTGTGTTCTTTTCTAAGCAGATGGAAATTACTGTCAT     14622
Pachon8      AATACAATTGTGATTAAGTTAATGTGTGTTCTTTTCTAAGCAGATGGAAATTACTGTCAT     14622
Pachon15     AATACAATTGTGATTAAGTTAATGTGTGTTCTTTTCTAAGCAGATGGAAATTACTGTCAT     14622
             ************************************************************

Rascon4      TAACTGACGATACACTCAGATCCTTCTTTCAAAATAAAGTGGTTTGAAATCACCCTAAAA     14628
Surface      TAACTGACGATACACTCAGATCCTTCTTTCAAAATAAAGTGGTTTGAAATCACCCTAAAA     14703
Rascon8      TAACTGACGATACACTCAGATCCTTCTTTCAAAATAAAGTGGTTTGAAATCACCCTAAAA     14680
Rascon2      TAACTGACGATACACTCAGATCCTTCTTTCAAAATAAAGTGGTTTGAAATCACCCTAAAA     14685
Rascon15     TAACTGACGATACACTCAGATCCTTCTTTCAAAATAAAGTGGTTTGAAATCACCCTAAAA     14686
Rascon13     TAACTGACGATACACTCAGATCCTTCTTTCAAAATAAAGTGGTTTGAAATCACCCTAAAA     14694
Rascon6      TAACTGACGATACACTCAGATCCTTCTTTCAAAATAAAGTGGTTTGAAATCACCCTAAAA     14700
Pachon14     TAACTGACGATACACTCAGATCCTTCTTTCAAAATAAAGTGGTTTGAAATCACCCTAAAA     14675
Pachon9      TAACTGACGATACACTCAGATCCTTCTTTCAAAATAAAGTGGTTTGAAATCACCCTAAAA     14684
Pachon17     TAACTGACGATACACTCAGATCCTTCTTTCAAAATAAAGTGGTTTGAAATCACCCTAAAA     14682
Pachon12     TAACTGACGATACACTCAGATCCTTCTTTCAAAATAAAGTGGTTTGAAATCACCCTAAAA     14664
Pachon11     TAACTGACGATACACTCAGATCCTTCTTTCAAAATAAAGTGGTTTGAAATCACCCTAAAA     14681
Pachon7      TAACTGACGATACACTCAGATCCTTCTTTCAAAATAAAGTGGTTTGAAATCACCCTAAAA     14682
Pachon3      TAACTGACGATACACTCAGATCCTTCTTTCAAAATAAAGTGGTTTGAAATCACCCTAAAA     14682
Pachon8      TAACTGACGATACACTCAGATCCTTCTTTCAAAATAAAGTGGTTTGAAATCACCCTAAAA     14682
Pachon15     TAACTGACGATACACTCAGATCCTTCTTTCAAAATAAAGTGGTTTGAAATCACCCTAAAA     14682
             ************************************************************

Rascon4      TA---------TCCTTCATTTTTTCAGGATTCTACTCTCTTATAAACATCATTCCCTGTC     14679
Surface      TATTTATGCTGTCCTTCATTTTTTCAGGATTCTACTCTCTTATAAACGTCATTCCCTGTC     14763
Rascon8      TATTTATGCTGTCCTTCATTTTTTCAGGATTCTACTCTCTTATAAACGTCATTCCCTGTC     14740
Rascon2      TATTTATGCTGTCCTTCATTTTTTCAGGATTCTACTCTCTTATAAACATCATTCCCTGTC     14745
Rascon15     TA---------TCCTTCATTTTTTCAGGATTCTACTCTCTTATAAACGTCATTCCCTGTC     14737
Rascon13     TATTTATGCTGTCCTTCATTTTTTCAGGATTCTACTCTCTTATAAACGTCATTCCCTGTC     14754
Rascon6      TATTTATGCTGTCCTTCATTTTTTCAGGATTCTACTCTCTTATAAACGTCATTCCCTGTC     14760
Pachon14     TATTTATGCTGTCCTTCATTTTTTCAGGATTCTACTCTCTTATAAACGTCATTCCCTGTC     14735
Pachon9      TATTTATGCTGTCCTTCATTTTTTCAGGATTCTACTCTCTTATAAACGTCATTCCCTGTC     14744
Pachon17     TATTTATGCTGTCCTTCATTTTTTCAGGATTCTACTCTCTTATAAACGTCATTCCCTGTC     14742
Pachon12     TATTTATGCTGTCCTTCATTTTTTCAGGATTCTACTCTCTTATAAACGTCATTCCCTGTC     14724
Pachon11     TATTTATGCTGTCCTTCATTTTTTCAGGATTCTACTCTCTTATAAACGTCATTCCCTGTC     14741
Pachon7      TATTTATGCTGTCCTTCATTTTTTCAGGATTCTACTCTCTTATAAACGTCATTCCCTGTC     14742
Pachon3      TATTTATGCTGTCCTTCATTTTTTCAGGATTCTACTCTCTTATAAACGTCATTCCCTGTC     14742
Pachon8      TATTTATGCTGTCCTTCATTTTTTCAGGATTCTACTCTCTTATAAACGTCATTCCCTGTC     14742
Pachon15     TATTTATGCTGTCCTTCATTTTTTCAGGATTCTACTCTCTTATAAACGTCATTCCCTGTC     14742
             **         *********************************** *************

Rascon4      ATACCTACAATTTCATGTTTAAAGCCAGACAGCGTGCCCCGACCTTTTTAGTTCTACCAA     14739
Surface      ATACCTACAATTTCATGTTTAAAGCCAGACAGCGTGCCCCGACCTTTTTAGTTCTACAAA     14823
Rascon8      ATACCTACAATTTCATGTTTAAAGCCAGACAGCGTGCCCCGACCTTTTTAGTTCTACCAA     14800
Rascon2      ATACCTACAATTTCATGTTTAAAGCCAGACAGCGTGCCCCGACCTTTTTAGTTCTACCAA     14805
Rascon15     ATACCTACAATTTCATGTTTAAAGCCAGACAGCATGCCCCGACCTTTTTAGTTCTACCAA     14797
Rascon13     ATACCTACAATTTCATGTTTAAAGCCAGACAGCGTGCCCCGACCTTTTTAGTTCTACCAA     14814
Rascon6      ATACCTACAATTTCATGTTTAAAGCCAGACAGCATGCCCCGACCTTTTTAGTTCTACCAA     14820
Pachon14     ATACCTACAATTTCATGTTTAAAGCCAGACAGCGTGCCCCGACCTTTTTAGTTCTACCAA     14795
Pachon9      ATACCTACAATTTCATGTTTAAAGCCAGACAGCGTGCCCCGACCTTTTTAGTTCTACCAA     14804
Pachon17     ATACCTACAATTTCATGTTTAAAGCCAGACAGCGTGCCCCGACCTTTTTAGTTCTACCAA     14802
Pachon12     ATACCTACAATTTCATGTTTAAAGCCAGACAGCGTGCCCCGACCTTTTTAGTTCTACCAA     14784
Pachon11     ATACCTACAATTTCATGTTTAAAGCCAGACAGCGTGCCCCGACCTTTTTAGTTCTACCAA     14801
Pachon7      ATACCTACAATTTCATGTTTAAAGCCAGACAGCGTGCCCCGACCTTTTTAGTTCTACCAA     14802
Pachon3      ATACCTACAATTTCATGTTTAAAGCCAGACAGCGTGCCCCGACCTTTTTAGTTCTACCAA     14802
```

```
Pachon8      ATACCTACAATTTCATGTTTAAAGCCAGACAGCGTGCCCCGACCTTTTTAGTTCTACCAA      14802
Pachon15     ATACCTACAATTTCATGTTTAAAGCCAGACAGCGTGCCCCGACCTTTTTAGTTCTACCAA      14802
             ********************************* *********************** **

Rascon4      CAGCTTCAAACACTCAGTCTCACAGGAAGCTTTTATATCAAACATAAGAGGTGCCATAAA      14799 Grp
Surface      CAGCTTCAAACACTCAGTCTCACAGGAAGCTTTTATATCAAACATAAGAGGTGCCACAAA      14883 Intron1
Rascon8      CAGCTTCAAACACTCAGTCTCACAGGAAGCTTTTATATCAAACATAAGAGGTGCCATAAA      14860
Rascon2      CAGCTTCAAACACTCAGTCTCACAGGAAGCTTTTATATCAAACATAAGAGGTGCCATAAA      14865
Rascon15     CAGCTTCAAACACTCAGTCTCACAGGAAGCTTTTATATCAAACATAAGAGGTGCCATAAA      14857
Rascon13     CAGCTTCAAACACTCTGTCTCACAGGAAGCTTTTATATCAAACATAAGAGGTGCCATAAA      14874
Rascon6      CAGCTTCAAACACTCAGTCTCACAGGAAGCTTTTATATCAAACATAAGAGGTGCCATAAA      14880
Pachon14     CAGCTTCAAACACTCAGTCTCACAGGAAGCTTTTATATCAAACATAAGAGGTGCCATAAA      14855
Pachon9      CAGCTTCAAACACTCAGTCTCACAGGAAGCTTTTATATCAAACATAAGAGGTGCCATAAA      14864
Pachon17     CAGCTTCAAACACTCAGTCTCACAGGAAGCTTTTATATCAAACATAAGAGGTGCCATAAA      14862
Pachon12     CAGCTTCAAACACTCAGTCTCACAGGAAGCTTTTATATCAAACATAAGAGGTGCCATAAA      14844
Pachon11     CAGCTTCAAACACTCAGTCTCACAGGAAGCTTTTATATCAAACATAAGAGGTGCCATAAA      14861
Pachon7      CAGCTTCAAACACTCAGTCTCACAGGAAGCTTTTATATCAAACATAAGAGGTGCCATAAA      14862
Pachon3      CAGCTTCAAACACTCAGTCTCACAGGAAGCTTTTATATCAAACATAAGAGGTGCCATAAA      14862
Pachon8      CAGCTTCAAACACTCAGTCTCACAGGAAGCTTTTATATCAAACATAAGAGGTGCCATAAA      14862
Pachon15     CAGCTTCAAACACTCAGTCTCACAGGAAGCTTTTATATCAAACATAAGAGGTGCCATAAA      14862
             *************** **************************************** ***

Rascon4      ATATGGAGTGAACACAGTGTTACTAAACACATTGGGGAATCAGAAATGGTCTCAGAGTCT      14859
Surface      ATATGGAGTGAACACAGTGTTACTAAACACATTGGGGAATCAGAAATGGTCTCAGAGTCT      14943
Rascon8      ATATGGAGTGAACACAGTGTTACTAAACACATTGGGGAATCAGAAATGGTCTCAGAGTCT      14920
Rascon2      ATATGGAGTGAACACAGTGTTACTAAACACATTGGGGAATCAGAAATGGTCTCAGAGTCT      14925
Rascon15     ATATGGAGTGAACACAGTGTTACTAAACACATTGGGGAATCAGAAATGGTCTCAGAGTCT      14917
Rascon13     ATATGGAGTGAACACAGTGTTACTAAACACATTGGGGAATCAGAAATGGTCTCAGAGTCT      14934
Rascon6      ATATGGAGTGAACACAGTGTTACTAAACACATTGGGGAATCAGAAATGGTCTCAGAGTCT      14940
Pachon14     ATATGGAGTGAACACAGTGTTACTAAACACATTGGGGAATCAGAAATGGTCTCAGAGTCT      14915
Pachon9      ATATGGAGTGAACACAGTGTTACTAAACACATTGGGGAATCAGAAATGGTCTCAGAGTCT      14924
Pachon17     ATATGGAGTGAACACAGTGTTACTAAACACATTGGGGAATCAGAAATGGTCTCAGAGTCT      14922
Pachon12     ATATGGAGTGAACACAGTGTTACTAAACACATTGGGGAATCAGAAATGGTCTCAGAGTCT      14904
Pachon11     ATATGGAGTGAACACAGTGTTACTAAACACATTGGGGAATCAGAAATGGTCTCAGAGTCT      14921
Pachon7      ATATGGAGTGAACACAGTGTTACTAAACACATTGGGGAATCAGAAATGGTCTCAGAGTCT      14922
Pachon3      ATATGGAGTGAACACAGTGTTACTAAACACATTGGGGAATCAGAAATGGTCTCAGAGTCT      14922
Pachon8      ATATGGAGTGAACACAGTGTTACTAAACACATTGGGGAATCAGAAATGGTCTCAGAGTCT      14922
Pachon15     ATATGGAGTGAACACAGTGTTACTAAACACATTGGGGAATCAGAAATGGTCTCAGAGTCT      14922
             ************************************************************

Rascon4      ACTGCACACCAGCTCAGCTCCAATACAGCCATGGCTTACCGGAACCATCCAGAGTAGACA      14919
Surface      ACTGCACACCAGCTCAGCTCCAATACAGCCATGGCTTACCGGAACCATCCAGAGTAGACA      15003
Rascon8      ACTGCACACCAGCTCAGCTCCAATACAGCCATGGCTTACCGGAACCATCCAGAGTAGACA      14980
Rascon2      ACTGCACACCAGCTCAGCTCCAATACAGCCATGGCTTACCGGAACCATCCAGAGTAGACA      14985
Rascon15     ACTGCACACCAGCTCAGCTCCAATACAGCCATGGCTTACCGGAACCATCCAGAGTAGACA      14977
Rascon13     ACTGCACACCAGCTCAGCTCCAATACAGCCATGGCTTACCGGAACCATCCAGAGTAGACA      14994
Rascon6      ACTGCACACCAGCTCAGCTCCAATACAGCCATGGCTTACCGGAACCATCCAGAGTAGACA      15000
Pachon14     ACTGCACACCAGCTCAGCTCCAATACAGCCATGGCTTACCGGAACCATCCAGAGTAGACA      14975
Pachon9      ACTGCACACCAGCTCAGCTCCAATACAGCCATGGCTTACCGGAACCATCCAGAGTAGACA      14984
Pachon17     ACTGCACACCAGCTCAGCTCCAATACAGCCATGGCTTACCGGAACCATCCAGAGTAGACA      14982
Pachon12     ACTGCACACCAGCTCAGCTCCAATACAGCCATGGCTTACCGGAACCATCCAGAGTAGACA      14964
Pachon11     ACTGCACACCAGCTCAGCTCCAATACAGCCATGGCTTACCGGAACCATCCAGAGTAGACA      14981
Pachon7      ACTGCACACCAGCTCAGCTCCAATACAGCCATGGCTTACCGGAACCATCCAGAGTAGACA      14982
Pachon3      ACTGCACACCAGCTCAGCTCCAATACAGCCATGGCTTACCGGAACCATCCAGAGTAGACA      14982
Pachon8      ACTGCACACCAGCTCAGCTCCAATACAGCCATGGCTTACCGGAACCATCCAGAGTAGACA      14982
Pachon15     ACTGCACACCAGCTCAGCTCCAATACAGCCATGGCTTACCGGAACCATCCAGAGTAGACA      14982
             ************************************************************

Rascon4      AAATAGAGAAGATCTCATAAAGGAAAGGAGCTCCATGTCTGGAGGCAACAAACATGCACA      14979
Surface      AAATAGAGAAGATCTCATAAAGGAAAGGAGCTCCATGTCTGGAGGCAACAAACATGCACA      15063
Rascon8      AAATAGAGAAGATCTCATAAAGGAAAGGAGCTCCATGTCTGGAGGCAACAAACATGCACA      15040
Rascon2      AAATAGAGAAGATCTCATAAAGGAAAGGAGCTCCATGTCTGGAGGCAACAAACATGCACA      15045
Rascon15     AAATAGAGAAGATCTCATAAAGGAAAGGAGCTCCATGTCTGGAGGCAACAAACATGCACA      15037
Rascon13     AAATAGAGAAGATCTCATAAAGGAAAGGAGCTCCATGTCTGGAGGCAACAAACATGCACA      15054
Rascon6      AAATAGAGAAGATCTCATAAAGGAAAGAAGCTCCATGTCTGGAGGCAACAAACATGCACA      15060
Pachon14     AAATAGAGAAGATCTCATAAAGGAAAGGAGCTCCATGTCTGGAGGCAACAAACATGCACA      15035
Pachon9      AAATAGAGAAGATCTCATAAAGGAAAGGAGCTCCATGTCTGGAGGCAACAAACATGCACA      15044
Pachon17     AAATAGAGAAGATCTCATAAAGGAAAGGAGCTCCATGTCTGGAGGCAACAAACATGCACA      15042
Pachon12     AAATAGAGAAGATCTCATAAAGGAAAGGAGCTCCATGTCTGGAGGCAACAAACATGCACA      15024
Pachon11     AAATAGAGAAGATCTCATAAAGGAAAGGAGCTCCATGTCTGGAGGCAACAAACATGCACA      15041
Pachon7      AAATAGAGAAGATCTCATAAAGGAAAGGAGCTCCATGTCTGGAGGCAACAAACATGCACA      15042
Pachon3      AAATAGAGAAGATCTCATAAAGGAAAGGAGCTCCATGTCTGGAGGCAACAAACATGCACA      15042
Pachon8      AAATAGAGAAGATCTCATAAAGGAAAGGAGCTCCATGTCTGGAGGCAACAAACATGCACA      15042
Pachon15     AAATAGAGAAGATCTCATAAAGGAAAGGAGCTCCATGTCTGGAGGCAACAAACATGCACA      15042
             *************************** ********************************

Rascon4      GAACTTCATAATGTGTTTGGAAAGCCACCAAGCAACCGATACCCTATTATTTCTTAGTTC      15039
```

```
Surface      GAACTTCATAATGTGTTTGGAAAGCCACCAAGCAACCGATACCCTAATATTTCTTAGTTC   15123 Grp
Rascon8      GAACTTCATAATGTGTTTGGAAAGCCACCAAGCAACCGATACCCTATTATTTCTTAGTTC   15100 Intron1
Rascon2      GAACTTCATAATGTGTTTGGAAAGCCACCAAGCAACCGATACCCTATTATTTCTTAGTTC   15105
Rascon15     GAACTTCATAATGTGTTTGGAAAGCCACCAAGCAACCGATACCCTATTATTTCTTAGTTC   15097
Rascon13     GAACTTCATAATGTGTTTGGAAAGCCACCAAGCAACCGATACCCTATTATTTCTTAGTTC   15114
Rascon6      GAACTTCATAATGTGTTTGGAAAGCCACCAAGCAACCGATACCCTATTATTTCTTAGTTC   15120
Pachon14     GAACTTCATAATGTGTTTGGAAAGCCACCAAGCAACCGATACCCTATTATTTCTTAGTTC   15095
Pachon9      GAACTTCATAATGTGTTTGGAAAGCCACCAAGCAACCGATACCCTATTATTTCTTAGTTC   15104
Pachon17     GAACTTCATAATGTGTTTGGAAAGCCACCAAGCAACCGATACCCTATTATTTCTTAGTTC   15102
Pachon12     GAACTTCATAATGTGTTTGGAAAGCCACCAAGCAACCGATACCCTATTATTTCTTAGTTC   15084
Pachon11     GAACTTCATAATGTGTTTGGAAAGCCACCAAGCAACCGATACCCTATTATTTCTTAGTTC   15101
Pachon7      GAACTTCATAATGTGTTTGGAAAGCCACCAAGCAACCGATACCCTATTATTTCTTAGTTC   15102
Pachon3      GAACTTCATAATGTGTTTGGAAAGCCACCAAGCAACCGATACCCTATTATTTCTTAGTTC   15102
Pachon8      GAACTTCATAATGTGTTTGGAAAGCCACCAAGCAACCGATACCCTATTATTTCTTAGTTC   15102
Pachon15     GAACTTCATAATGTGTTTGGAAAGCCACCAAGCAACCGATACCCTATTATTTCTTAGTTC   15102
             ********************************************** *************

Rascon4      ATTTGGGTCCAGCATCAAATTTAGTATGATAAAAACTTATTAGGGTGAATTAATTTGTTT   15099
Surface      ATTTGGGTCCAGCATCAAATTTAGTATGATAAAAACTTATTAGGGTGAATTAATTTGTTT   15183
Rascon8      ATTTGGGTCCAGCATCAAATTTAGTATGATAAAAACTTATTAGGGTGAATTAATTTGTTT   15160
Rascon2      ATTTGGGTCCAGCATCAAATTTAGTATGATAAAAACTTATTAGGGTGAATTAATTTGTTT   15165
Rascon15     ATTTGGGTCCAGCATCAAATTTAGTATGATAAAAACTTATTAGGGTGAATTAATTTGTTT   15157
Rascon13     ATTTGGGTCCAGCATCAAATTTAGTATGATAAAAACTTATTAGGGTGAATTAATTTGTTT   15174
Rascon6      ATTTGGGTCCAGCATCAAATTTAGTATGATAAAAACTTATTAGGGTGAATTAATTTGTTT   15180
Pachon14     ATTTGGGTCCAGCATCAAATTTAGTATGATAAAAACTTATTAGGGTGAATTAATTTGTTT   15155
Pachon9      ATTTGGGTCCAGCATCAAATTTAGTATGATAAAAACTTATTAGGGTGAATTAATTTGTTT   15164
Pachon17     ATTTGGGTCCAGCATCAAATTTAGTATGATAAAAACTTATTAGGGTGAATTAATTTGTTT   15162
Pachon12     ATTTGGGTCCAGCATCAAATTTAGTATGATAAAAACTTATTAGGGTGAATTAATTTGTTT   15144
Pachon11     ATTTGGGTCCAGCATCAAATTTAGTATGATAAAAACTTATTAGGGTGAATTAATTTGTTT   15161
Pachon7      ATTTGGGTCCAGCATCAAATTTAGTATGATAAAAACTTATTAGGGTGAATTAATTTGTTT   15162
Pachon3      ATTTGGGTCCAGCATCAAATTTAGTATGATAAAAACTTATTAGGGTGAATTAATTTGTTT   15162
Pachon8      ATTTGGGTCCAGCATCAAATTTAGTATGATAAAAACTTATTAGGGTGAATTAATTTGTTT   15162
Pachon15     ATTTGGGTCCAGCATCAAATTTAGTATGATAAAAACTTATTAGGGTGAATTAATTTGTTT   15162
             ************************************************************

Rascon4      TGGTAAATACCAAAAGTAGTCTTGATTTATTAGATTTAGCCTAAATAGATATTTACATTT   15159
Surface      TGGTAAATACCAAAAGTAGTCTTGATTTATTAGATTTAGCCTAAATAGATATTTACATTT   15243
Rascon8      TGGTAAATACCAAAAGTAGTCTTGATTTATTAGATTTAGCCTAAATAGATATTTACATTT   15220
Rascon2      TGGTAAATACCAAAAGTAGTCTTGATTTATTAGATTTAGCCTAAATAGATATTTACATTT   15225
Rascon15     TGGTAAATACCAAAAGTAGTCTTGATTTATTAGATTTAGCCTAAATAGATATTTACATTT   15217
Rascon13     TGGTAAATACCAAAAGTAGTCTTGATTTATTAGATTTAGCCTAAATAGATATTTACATTT   15234
Rascon6      TGGTAAATACCAAAAGTAGTCTTGATTTATTAGATTTAGCCTAAATAGATATTTACATTT   15240
Pachon14     TGGTAAATACCAAAAGTAGTCTTGATTTATTAGATTTAGCCTAAATAGATATTTACATTT   15215
Pachon9      TGGTAAATACCAAAAGTAGTCTTGATTTATTAGATTTAGCCTAAATAGATATTTACATTT   15224
Pachon17     TGGTAAATACCAAAAGTAGTCTTGATTTATTAGATTTAGCCTAAATAGATATTTACATTT   15222
Pachon12     TGGTAAATACCAAAAGTAGTCTTGATTTATTAGATTTAGCCTAAATAGATATTTACATTT   15204
Pachon11     TGGTAAATACCAAAAGTAGTCTTGATTTATTAGATTTAGCCTAAATAGATATTTACATTT   15221
Pachon7      TGGTAAATACCAAAAGTAGTCTTGATTTATTAGATTTAGCCTAAATAGATATTTACATTT   15222
Pachon3      TGGTAAATACCAAAAGTAGTCTTGATTTATTAGATTTAGCCTAAATAGATATTTACATTT   15222
Pachon8      TGGTAAATACCAAAAGTAGTCTTGATTTATTAGATTTAGCCTAAATAGATATTTACATTT   15222
Pachon15     TGGTAAATACCAAAAGTAGTCTTGATTTATTAGATTTAGCCTAAATAGATATTTACATTT   15222
             ************************************************************

Rascon4      AAGACATGTGGTCAATATGATCAGTTTGATTAATTAAACATTTAGACAGAGTTAATAAGA   15219
Surface      AAGACATGTGGTCAATATGATCAGTTTGATTAATTAAACATTTAGACAGAGTTAATAAGA   15303
Rascon8      AAGACATGTGGTCAATATGATCAGTTTGATTAATTAAACATTTAGACAGAGTTAATAAGA   15280
Rascon2      AAGACATGTGGTCAATATGATCAGTTTGATTAATTAAACATTTAGACAGAGTTAATAAGA   15285
Rascon15     AAGACATGTGGTCAATATGATCAGTTTGATTAATTAAACATTTAGACAGAGTTAATAAGA   15277
Rascon13     AAGACATGTGGTCAATATGATCAGTTTGATTAATTAAACATTTAGACAGAGTTAATAAGA   15294
Rascon6      AAGACATGTGGTCAATATGATCAGTTTGATTAATTAAACATTTAGACAGAGTTAATAAGA   15300
Pachon14     AAGACATGTGGTCAATATGATCAGTTTGATTAATTAAACATTTAGACAGAGTTAATAAGA   15275
Pachon9      AAGACATGTGGTCAATATGATCAGTTTGATTAATTAAACATTTAGACAGAGTTAATAAGA   15284
Pachon17     AAGACATGTGGTCAATATGATCAGTTTGATTAATTAAACATTTAGACAGAGTTAATAAGA   15282
Pachon12     AAGACATGTGGTCAATATGATCAGTTTGATTAATTAAACATTTAGACAGAGTTAATAAGA   15264
Pachon11     AAGACATGTGGTCAATATGATCAGTTTGATTAATTAAACATTTAGACAGAGTTAATAAGA   15281
Pachon7      AAGACATGTGGTCAATATGATCAGTTTGATTAATTAAACATTTAGACAGAGTTAATAAGA   15282
Pachon3      AAGACATGTGGTCAATATGATCAGTTTGATTAATTAAACATTTAGACAGAGTTAATAAGA   15282
Pachon8      AAGACATGTGGTCAATATGATCAGTTTGATTAATTAAACATTTAGACAGAGTTAATAAGA   15282
Pachon15     AAGACATGTGGTCAATATGATCAGTTTGATTAATTAAACATTTAGACAGAGTTAATAAGA   15282
             ************************************************************

Rascon4      AGTTTAAAATTCACCTACTGTATATTTACAATGTGTAAAAAATGTAAAAAAAAAAAAATA   15279
Surface      AGTTTAAAATTCACCTACTGTATATTTACAATGTGTAAAAAATGTAAAAAAAAAAAAATA   15363
Rascon8      AGTTTAAAATTCACCTACTGTATATTTACAATGTGTAAAAAATGTAAAAAAAAAAAAATA   15340
Rascon2      AGTTTAAAATTCACCTACTGTATATTTACAATGTGTAAAAAATGTAAAAAAAAAAAAATA   15345
Rascon15     AGTTTAAAATTCACCTACTGTATATTTACAATGTGTAAAAAATGTAAAAAAAAAAAAATA   15337
Rascon13     AGTTTAAAATTCACCTACTGTATATTTACAATGTGTAAAAAATGTAAAAAAAAAAAAATA   15354
```

```
Rascon6      AGTTTAAAATTCACCTACTGTATATTTACAATGTGTAAAAAATGTAAAAAAAAAAAAAATA      15360
Pachon14     AGTTTAAAATTCACCTACTGTATATTTACAATGTGTAAAAAATGTAAAAAAAAAAAAAATA      15335
Pachon9      AGTTTAAAATTCACCTACTGTATATTTACAATGTGTAAAAAATGTAAAAAAAAAAAAAATA      15344
Pachon17     AGTTTAAAATTCACCTACTGTATATTTACAATGTGTAAAAAATGTAAAAAAAAAAAAAATA      15342
Pachon12     AGTTTAAAATTCACCTACTGTATATTTACAATGTGTAAAAAATGTAAAAAAAAAAAAAATA      15324
Pachon11     AGTTTAAAATTCACCTACTGTATATTTACAATGTGTAAAAAATGTAAAAAAAAAAAAAATA      15341
Pachon7      AGTTTAAAATTCACCTACTGTATATTTACAATGTGTAAAAAATGTAAAAAAAAAAAAAATA      15342
Pachon3      AGTTTAAAATTCACCTACTGTATATTTACAATGTGTAAAAAATGTAAAAAAAAAAAAAATA      15342
Pachon8      AGTTTAAAATTCACCTACTGTATATTTACAATGTGTAAAAAATGTAAAAAAAAAAAAAATA      15342
Pachon15     AGTTTAAAATTCACCTACTGTATATTTACAATGTGTAAAAAATGTAAAAAAAAAAAAAATA      15342
             ************************************************************

Rascon4      AATACATATATATATATATATATATTATATATATATAATATATATATATATATATATATATA      15339 Grp
Surface      AATACATATATATATATATATATATTATATATATATAATATATATATATATATATATATATA      15423 Intron1
Rascon8      AATACATATATATATATATATATATTATATATATATAATATATATATATATATATATATATA      15400
Rascon2      AATACATATATATATATATATATATTATATATATATAATATATATATATATATATATATATA      15405
Rascon15     AATACATATATATATATATATATATTATATATATATAATATATATATATATATATATATATA      15397
Rascon13     AATACATATATATATATATATATATTATATATATATAATATATATATATATATATATATATA      15414
Rascon6      AATACATATATATATATATATATATTATATATATATAATATATATATATATATATATATATA      15420
Pachon14     AATACATATATATATATATATATATTATATATATATAATATATATATATATATATATATATA      15395
Pachon9      AATACATATATATATATATATATATTATATATATATAATATATATATATATATATATATATA      15404
Pachon17     AATACATATATATATATATATATATTATATATATATAATATATATATATATATATATATATA      15402
Pachon12     AATACATATATATATATATATATATTATATATATATAATATATATATATATATATATATATA      15384
Pachon11     AATACATATATATATATATATATATTATATATATATAATATATATATATATATATATATATA      15401
Pachon7      AATACATATATATATATATATATATTATATATATATAATATATATATATATATATATATATA      15402
Pachon3      AATACATATATATATATATATATATTATATATATATAATATATATATATATATATATATATA      15402
Pachon8      AATACATATATATATATATATATATTATATATATATAATATATATATATATATATATATATA      15402
Pachon15     AATACATATATATATATATATATATTATATATATATAATATATATATATATATATATATATA      15402
             ************************************************************

Rascon4      TATATATATATTATATATATATATATATCCACAACAGTTAGAATGGTGTGTAGGTATATAGA      15399
Surface      TATATATATATTATATATATATATATATCCACAACAGTTAGAATGGTGTGTAGGTATGTAGA      15483
Rascon8      TATATATATATTATATATATATATATATCCACAACAGTTAGAATGGTGTGTAGGTATGTAGA      15460
Rascon2      TATATATATA-TATATATATATATATATCCACAACAGTTAGAATGGTGTGTAGGTATGTAGA      15464
Rascon15     TATATATATATTATATATATATATATATCCACAACAGTTAGAATGGTGTGTAGGTATGTAGA      15457
Rascon13     TATATATATATTATATATATATATATATCCACAACAGTTAGAATGGTGTGTAGGTATGTAGA      15474
Rascon6      TATATATATATTATATATATATATATATCCACAACAGTTAGAATGGTGTGTAGGTATGTAGA      15480
Pachon14     TATATATATATTATATATATATATATATCCACAACAGTTAGAATGGTGTGTAGGTATGTAGA      15455
Pachon9      TATATATATATTATATATATATATATATCCACAACAGTTAGAATGGTGTGTAGGTATGTAGA      15464
Pachon17     TATATATATATTATATATATATATATATCCACAACAGTTAGAATGGTGTGTAGGTATGTAGA      15462
Pachon12     TATATATATATTATATATATATATATATCCACAACAGTTAGAATGGTGTGTAGGTATGTAGA      15444
Pachon11     TATATATATATTATATATATATATATATCCACAACAGTTAGAATGGTGTGTAGGTATGTAGA      15461
Pachon7      TATATATATATTATATATATATATATATCCACAACAGTTAGAATGGTGTGTAGGTATGTAGA      15462
Pachon3      TATATATATATTATATATATATATATATCCACAACAGTTAGAATGGTGTGTAGGTATGTAGA      15462
Pachon8      TATATATATATTATATATATATATATATCCACAACAGTTAGAATGGTGTGTAGGTATGTAGA      15462
Pachon15     TATATATATATTATATATATATATATATCCACAACAGTTAGAATGGTGTGTAGGTATGTAGA      15462
             ********** ******************************************** ****

Rascon4      AACCTTCTCACTTAATG--TTTTTTTTTTCTCTATTTATATTATTTTCTACATTGTAAAT      15457
Surface      AACCTTCTCACTTAATGTTTTTTTTTTTTCTCTATTTATATTATTTTCTACATTGTAAAT      15543
Rascon8      AACCTTCTCACTTAATGTTTTTTTTTTTTCTCTATTTATATAATTTTCTACATTGTAAAT      15520
Rascon2      AACCTTCTCACTTAATG--TTTTTTTTTTCTCTATTTATATTATTTTCTACATTGTAAAT      15522
Rascon15     AACCTTCTCACTTAATGTTTTTTTTTTTTCTCTATTTATATTATTTTCTACATTGTAAAT      15517
Rascon13     AACCTTCTCACTTAATG---TTTTTTTTTCTCTATTTATATAATTTTCTACATTGTAAAT      15531
Rascon6      AACCTTCTCACTTAATGTTTTTTTTTTTTCTCTATTTATATTATTTTCTACATTGTAAAT      15540
Pachon14     AACCTTCTCACTTAATGTTTTTTTTTTTTCTCTATTTATATTATTTTCTACATTGTAAAT      15515
Pachon9      AACCTTCTCACTTAATGTTTTTTTTTTTTCTCTATTTATATTATTTTCTACATTGTAAAT      15524
Pachon17     AACCTTCTCACTTAATGTTTTTTTTTTTTCTCTATTTATATTATTTTCTACATTGTAAAT      15522
Pachon12     AACCTTCTCACTTAATGTTTTTTTTTTTTCTCTATTTATATTATTTTCTACATTGTAAAT      15504
Pachon11     AACCTTCTCACTTAATGTTTTTTTTTTTTCTCTATTTATATTATTTTCTACATTGTAAAT      15521
Pachon7      AACCTTCTCACTTAATGTTTTTTTTTTTTCTCTATTTATATTATTTTCTACATTGTAAAT      15522
Pachon3      AACCTTCTCACTTAATGTTTTTTTTTTTTCTCTATTTATATTATTTTCTACATTGTAAAT      15522
Pachon8      AACCTTCTCACTTAATGTTTTTTTTTTTTCTCTATTTATATTATTTTCTACATTGTAAAT      15522
Pachon15     AACCTTCTCACTTAATGTTTTTTTTTTTTCTCTATTTATATTATTTTCTACATTGTAAAT      15522
             *****************   ********************* ******************

Rascon4      TAATACTGGAGACATCAAAACTATGAAGGAATGCAATT      15495
Surface      TGATACTGGAGACATCAAAACTATGAAGGAATGCAATT      15581
Rascon8      TAATACTGGAGACATCAAAACTATGAAGGAATGCAATT      15558
Rascon2      TAATACTGGAGACATCAAAACTATGAAGGAATGCAATT      15560
Rascon15     TGATACTGGAGACATCAAAACTATGAAGGAATGCAATT      15555
Rascon13     TGATACTGGAGACATCAAAACTATGAAGGAATGCAATT      15569
Rascon6      TGATACTGGAGACATCAAAACTATGAAGGAATGCAATT      15578
Pachon14     TGATACTGGAGACATCAAAACTATGAAGGAATGCAATT      15553
Pachon9      TGATACTGGAGACATCAAAACTATGAAGGAATGCAATT      15562
Pachon17     TGATACTGGAGACATCAAAACTATGAAGGAATGCAATT      15560
Pachon12     TGATACTGGAGACATCAAAACTATGAAGGAATGCAATT      15542
```

```
Pachon11      TGATACTGGAGACATCAAAACTATGAAGGAATGCAATT       15559
Pachon7       TGATACTGGAGACATCAAAACTATGAAGGAATGCAATT       15560
Pachon3       TGATACTGGAGACATCAAAACTATGAAGGAATGCAATT       15560
Pachon8       TGATACTGGAGACATCAAAACTATGAAGGAATGCAATT       15560
Pachon15      TGATACTGGAGACATCAAAACTATGAAGGAATGCAATT       15560
              * ************************************
```
